# Genetic history of Rus’

**DOI:** 10.64898/2025.12.30.695215

**Authors:** Tatiana V. Andreeva, Svetlana S. Kunizheva, Alexandra B. Malyarchuk, Fedor E. Gusev, Tatiana V. Ustkachkintseva, Irina Yu. Adrianova, Gleb S. Dotsenko, Anna D. Soshkina, Irina L. Kuznetsova, Natalia A. Dudko, Maria Yu. Plotnikova, Elizaveta V. Rozhdestvenskikh, Andrey D. Manakhov, Asya V. Engovatova, Vladimir V. Sedov, Natalia A. Birkina, Olga V. Zelentsova, Ekaterina O. Chelogaeva, Anna M. Krasnikova, Igor Y. Strikalov, Anna V. Rasskazova, Alexander S. Syrovatko, Pavel D. Manakhov, Maria V. Dobrovolskaya, Alexandra P. Buzhilova, Nikolay A. Makarov, Evgeny I. Rogaev

**Affiliations:** Center for Genetics and Life Science, Sirius University of Science and Technology; Sochi, Russia, 354340; Laboratory of Evolutionary Genomics, Department of Genomics and Human Genetics, Vavilov Institute of General Genetics, Russian Academy of Sciences; Moscow, Russia, 119333; Faculty of Biology, Centre for Genetics and Genetic Technologies, Lomonosov Moscow State University; Moscow, Russia, 119192; Institute of Archaeology, Russian Academy of Sciences; Moscow, Russia, 117292; State Historical Museum; Moscow, Russia, 109012; Institute of Ethnology and Anthropology, Russian Academy of Sciences; Moscow, Russia, 119334; Institute of Geography, Russian Academy of Sciences; Moscow, Russia, 119017; Research Institute and Museum of Anthropology, Lomonosov Moscow State University; Moscow, Russia, 125009; Department of Psychiatry, Umass Chan Medical School; Shrewsbury, MA, USA, 01545

## Abstract

The foundation of the first ancient Rus’ state occurred as a result of the consolidation of diverse communities inhabiting Eastern Europe during the second half of the first millennium CE. Historical sources imply that these communities mostly include East Slavs, whose settlement across a vast territory led to the emergence of the East Slavic/Rus’ culture within the Rus’ state. We generated genomic data for 200 medieval individuals from different locations to elucidate the origin and genetic structure of the Rus’ population during the early stages of the state formation. Our findings reveal a genetic continuum predominantly shaped by two key genetic groups: a broad Slavic-related continuity of different genetic subclusters of Rus’ occupying the enormous European Plain area, and a Fenno-Ugrian (Uralic)-related component in the Northern Rus’ region. Importantly, both groups have a shared genetic substrate inherited from preceding ancient Baltic region populations. To scale Scandinavian ("Viking") heritage, we traced minor Scandinavian genetic lineages that did not make up the dominating genetic stratum of the early Rus’ state. Our study presents the first comprehensive genomic image of the medieval Rus’, highlighting the intricate cultural and genetic interactions between Slavic, Fenno-Ugrian, and other groups that formed the first Rus’ state affecting Europe’s history.

## Introduction

The Rus’ (ancient Rus’) is known as an immense medieval state which unified vast territories from Lake Ladoga to the Middle Dnieper with variegated ethnic landscape and high population mobility^1^. According to traditional accounts the state was founded in 862, when Varangian prince Rurik became the ruler of the Novgorod and the founder of the dynasty that later ruled Rus’ principalities. Slavic expansion to the northern and northeastern territory of Eastern Europe was one of the fundamental factors that contributed to the formation of the Rus’ state and influenced its territorial boundaries^1^. *The Tale of Bygone years* (*Primary Chronicle*)^2,3^, the principal source on the history of ancient Rus’, depict the East Europe territories inhabited by Slavic ethnic units, commonly referred to as “tribes”, each with distinct names (Polyanians, Drevlians, Volynians, Severians, Radimichians, Vyatichians, Krivichians, Ilmen Slovenes, Dregovichians, Polochans, Ulichians, Dulebes, Tivercians, and White Croatians), as well as non-Slavic entities, or "inii iazytse" (other languages). The chronicler identifies the Merians, Chud’, Ves’, and Muroma as the initial inhabitants of the northern and northeastern regions of the Rus’^1,4–7^. Numerous archaeological data reflect consolidation of Rus’ in the 10th-11th centuries CE as the formation of the Rus’ archaeological culture and its spread over extensive territories with some regional differences^8,9^, which was accompanied by the spreading of the Rus’ language (ancient Rus’ language) and the Cyrillic alphabet^10^.

Currently, there is no consensus among historians and archaeologists about the origin and degree of community of the Slavic ethnic units in the Rus’ area during the late first and early second millennium CE^5,7,11,12^. Their interaction with Baltic and Fenno-Ugrian (Uralic) groups inhabiting the western, northern, and eastern territory of Eastern Europe, along with nomadic Turkic-speaking southern groups, is poorly documented in the written sources and likely exhibited variability in its forms and dynamics across different regions^13–17^. The consolidation of the Rus’ state was driven by the northeastern Slavic expansion during the 10th-12th centuries CE into regions historically inhabited by Fenno-Ugrian (Uralic) ethnic communities^16,17^. A critical region for studying the Slavic-Fenno-Ugrian interaction is the area around Beloozero, inhabited, according to the *Primary Chronicle*, by medieval Ves’ people. The Beloozero region and its surrounding territory in present-day Vologda and Arkhangelsk regions thus represent a unique context to genetically trace the interaction between two distinct linguistic and cultural groups that shaped early Rus’. The Slavic settlement resulted in a substantial transformation of the cultural landscape in this area. According to archaeological data, the material culture of all ancient Rus’ urban and rural necropolises in the 10th-13th centuries CE shared cultural uniqueness that spread in vast areas but had specific features in funeral ritual and costume related to local or ethnic identity^8,14^.

The practice of cremation among the early Slavs and ancient Rus’ people persisted until the late 10th century CE, and presents a considerable challenge in obtaining genetic data from samples of the pre-11th century period. Beyond that, except for a few cemeteries^18–20^, there is currently no data on the gene pool of the early medieval population of the whole Rus’ territory, to provide a comprehensive understanding of the genetic makeup and origins of the ancient Rus’ people. In this regard, the genetic characteristics of the people of Rus’ is a key topic for both historians and archaeologists, and here we perform the first comprehensive attempt to characterize the medieval people of Rus’.

Importantly, there are several historiographical names for this Eastern European medieval state that our work is devoted to. Over time, different terms have predominated in the literature. Here, we use the generalised term “Rus’” to delineate the cultural community as well as the geographical area encompassed by our research. Our sample includes anthropological material covering a wide area and a broad time period of the state’s formation and existence, both as a whole state and as several distant principalities. In this study we also adopted the next terminology. The term “East Slavic/Rus’” refers to ancient Rus’ archaeological culture. The term “Slavic-related” refers to populations or individuals exhibiting genetic similarities to contemporary Slavic-speaking populations, while “Fenno-Ugrian (Uralic)” describes those with genetic similarities to present-day Uralic-speaking populations.

## Results

In order to elucidate the origin and history of the ancient Rus’ people, a sample set of individuals from archaeological material dating mainly from the 10th to the 13th centuries CE was combined and analyzed (Fig.1). For the whole genome sequencing we explored bone samples of two hundred individuals from the burials of Rus’/East Slavic type, Fenno-Ugrian type burials, and mixed type burials from the ancient Rus’ area and neighboring regions, tracing back to the formation of the Rus’ state (Fig. 1). An average whole genome coverage for 173 samples ranged from 0.02x to 15.8x (mean genome coverage 1.6x) (Supplementary Tables 1-3). Complete mitochondrial genome sequences and mitochondrial haplogroups were ascribed for all two hundred tested individuals (Supplementary Table 4). The sample set encompass five geographical regions of medieval Rus’: 1) Northern Rus’ (Beloozero and Poonezhye regions) (n=83); 2) North-Western Rus’ Novgorod (n=16); 3) Western Rus’ Dnieper-Dvina region (n=7); 4) Central Rus’ Volga-Oka region (include historical North-Eastern Rus’) (n=54), and 5) Southern Rus’ (n=29) (Fig. 1). To conduct a comprehensive analysis within a broad geographical and temporal context, we also collected samples of synchronous neighbouring non-Rus’/non-Slavic groups from six archaeological sites. These include burial sites from the 12th to 14th centuries CE associated with the Fenno-Ugrian Muroma and Mordva, along with Volga Bulgarians and Eastern-Baltic Latgalian culture individuals, which dates to the 8th-10th centuries CE. Our novel genomic data for the samples derived from the northern to southern regions of ancient Rus’ addresses a significant gap in the genetic record of medieval Eastern Europe.

**Fig. 1.**
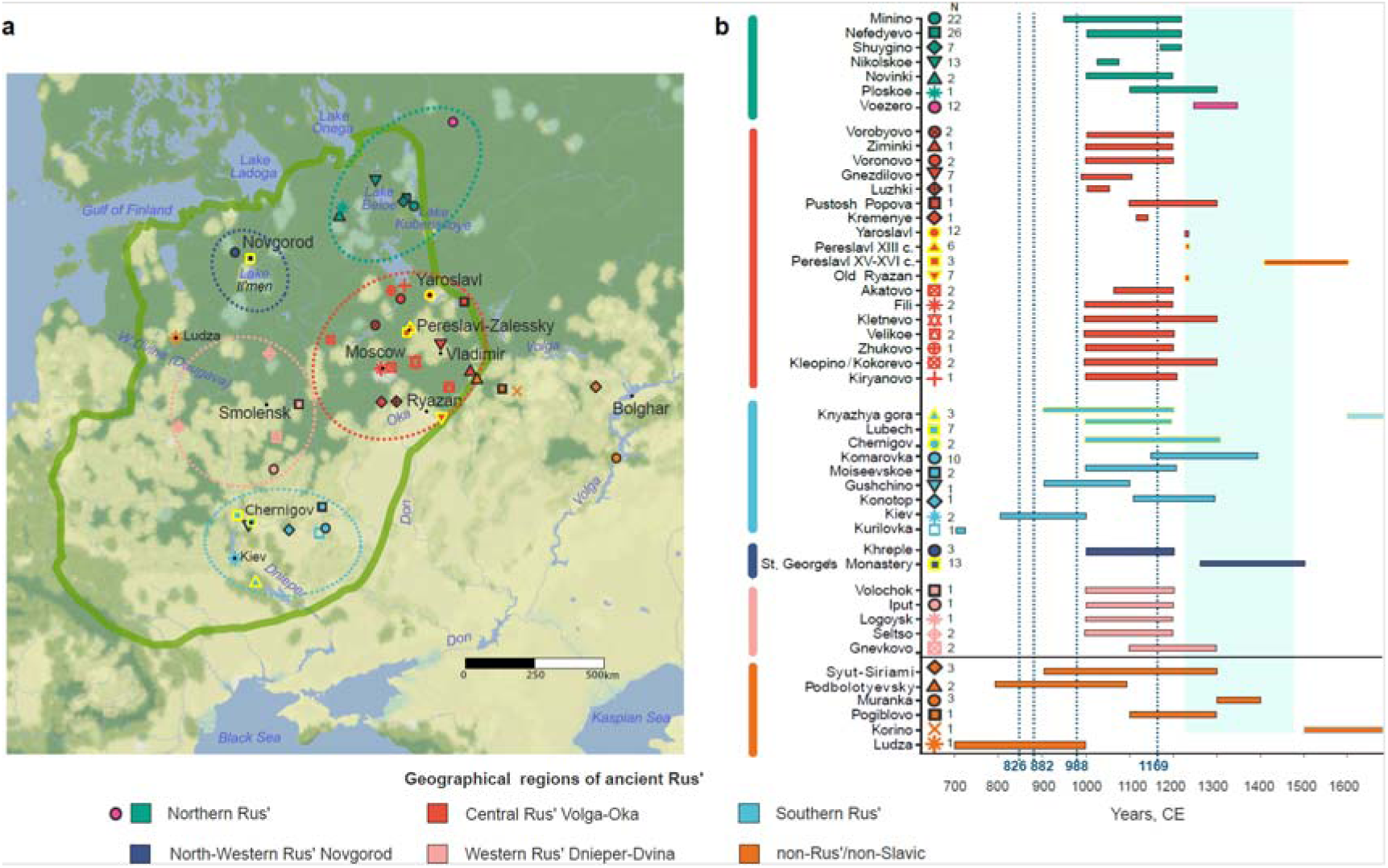
Overview of medieval individuals analysed in this study. (a) Geographical locations of archaeological sites within the ancient Rus’ area and neighbouring territories. The green line corresponds to the approximate border of Rus’ at the end of 11th century - beginning of 12th century CE^92^. (b) Chronological distribution of the 200 ancient genomes sequenced and reported in this study. Dotted lines correspond to the ancient Rus’ establishment (862 CE), the Rus’ capital moving from Novgorod to Kiev (882 CE), Christianization of Rus’ (988 CE), the Rus’ capital moving from Kiev to Vladimir (1169 CE); the blue area corresponds to the period of Mongol invasion of Rus’. The burials in the Northern Rus’ region are marked in green (Beloozero region) and dark pink (Poonezhye region); Central Rus’ Volga-Oka region is in red; the Southern Rus’ region is in light blue; the North-Western Rus’ Novgorod is in dark blue; the Western Rus’ Dnieper-Dvina region is in pink; the burials of the non-Rus’/non-Slavic groups are in orange.

### The genetic structure of Rus’ populations

Principal component analysis (PCA) of medieval Rus’ individuals projected onto present-day Eurasian and European individuals shows that ancient Rus’ samples predominantly align with present-day Europeans with only a few exceptions (Fig. 2, Supplementary Fig. 29). The majority of Rus’ samples are organized in two main genetic clusters (Fig. 3). The largest cluster, which we termed Ancient Major Rus’ genetic cluster (n = 75), lies within genetic variations of present-day Slavic populations. It includes individuals predominantly from Slavic-associated burials from Central Volga-Oka, Western Dnieper-Dvina, North-Western (Novgorod), and Southern Rus’ regions of 11th-13th centuries CE and from burials of late medieval period (14th-17th centuries CE). This Slavic-related genetic cluster also includes several samples from the northern geographic area of ancient Rus’ (Beloozero region): nine samples from the Nikolskoe burial site (attributed to ancient East Slavic/Rus’ culture) and two samples from the burial site of Minino (attributed to a blend of Slavic and Finnish elements). The second genetic cluster, termed Ancient North Rus’ (n = 54), comprises the majority of samples from Northern Rus’ burials. These burials exhibited a blend of Slavic and Finnish cultural elements. Most individuals from the Northern Rus’ burials are distant from present-day Slavic individuals but drawn toward the present-day Vologda and Arkhangelsk populations along with the present-day Fenno-Ugian (Uralic) groups of Finns, Veps, and Karelians representing the so-called Baltic Finnic people^21,22^. Seven individuals are positioned between the two genetic clusters and further named Rus_Middle. Single samples from the Southern Rus’ region fall outside the European genetic variability and are positioned within East Asian or Caucasian populations (Fig. 2).

**Fig. 2.**
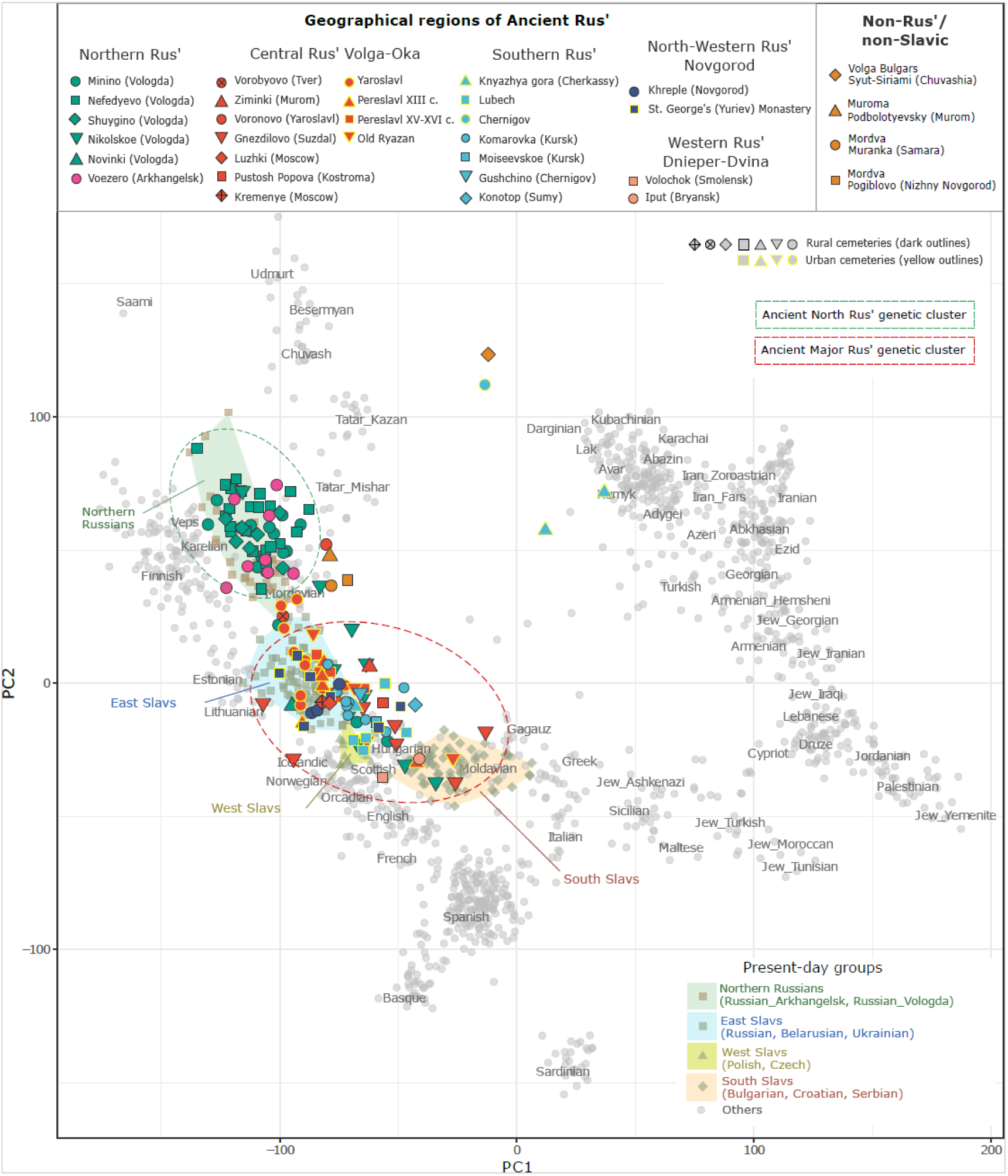
Principal component analysis (PCA). PCA plot visualizing 144 newly reported samples from different geographical regions of ancient Rus’ and neighbouring contemporary groups. Only samples with at least 10,000 SNPs are included in this analysis. Ancient individuals are projected on present-day Western Eurasian individuals of the Human Origin dataset supplemented by present-day Slavs from the Simons Genome Diversity Project^59^, Finnish from the 1000 Genome Project^60^ and Serbians^61^ (grey points). This present-day set includes Northern Russians, East, West, and South Slavs (greyish squares, triangles, and diamonds combined by coloured polygons, see legend for Present-day groups). The labels for the ancient individuals correspond to their burial sites, indicating the site name and/or neighbouring town (see also Fig. 1). Ancient individuals colors are as follows (see also legend on the top): green and dark-pink, samples from Northern Rus’ region; red, samples from Central Rus’ region, light blue, samples from Southern Rus’ region, dark-blue, samples from North-Western Rus’ Novgorod; peach, samples from Western Rus’; orange, samples from Volga Bulgars, Mordva, and Muroma burials. Samples from urban cemeteries have yellow outlines, samples from rural cemeteries have dark outlines.

**Fig. 3.**
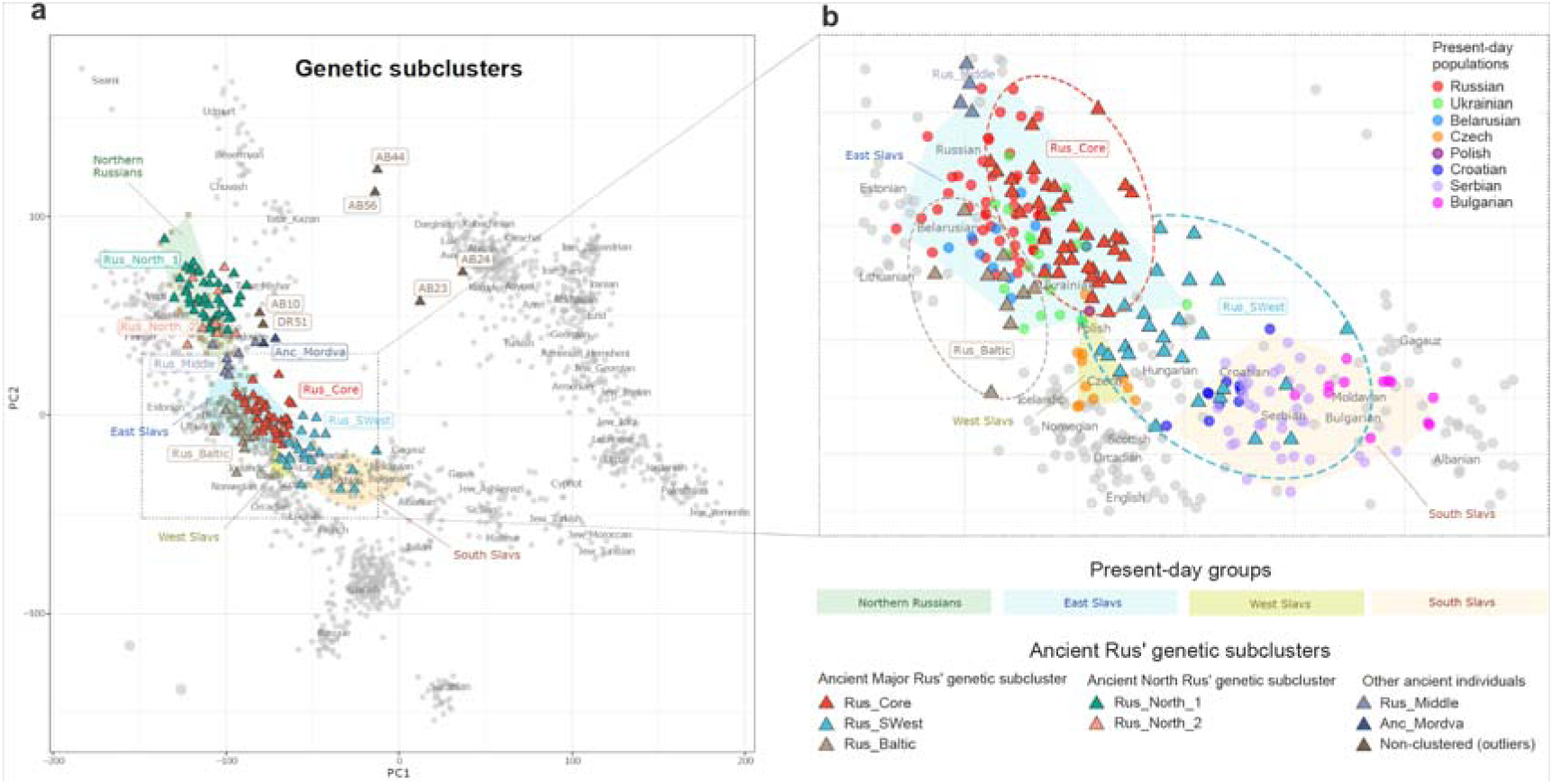
Principal component analysis (PCA) for Ancient Rus’ genetic subclusters. The coloured points for newly reported ancient Rus’ individuals (triangles) correspond to genetic subclusters used in subsequent analysis: Rus_North_1, Rus_North_2, Rus_Middle, Rus_Core, Rus_Baltic, Rus_SWest, Anc_Rus and outliers. (a) PCA plot visualising newly reported ancient Rus’ samples. The labels for the ancient individuals correspond to their genetic subclusters (see also legend). (b) Zoomed in image of the Ancient Major Rus’ genetic subclusters is provided on the right plots for greater detail and to show present-day Slavic populations.

Individuals of non-Rus’/non-Slavic cultures are distributed both within and outside of present-day European genetic variation (Fig. 2, Supplementary Fig. 29). Among them, ancient Mordva and Muroma Fenno-Ugrian subjects fall in the close proximity of Ancient North Rus’ cluster, and the Volga Bulgarian has East Asian affinity. This analysis reveals the structured genetic diversity of ancient Rus’ population and their neighbors, emphasizing both geographic and cultural factors.

Unsupervised ADMIXTURE analyses of these novel genomic data along with genetic profiles of 2304 present-day and ancient subjects at K=7 indicated that ancestry composition of ancient Rus’ individuals is predominantly formed by European western and eastern hunter-gatherers (HG) and European Neolithic farmers (Fig. 4 , Supplementary Fig. 32, 33). The distribution of the HG components are similar in both Ancient North Rus’ and Ancient Major Rus’ genetic clusters, but they differ in European Neolithic farmers and ancient Siberian component proportions. The contribution of European Neolithic farmers is high in the Ancient Major Rus’ samples and progressively decreases from South to North regions. In contrast, the ancient Siberian component, which is maximised in present-day Nganasans, is minor in the Ancient Major Rus’ but notable in the Ancient North Rus’. The East Asian component (related to present-day Hans) predominantly appears among genetic outliers, particularly from Southern Rus’ region, Volga Bulgars, and ancient Mordva samples at varying low frequencies (Fig. 4). These findings indicate the presence of genetic gradients between Slavic-related individuals, who fall into the Ancient Major Rus’ genetic cluster, and Finno-Ugric (Uralic) groups, represented by individuals from the Ancient North Rus’ genetic cluster as well as by ancient Mordva (Anc_Mordva) and Muroma subjects. This genetic difference is consistent with historical and archaeological data^23,24^.

**Fig. 4.**
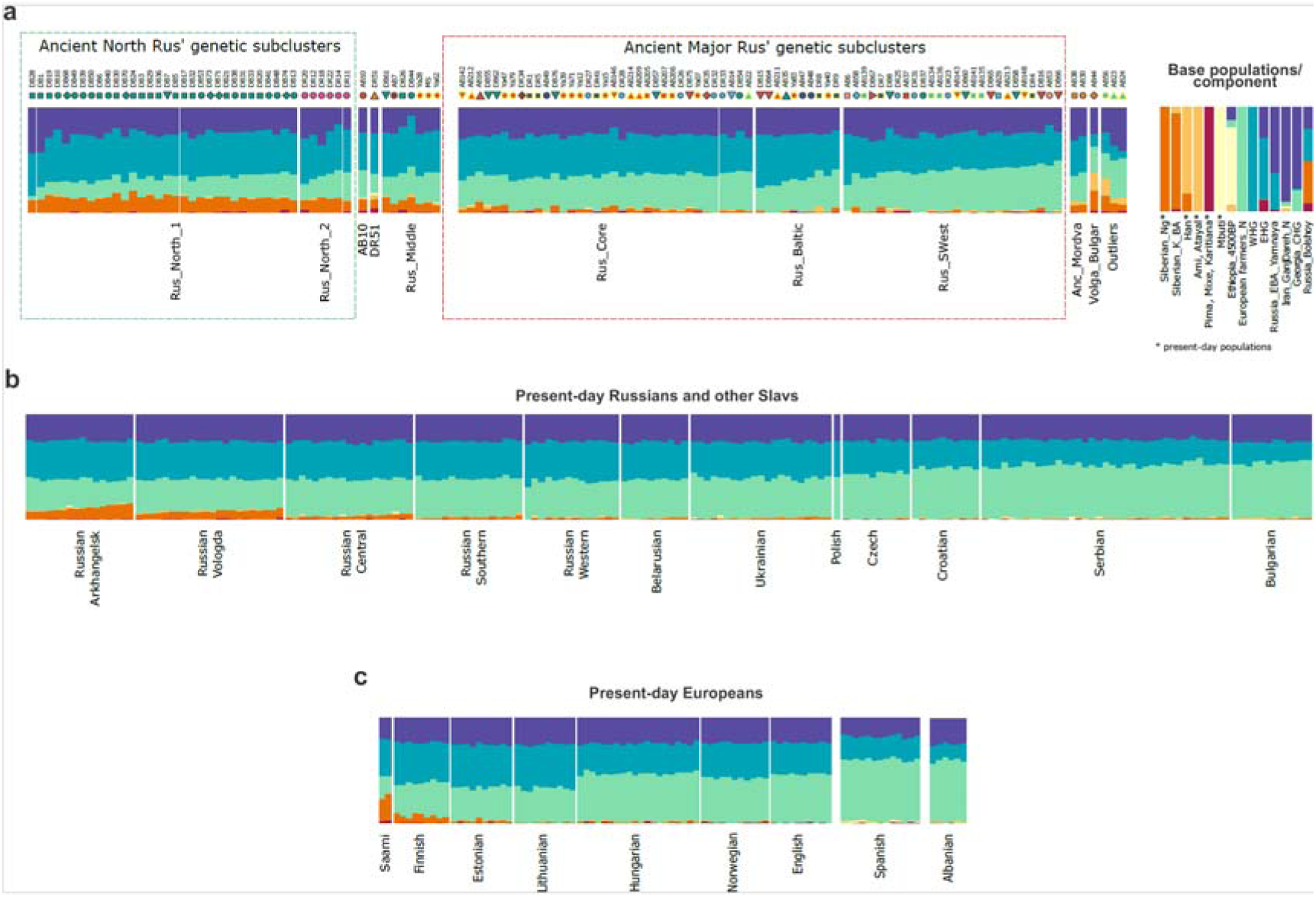
ADMIXTURE results (a) Unsupervised ADMIXTURE results at K = 7 for newly reported ancient Rus’ and neighbouring people, excluding genetically related individuals. Base present-day (marked by an asterisk) and ancient groups (no asterisk) are shown on the top. Labels for ancient individuals correspond to Fig. 1, 2. Siberian_Ng corresponds to present-day Nganasans and Siberian_K_BA corresponds to Russia_Krasnoyarsk BA samples. (b, c) Unsupervised ADMIXTURE results at K = 7 for present-day Russians and Slavic individuals (b), along with some representatives of present-day European populations (c). See also Supplementary Fig. 32 for a complete set of tested ancient and present-day groups and Supplementary Fig. 33 for previously published medieval groups.

Outgroup *f*3-statistics (see Supplementary Informations 2.3) revealed that from the Bronze Age (BA) to Medieval Age (MA) periods the ancient groups from the Baltic region had the highest *f*3 estimates for these two ancient Rus’ genetic clusters (Supplementary Table 8, Supplementary Table 11, Fig. 5, 6). Interestingly, Ancient North Rus’ and Ancient Major Rus’ genetic clusters have no substantial differences in allele sharing rate with all tested BA groups. However, *f*3-analysis of subsequent Iron Age (IA) populations shows that the Ancient North Rus’ cluster had more genetic affinity with IA Finland_Saami and Finland_Levanluchta^22^, than the Ancient Major Rus’ cluster do, highlighting some disparities in their post-BA genetic substratum (Fig. 6).

**Fig. 5.**
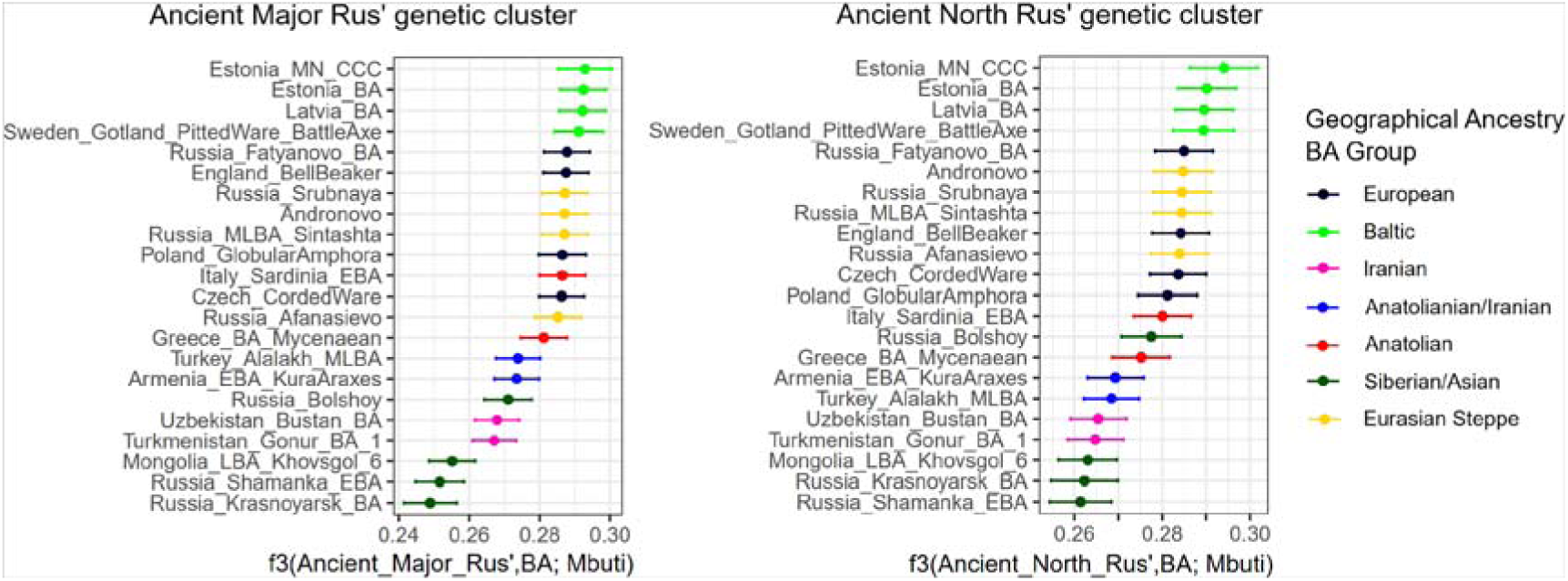
Outgroup f3-statistics show shared genetic drift between two Rus’ genetic clusters and BA groups. Each plot displays the actual values of the f3 statistics (± 3 standard errors) between Rus’ genetic clusters and tested BA groups. Color-coding represents geographical regions where BA groups originated.

**Fig. 6.**
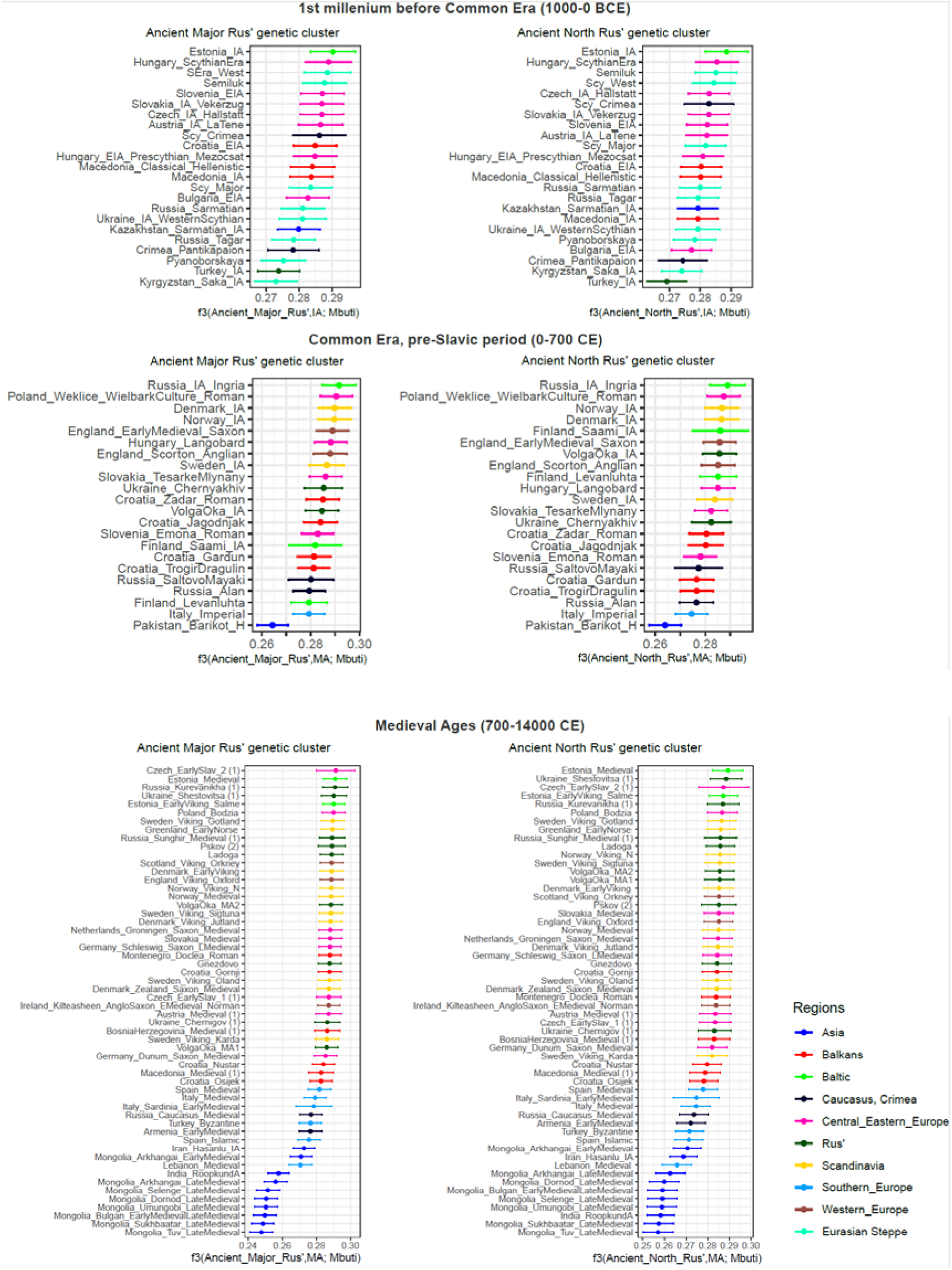
Outgroup f3-statistics displaying shared genetic drift between two Rus’ genetic clusters and BA groups. Each plot displays the actual values of the f3 statistics (± 3 standard errors) between Rus’ genetic clusters and tested groups from the IA to MA, subdivided into three temporal periods: first millennium BCE, pre-Slavic period of CE, and MA. Color-coding represents geographical regions of the tested CE groups.

To trace uniparental heritage among the Rus’ inhabitants, we performed analysis of deep Y-chromosome and mtDNA lineages. Analysis of the Y-chromosome haplogroups showed differences in the distribution of paternal lineages between two ancient Rus’ genetic clusters (Supplementary Informations S2.7). The predominant Y-chromosome lineages identified among the individuals of the Ancient Major Rus’ genetic cluster belong to haplogroup R1a (R1a-Z282), representing 63% of the male samples, and are now prevalent among present-day Eastern European and Slavic populations (Supplementary Fig. 55). Other haplogroups, including E1b, I2a, N1a, R1b, and I1a, emerged with much lower frequencies (Supplementary Fig. 61). In contrast, most individuals (81%) of the Ancient North Rus’ genetic cluster belonged to Y-chromosome clade N1a (Supplementary Fig. 61), which is distributed throughout Northern Eurasia and has the greatest frequency among present-day populations of Fennoscandia, the Uralic region, and native Siberian groups^25^. Alongside N1a, Y-chromosome R1a, I1, and I2 lineages were found with low frequency in the genetic Ancient North Rus’ cluster (Supplementary Fig. 61). The predominant paternal lineage found in the Ancient Major Rus’ cluster is R1a-Z282, specified further as the R1a-Z280 and R-PF6155 sub-branches. These subbranches are today most prevalent among present-day East Slavic groups, indicating a direct genetic continuity between the medieval Rus’ population and present-day East Slavs. Comparative analysis of deep Y-chromosome lineages is also described in Supplementary Informations 2.7.2.

The majority of male individuals of the Northern Rus’ community belong to the Y-chromosome haplogroup N-FT68246 (N1a1a1a1a2a1a1a), indicating a strong patrilineal lineage and underscoring the importance of a “founder” father’s lineage in shaping this northern population. Notably, this paternal lineage presumably dates to the first centuries CE (International Society of Genetic Genealogy, http://www.isogg.org). Remarkably, this deeply defined lineage is currently found solely among Russian populations (Supplementary Fig. 62). The second lineage identified in the Northern Rus’ region is N-Z35049 (N1a2b2a1a). Currently, this Y-chromosome haplogroup is also found exclusively among present-day Russians (Supplementary Fig. 62, (http://www.isogg.org).

Ancient Rus’ mitochondrial DNA (mtDNA) belonged to a variety of haplogroups, primarily of Western Eurasian mitochondrial lineages (H, HV, I, J1, K, R1, T, U, W, and X) (Supplementary Table 4, Supplementary Fig. 56). Only seven of the 200 tested individuals from the Rus’ and neighbouring populations had haplogroups M, A, and D, which are common in present-day Eastern Eurasia. It is noteworthy that over half of the subjects in both North and Central Rus’ genetic clusters are represented by haplogroups H and U5 (Supplementary Fig. 56). Overall, these two Rus’ genetic clusters share a similar distribution of common mtDNA haplogroups. This contrasts with notable differences in the distribution of paternal Y-chromosome lineages between them, with N1a predominating among samples of Ancient North Rus’ and R1a among samples of Ancient Major Rus’. The match between deep mtDNA haplotypes and complete mtDNA sequences revealed direct ancestral linkages between certain people through maternal lineages from the same or distinct cultures and epochs (Supplementary Informations 2.7.1, Supplementary Figure 58).

### Genetic heterogeneity of Slavic-related genetic pool of Rus’

Using PCA data supplemented by ADMIXTURE analysis results (Fig. 2, 4), three genetic subclusters (genetic groups) are identified within the Slavic-related Ancient Major Rus’ genetic cluster (Fig. 3). (1) The Rus_Core samples exhibit genetic profiles overlapping with that of present-day East Slavs (Russians, Belarusians, and Ukrainians) while maintaining relative distinction from other present-day European populations. (2) The Rus_Baltic genetic subcluster shifts away from the Rus_Core cluster towards the present-day East Baltic groups. (3) The Rus_SWest genetic subcluster exhibited greater overlap with present-day individuals of Central-Eastern Europe and South Slavs (Balkan region) (Fig. 3, Fig. 4, Supplementary Fig. 30).

Interestingly, we found no substantial geographic-genetic correlations for the Rus’ genetic subclusters. For example, individuals from the geographically close locations and from the same Rus’ towns (Yaroslavl, Old Ryazan, Pereslavl, and Novgorod) are presented in different genetic subclusters. Individuals from military elite (Druzhina) barrows at Gnezdilovo (Suzdal, Volga-Oka region) and Nikolskoe (Beloozero region) are also placed within different subclusters of the Ancient Major Rus’ genetic cluster (Supplementary Fig. 34). Overall, this observation suggests that the geographically separated Rus’ communities (referred to as several medieval Slavic-related “tribes”)^2,3^ are not genetically distinct groups. They form one heterogeneous genetic continuity pool, ancestral to present-day East Slavs.

Finally, we specify the possible genetic links of each Ancient Major Rus’ genetic subcluster, stressing their similarities with geographically and temporally suitable populations using PCA. All three genetic subclusters of the Ancient Major Rus’ were positioned mainly in proximity to European IA and MA populations on the PCA plot.

However, subjects from Medieval Slavic burials of Central Europe (Slovakia, Croatia, and Austria)^26,27^ are situated within the Rus_Core subcluster (Supplementary Fig. 30). On the PCA plot, some MA and IA samples originating from the Scandinavian region, as well as from the MA burials attributed to Saxons, Vikings, and Slavs from the territory of present-day Germany, Poland, Croatia, and some individuals from Scandinavian-type burials from the ancient Rus’ area^19,26,28,29^ are also positioned alongside subjects of the Rus_SWest subcluster (Supplementary Fig. 30).

Further, we traced deeper ancestry of each genetic subcluster of the Ancient Major Rus’ group through *f*3- and *f*4-statistics and *qpAdm* analysis with ancient populations from the early BA period up to MA (Supplementary Informations S2.3, S2.4).

First, we tested Ancient Major Rus subclusters with groups of BA periods. We observed a high affinity of all subclusters (Rus_Core, Rus_Baltic and Rus_SWest) with Baltic region populations of the BA period (Fig. 5, 6, Supplementary Table 8, 9, 10, 11, Supplementary Fig. 36, 37). Despite the geographic proximity of the BA Fatyanovo group to Rus’ area, it did not exhibit a significantly higher number of shared alleles with any of the Ancient Major Rus’ genetic subclusters (Supplementary Fig. 39, Supplementary Table 13).

In *qpAdm* models based on BA groups (see details in Supplementary Informations S2.4), all Ancient Major Rus’ subclusters could be modelled as a two- or three-source predominantly of Baltic ancestry component (60-82%, represented by Latvia_BA). In all three subclusters this BA Baltic ancestry component is admixed with European/Anatolian-related component (up to 32%, represented by Greece_BA_Mycenaean) and with minimum percentage (less than 8%) if else of ancient Siberian ancestry (Russia_Krasnoyarsk_BA) component (Supplementary Informations S2.4, Fig. 7).

**Fig. 7.**
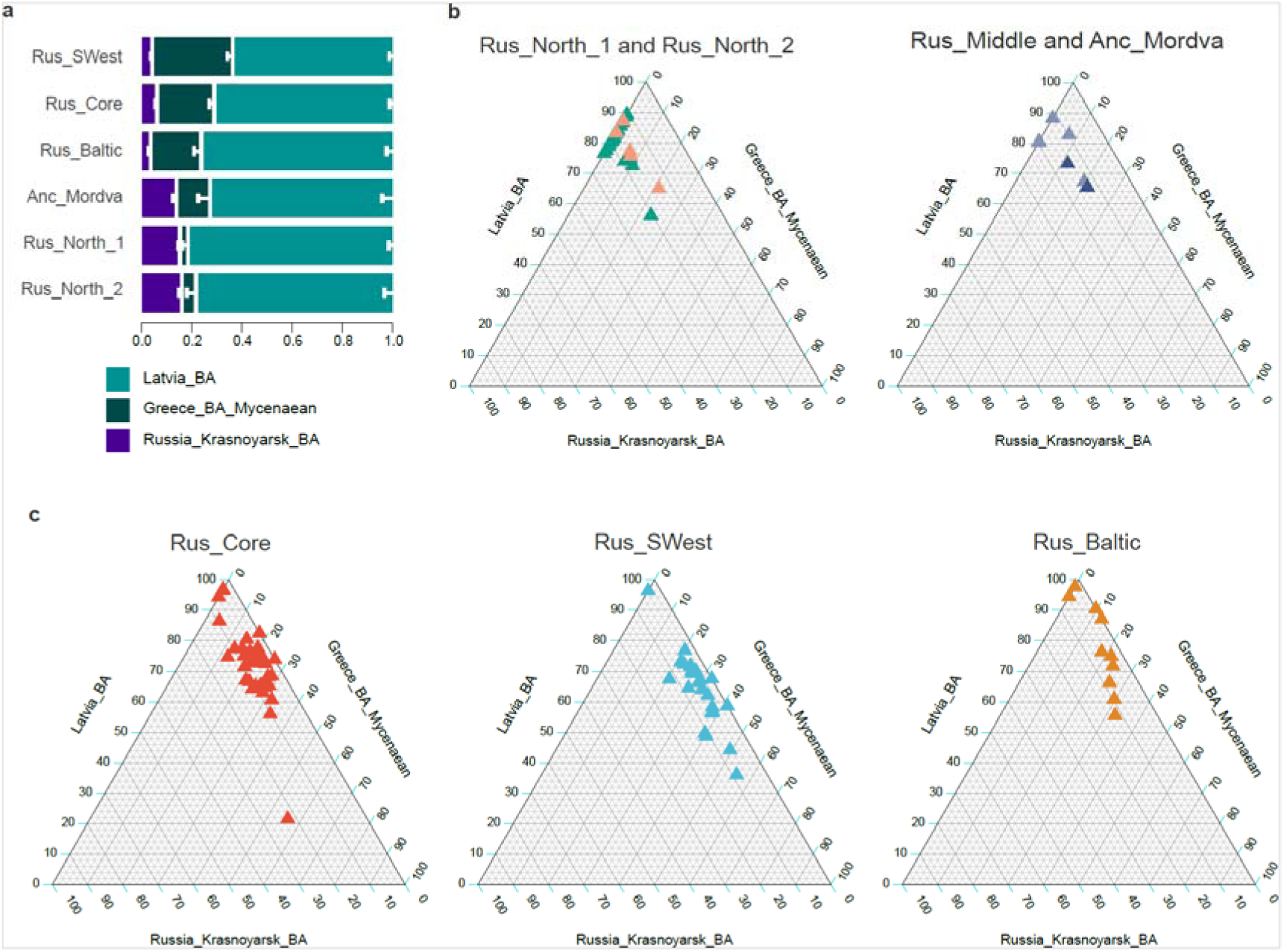
Three-source BA-based qpAdm model elucidates an hypothetical history of admixture among the ancient Rus’ people during the 10th-13th centuries CE. (a) Inferred proximal qpAdm models for the Rus’ genetic subclusters using Latvia_BA, Greece_BA_Myceniana, and Russia_Krasnoyarsk_BA sources. Each bar represents admixture proportions of the BA sources. (b) Ternary plot for a three-way admixture model per individual of Ancient North Rus’ genetic subclusters using Latvia_BA, Greece_BA_Myceniana, and Russia_Krasnoyarsk_BA sources (c) Ternary plot for a three-way admixture model per individual of Ancient Major Rus’ genetic subclusters using Latvia_BA, Greece_BA_Myceniana, and Russia_Krasnoyarsk_BA sources. See also Supplementary Informations S2.4 and Supplementary Fig. 49a. The colour and shape of the points correspond to Fig. 3.

However, these genetic subclusters have different proportions of these basic ancestral components according to the *qpAdm* modeling (Fig. 7), which align with the ADMIXTURE analysis (Fig. 4). The BA European/Anatolian-related component proportion is highest in the Rus_SWest subcluster while the BA Baltic component is maximized in the Rus_Baltic subcluster with only a minimal ancient Siberian influence in both these subclusters (Fig. 7).

Next, we compared the Ancient Major Rus subclusters with the post-BA populations. During the post-BA period before CE, all local populations of the forest-steppe and forest zones of Rus’ area practiced cremation, leaving no anthropological material available for genetic analysis. Therefore, we used for further analysis the available samples dated from the beginning of CE and later. Firstly, outgroup *f*3-statistics in the form *f*3(Ancient Rus’; Test_Pop; Mbuti) revealed that all three Ancient Major Rus’ genetic subclusters show the greatest genetic affinity with the Baltic region population (Russia_IA_Ingria dated to 1st-2nd centuries CE) compared to other European groups of the same chronological period from the beginning of CE (Fig. 8, Supplementary Fig. 38). Next, the *f*4-statistics indicated that all three genetic subclusters of Ancient Major Rus’ are cladal with ancient populations (from the beginning of CE) of Eastern-European ancestry, not Northwestern or Southern European ones (Supplementary Fig. 42, Supplementary Informations S2.3.1).

**Fig. 8.**
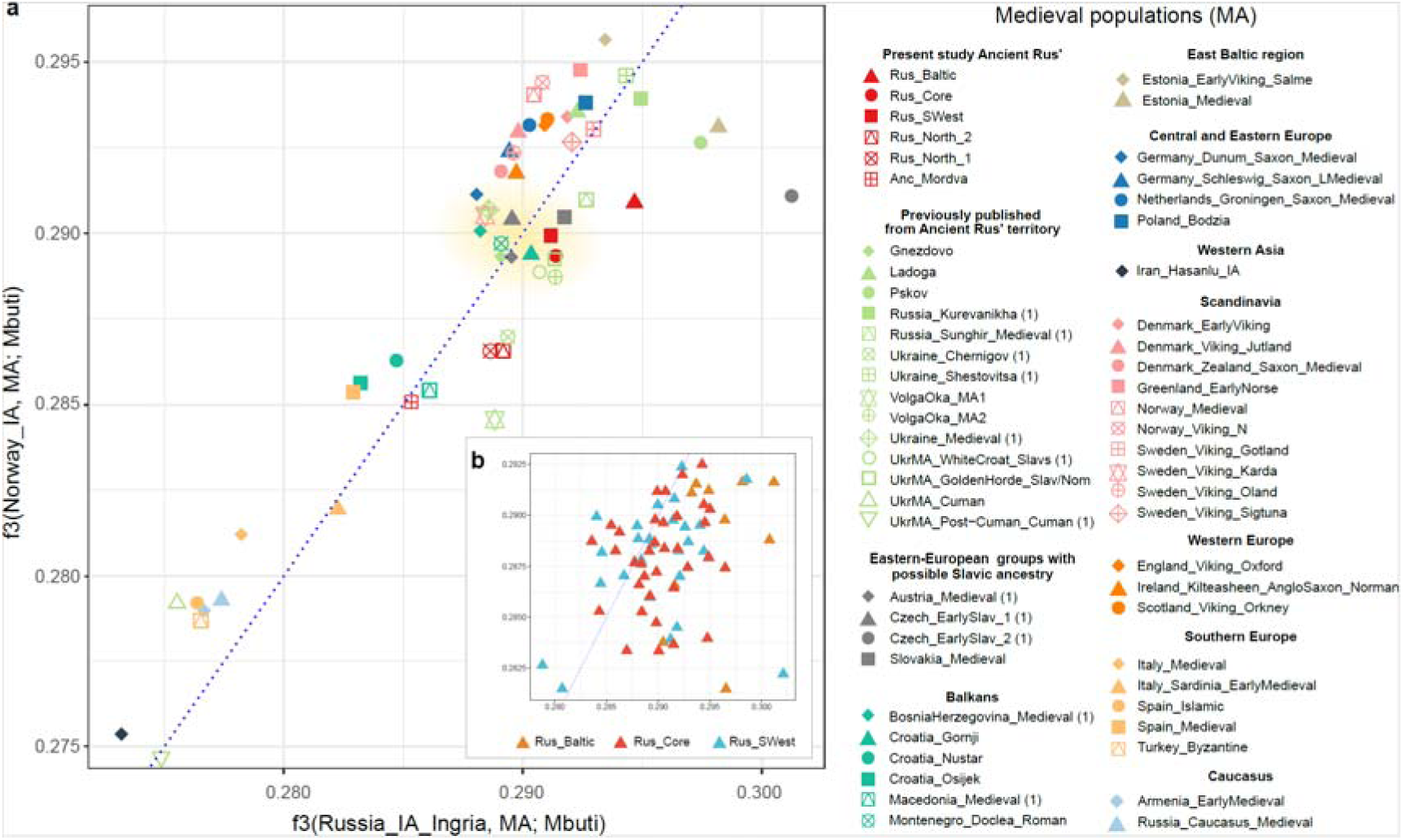
Genetic affinity of ancient Rus’ individuals. Outgroup *f*3-statistic results of form *f*3(IA, MA; Mbuti) plotted against each other: Baltic region IA group (Russia_IA_Ingria, X axis) and Scandinavian IA group (Norway_IA, Y axis). The colour and shape of the points are provided in the figure legends and correspond to Medieval populations (a) and individuals of Ancient Major Rus’ genetic clusters (b). The number 1 is shown in parenthesis for a single-sample group. The putative area of the ancient Slavic-related genetic core on the plot is highlighted in yellow (a). See also Supplementary Information S2.3.1 and Supplementary Fig. 38.

Furthermore, the *f*3-statistics results demonstrate connections between the Rus_Core and Rus_SWest subclusters and individuals from Slavic-related burials beyond the territory of ancient Rus’ (Fig. 8). Notably, in confirmation of our PCA results (Supplementary Fig. 30, 35), several individuals from non-Slavic MA burials from Southern Sweden^30^ and Lower Saxony^31^ (9th-11th centuries CE) exhibit a greater genetic affinity with Slavic-related groups than with Scandinavians or other MA European populations (Fig. 8). Presumably, these MA individuals could represent European groups genetically linked to ancient Slavic-related populations within a common Slavic-related genetic core (Fig. 8)

Taken together, the genetic pool of all three Ancient Major Rus’ subclusters is primarily based on the Baltic region ancestral BA substrate exhibiting also ties to some other geographically broader MA groups.

### Genetic structure of Northern Rus’ genetic pool

The samples from medieval Beloozero dated to the 10th-13th centuries СЕ and Poonezhye regions dated to the 13th-14th centuries СЕ (Fig. 1) form separate Ancient North Rus’ genetic cluster on PCA (excluding a few samples that were genetically aligned with the Ancient Major Rus’ cluster) (Fig. 2, 3). These two Ancient North Rus’ groups are further designated as Rus_North_1 (Beloozero region individuals) and Rus_North_2 (Poonezhye region individuals) genetic subclusters. These two genetic subclusters are overlapped on the PCA plot (Fig. 3a) but exhibit some differences in their ancestry composition, with a higher proportion of ancestry from European Neolithic farmers in the Rus_North_2 than in the Rus_North_1 subcluster (Fig. 4a).

In comparative analysis with groups of BA groups, both Rus_North_1 and Rus_North_2, according to the *f*3- and *f*4-statistics, share the highest genetic affinity with Baltic region groups through all BA periods, similar to all Ancient Major Rus’ genetic subclusters (Supplementary Table 9, 13, Supplementary Fig. 36, 37).

By testing two- and three-way *qpAdm* models based on BA groups, we revealed that, according to the best fitted models, both Rus_North_1 and Rus_North_2 genetic subclusters received the majority of their ancestry (approximately 80%) from the late BA Baltic region group (Latvia_BA) (Fig. 7, see details in Supplementary Informations 2.4). Altogether our data demonstrate that both Ancient North Rus’ and Ancient Major Rus’ genetic clusters have genetic affinity to the Baltic ancestry BA populations. However, in contrast to the Ancient Major Rus’ genetic subclusters, which show a visible European Anatolian-related ancestry component, only several individuals from the North_Rus_1 and North_Rus_2 subclusters exhibit Anatolian-related ancestry (Supplementary Table ) (Fig. 7b). Our analysis also reveals that both Ancient Northern Rus’ subclusters harbor a notable proportion of ancient Siberian ancestry (Russia_Krasnoyarsk_BA), 10-23% in individual samples (Fig. 7b) most enriched in present-day Samoyedic-speaking Nganasans^32^. To trace the origin of the ancient Siberian ancestry involved in the formation of ancient Northern Rus’ populations, we conducted *f*4-statistics to test the affinity with ancient Siberian sources (among the previously published ancient genomic data). Our analysis reveals a greater genetic affinity of both Ancient North Rus’ genetic subclusters to ancient BA Siberian groups compared to the geographically proximate BA Bolshoy Oleni Ostrov population (Supplementary Fig. 41). This finding underscores that the influx of Siberian ancestry into the Northern Rus’ population occurred through independent migration waves from Siberia, rather than from Northeastern Europe, represented by Bolshoy Oleni Ostrov, where the earliest presence of the Siberian ancestry in Northeastern Europe has been discovered^33^.

Next, we tested the ancient populations dated from the beginning of CE and later. We observed that only one ancient group, VolgaOka_IA (from Bolshoye Davydovskoye 2 from the Suzdal region)^18^, dated to the 3rd to 4th centuries CE, was positioned on the PCA plot among the samples of the Ancient North Rus’ genetic subclusters (Supplementary Fig. 30). However, *f*4-statistics results showed that among the European populations of the first millennium CE, the VolgaOka_IA group shared a higher number of alleles with Ancient North Rus’ individuals exclusively in tests involving Western European groups. In contrast, the Baltic groups (Russia_IA_Ingria, 1st-3rd centuries CE, and Estonia_Medieval.SG, 12th-16th centuries CE) exhibited significantly higher estimates of shared alleles with the Ancient North Rus’ genetic subclusters when compared to the VolgaOka_IA group (3rd-4th centuries CE) (Supplementary Table 14). These findings provide further evidence that the primary genetic substratum of both the Rus_North_1 and Rus_North_2 genetic subclusters was identical to that of the Ancient Major Rus’ cluster and genetically related to ancient Baltic region groups. However, unlike the Ancient Major Rus’ genetic subclusters, the Ancient North Rus’ subclusters incorporate a substantial proportion of ancient Siberian ancestry.

Next, in Ancient North Rus’ groups, we identified numerous individuals with close biological kinship (see Supplementary Informations S2.8, Supplementary Table 30, 31).

We combined data from autosomal, uniparentally inherited markers (mitochondrial DNA and Y-chromosome lineages), along with archaeological dates of burials, age-at-death, and genetic sex of tested individuals to establish small initial pedigrees (Supplementary Fig. 64). Then, we incrementally expanded them, resulting ultimately in genealogy reconstruction of the large pedigree of the Northern Rus’ community, in total, spanning seven generations and comprising 48 individuals from the Nikolskoe, Minino, Nefedyevo, and Shuygino burial sites (Fig. 9a). The majority of male individuals of this Northern Rus’ community belong to the Y-chromosome haplogroup N-FT68246 (N1a1a1a1a2a1a1a), indicating a strong patrilineal lineage and underscoring the importance of a “founder” father’s lineage in shaping this northern population. Notably, this paternal lineage presumably dates to the first centuries CE (International Society of Genetic Genealogy, http://www.isogg.org). Remarkably, this deeply defined lineage is currently found solely among Russian populations (Supplementary Fig. 62).

**Fig. 9.**
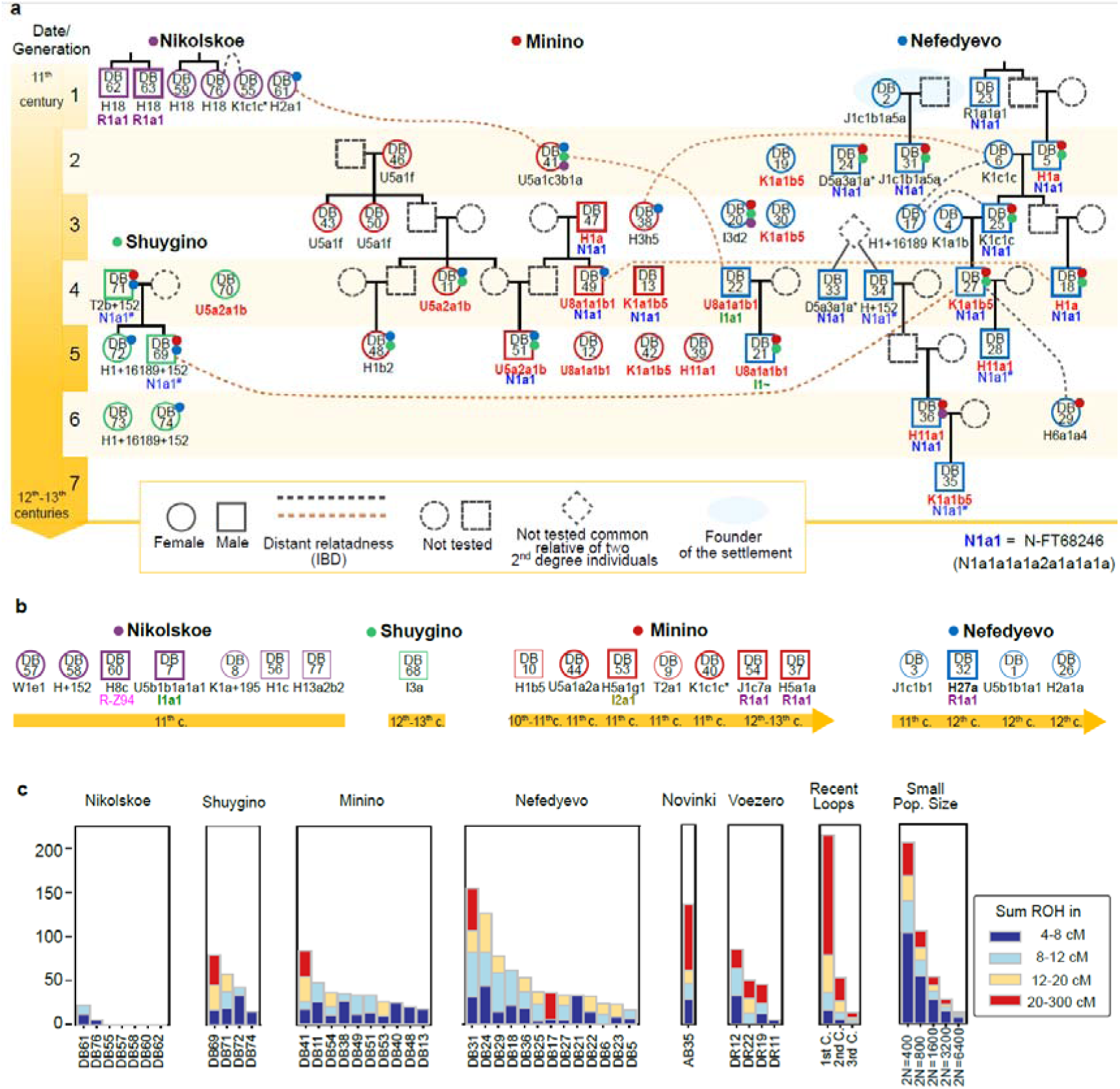
Genealogical network and runs of homozygosity (ROH) for ancient individuals from the Northern Rus’ region. The reconstructed pedigrees for the Nikolskoe (purple), Minino (red), Nefedyevo (blue), and Shuygino (green) groups, extending to the 7th generation. The reconstructions are based on a comprehensive analysis of results from READv2, identity by descent (IBD), and mtDNA assessments, as well as biological age and sample dating. Black dashed lines indicate distant relationships (4th to 6th degree) within the same burial site, while brown dashed lines illustrate an example of the distant relationships between subjects from different burials, with other inter-burial relationships not shown to facilitate readability. The colored dots on the samples indicate the individuals from whom the burial site has been identified as genetic relatives (4th to 6th degree) for the marked sample: red, Minino; blue, Nefedyevo; green - Shuygino; plum - Nikolskoe. Y-chromosome and mtDNA haplogroups are provided for each sample. mtDNA haplogroups of individuals who share identical mtDNA sequences are indicated in red. The symbol * denotes non-identical mtDNA sequences of the sample mtDNA haplogroup. The symbol # denotes low-covered samples, and samples whose Y-chromosome haplogroup was ascertained based on the reconstructed pedigree. Although the kinship between DB35 and DB36 was predicted to be parent/offspring, it was impossible to identify which of them is a father and which is a son. For complete data, see Supplementary Informations S2.6, S2.8 and Supplementary Tables 28, 30, 31. (a) mtDNA and Y chromosome haplogroups for Northern Rus’ individuals whose family ties have not been found, along with a chronology of their burial site. Thin lines indicate low-coverage data samples. (b) Distributions of inferred runs of homozygosity longer than 4 cM (ROHs) for Northern Rus’ individuals. Each sample is represented by stacked vertical bars grouped in four length colour-coded classes: 4–8cM (dark blue), 8–12cM (light blue), 12–20cM (yellow), and more than 20cM (red). The expected ROH length distributions for specific parental relationships and demographic scenarios are depicted on Recent Loops and Small Population Size panels.

Evidence of biological relatedness was also found among the individuals of the Poonezhye population of the Ancient North Rus’ (Rus_North_2), where four pairs of first-degree relatives and several mitochondrial lineages of relatedness were determined among eight out of ten tested individuals (Supplementary Fig. 65). It is noteworthy that two Y-chromosome branches were identified among the tested individuals from Voezero, both of the N1a haplogroup. Two individuals with distant IBD-connections (DR11 and DR12) shared a Y-chromosome lineage similar to those among the Beloozero region individuals (the longest identified haplogroup is N-Z1927 (N1a1a1a1a2a1a1a) (Supplementary Table 4). The second lineage, N-Z35049 (N1a2b2a1a) was identified in the child from the Voezero family group (Supplementary Fig. 62, http://www.isogg.org).

To clarify further the genetic structure of medieval Northern Rus’ communities, we computed runs of homozygosity (ROH). We observed a notable difference in the genetic structure of the Ancient North Rus’ populations compared to those of the Ancient Major Rus’ groups (Supplementary Fig. 51). We identified a pattern of runs of homozygosity that suggests increased background relatedness attributable to the small size of the Northern Rus’ populations. Furthermore, evidence of direct consanguinity was present in the Northern Rus’ region (Fig. 9c).

### Genetic interactions in a process of formation of the Rus’ state

Our data provide evidence for genetic interactions between different genetic groups within a broad Rus’ area. A group of seven individuals that we designated as Rus_Middle positioned on the PCA plot between the Ancient Major Rus’ and Ancient North Rus’ genetic clusters (Fig. 3a). These individuals originated from different geographic regions of Rus’; they were inhabitants of the Northern Rus’ (Nefedyevo, Minino, and Nikolskoe) and Central Rus’ regions (Yaroslavl and Tver regions). According to the ADMIXTURE analysis (Fig. 4), Rus_Middle individuals demonstrate intermediate genetic profiles between the Ancient Major Rus’ and the Ancient North Rus’ genetic clusters. The significantly negative admixture-*f*3 statistics *f*3(Rus_Middle; Ancient_North_Rus, Ancient_Major_Rus) further supported the classification of the Rus_Middle genetic group as an admixture between the Ancient Major Rus’ and Ancient North Rus’ (Supplementary Table 16). These data suppose genetic admixture occurring between some communities of the Northern and Central Rus’ regions, representing Fenno-Ugrian and Slavic groups, respectively.

Direct interaction between the representatives of the Ancient Major Rus’ and Ancient North Rus’ genetic clusters is also confirmed by shared IBD fragments (Supplementary Table 31, Supplementary Fig. 53). In the Minino burials, characterized by predominant Ancient North Rus’ genetic profiles, we also found individuals with Slavic-related genetic profiles of the Ancient Major Rus’ cluster (DB37 and DB54) (Table 1, Fig. 2). Vice versa, in the Slavic mound at the Nikolskoe burial site characterized by the predominant Ancient Major Rus’ genetic profile, we found an individual of the Ancient North Rus’ genetic profile (DB7) (Supplementary Table 4, Fig. 2). In addition, we revealed a four-five degree relatedness between the member of the Minino’s kin (DB41) and the person from the Nikolskoe burial (DB61) in the Northern Rus’ region. Altogether, these results imply the coexistence and direct interaction of two genetically and culturally distant groups within the Northern Rus’ region.

**Table 1.**
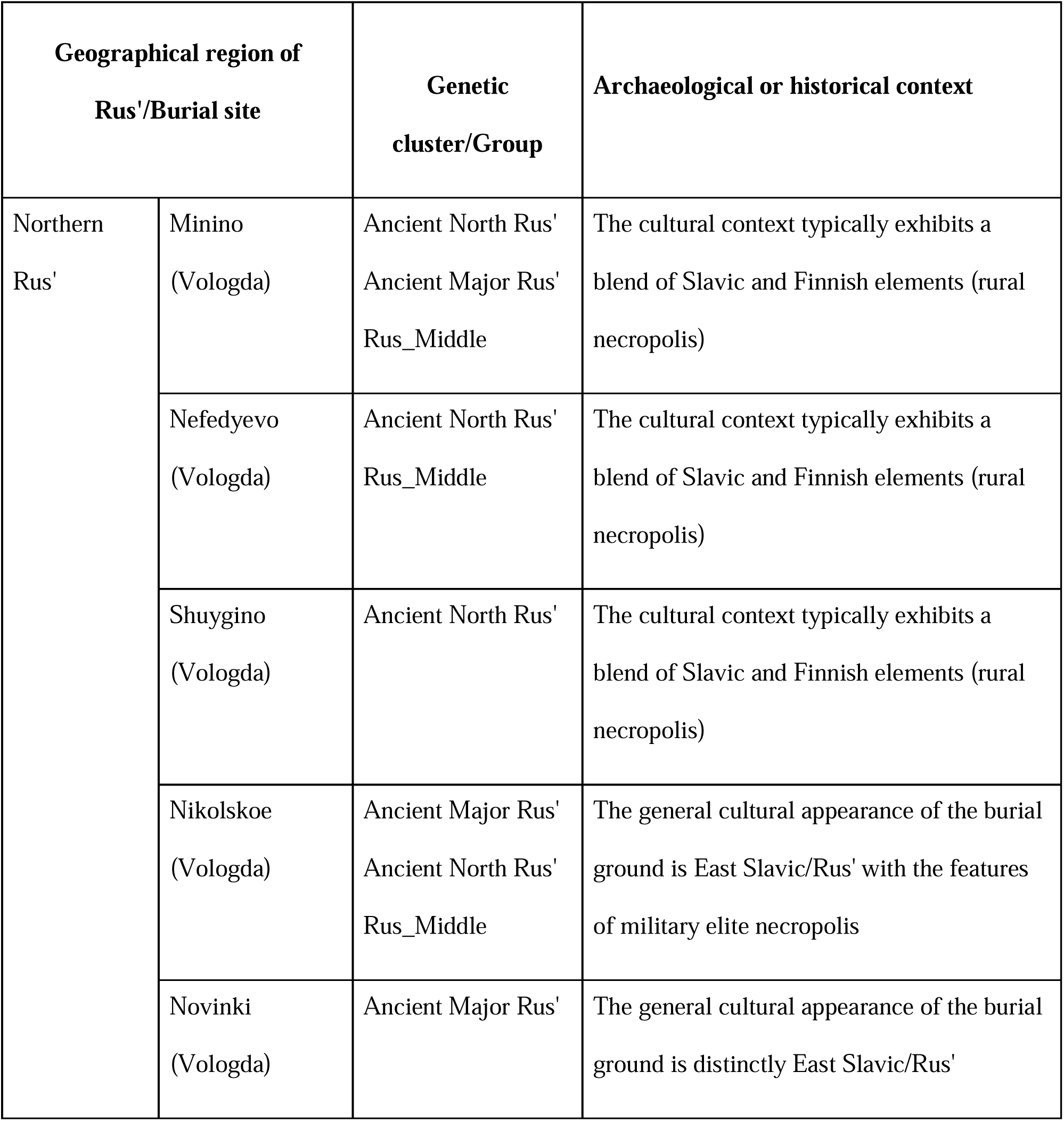
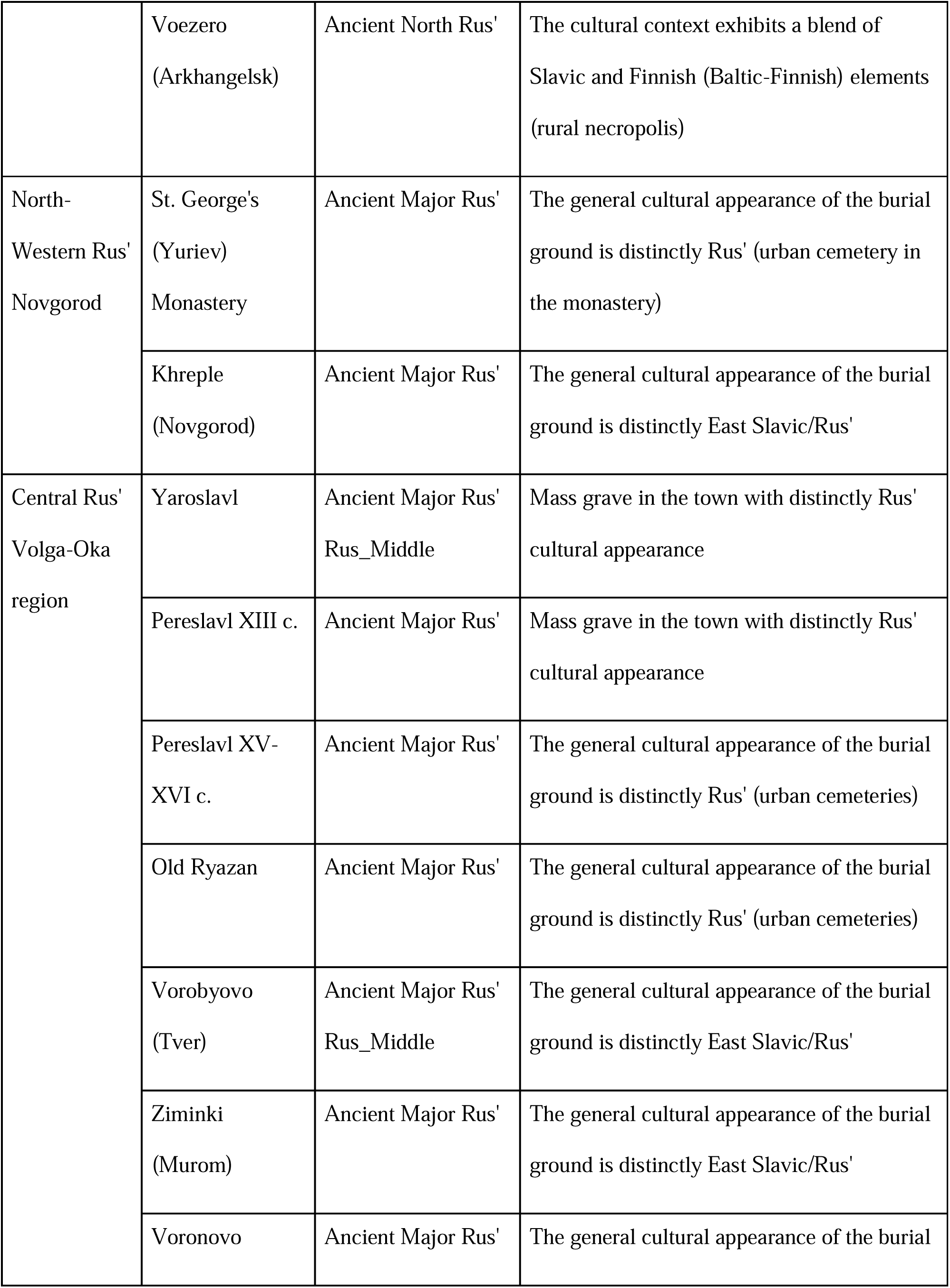
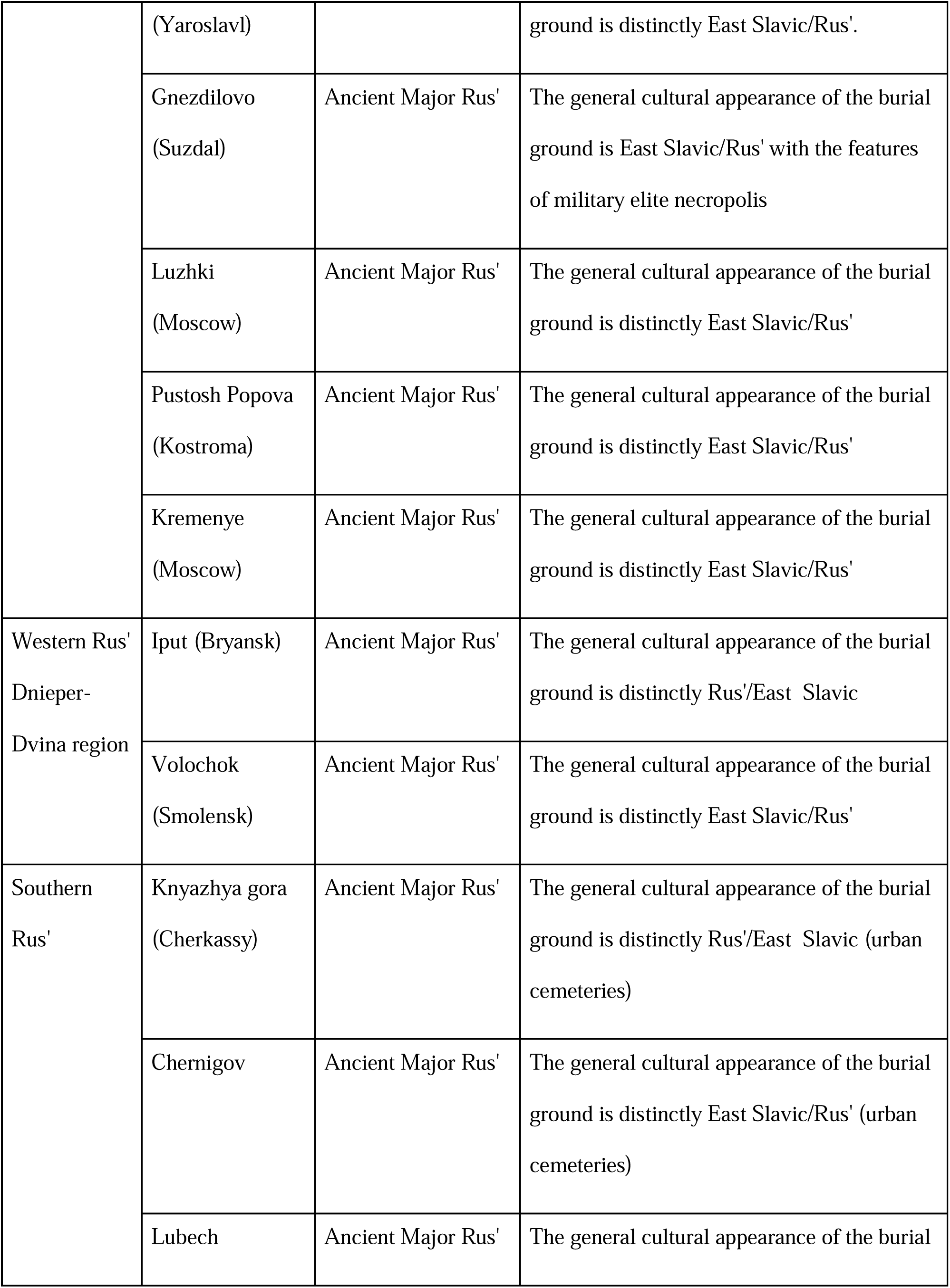
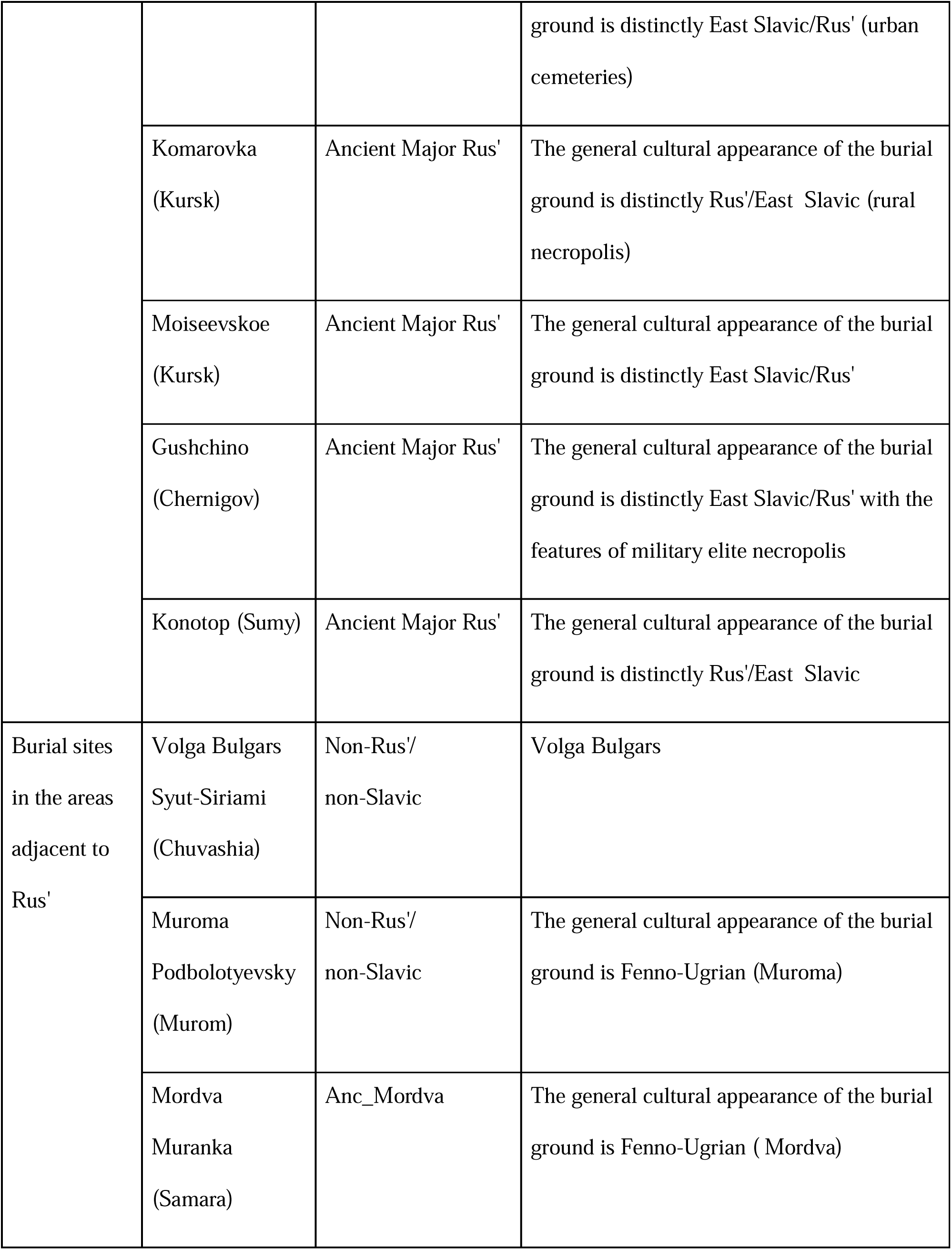
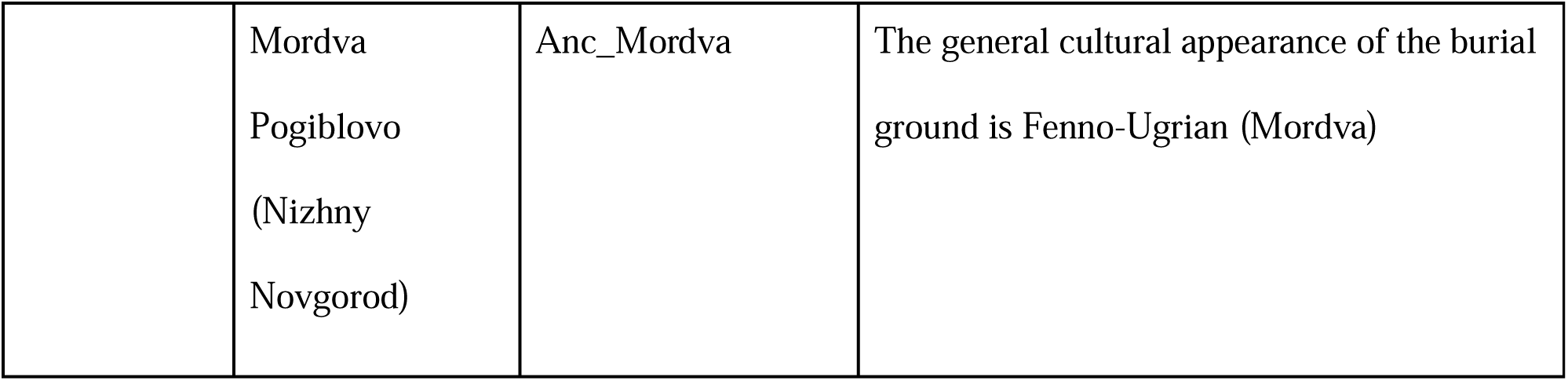
The table displays exclusively the samples from our dataset that were selected and utilised in the population analysis

Next, we tested possible genetic connections between inhabitants of geographically distant Rus’ regions and neighbouring areas. We employed IBD shared fragments analysis, ADMIXTURE, and PCA results (Fig. 2, 3, Supplementary Fig. 54, Supplementary Table 28, Supplementary Informations 2.6) along with uniparental Y-chromosome and mtDNA haplogroup data (Supplementary Table 4) to search for the presumable genetic links and affinity. We found common IBD fragments between some Northern Rus’ inhabitants and individuals from medieval Yaroslavl, Pereslavl, and Old Ryazan (Supplementary Fig. 54). We also identified common IBD fragments between Mordva individual (AB30) from the Samara region and several Northern Rus’ subjects. These individuals represent two different Fenno-Ugrian groups from geographically distant regions. Among individuals from Slavic-related burials, IBD connections were revealed between several individuals from distant Southern Rus’ burials (AB22 and AB29), as well as between two individuals from geographically distant urban cemeteries of Novgorod and Pereslavl (Supplementary Fig. 54, Supplementary Table 28, 29, Supplementary Informations 2.6). These results provide evidence of interactions and long-distance mobility within the Rus’ area.

In addition, in the burials attributed to the East Slavs in the Southern Rus’ towns, we revealed genetic outliers (see Supplementary Informations S2.2.3). Despite their striking genetic differences from the remaining Rus’ area inhabitants, all the genetic outliers were buried according to the East Slavic/Rus’ burial practice, suggesting integration of these individuals into the local Rus’ society.

Finally, we examined previously published groups of European medieval individuals to identify their presumable links to the Rus’ inhabitants. Screening of medieval European samples for shared IBD fragments within our dataset revealed several pairs exhibiting IBD connections between Rus’ and other Europeans described in more detail in Supplementary Informations 2.6, Supplementary Fig. 54. A single shared IBD was found in an early medieval sample from Croatia (I26748)^26^ and the subject from Yaroslavl (Ancient Major Rus’ cluster). We paid special attention to looking for probable Scandinavian genetic contributions to the ancient Rus’ population, which can be suggested from the *Primary Chronicle* and archaeological data^2,3^. We analyzed our Rus’ samples in comparison to a large set of previously published genomic data designated as Viking Era (VKE) samples (8th-11th centuries CE)^19^ (see Supplementary Informations S2.2.1). An individual (VK56) from the Scandinavian western coast of Gotland Island is connected to two related individuals from the Ancient North Rus’ genetic cluster (Supplementary Fig. 35), implying a potential connection between this Scandinavian site and the Northern Rus’ . We also detected a single shared IBD segment between the Scandinavian sample VK505^19^ and another Northern Rus’ sample (Supplementary Fig. 54). Importantly, the low number and the length of the aforementioned shared genomic segments show that these individuals have been distanced by numerous generations from a common ancestor who lived several centuries before the Viking Era.

Next, we observed that some of the VKE samples appear to be positioned within Ancient Major Rus’ genetic clusters on the PCA plot (mostly the Rus_SWest and Rus_Baltic genetic subclusters) (Supplementary Fig. 35). This VKE group represents a heterogeneous set of individuals from Scandinavia and Rus’ areas^23^. When focusing only on Viking groups with a predominance of Scandinavian ancestry and originating from present-day Danish and Swedish regions (see Supplementary Informations S2.2.1, Supplementary Fig. 30), we identified that the Viking samples form a separate cluster that is just partially overlapped with Rus_Baltic and Rus_SWest subclusters. The differences between Vikings and Rus’ populations were basically confirmed by outgroup *f*3-statistics (Supplementary Table 12, Fig. 8, Supplementary Fig. 38). Overall, our analysis revealed that only a limited number of individuals from various regions of ancient Rus’ exhibit possible genetic traces of Scandinavian Viking ancestry. Notably, no genetic similarities were identified between the genetic profiles of individuals from the barrows of the military elite (Druzhina warriors) in the Gnezdilovo and Nikolskoe mounds and those of Scandinavian Vikings (Supplementary Fig. 34b, Supplementary Fig. 35, see Supplementary Informations S2.2.1). This finding does not support the hypothesis of a likely Scandinavian ancestry for the Druzhina warriors buried in at least these two studied burial sites (Gnezdilovo and Nikolskoe).

### The genetic legacy of the Rus’ ancestries in present-day populations

On the PCA plot, the majority of Rus’ individuals are positioned within clusters of present-day Russians/ East Slavs (Fig. 3). A gradient-like shift with decreasing ancient Siberian ancestry and increasing the European Neolithic farmers’ component along the northeast-to-southwest direction was revealed by unsupervised ADMIXTURE analysis both among the ancient Rus’ and present-day Russians and other Slavic populations of Europe (Fig. 4b).

We performed outgroup *f*3-statistics analysis to measure allele sharing between ancient Rus’ and present-day individuals (Supplementary Table 17-20). We observed differences in allele sharing between two Rus’ genetic clusters. While the individuals of Ancient Major Rus’ subclusters (Rus_Core, Rus_Baltic, Rus_SWest) exhibit genetic affinities with all present-day East Slavic and Baltic (Lithuanian, Estonian) populations, the Ancient North Rus’ subclusters (Rus_North_1, Rus_North_2) demonstrate greater genetic drift with Northern European Fenno-Ugric groups (Veps, Karelians, Saami) and Northern Russians from the Arkhangelsk region (Supplementary Table 18, Supplementary Fig. 43, 44). These data were supported by analysis of the f4-statistics (Supplementary Table 22, Supplementary Fig. 47, 48). F4-cladality test results (Supplementary Fig. 48) suggest that individuals from both Ancient North Rus’ subclusters likely represent a community that shares a common ancestor with Veps, Saami and contemporary Northern Russians from Arkhangelsk. Notable, the testing of present-day Baltic Finnic groups (Estonian, Finnish, Karelian, Estonian, Veps) provides further evidence of genetic differences of North_1 and North_2 subclusters, as initially revealed by ADMIXTURE analysis and qpAdm modeling (Supplementary Table 25f, Supplementary Fig. 49a, Fig. 4). We revealed distinct patterns of genetic affinity with present-day Baltic Finnic groups. Specifically, Rus_North_1, unlike Rus_North_2, demonstrated a significant connection to present-day Karelian and Finnish populations (Supplementary Fig. 48).

Overall, our results indicate genetic continuity between ancient and present-day East Slavic groups. Ancient northern Rus’ inhabitants had notable affinity to several present-day Fenno-Ugrian (Uralic) groups from the northern populations of Russia.

### Phenotypes

Medieval chronicles depicted the ancient Rus’ population (Eastern Slavs) as tall, fair-skinned, light-haired (blonde to brown), and blue-eyed^34–37^ (see Supplementary Informations S2.9.1) To examine the probable pigmentation phenotypes in the ancient Rus’ groups, we explored whole genome analysis along with HIrisPlex multiplex and *HERC2* rs12913832 single marker assays that we developed for highly degraded DNA (see *Materials and Methods*, Supplementary Informations S2.9). As a result, many medieval Rus’ inhabitants were predicted to have prevalently blue eyes, light hair (blond to light brown), and all tested individuals had skin tones ranging from pale to intermediate (Supplementary Fig. 66, Supplementary Table 35). These traits were prevalent in both Northern Rus’ and Central Rus’ Volga-Oka regions. These findings are congruent with the descriptions made by medieval authors^34–37^ (Supplementary Information S2.9.1)

## Discussion

The second half and end of the first millennium CE is a remarkable period in European history, defined by the extensive migration of Slavic peoples across enormous territory and the establishment of the first Slavic states or kingdoms^38^. The genetic data for the populations that established these primary medieval states is sparse. The consolidation of people of numerous East Slavic local groups mentioned in the *Primary Chronicle*, along with their presumed contacts with their neighbours from Eastern and Northern Europe, culminated in the establishment of the Rus’ state, which eventually united various principalities and cities.

For the archeo-genomic study, we used a large sample set from Slavic and non-Slavic burials of the early period of Rus’ establishment. Our findings reveal clear genetic continuity between the inhabitants of ancient Rus’ and contemporary Russians across the broad territory of Eastern Europe. Their genetic profiles are substantially different from those of non-Slavic mediaeval and present-day European populations (Fig. 2, Fig. 4).

We elucidated the complex structure of the ancient populations that settled in different locations of the emerging Rus’ state including its northern periphery. We identified two genetically distinct groups: the Ancient Major Rus’ and Ancient North Rus’ genetic clusters, which represent the foundational communities that contributed to the formation of the Rus’ state. We can assume that the first group corresponds to the East Slavic groups described in the *Primary Chronicle*, which have long been regarded as the principal communities in the early history of ancient Rus’. The second ones mentioned in the *Primary Chronicle* as non-Slavic groups, may be associated with our Ancient North Rus’ cluster, which is linked to Fenno-Ugrian (Uralic) populations. We have demonstrated the genetic heterogeneity of ancient East Slavic groups within Rus’, revealing that the Slavic-related component of Rus’ population comprises at least three genetically different subclusters: Rus_Core, Rus_Baltic, and Rus_SWest. Notably, individuals assigned to specific genetic subclusters are originated from diverse geographic regions (Supplementary Fig. 34). Thus, we provide genetic evidence that the Slavic communities referenced in the *Primary Chronicle* as early Slavic units despite their separate territorial locations are represented by individuals with the shared heterogenous genetic pool apparently of common origin by the time they united to form the ancient Rus’ state, as some historians had previously suggested^5,7,12^. According to our data, the ancestry of all these groups was based on the common Baltic (or Balto-Slavic) genetic substratum that was present in Eastern Europe and the Baltic region at least from the BA. This ancestry is maximal among the Rus_Baltic subcluster’s individuals. The minor influence of other ancient populations, such as those from Central and Western Europe and Uralic groups who carried ancient Siberian component are also contributed to some genetic profile variabilities of Ancient Rus’ genetic subclusters.

People of medieval Northern Rus’ population of both Rus_North_1 (that can be termed as “Beloozerians”) and Rus_North_2 genetic subclusters could represent groups originated from chronicler non-Slavic Chud’ or Ves’ ethnic units which settled on the northeastern periphery of Rus’ on the northern portages and river routes. These people culturally demonstrate the Finno-Slavic blend and had Fenno-Ugrian (Uralic) genetic profiles. The high rate of genetic affinity of the Ancient North Rus’ individuals with present-day Vepses (Supplementary Fig. 43) is in line with the hypothesis that the Vepses are related by genetic origin to the legendary Ves’ and Chud’. Overall, the Ancient North Rus’ genetic groups are based on the same substrate related to ancient Baltic region populations as Ancient Major Rus’ do. It is important to note that our analysis identified that each tested northern region (Beloozero and Poonezhye) groups represent relatively isolated endogamic populations (Fig. 9).

Northern Rus’ Beloozero region remained beyond the sphere of expansion of the East Slavs until the 10th century CE^39^. For the first time we performed here a trace of primary interaction between the Slavic and Fenno-Ugrian (Uralic) peoples. In particular, Nikolskoe burial site in the Northern Rus’ Beloozero region dated to the 11th century CE has a distinctly ancient Rus’ East Slavic cultural appearance and is represented mostly by individuals with Slavic-related genetic profiles. This differentiates Nikolskoe from the other Beloozero burial grounds, which exhibited a blend of Slavic and Finnish cultural elements, and a predominantly non-Slavic genetic profile of Ancient North Rus’ genetic cluster (Fig. 2). Furthermore, individuals of admixtured profiles (Rus_Middle group) were revealed as in Northern Rus’ as in the Central Rus’ burial sites. This finding supports the concept of the involvement of people of Fenno-Ugrian (Uralic) origin along with East Slavs migrating and integrating with these groups in the northeastern territories, contributing to the establishment of the whole ancient Rus’ community.

Our analysis demonstrates that the ancient Rus’ and Scandinavian Viking genetic pools had no overlap. Nonetheless, a few individuals from various regions of Rus’ exhibit genetic traces of Viking ancestry (Supplementary Fig. 30). It is of note that in Ladoga burial sites (10th-13th centuries CE, North-Western Rus’ region^19^), showing Viking culture attributes, most tested individuals had Scandinavian-type genetic profiles (Supplementary Fig. 30, Supplementary Fig. 49b). Overall, our analysis of a comprehensive set of ancient Rus’ culture samples collected from a broad geographic area did not indicate a significant influence of Scandinavian (Viking) genetic ancestry on the ancient Rus’ population as a whole.

Taken together, our findings provide the first insight into the genetic structure of various groups or ethnic units that coalesced in the second part of the first millennium CE to establish the East Slavic/Rus’ cultural community within the Rus’ state and participated in colonisation of the Northern regions. Because cremation was common prior to Christization of the medieval Rus’, CE samples from before the founding of the Rus’ state are quite scarce. The examination of such samples will be critical in providing a full view of historical events that underpin the early genesis and primary settlements of East Slavs and other communities on the East European Plain.

## Materials and Methods

### Archaeological Material

A total of 200 anthropological samples were collected from 46 archaeological sites across the East European Plain (Fig. 1). There were 36 burial sites with East Slavic/Rus’ culture attribution from different geographical regions, 4 burial sites whose cultural context exhibits a blend of Slavic and Finnish elements from the Northern Rus’ region, and 6 burial sites of other cultural groups adjacent to the ancient Rus’ area (Mordva, Muroma, Volga Bulgars, and Latgalians).

Our samples represent five geographical regions of Rus’: 1) Northern Rus’ (Beloozero and Poonezhye regions); 2) North-Western Rus’ Novgorod region; 3) Western Rus’ Dnieper-Dvina region 4) Central Rus’ Volga-Oka region; 5) Southern Rus’ region.

For the Northern Rus’ we report data on 86 individuals from seven necropolise. Among them, five represent the Beloozero region: Minino (n=22), Shuygino (n=7), Nefedyevo (n=26), Nikolskoe (n=13). One burial site is from the Poonezhye region, Voezero (n=12). And two other Northern Rus’ burial sites: Novinki (n=2) and Ploskoe (n=1). The Central Rus’ Volg-Oka region is represented by 54 samples from seventeen archaeological sites: three urban cemeteries from Yaroslavl (n=12), Pereslavl-Zalessky (n=9), Old Ryazan (n=7), and fourteen barrows and ground burial sites in the rural areas: Gnezdilovo (n=7), Luzhki (n=1), Akatovo (n=2), Velikoe (n=2), Vorobyovo (n=2), Voronovo (n=2), Zhukovo (n=1), Ziminki (n=1), Kiryanovo (n=1), Kleopino/Kokorevo (n=2), Kletnevo (n=1), Pustosh Popova (n=1), Kremenye (n=1), and Fili (n=2). North-Western Rus’ Novgorod region is represented by 16 samples from two sites: St. George’s (Yuryev) Monastery in Novgorod (n=13) and Khreple burial ground (n=3). The Western Rus’ Dnieper-Dvina region is represented by 7 samples from 5 burial sites: Volochok (n=1), Gnevkovo (n=2), Iput (n=1), Seltso (n=2), and Logoysk (n=1). The Southern Rus’ region is represented by 26 samples collected from the nine sites, four urban ones, namely Knyazhya gora (n=3), Chernigov (n=2), Lubech (n=7), Kiev (n=2) and five rural burials: Komarovka (n=10), Kurilovka (n=1), Gushchino (n=1), Moiseevskoe (n=2), and Konotop (n=1). In order to compare the results with geographical and temporal context, we also included representatives of synchronous neighbouring non-Rus’ and non-Slavic groups from 6 archaeological sites, such as the burial sites of Muroma (Podbolotyevsky), Mordva (Korino, Muranka, and Pogiblovo), Volga Bulgarians (Syut-Siriami), and Latgalians (Lucinsky).

### Permissions for archaeological research

Each sample used in this study is managed by an archaeologist, museum curator, or researcher who is the author of the paper. Archaeological samples were excavated by archaeologists in the territory of the former Russian Empire and the former Soviet Union before 1991, as well as in the territory of the Russian Federation. All samples of anthropological materials used in this study were obtained from the collections of the Museum of Anthropology at Lomonosov Moscow State University, from the Institute of Archaeology, Russian Academy of Sciences or were provided by archaeologists. The detailed information on each sample is provided in the Supplementary Information 1.

These include the organizations that conducted the excavations, the head of archaeological expeditions, primary information regarding burials, which is documented in the official reports of archaeological data, and appropriated references.

### Radiocarbon data

New radiocarbon dating for this study was performed on the bone and tooth fragment (Supplementary Materials S1.4). The results were calibrated according to the IntCal20 atmospheric curve^40^.

### DNA extraction library preparation and sequencing

DNA was extracted from petrous bones, auditory ossicles, or teeth as described previously^41^. 50-200 mg of tooth or bone powder were incubated for two to four hours at 56°C in the PrepFiler BTA buffer (Thermo Fisher Scientific) with 13 mmol DTT and 0.6 mg/ml proteinase K at a temperature of 56°C for 2-4 hours. After that, 10 volumes of Qiagen PNI buffer were combined with lysate supernatant before it was applied to the MinElute Spin column (QIAGEN). After being purified with Buffer PE (QIAGEN), DNA was eluted from the column using 30 mkl of Buffer EB (QIAGEN). A negative control was used in each cycle of DNA extraction. The High Sensitivity DNA Chip was used to evaluate DNA and negative controls with the Bioanalyzer 2100 (Agilent).

The resulting DNA was used to prepare libraries of single-stranded DNA fragments^42^. Some samples with the best DNA preservation (according to results of test low-coverage sequencing) were additionally treated with PreCR® Repair Mix (New England Biolab) before library preparation. A total of 303 genomic libraries were prepared. Libraries were firstly sequenced on Illumina MiSeqFGx and then on Illumina NovoSeq6000 in single-read mode.

### Bioinformatics analysis

We used bcl2fastq (v2.20) for demultiplexing and base calling of sequencing runs. Sequencing adapter sequences were trimmed from the raw reads and reads shorter than 25 nucleotides were removed from further analyses using the AdapterRemoval (v2.3.1)^43^.

BWA software package version BWA software package (0.7.17)^44^ with the parameters -n 0.01 -o 2 -l 16500 was used to map reads against both the human reference genome (hg19/GRCh37) and the revised Cambridge Reference Sequence (rCRS, NC_012920). Duplicates were marked by the MarkDuplicates function from GATK (v4.2.5.0) software package. Analysis of the postmortem DNA damage patterns and rescaling the base quality scores specific to the damage patterns of ancient DNA were performed using the mapDamage2 (v2.2.1) software^45^. We used Shmutzi (v1.5.5.5) to estimate contamination level based on mtDNA conflicting signals^46^ and ANGSD (v0.937) to estimate contamination on the X chromosome for individuals identified as males^47^. To determine the genetic sex of each individual, we calculated the ratio of reads mapped to the X and Y chromosomes to reads mapped to autosomes, and additionally the karyo_RxRy (v1.0.1) software was used^48^.

Mitochondrial haplogroup was assigned using Haplogrep v. 2.1.20^49,50^ and further supplemented with data from PhyloTree Build 17^51^ and Yfull MTree (MTree v. 1.02.22369, https://www.yfull.com/ ). Phylogenetic analysis of the mtDNA nucleotide sequences was performed using the median network method^52^, implemented in the mtPhyl v2.8 software package^53^. To assess the affinity of our samples with any previously published present-day and ancient samples searched for matching mtDNA sequences through the GenBank (www.ncbi.nlm.nih.gov), AmtDB (https://amtdb.org) and AADR^53^ databases. For male individuals the Y chromosome haplogroup was ascertained by the yHaplo program^54^ based on informative Y chromosome markers from ISOGG (v.11, April 2016) and Yfull YTree (v. 12.04.00). Every variant found on the Y chromosome and mtDNA was examined in the IGV v. 2.4.10 browser^55^. It should be noted that low or missing coverage of some Y-chromosome sites can lead to less precise and incomplete haplogroup assignments.

A pseudo-haploid dataset was generated by randomly selected one read per SNP of 1240K SNP panel^56^ presented in The Allen ancient DNA Resource v. 54.1^57^. We used pileupCaller (from sequenceTools v1.4.0, https://github.com/stschiff/sequenceTools/tree/master) with randomHaploid option to choose an allele at a given SNP position. Only samples with more than 10,000 SNPs were used in subsequent analysis.

We performed PCA using the smartpca tools from the EIGENSOFT (v7.2.1) package^58^. Ancient samples were projected onto the first two principal components of the present-day individuals from Human Origin (HO) dataset supplemented by present-day Slavs from the Simons Genome Diversity Project^59^, Finnish from the 1000 Genome Project^60^ and Serbians^61^. We perform an initial PCA analysis using a set of 175 Eurasian populations.

We found our ancient Rus’ samples clustered predominantly with present-day European populations, therefore the second PCA was performed using a set of 78 European-associated populations. We also projected our medieval individuals, along with previously published ancient individuals of the Iron Age and the Middle Ages from the Russian Plain and adjacent regions of Europe and Caucasus. See Supplementary Table 6 for a sample list used in analyses.

*f*3- and *f*4-statistics were calculated with the AdmixTools v. 7.0.1 software package^62^ using admixr version 0.9.1^63^. To measure allele sharing between our analysis samples and other ancient or present-day populations (Test), we calculated outgroup *f*3-statistics of the form *f3(*Anc_Rus, Test; Mbuti). We used ancient Rus’ samples merged with Human Origin dataset calculation of *f*3-outgroup statistics when tested present-day populations, and our ancient Rus’ samples merged to 1240K dataset when analysed ancient previously published groups. Higher estimates of *f*3 indicate higher levels of allele sharing. Samples with >20000 SNPs from the 1240K AADR panel were included in the analysis. We used previously published groups of at least three genomes, but also included some single genomes in case of their significance in testing of Rus’ samples (e.g. Finland_Levanluhta, Saami_IA, Czech_EarlySlavs).

We performed admixture-*f*3 statistics in the form of *f3*(Source 1, Source 2; Rus_Middle) using ancient Rus’ genetic clusters as plausible ancestral sources to explore the potential ancestry of Rus_Middle groups. Significantly negative (Z[<[−3) admixture-*f*3 estimates indicating that the gene pool Rus_Middle can be modelled as the result of the admixture between Source 1 and Source 2.

To measure the genetic affinity of our Ancient Rus’ genetic cluster to preceding or contemporaneous ancient populations we calculated *f*4-statistics of form *f*4(Ancient_Rus’, Mbuti; Ancient_pops, Ancient_pops). The option *inbreed* was set to YES in all f4-statistics calculations. To test cladality with present-day populations we calculated *f*4-statistics of form f4(Mbuti, Modern_pops;Ancient_Rus’,Target), where Modern_pops are present-day Eurasian and Northern-African populations, and Target is a set of geographically and genetically appropriated present-day Eastern-European populations. We filtered out transitions for the f4-test with present-day populations.

We estimated ancestry proportions using *qpAdm* software from AdmixTools v. 7.0.1^62^ with the allsnps: YES option and 1240K dataset. Samples with less than 20000 SNPs were excluded from analysis (see Supplementary Informations S2.4 for details).

We performed model-based clustering analysis using ADMIXTURE (v1.3.0)^64^ in unsupervised mode on a dataset containing 2377 present-day and ancient individuals (see Supplementary Table 6). We used samples with > 10000 SNPs from the 1240K AADR panel^57^, supplemented by a set of previously published ancient samples from Rus’ and neighbouring regions^18,65^. We filtered out transitions and selected the single individual with the greatest number of SNPs for every kinship pair. We used PLINK (v2.00a2.3)^66^ to exclude variants present in only one individual and performed pruning for linkage disequilibrium with the parameter --indep-pairwise 200 5 0.5. We performed 20 runs with random starting values for K in range from 2 to 12. The K value with the lowest cross-validation error was 7.

### Analysis of biological relatedness

We combined nuclear DNA, mitochondrial genome, and Y-chromosome haplogroup analysis data to identify the possible genetic kinship among the individuals of the Ancient Rus’ territory. Relationship estimation from ancient DNA (READv2) software^67^ was used to investigate kinship between the individuals. For the first run, we analysed the whole set of samples. We found a substantial proportion of relatives among the Northern Rus’ individuals in comparison to other tested samples. Then we run READv2 separately for Northern Rus’ and all other samples. We used --window_size 20000000 for Northern samples analysis to differentiate parent-offspring and sibling pairs. In all tests we restricted the analysis to transversion only and filtered out non-polymorphic and low-frequency variants (maf<0.01).

To detect relatives at a more distant degree we performed IBD analysis for samples with average mapping coverage higher than x0.25. IBD analysis consisted of the following stages: 1) BAM files filtration by mapping quality (MQ>30, read length > 30 bases, SAMtools (v1.12) program package^68^, 2) genotypes calling and SNPs filtration by quality (QUAL≥30, BCFtools (v1.9) program package^68^, 3) imputation and phasing (GLIMPSE v2.0 program)^69,70^, 4) IBD search (ancIBD program)^71^, and 5) IBD analysis (R and Python scripts). Genetic variation data from the 1000 Genomes Project (2505 individuals) were applied as a reference panel for imputation and phasing^60^ SNPs from 1,240,000 panel^57^ were used as markers for IBD calling. IBD analysis was performed for fragments longer than 8, 10, 12, 14, 16, 20, 24, 28 and 32 centimorgan.

IBD analysis of kinship was performed as described previously^71,72^. We simulated IBD distribution for different types of relationships (100 individuals per type of relationship) using the PEDSIM program^73^. For further analysis, we calculated the proportion of genome shared IBD by relatives (IBDP)^74,75^ between pairs of individuals as the total length of the genome covered by shared IBD fragments longer than 8 cM divided by the length of the diploid genome and expressed as a percentage. Then we plotted IBDP for simulated data as well as for studied individuals (Supplementary Fig. 52). The IBDP results for ancient individuals were also visualized as a phylogenetic tree (Supplementary Fig. 53). The phylogenetic tree was constructed using an unweighted pair group method with arithmetic mean (UPGMA) using R ape and phangorn packages.

To investigate distant biological relatedness, we also tested for IBD sharing among our ancient Rus’ samples and a set of 438 previously published individuals who are broadly contemporaneous and geographically close^18–20,27,29,61,65,76–79^. We only consider samples with >400,000 SNPs covered and shared IBD segments >16 cM to prevent false-positive relationships. IBD connection network analysis was performed using the ForceAtlas2 algorithm^80^ implemented in the iGraph R package (undirected graph, 1000 iterations, gravity=1000). The graph edges (links) were weighted by the maximum IBD length values obtained in the ancIBD analysis.

We calculated runs of homozygosity (ROHs) with software hapROH (v. 0.1a4)^81^ using a haplotype reference panel of 5008 phased haplotypes from the 1000 Genomes Project dataset (Phase 3, release 20130502)^60^. A total of 83 pseudohaploid data samples from our study with at least 0.3× whole-genome sequencing coverage were analysed. We counted all ROH blocks longer than 4, 8, 12, and 20 cM for each individual from different regions of Ancient Rus’ and neighboring non-Rus’/non-Slavic samples (Supplementary Table 27, Supplementary Fig. 51).

### Phenotyping

To analyze variations linked to phenotypic traits in the ancient Rus’ population, we utilized several approaches to determine genotypes of phenotype-related HirisPlex-S markers^82^. Initially, 41 HirisPlex-S markers were examined in samples with an average genome coverage >0.4x. Next, for individuals lacking data for phenotypic SNP markers from whole genome sequencing data due to insufficient genome coverage, we utilized targeted sequencing of 24 HIrisPlex markers^83^. We used 1.5 - 2.5 μl of total DNA and conducted multiplex amplification employing our laboratory-designed set of oligonucleotides for multiplex PCR, specifically targeting HIrisPlex markers for highly degraded DNA^84^. Multiplex PCR products were used for library preparation with the TruSeq DNA PCR Free DNA kit (Illumina) and *TruSeq Dual Indexes* adapters (Illumina). Libraries were sequenced on the Illumina MiSeqFGx in the 75+75 PE mode. The BWA MEM method^85^ was employed to align reads to the human reference genome (hg19/GRCh37). All phenotypic variants were examined using the IGV v. 2.4.10 browser^55^. We estimated the marker’s genotype as "NA" if covered only by one read or none. If an allele in the marker position presented by less than 20% reads, the genotype was classified as homozygous. To confirm or further specify in low-coverage samples the genotype of the *HERC2* rs12913832 marker, crucial for eye color prediction, we additionally employed genomic data for two markers, rs1129038 and rs12916300, which are in linkage disequilibrium with the rs12913832 marker^86^. Additionally, the *HERC2* rs12913832 single marker pyrosequencing assay^86^ was used for genotyping.

The HIrisPlex-S web tool was used to calculate phenotype probabilities and to predict eye, hair, and skin color (https://hirisplex.erasmusmc.nl/).

### Graphic craniofacial reconstruction of ancient individuals

The reconstruction followed the method developed by M.M. Gerasimov^87^, with additions and refinements suggested by various authors. Soft tissue thicknesses were determined using the Global T-Table, which includes data from over 19,500 adults^88^. The nasal profile was reconstructed using the method of G.V. Lebedinskaya, which is largely consistent with that of Prokopec and Ubelaker^89,90^. Nasal width was estimated using a regression equation^91^. Eye, hair and skin pigmentation were determined from genetic analysis data (see Supplementary Fig. 65).

## Data availability

The genomic data are available in the NCBI SRA under accession number PRJNA1337425. All other data needed to evaluate the conclusions in this paper are present in the paper and/or the Supplementary Materials.

## Acknowledgements

We express gratitude to Konkov A.S. for discussions and for help in the reference list preparation. We thank Protasova M.S. for technical help in manuscript preparation.

## Author Contributions

Conceptualization: E.I.R., N.A.M., A.P.B., and T.V.A. Supervision: E.I.R. Methodology:

E.I.R. and T.V.A. Resources: N.A.M., A.P.B., M.V.D., A.V.E., V.V.S., O.V.Z., P.D.M.,

N.A.D, and A.S.S. Investigation: T.V.A., A.B.M., N.A.D., M.Yu.P., I.L.K., A.D.S.,

A.D.M., G.S.D., T.V.U., and E.I.R. Visualization: T.V.A., F.E.G., G.S.D., A.V.R, I.L.K.,

I.Yu.A., S.S.K. and E.I.R. Software: F.E.G., G.S.D., T.V.U., and T.V.A. Project administration: E.I.R., N.A.M, A.P.B, and S.S.K. Writing—original draft: T.V.A., S.S.K., F.E.G., I.Yu.A., A.P.B., N.A.M., and E.I.R. Writing—review and editing: all authors.

