## Supplementary Figures for "Genetic history of Rus’"

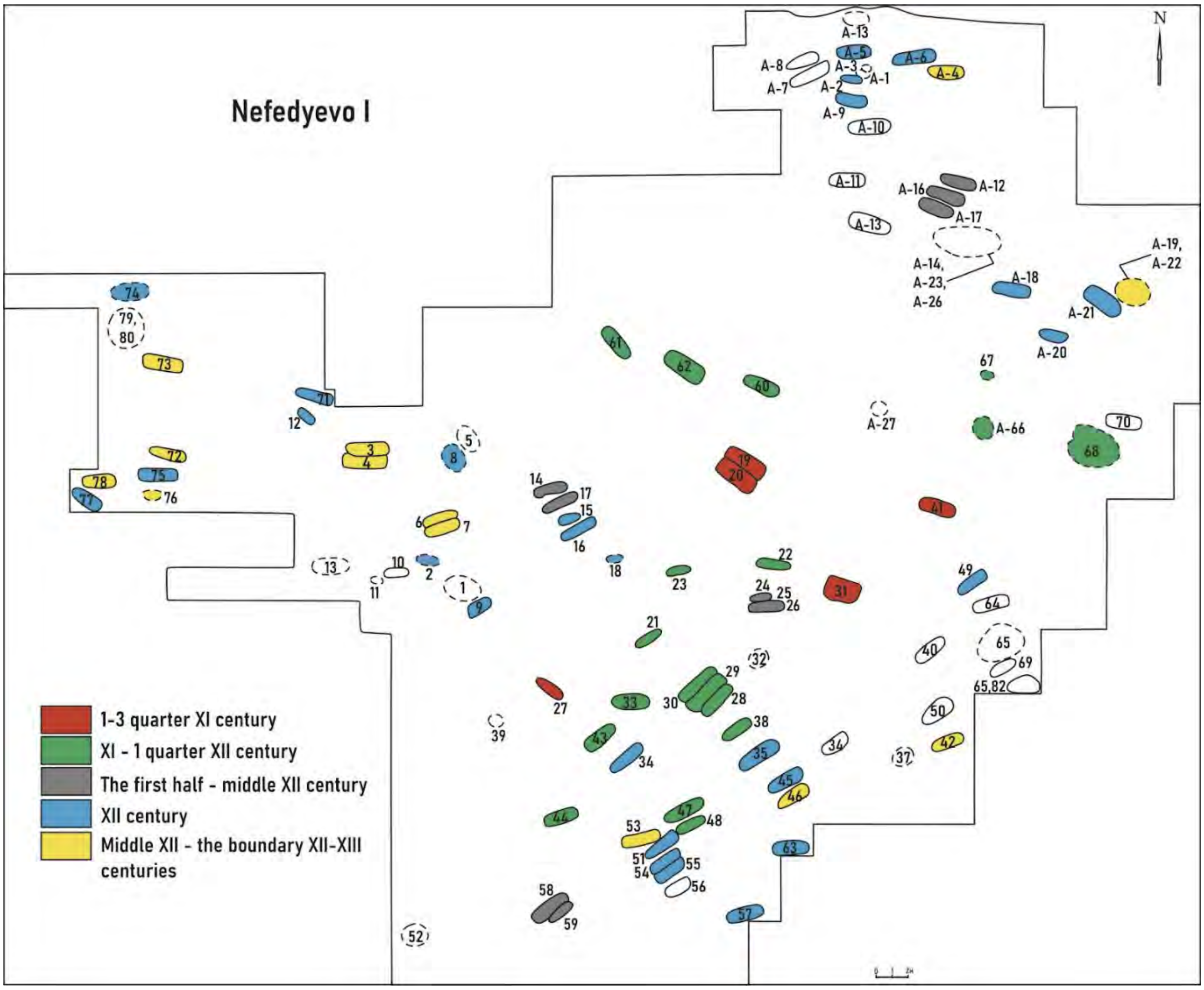


**Supplementary Fig. 1. The plan of the Nefedyevo I burial site.**

The location of Nefedyevo IB and Nefedyevo IA, and the designations of the burial dates. The burials of Nefedyevo IA marked under the letter A. The burials of Nefedyevo IB are marked without the letters. Burials from different periods are marked with different colors shown on the left.

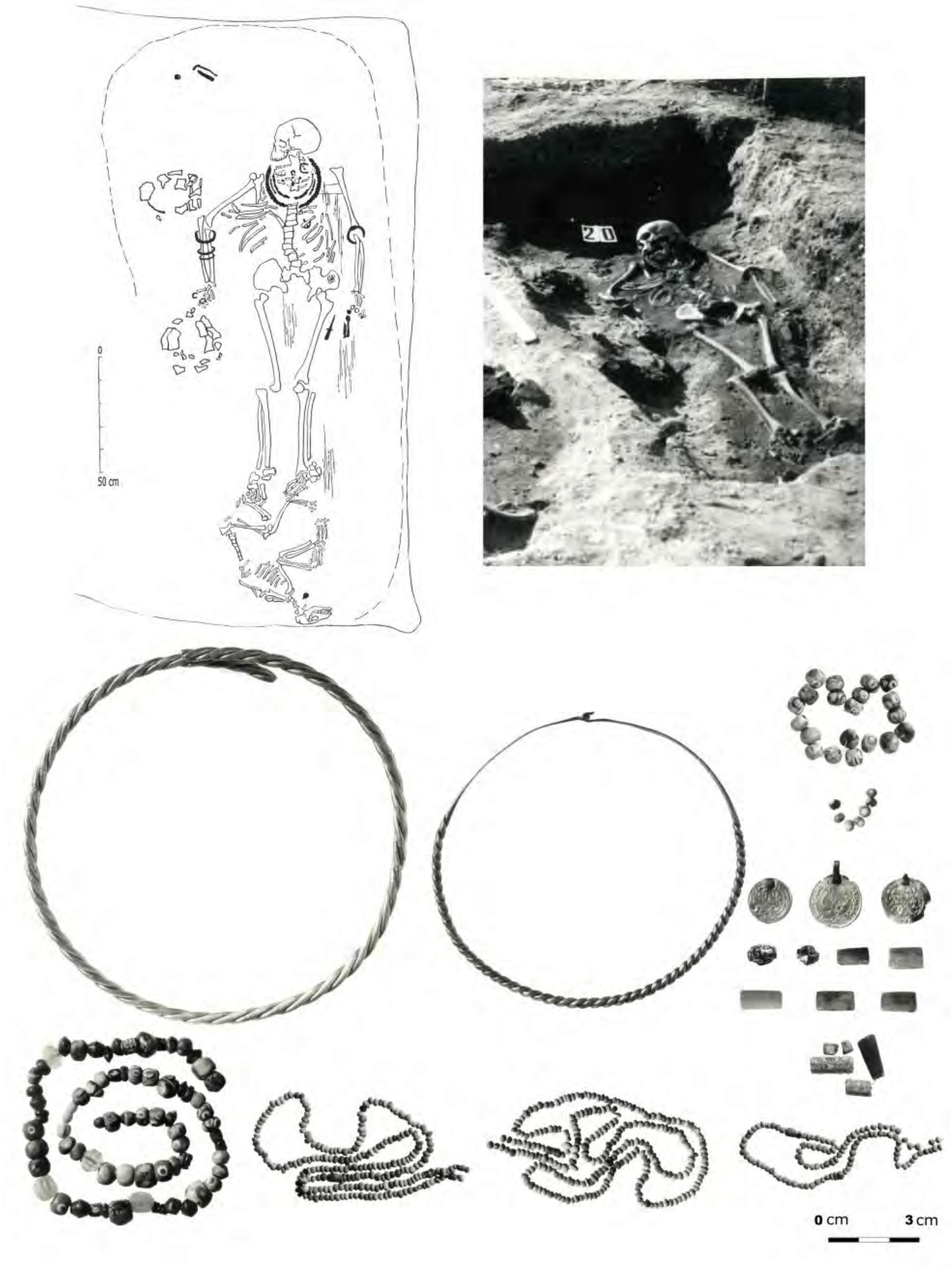


**Supplementary Fig. 2. The Nefedyevo IB burial site. Burial 20 (DB2).**

The human remains of the individual DB2 (top right), the skeleton scheme of the DB2 individual (top left) and the grave goods found at Burial 20 (below). Photo by N.A. Makarov.

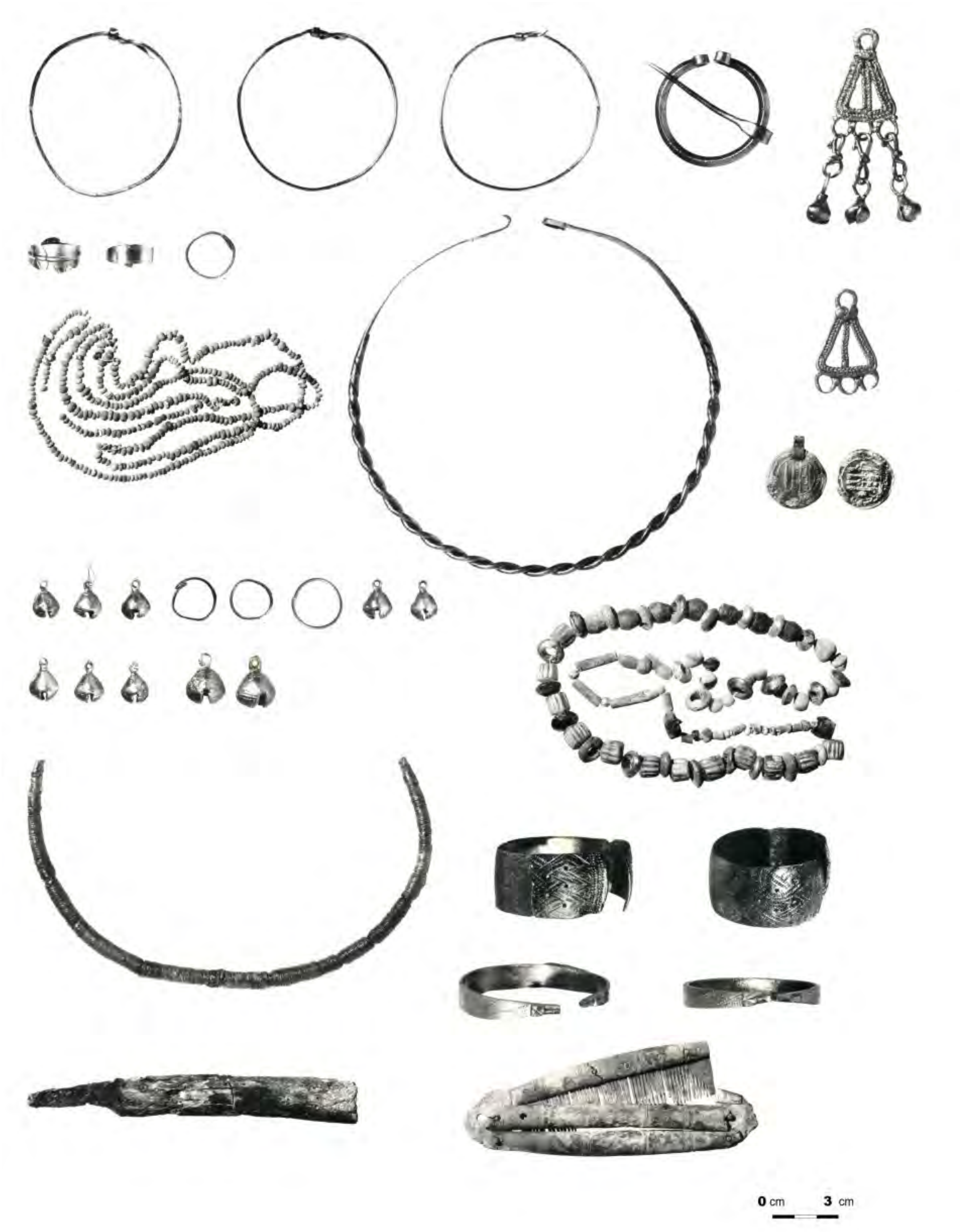


**Supplementary Fig. 3. The grave goods at Burial 41 (DB3), the Nefedyevo IB burial site.** Photo by N.A. Makarov.

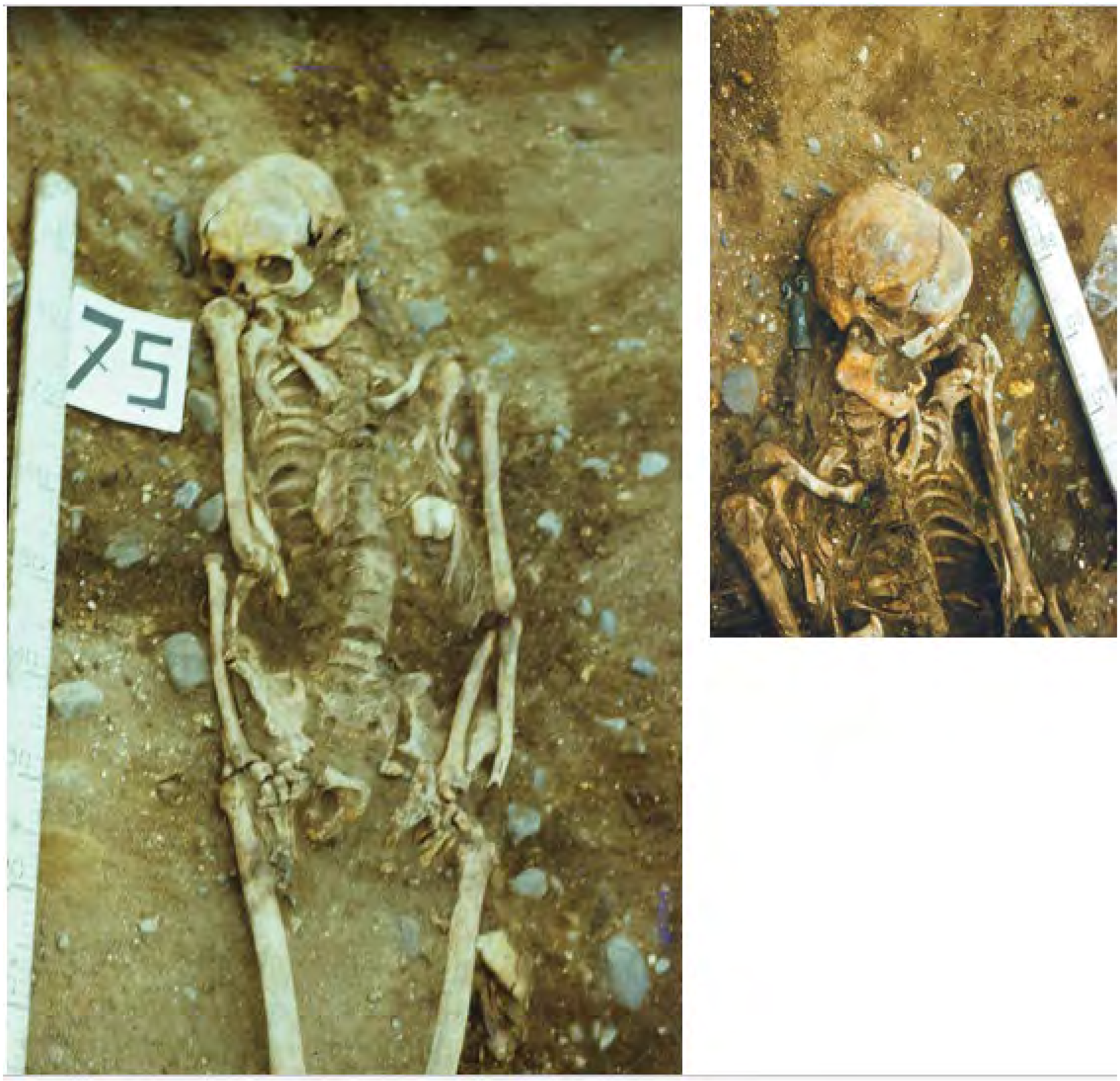


**Supplementary Fig. 4. The Nefedyevo IB burial site. Burial 75 (DB34).**

The human remains of the individual DB34 (left) and DB34 individual's skull with grave goods (right). Photo by N.A. Makarov**.**

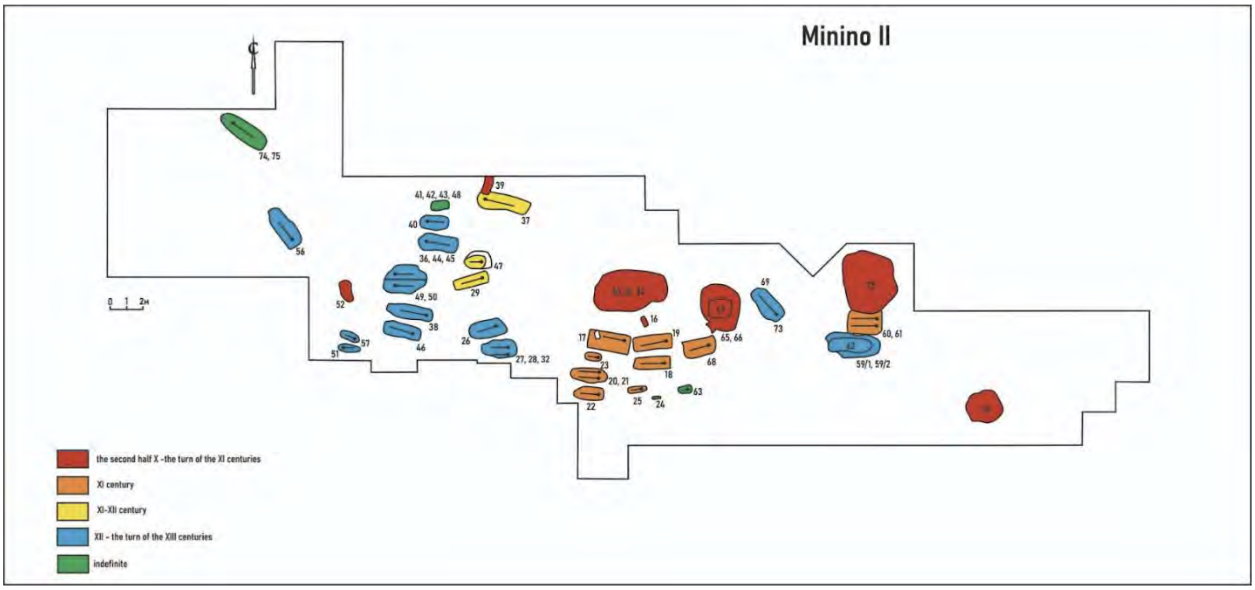


**Supplementary Fig. 5. The plan of the Minino II burial site (modified from Makarov et al, 2007).**

Burials from different periods are marked with different colors shown on the left.

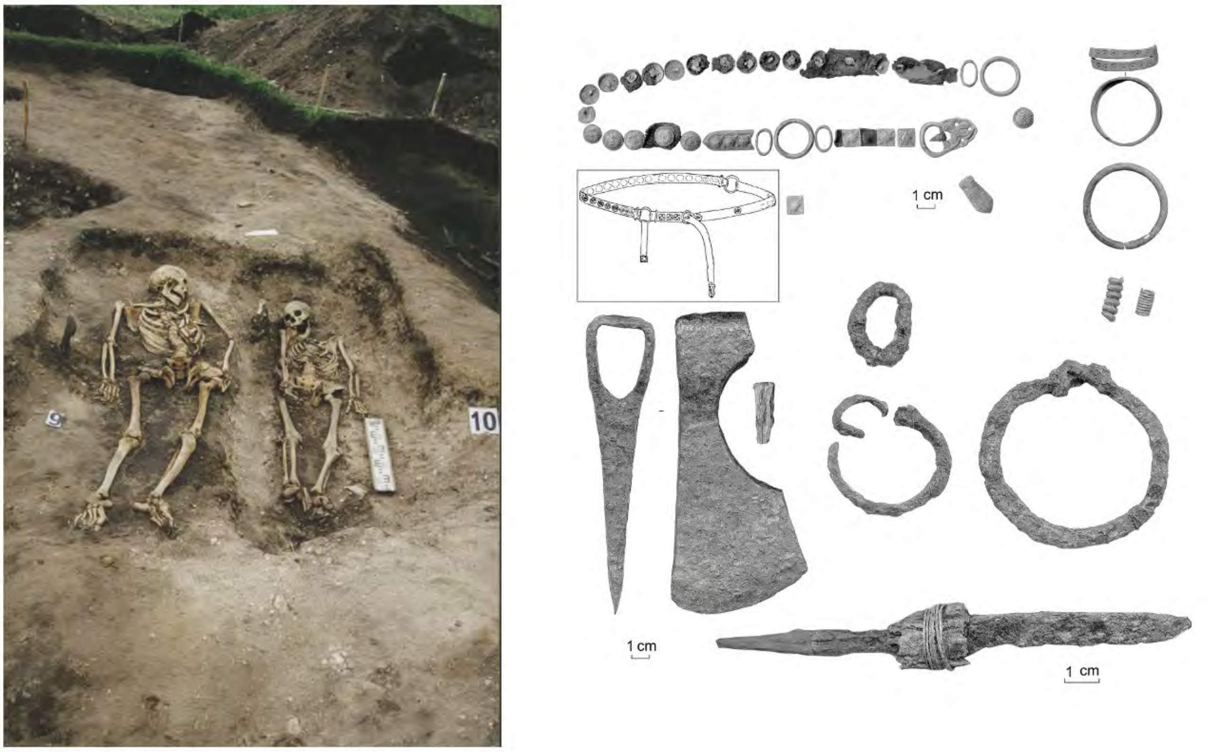


**Supplementary Fig. 6. The Minino II burial site. Burial 9 (DB9) and Burial 10.**

The human remains of the individual DB9 (left) and the female individual from the Burial 10 (not included in this study), and the grave goods found at Burial 9 (right). Photo by N.A. Makarov).

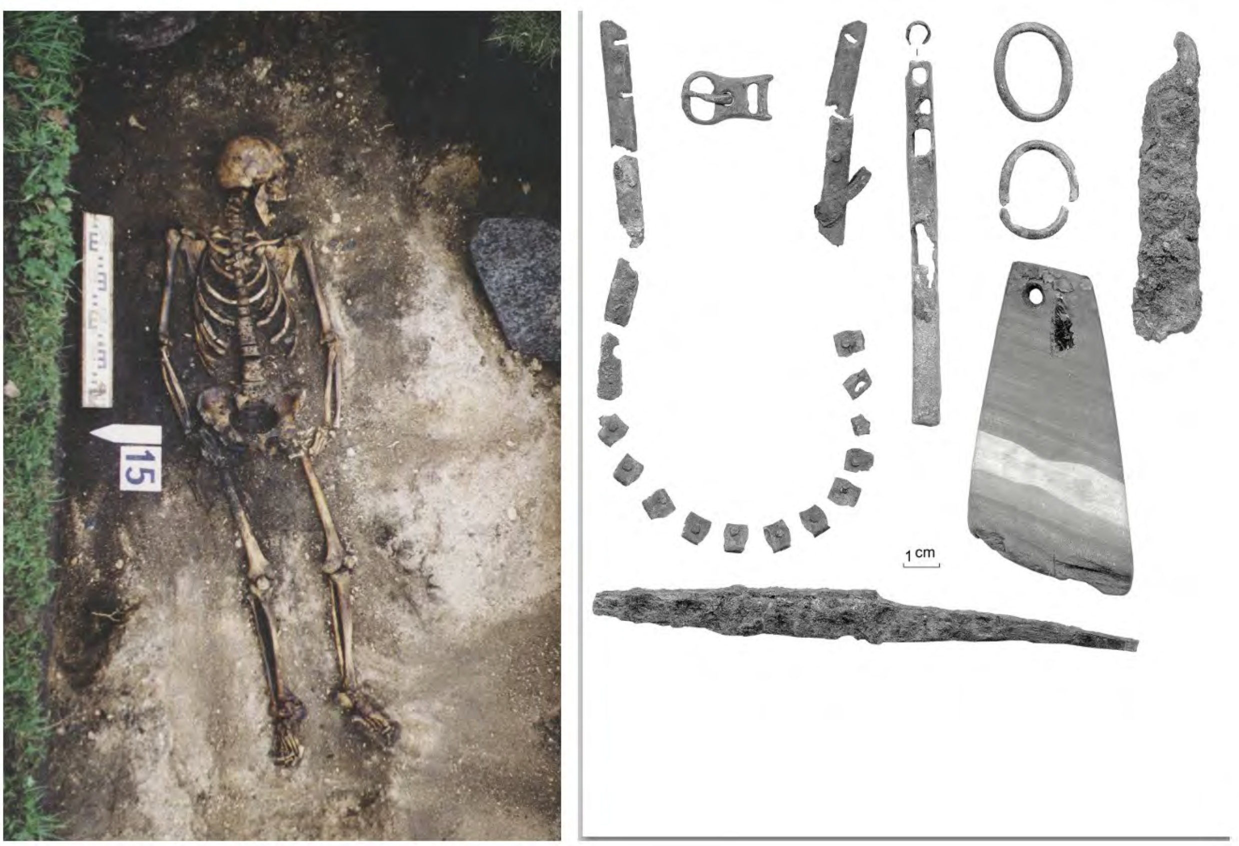


**Supplementary Fig. 7. The Minino II burialsite. Burial 15 (DB10).**

The human remains of the individual DB10 (left), and the grave goods found at Burial 15 (right). Photo by N.A. Makarov.

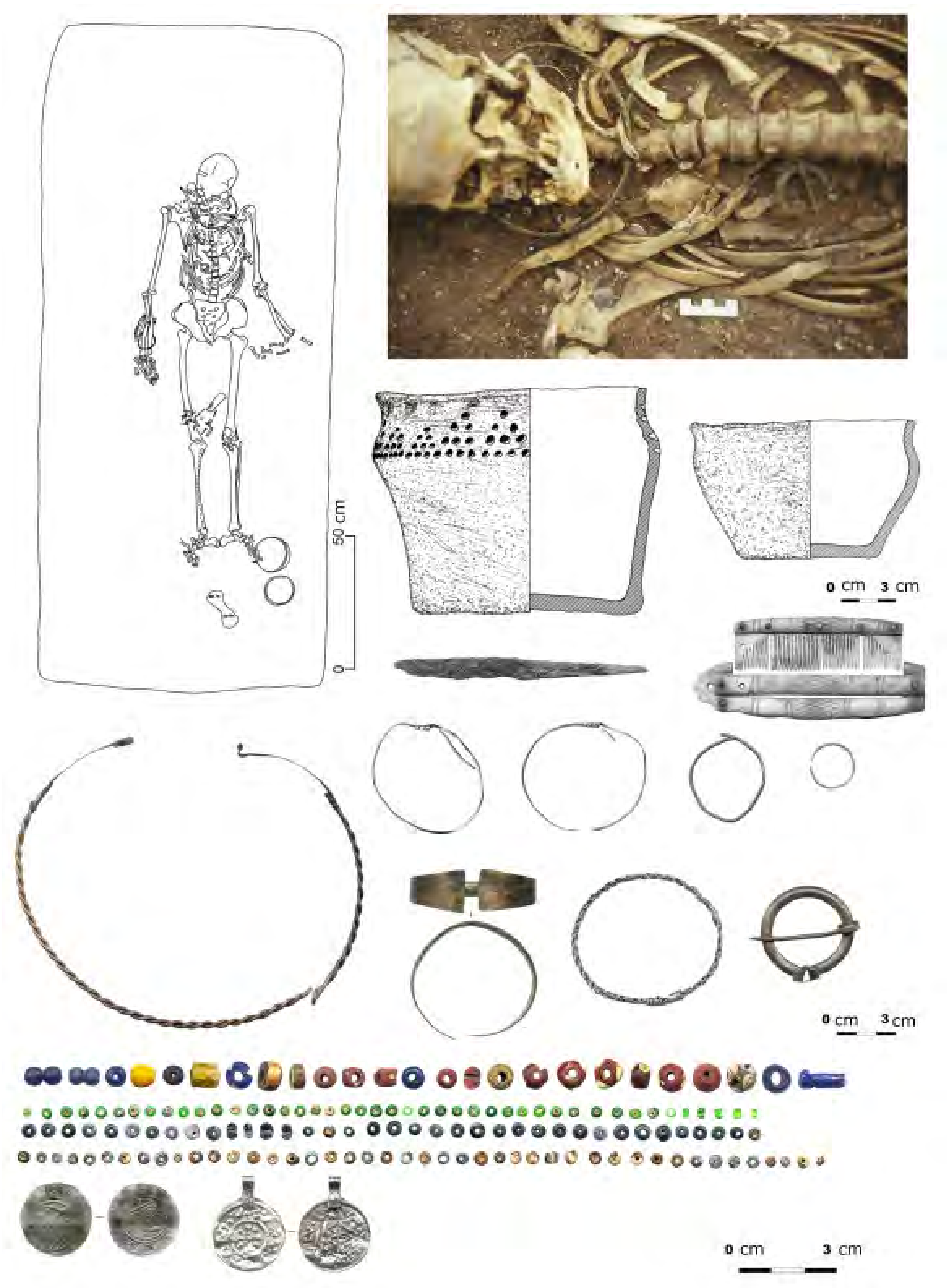


**Supplementary Fig. 8. The Minino II burial site. Burial 17 (DB41).**

The human remains of the individual DB41 (top right), the skeleton scheme of the DB41 individual (top left) and the grave goods found at Burial 17 (below). Photo by N.A. Makarov.

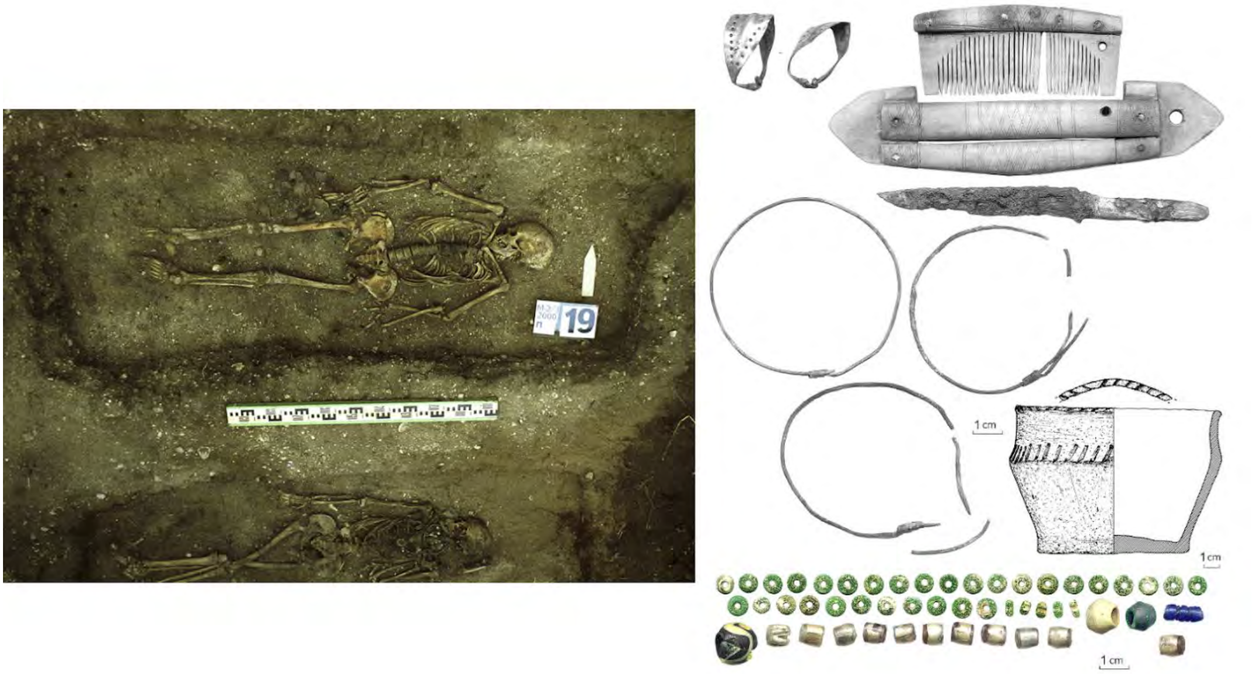


**Supplementary Fig. 9. The Minino II burial site. Burial 19 (DB42).**

The human remains of the individual DB42 (left), and the grave goods found at Burial 19 (right). Photo by N.A. Makarov.

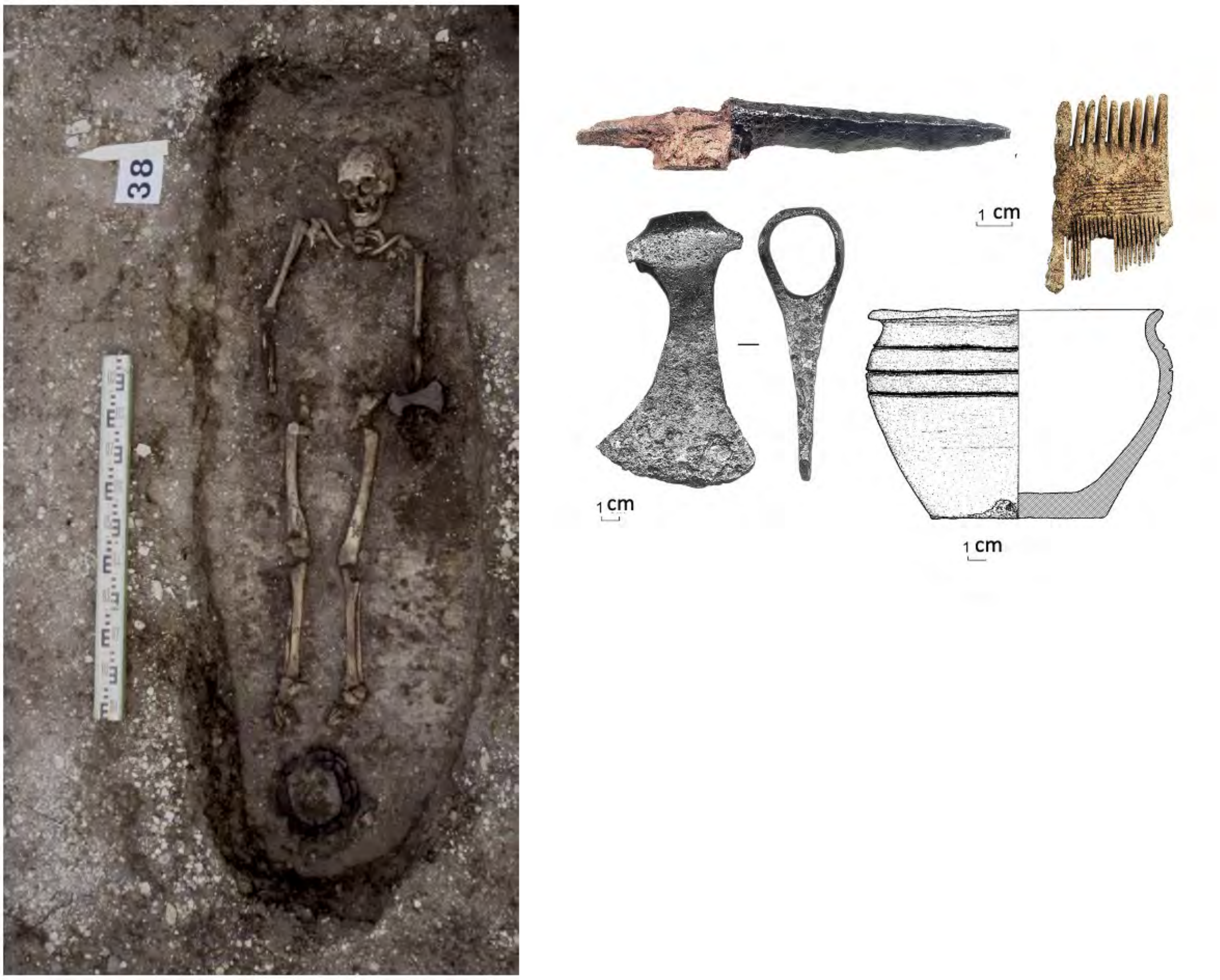


**Supplementary Fig. 10.  The Minino II burial site. Burial 38 (DB47).**

The human remains of the individual DB47 (left), and the grave goods found at Burial 38 (right). Photo by N.A. Makarov.

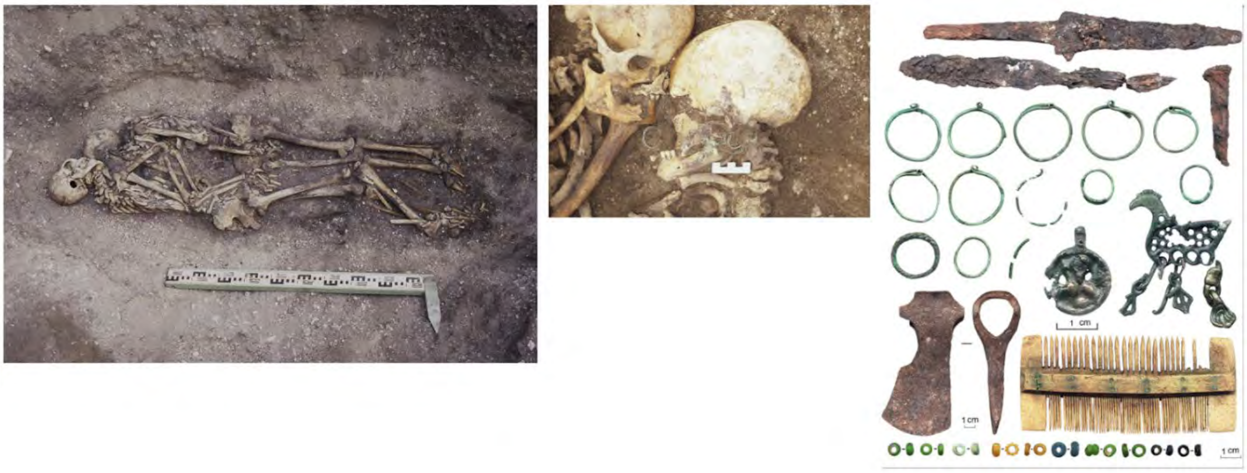

**Supplementary Fig. 11.  The Minino II burial site. Burial 48 (DB12).**

The human remains of the DB12 (left) and an individual of man (not included in this study), the skulls of these individuals (center) and the grave goods found at Burial 48 (right). Photo by N.A. Makarov.

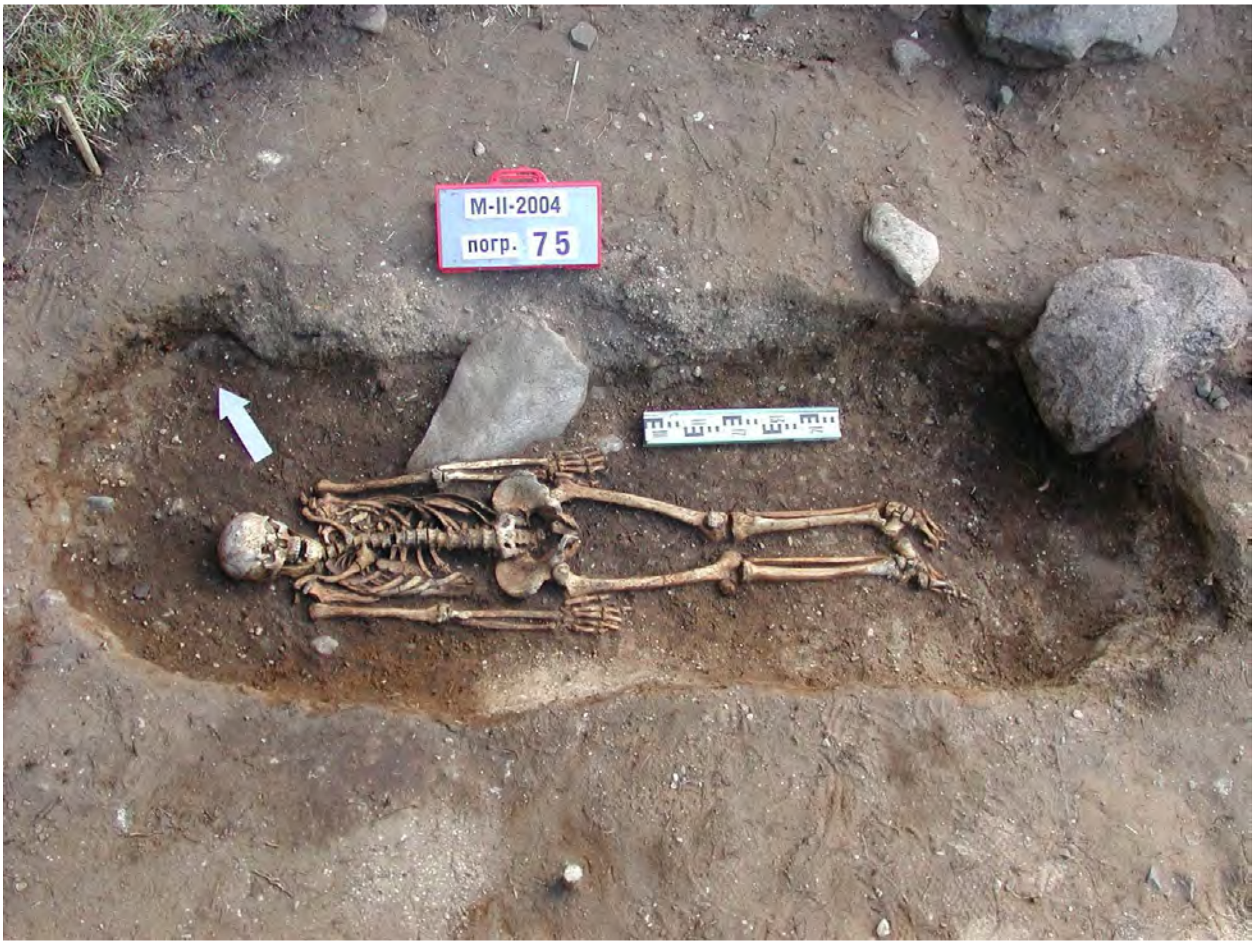


**Supplementary Fig. 12. The Minino II burial site. Burial 75 (DB54).**

The human remains of the individual. Photo by N.A. Makarov.

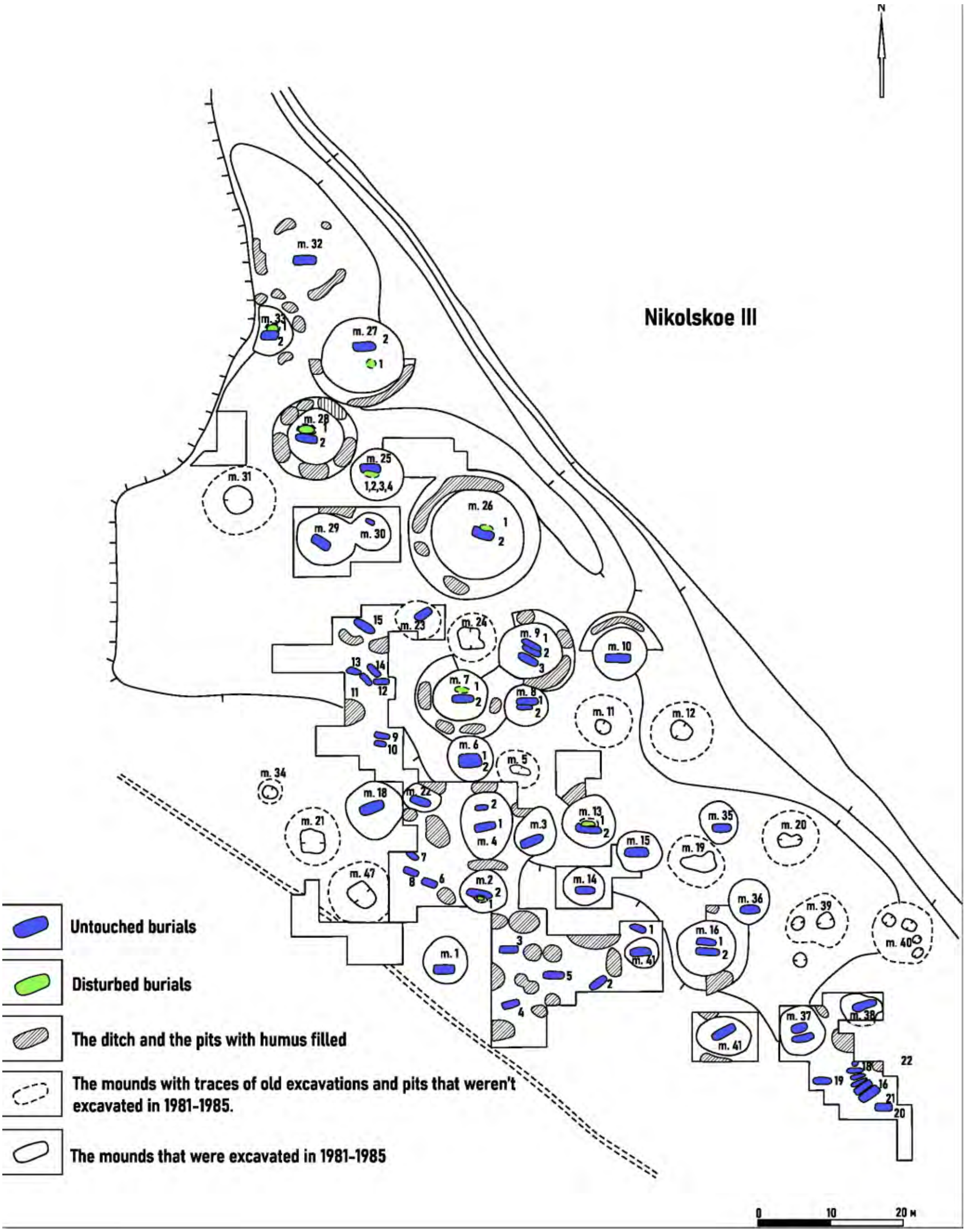


**Supplementary Fig. 13. The plan of the Nikolskoe III burial site.**

The location of individuals taken for this study (in green).The degree of preservation of the burial is shown on the left (modified from [24])

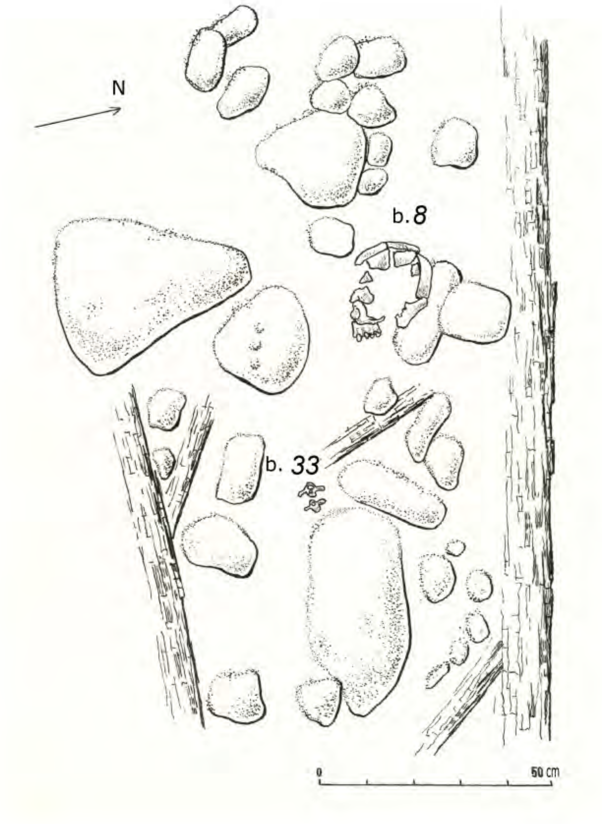


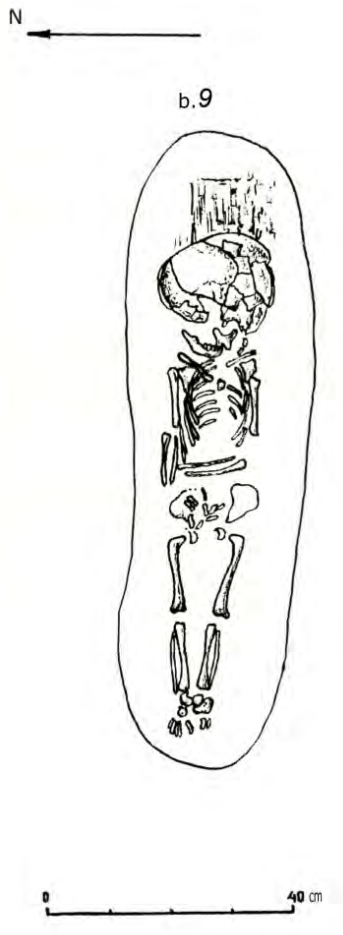


**Supplementary Fig. 14. The Voezero burial site. Burial 8 (DR12) and Burial 9 (DR13).**

The skeleton scheme of the DR12 individual (left) and the skeleton scheme of the DR13 individual (right)[25].

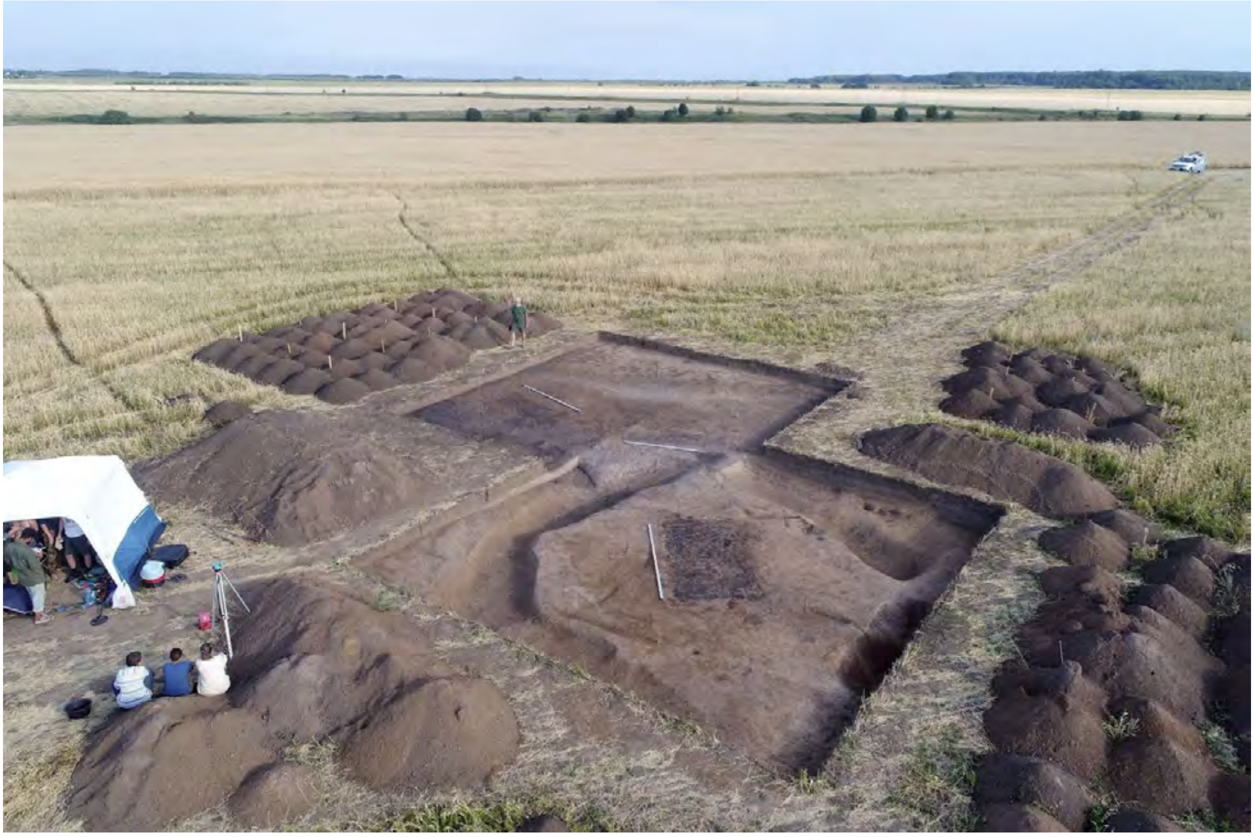

**Supplementary Figure 15. Gnezdilovo 12 burial site. Burial 19.**[28]

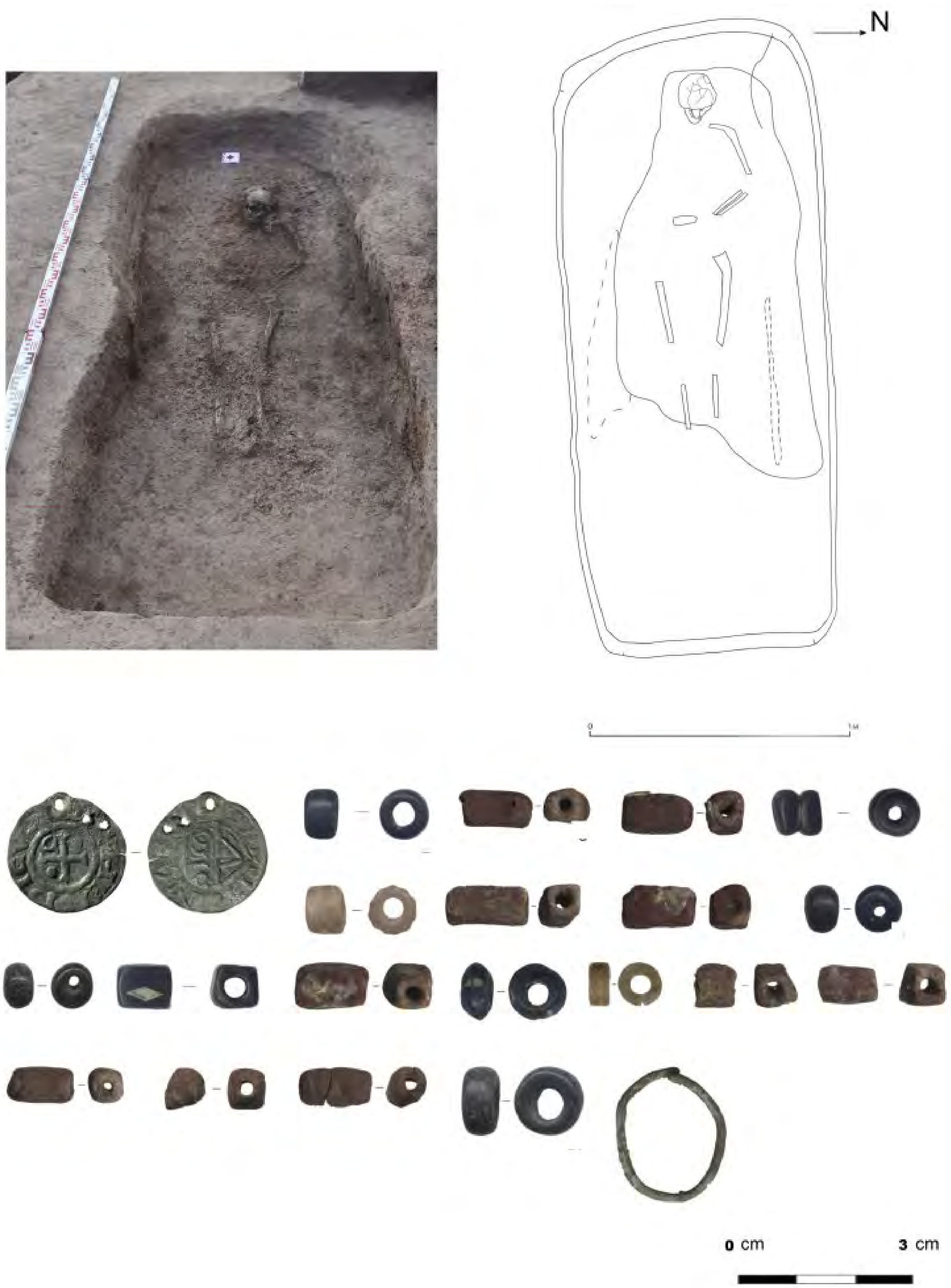


**Supplementary Fig. 16. The Gnezdilovo 12 burial site. Burial 5 (DB16).**

The human remains of the individual DB16 (top left), the skeleton scheme of the DB16 individual (top right) and the grave goods found at Burial 5 (below)[28].

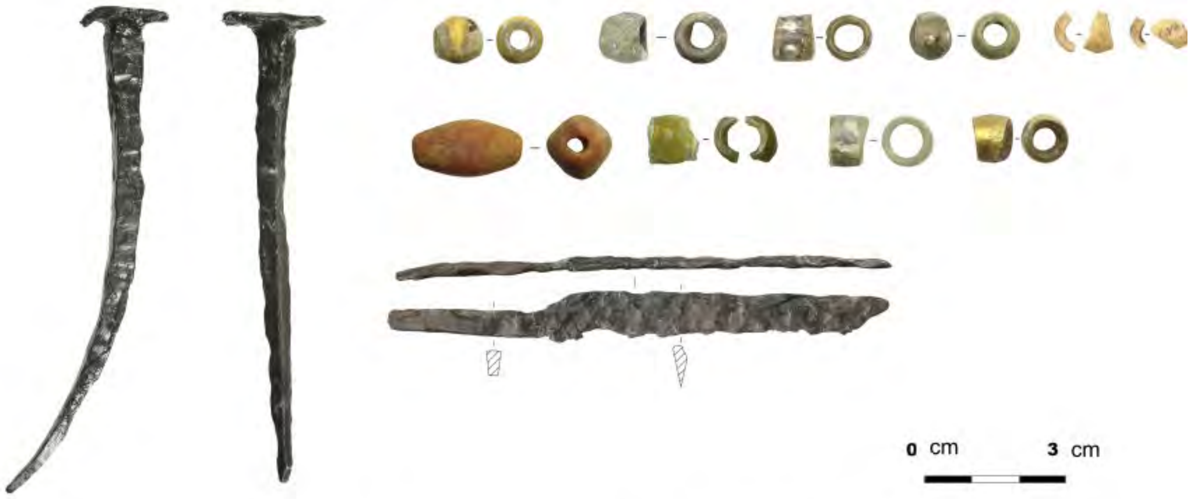


Gnezdilovo 12. Burial 16 (DB64)

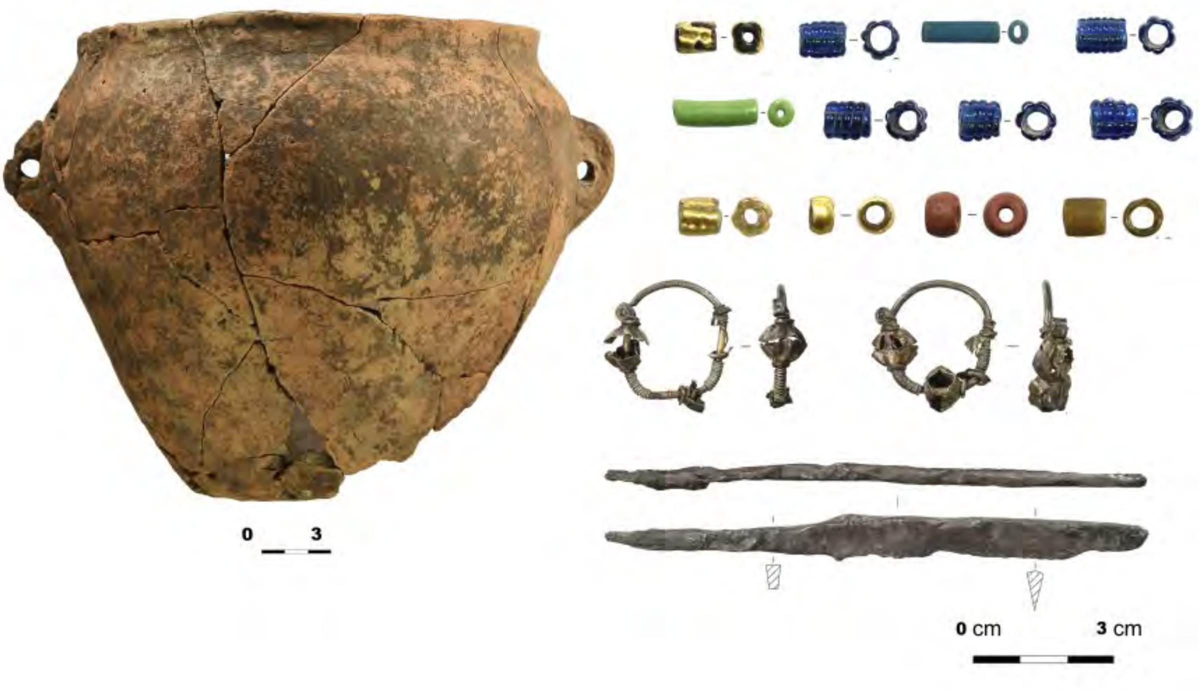


Gnezdilovo 12. Burial 17 (DB65)

**Supplementary Fig. 17. The Gnezdilovo 12 burial site. Burial 16 (DB64) and Burial 17 (DB65).**

The grave goods are found at Burial 16 (top) and Burial 17 (below). Photo by N.A. Makarov

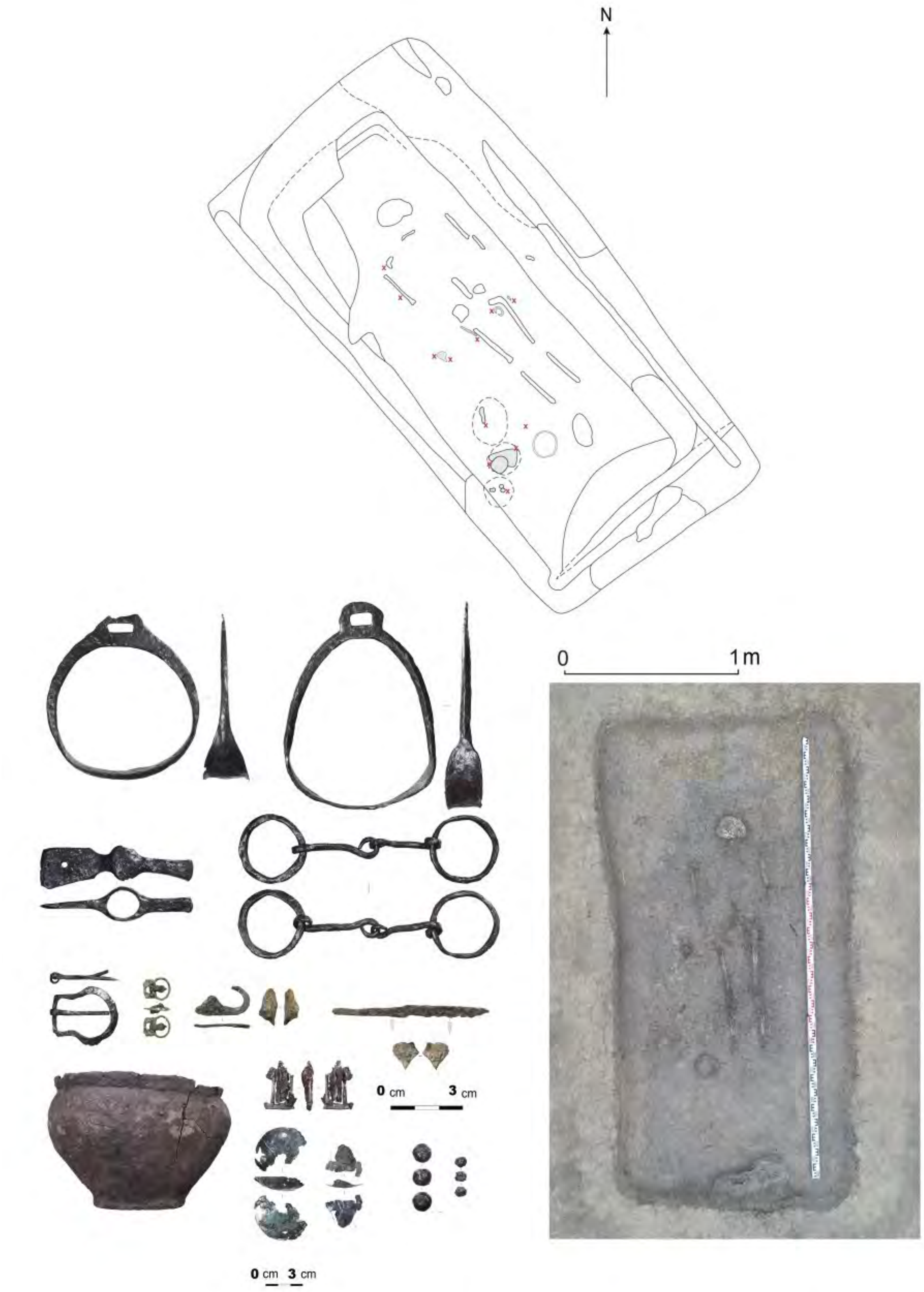


**Supplementary Fig. 18. The Gnezdilovo 12 burial site. Burial 19 (DB66).**

The skeleton scheme of the DB66 individual (top), human remains of the individual DB66 (below right) and the grave goods found at Burial 19 (below left)[28].

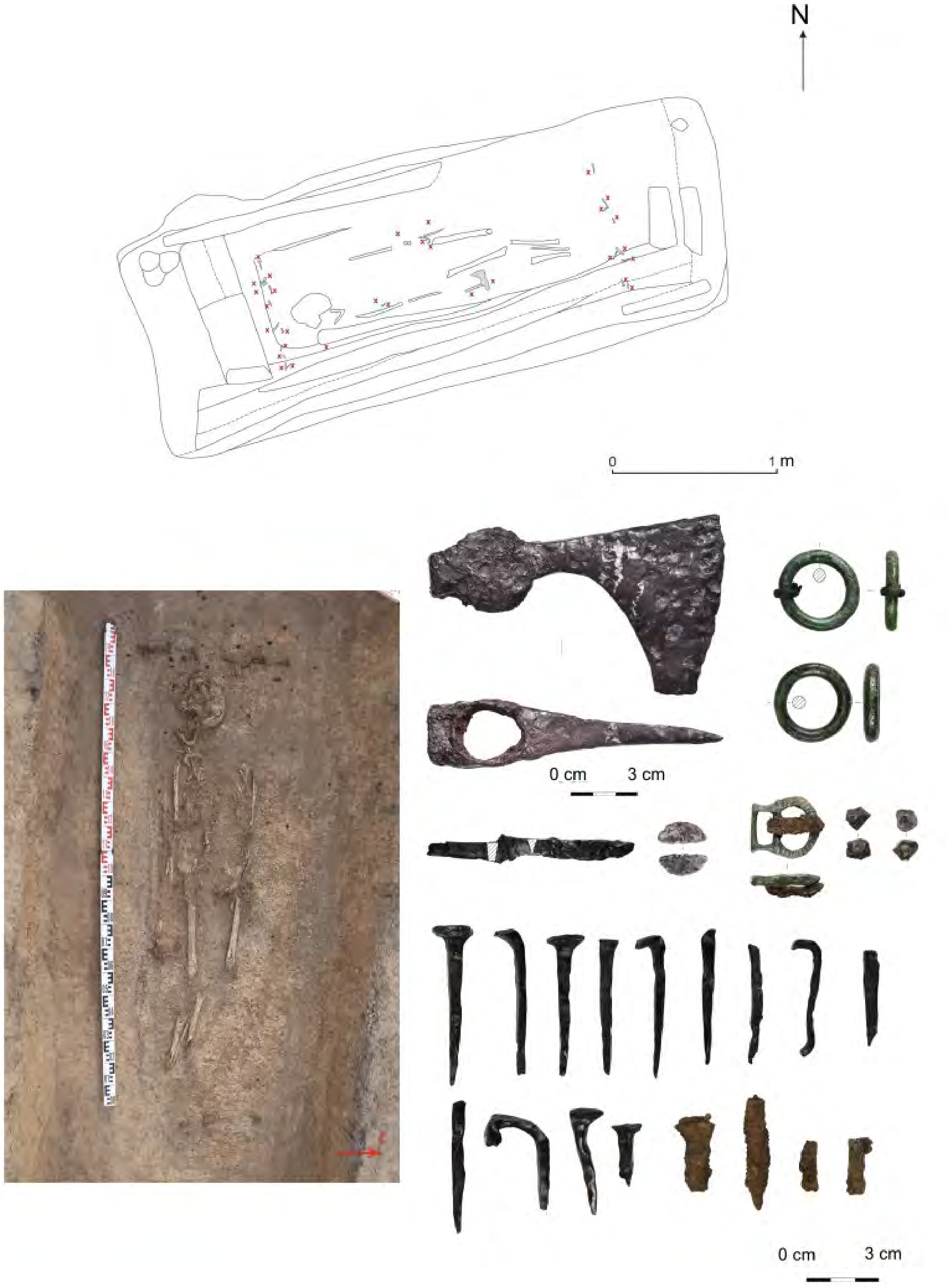


**Supplementary Fig. 19. The Gnezdilovo 12 burial site. Burial 24 (DB75).**

The skeleton scheme of the DB75 individual (top), the human remains of the individual DB75 (below left) and the grave goods found at Burial 24 (below right). Photo by N.A. Makarov.

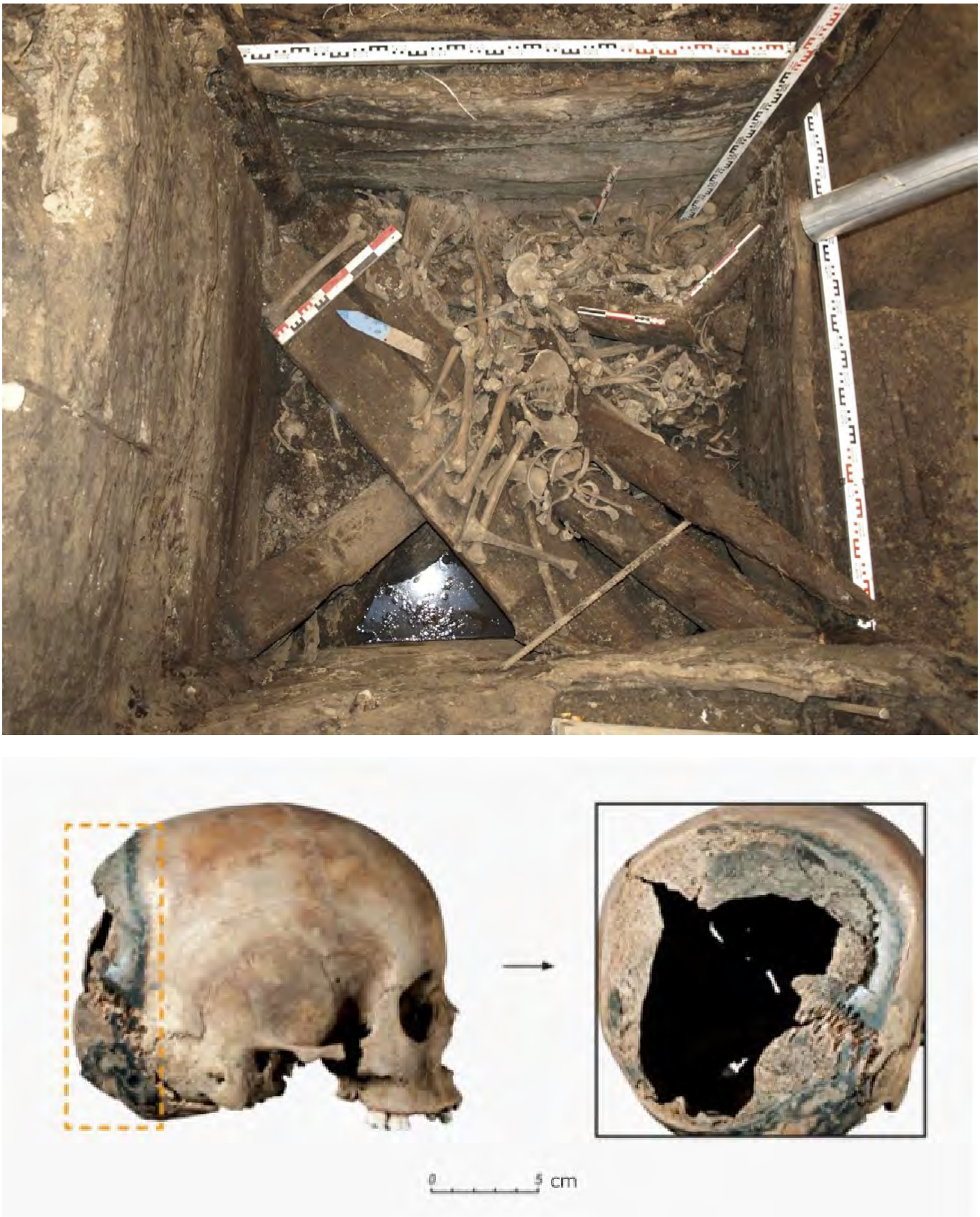


**Supplementary Fig. 20. The Yaroslavl Kremlin burial site. Burial № 110. Ya15.**

Collective burial in a well. Top view from the southwest. Tier 10. Scaffolds were thrown into the well, with human and animal bones piled between them (top), skull of the Ya15 (bottom left), the area of the charring in the posterior part of the cerebral region of the skull (bottom right). Photo by A.V. Engovatova.

.

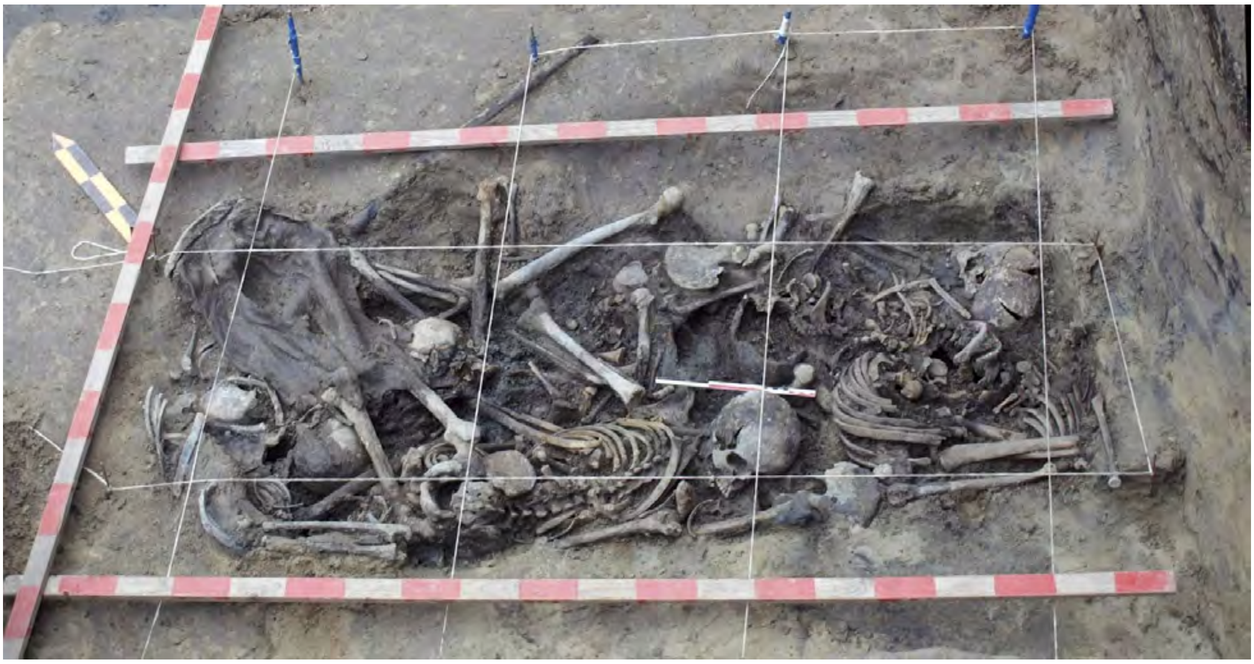

**Supplementary Fig. 21. The Yaroslavl Kremlin burial site. Burial № 76. MS.**

Collective burial (mass graves). Tiers 1-2. Top view from the southwest. MS is wearing leather boots. Photo by A.V. Engovatova.

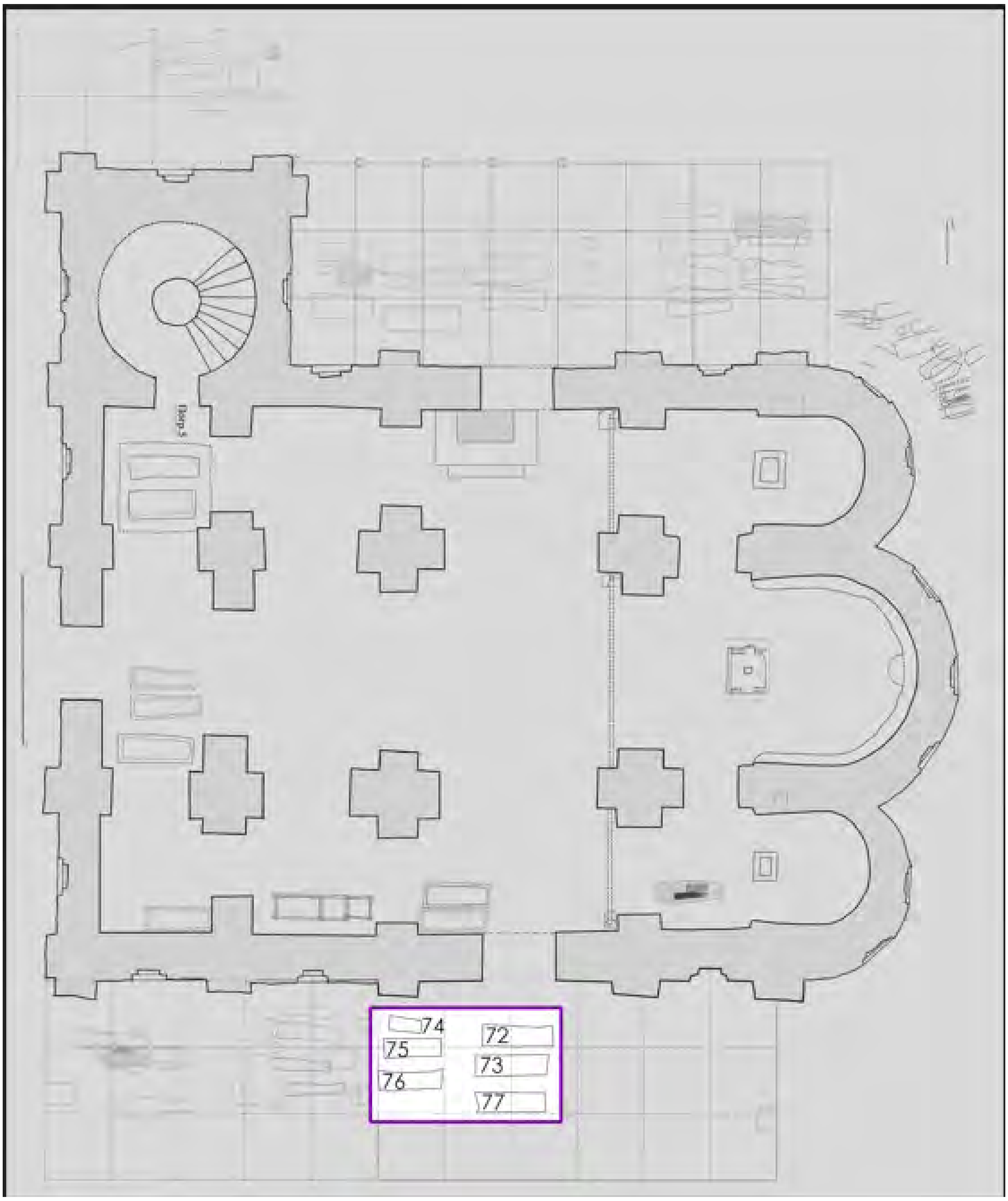


**Supplementary Fig. 22. The necropolis of the St. George (Yuriev) Monastery, around St. George's Cathedral.**

The general plan of the investigated part with the designation of burials 72-77. Drawing by Sedov V.V., Vdovichenko M.V.

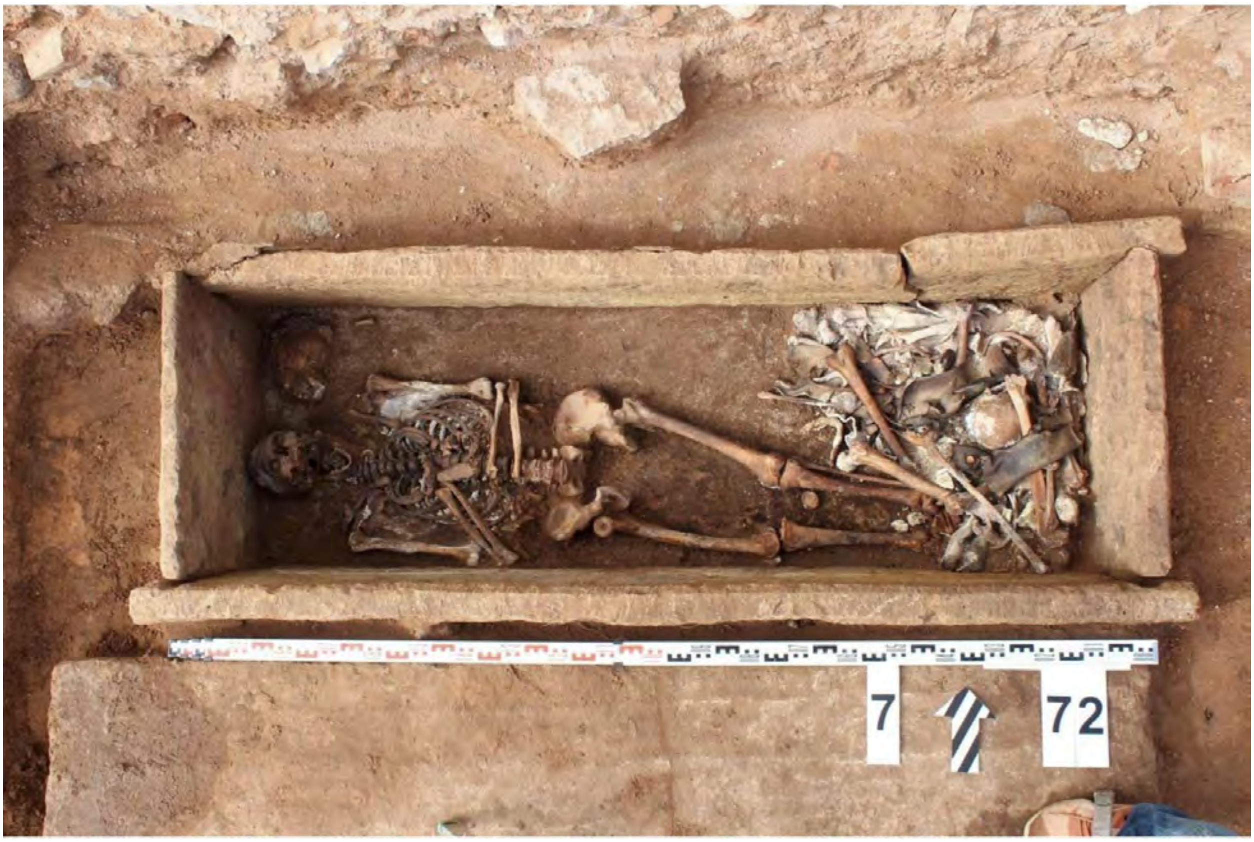


**Supplementary Fig. 23. The necropolis of the St. George (Yuriev) Monastery. Burial 72. Sarcophagus 7.**
View of the upper level. Photo provided by V.V. Sedov.

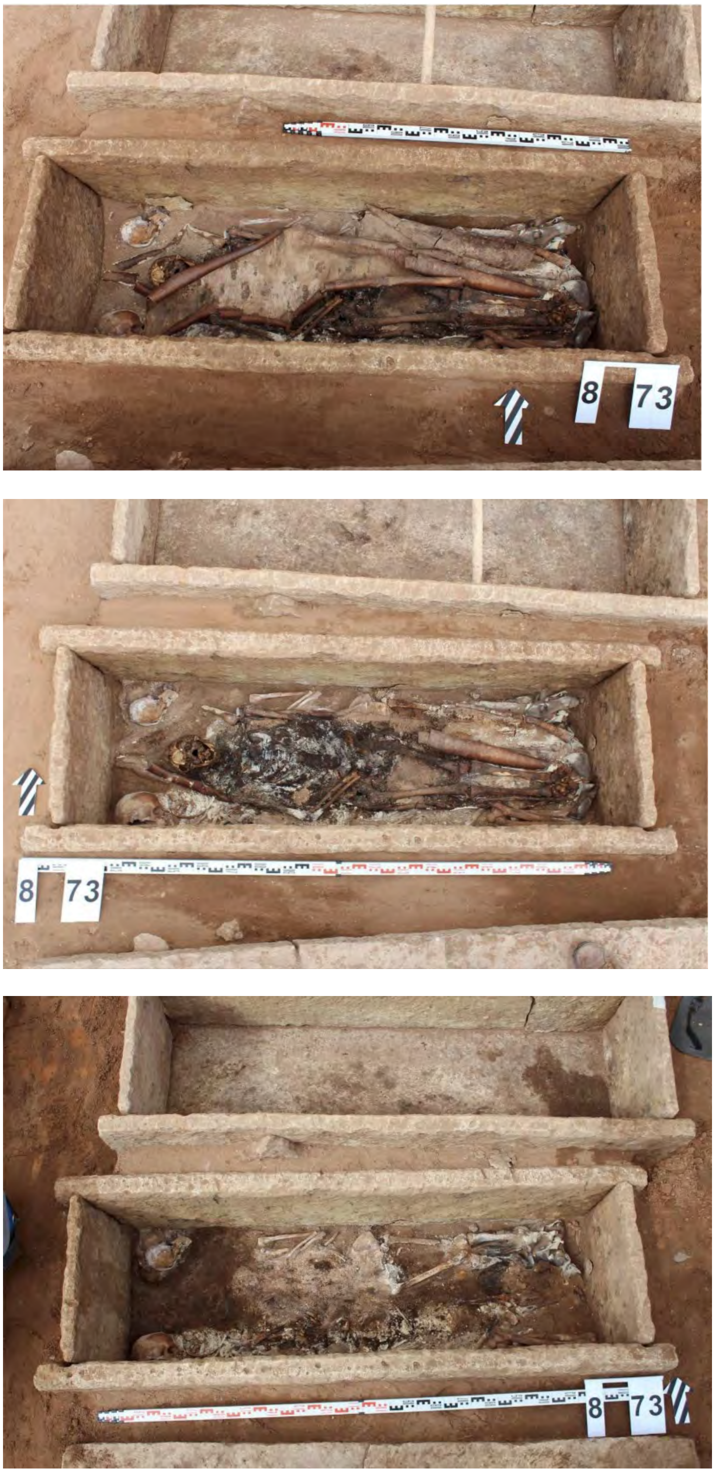


**Supplementary Fig. 24. The necropolis of the St. George (Yuriev) Monastery. Burial 73. Sarcophagus 8.**
Top view of the upper level of birch bark (top), top view of the level with partial removal in the eastern part of the sarcophagus (central), top view of the lower level (bottom). Photo provided by V.V. Sedov.

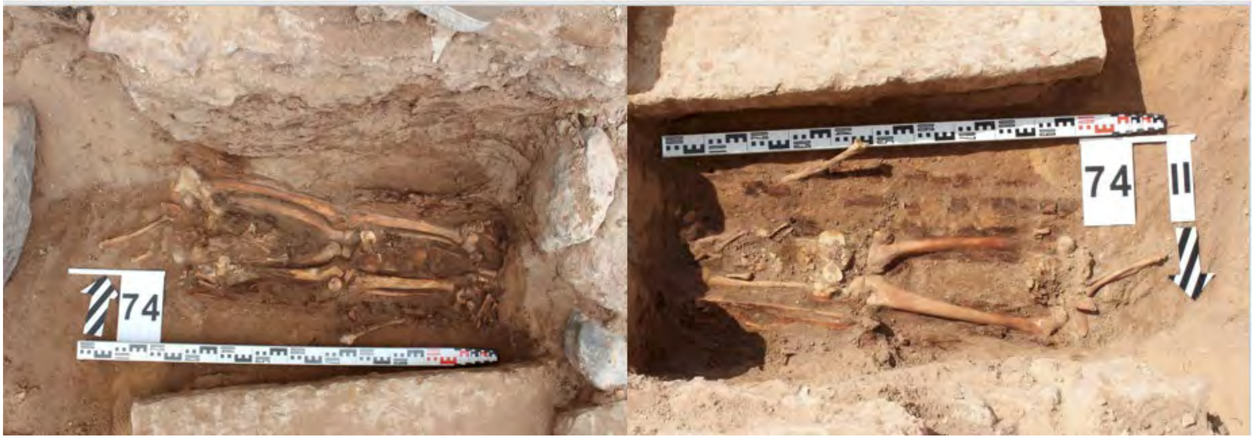


**Supplementary Fig. 25. The necropolis of the St. George (Yuriev) Monastery. Burial 74.** Ground-level. View of the upper level (left), view of lower level (right). Photo provided by V.V. Sedov.

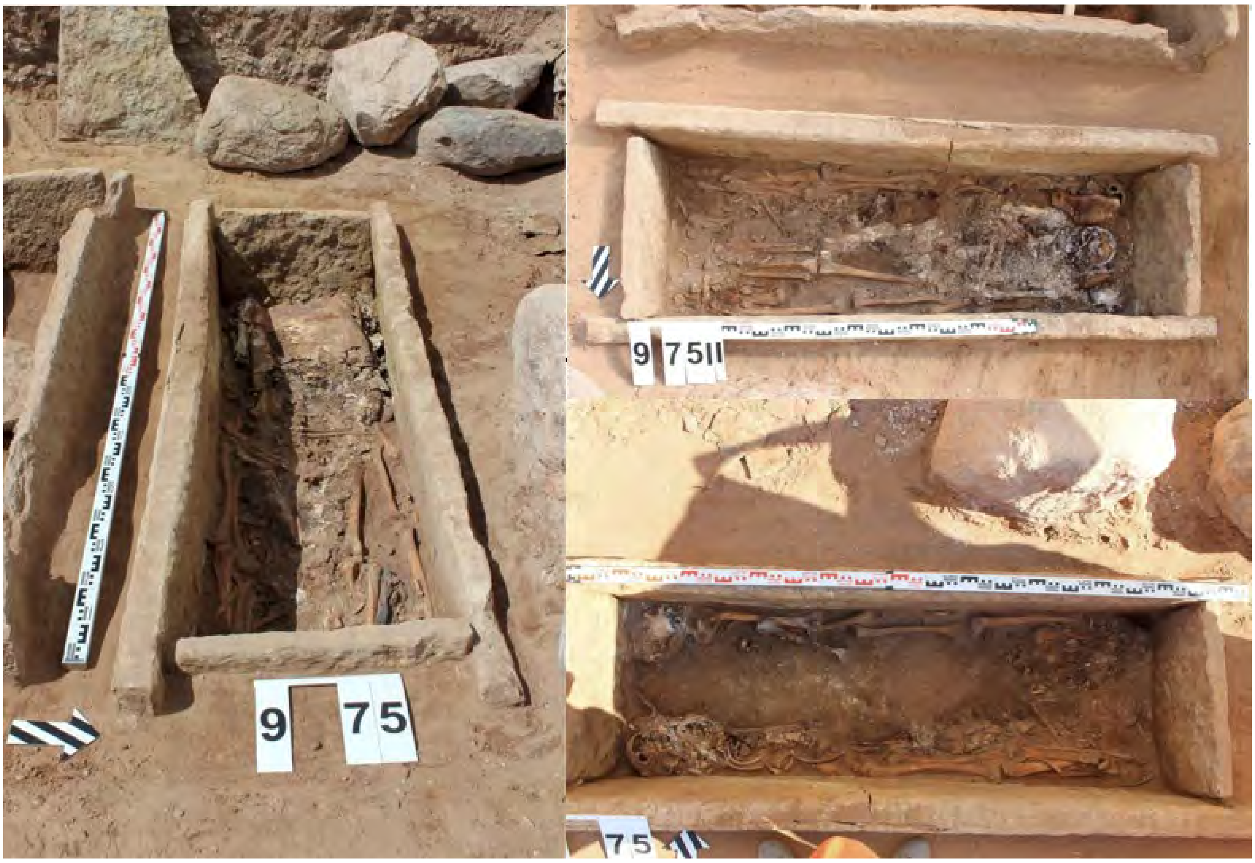


**Supplementary Fig. 26. The necropolis of the St. George (Yuriev) Monastery. Burial 75. Sarcophagus 9.**

View from the east to the upper level (left), view of the level after removing the birch bark (right top), view of the lower level (right below). Photo provided by V.V. Sedov.

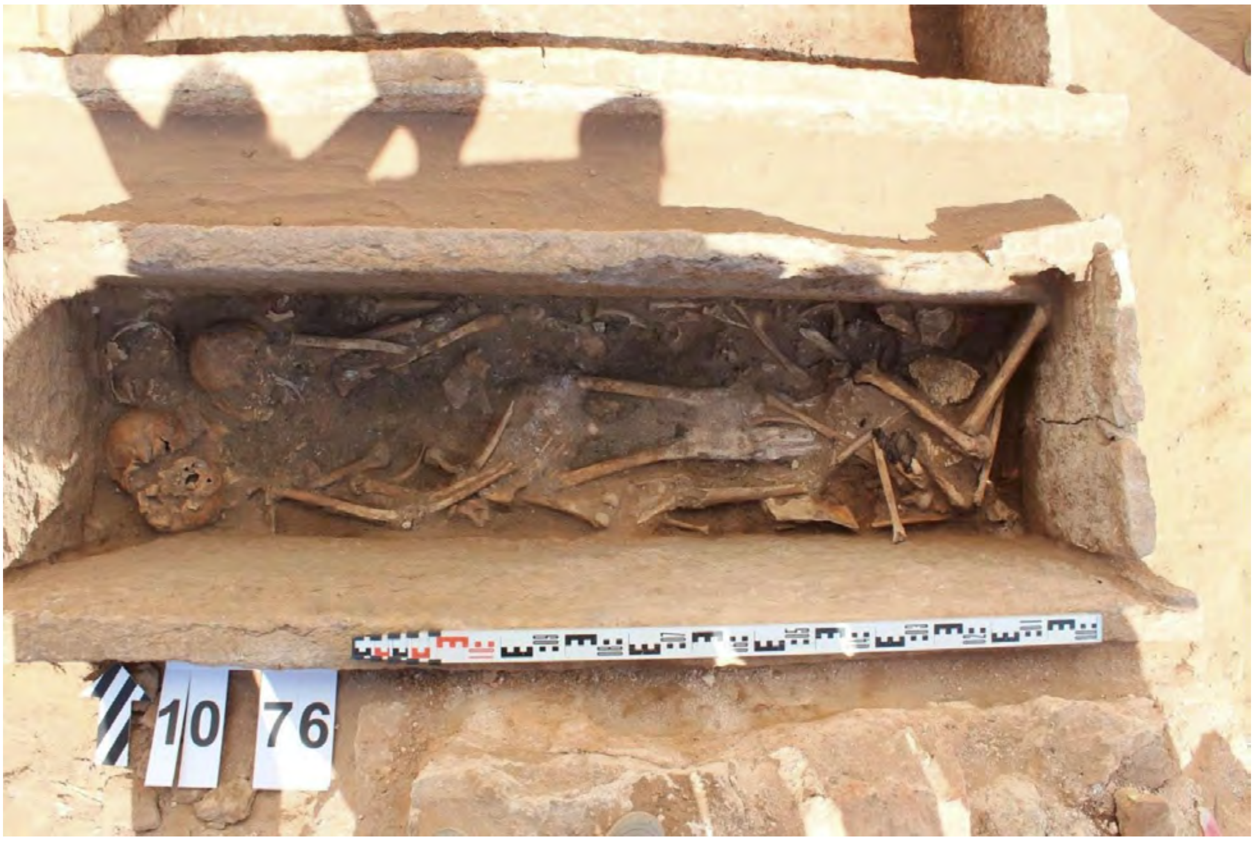


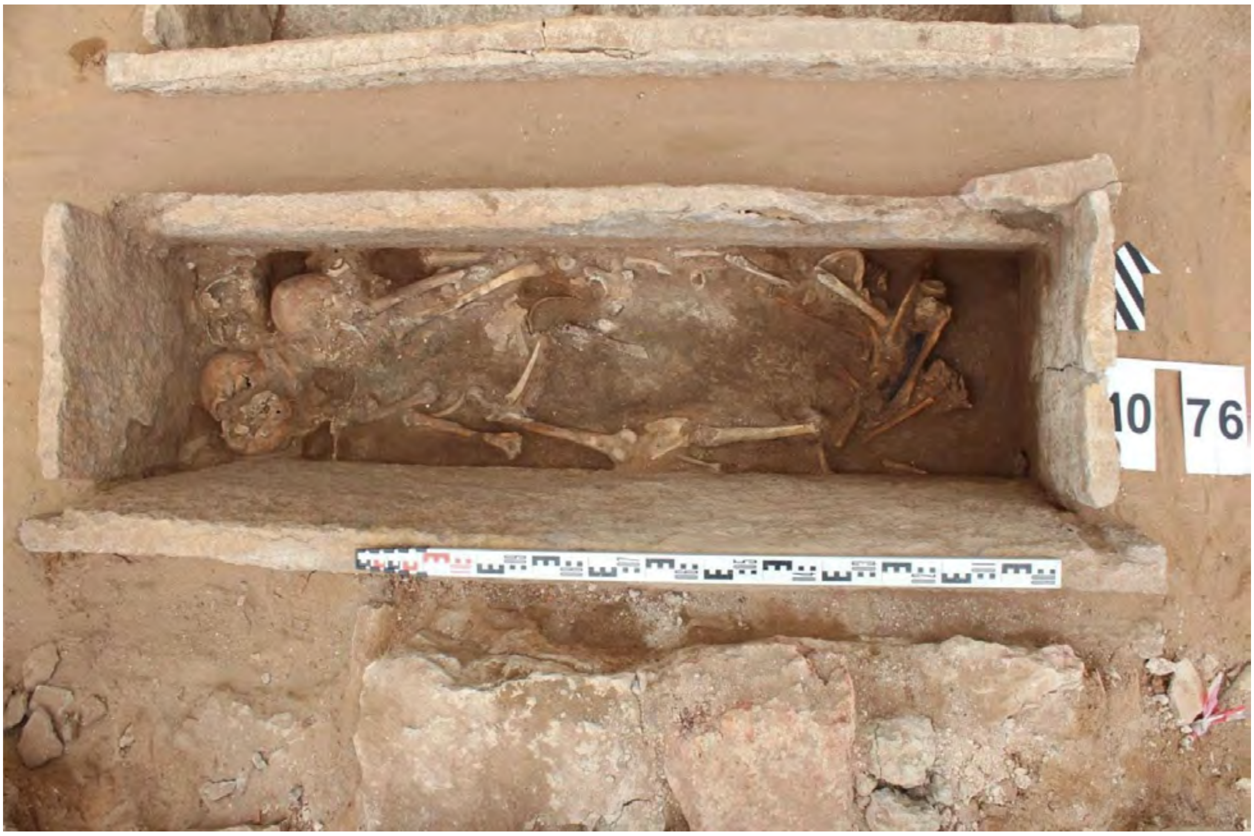


**Supplementary Fig. 27. The necropolis of the St. George (Yuriev) Monastery. Burial 76.**

View of the upper level (top), view of the lower level (below). Photo provided by V.V. Sedov.

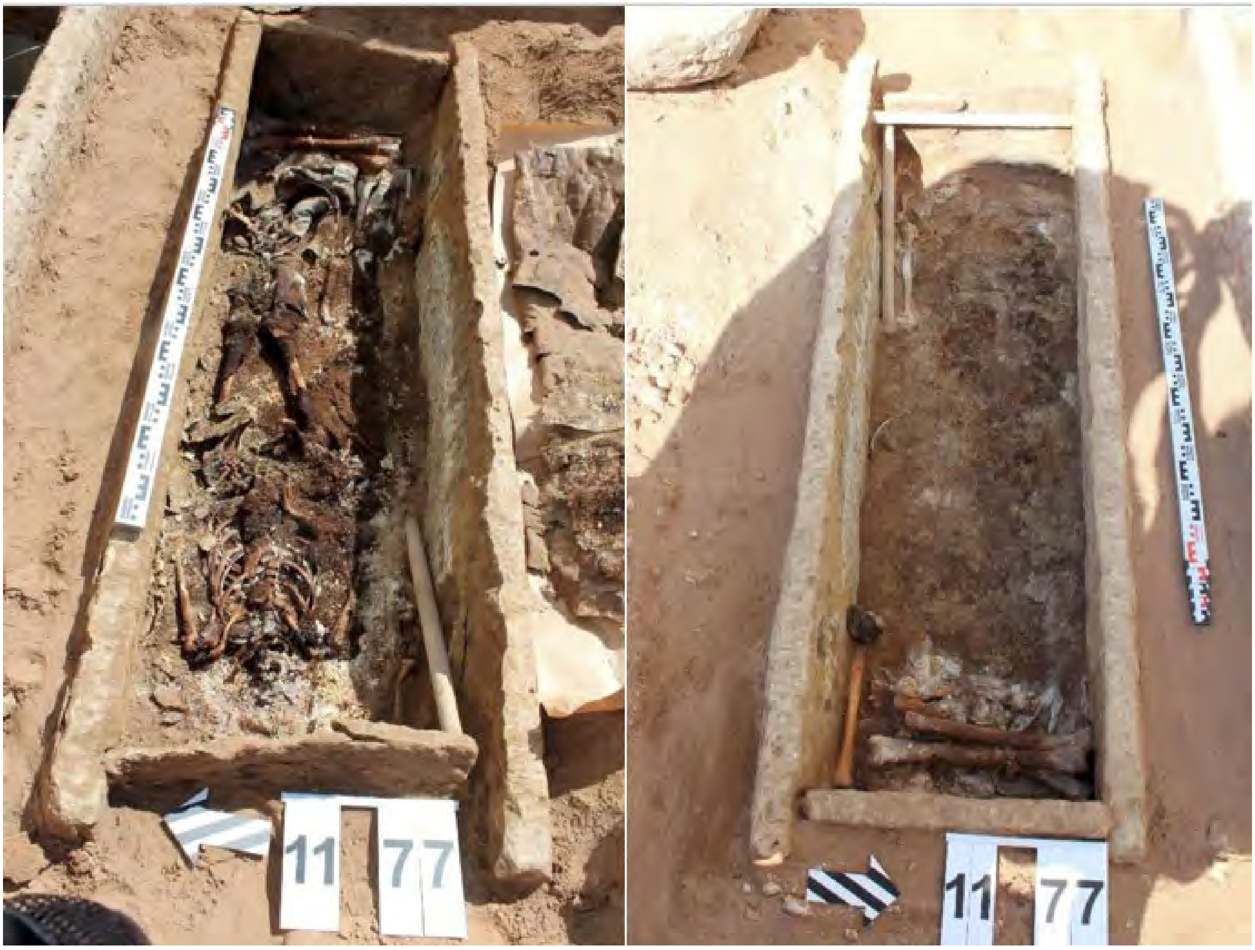


**Supplementary Fig. 28. The necropolis of the St. George (Yuriev) Monastery. Burial 77.**

View from the west to the upper level (left), view from the east to the lower level (right). Photo provided by V.V. Sedov.

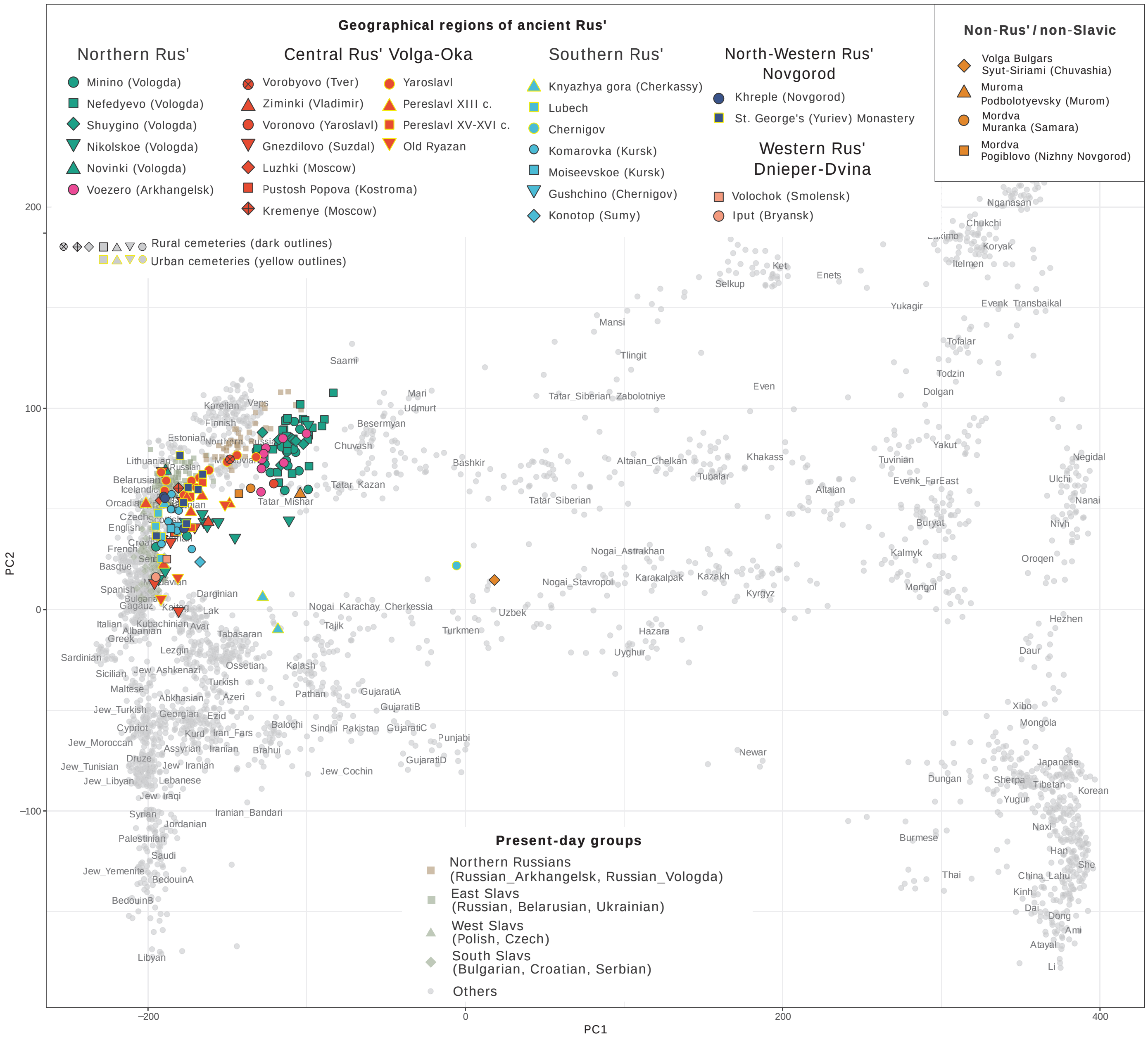

Geographical regions of ancient Rus'
Northern Rus'
Minino (Vologda)
Nefedyevo (Vologda)
Shuygino (Vologda)
Nikolskoe (Vologda)
Novinki (Vologda)
Voezero (Arkhangelsk)
Central Rus' Volga-Oka
Vorobyovo (Tver)
Ziminki (Vladimir)
Voronovo (Yaroslavl)
Gnezdilovo (Suzdal)
Luzhki (Moscow)
Pustosh Popova (Kostroma)
Kremenye (Moscow)
Yaroslavl
Pereslavl XIII c.
Pereslavl XV-XVI c.
Old Ryazan
Southern Rus'
Knyazhya gora (Cherkassy)
Lubech
Chernigov
Komarovka (Kursk)
Moiseevskoe (Kursk)
Gushchino (Chernigov)
Konotop (Sumy)
North-Western Rus' Novgorod
Khreple (Novgorod)
St. George's (Yuriev) Monastery
Western Rus' Dnieper-Dvina
Volochok (Smolensk)
Iput (Bryansk)
Non-Rus' / non-Slavic
Volga Bulgars Syut-Siriami (Chuvashia)
Muroma Podbolotyevsky (Murom)
Mordva Muranka (Samara)
Mordva Pogiblovo (Nizhny Novgorod)
Rural cemeteries (dark outlines)
Urban cemeteries (yellow outlines)
Present-day groups
Northern Russians (Russian_Arkhangelsk, Russian_Vologda)
East Slavs (Russian, Belarusian, Ukrainian)
West Slavs (Polish, Czech)
South Slavs (Bulgarian, Croatian, Serbian)
Others
PC1
PC2
200
100
0
−100
−200
0
200
400

**Supplementary Figure 29. Principal component analysis (PCA).**

PCA plot visualizing 144 newly reported samples from different geographical regions of ancient Rus' and neighboring contemporary tribes. Ancient individuals are projected on present-day Eurasians individuals of the Human Origin dataset supplemented by present-day Slavs from the Simons Genome Diversity Project[159], Finnish from the 1000 Genome Project[160] and Serbians[107] (grey points). This present-day set includes Northern Russians, East, West, and South Slavs (grayish squares, triangles, and diamonds combined by colored polygons, see legend for Present-day groups). The labels for the ancient individuals correspond to their burial sites, indicating the site name and/or neighboring town (see also Fig. 1, Fig. 2). Ancient individuals' colors are as follows (see also legend on the top): green and dark-pink, samples from Northern Rus' region; red, samples from Central Rus' region, light blue, samples from Southern Rus' region, dark-blue, samples from North-Western Rus' Novgorod; peach, samples from Western Rus'; orange, samples from Volga Bulgars, Mordva, and Muroma burials. Samples from urban cemeteries have yellow outlines, samples from rural cemeteries have dark outlines.

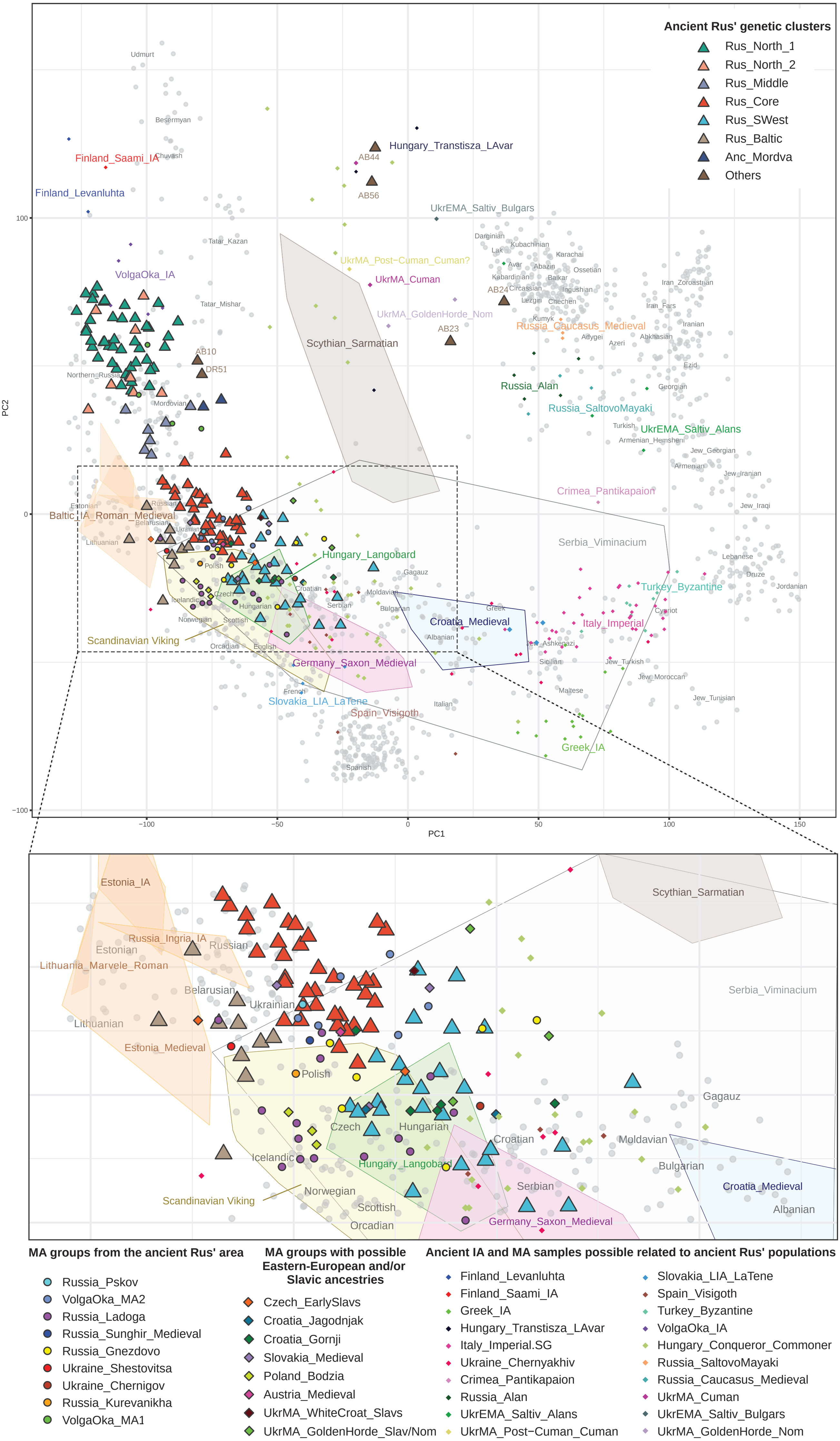
Ancient Rus' genetic clusters
Rus_North_1
Rus_North_2
Rus_Middle
Rus_Core
Rus_SWest
Rus_Baltic
Anc_Mordva
Others
PC2
PC1
100
0
−100
−100
−50
0
50
100
150
Udmurt
Besermyan
Chuvash
Finland_Saami_IA
Finland_Levanluhta
Tatar_Kazan
VolgaOka_IA
Tatar_Mishar
AB10
DR51
Northern_Russia
Mordovian
Hungary_Transtisza_LAvar
AB44
AB56
UkrEMA_Saltiv_Bulgars
UkrMA_Post−Cuman_Cuman?
UkrMA_Cuman
UkrMA_GoldenHorde_Nom
AB23
AB24
Scythian_Sarmatian
Russia_Caucasus_Medieval
Russia_Alan
Russia_SaltovoMayaki
UkrEMA_Saltiv_Alans
Crimea_Pantikapaion
Serbia_Viminacium
Hungary_Langobard
Turkey_Byzantine
Italy_Imperial
Croatia_Medieval
Germany_Saxon_Medieval
Slovakia_LIA_LaTene
Spain_Visigoth
Greek_IA
Scandinavian Viking
Baltic_IA_Roman_Medieval
Estonian
Russian
Belarusian
Ukrainian
Lithuanian
Polish
Czech
Icelandic
Hungarian
Croatian
Serbian
Moldavian
Bulgarian
Gagauz
Norwegian
Scottish
Orcadian
English
French
Italian
Spanish
Albanian
Greek
Sicilian
Maltese
Jew_Ashkenazi
Jew_Turkish
Jew_Moroccan
Jew_Tunisian
Cypriot
Lebanese
Druze
Jordanian
Jew_Iraqi
Jew_Iranian
Armenian
Jew_Georgian
Armenian_Hemsheni
Turkish
Georgian
Ezid
Iranian
Iran_Fars
Iran_Zoroastrian
Abkhasian
Azeri
Adygei
Kumyk
Lezgin
Chechen
Circassian
Ingushian
Kabardinian
Balkar
Avar
Abazin
Ossetian
Lak
Kubachinian
Karachai
Darginian
Estonia_IA
Russia_Ingria_IA
Lithuania_Marvele_Roman
Estonia_Medieval
MA groups from the ancient Rus' area
Russia_Pskov
VolgaOka_MA2
Russia_Ladoga
Russia_Sunghir_Medieval
Russia_Gnezdovo
Ukraine_Shestovitsa
Ukraine_Chernigov
Russia_Kurevanikha
VolgaOka_MA1
MA groups with possible Eastern-European and/or Slavic ancestries
Czech_EarlySlavs
Croatia_Jagodnjak
Croatia_Gornji
Slovakia_Medieval
Poland_Bodzia
Austria_Medieval
UkrMA_WhiteCroat_Slavs
UkrMA_GoldenHorde_Slav/Nom
Ancient IA and MA samples possible related to ancient Rus' populations
Finland_Levanluhta
Finland_Saami_IA
Greek_IA
Hungary_Transtisza_LAvar
Italy_Imperial.SG
Ukraine_Chernyakhiv
Crimea_Pantikapaion
Russia_Alan
UkrEMA_Saltiv_Alans
UkrMA_Post−Cuman_Cuman
Slovakia_LIA_LaTene
Spain_Visigoth
Turkey_Byzantine
VolgaOka_IA
Hungary_Conqueror_Commoner
Russia_SaltovoMayaki
Russia_Caucasus_Medieval
UkrMA_Cuman
UkrEMA_Saltiv_Bulgars
UkrMA_GoldenHorde_Nom

**Supplementary Fig. 30. Principal component analysis of Rus' individuals with previously published medieval samples.**
PCA analysis results of ancient individuals projected on present-day West-Eurasians (grey points) from the Human Origin dataset supplemented by present-day Slavs from the Simons Genome Diversity Project[159], Finnish from the 1000 Genome Project[160] and Serbians[107] (grey points). Ancient Rus' individuals (triangles) subdiveded into genetic clusters: Rus_North_1, Rus_North_2, Rus_Middle, Rus_Core, Rus_Baltic, Rus_SWest, Anc_Rus and outliers. Previously published ancient samples are presented by coloured diamonds (geographically and genetically appropriated populations), and by coloured points (samples from Ancient Rus' territory). To improve readability, following previously published ancient populations have been presented as colored polygons included all individuals in each population: brown, Scythian_Sarmatian (Kazakhstan_Sarmatian_IA, Russia_Sarmatian,Ukraine_IA_WesternScythian, Moldova_Glinoe_Scythian); orange, Baltic_IA_Roman_Medieval (Estonia_IA, Russia_Ingria_IA, Lithuania_Marvele_Roman); yellow, Scandinavian_vikings (Denmark_IA, Norway_IA, Sweden_IA, Estonia_Salme_EarlyViking, Denmark_Sealand_EarlyViking, Norway_Viking, Iceland_Viking); pink, Germany_Saxon_Medieval (Germany_Anderten_Saxon_Medieval, Germany_Drantum_Saxon_Medieval, Germany_Dunum_Saxon_Medieval); green, Hungary_Langobard; blue, Croatia_Medieval (Croatia_Medieval, Croatia_Nustar, Croatia_Sipar, Croatia_Gardun); light-grey, Serbia_Viminacium (Serbia_Viminacium_Roman_elite, Serbia_TimacumSlog, Serbia_Viminacium_Anat, Serbia_ViminaciumPecine, Serbia_ViminaciumPirivoj).

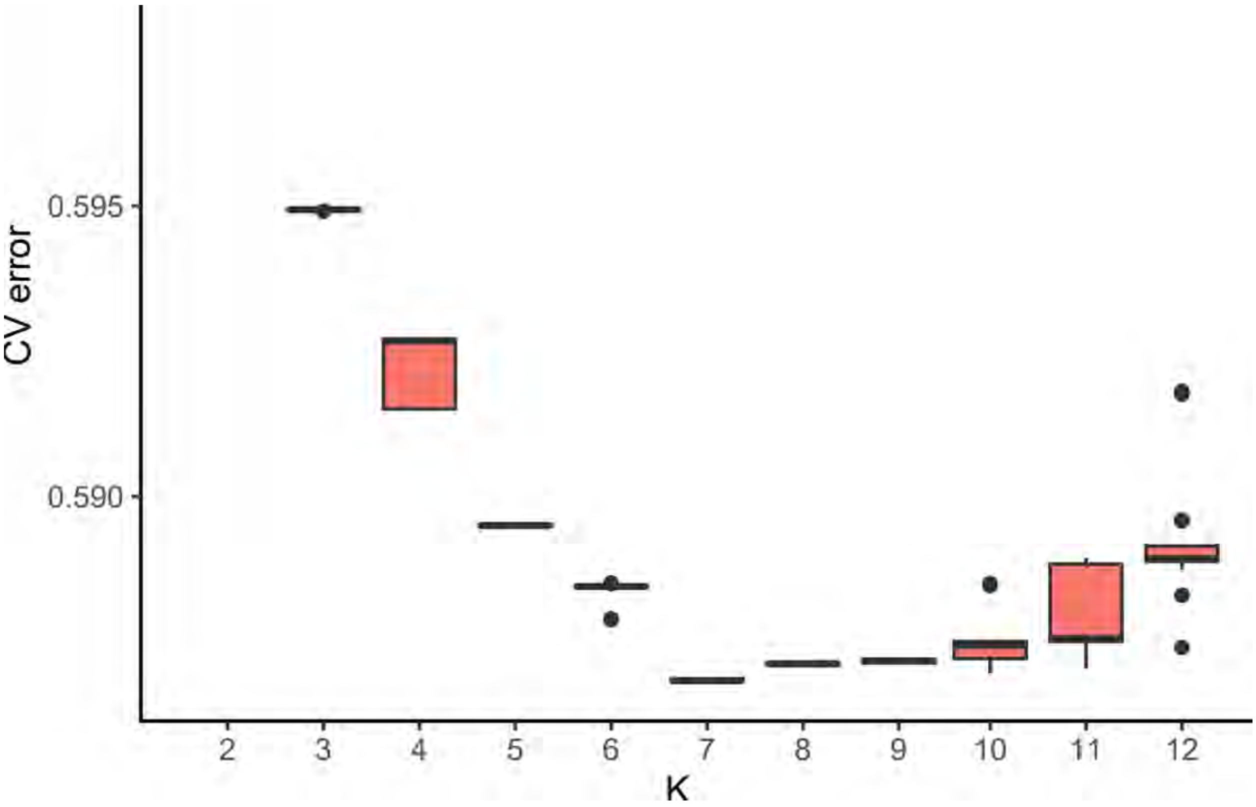


**Supplementary Fig. 31. Estimation of cross validation error for 20 runs of ADMIXTURE from K=2 through K=12.**
We chose K=7, as the value that minimizes the error.

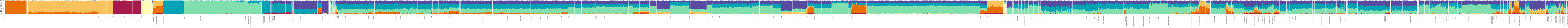

**Supplementary Fig. 32. ADMIXTURE analysis results for newly sequenced and previously published individuals (*K* = 8).**
See Supplementary Table 6 for a list of used samples.

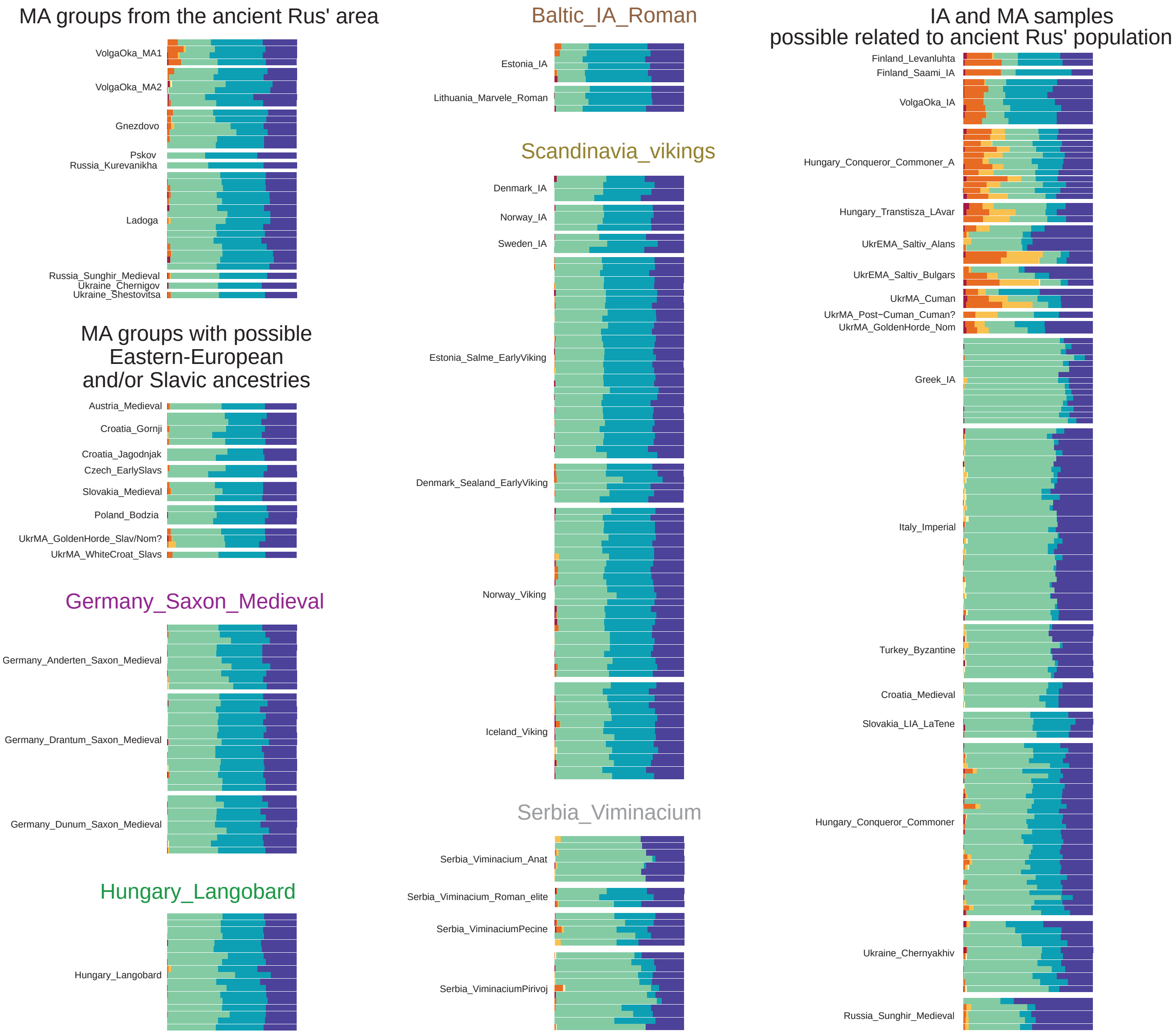
MA groups from the ancient Rus' area
VolgaOka_MA1
VolgaOka_MA2
Gnezdovo
Pskov
Russia_Kurevanikha
Ladoga
Russia_Sunghir_Medieval
Ukraine_Chernigov
Ukraine_Shestovitsa
MA groups with possible Eastern-European and/or Slavic ancestries
Austria_Medieval
Croatia_Gornji
Croatia_Jagodnjak
Czech_EarlySlavs
Slovakia_Medieval
Poland_Bodzia
UkrMA_GoldenHorde_Slav/Nom?
UkrMA_WhiteCroat_Slavs
Germany_Saxon_Medieval
Germany_Anderten_Saxon_Medieval
Germany_Drantum_Saxon_Medieval
Germany_Dunum_Saxon_Medieval
Hungary_Langobard
Hungary_Langobard
Baltic_IA_Roman
Estonia_IA
Lithuania_Marvele_Roman
Scandinavia_vikings
Denmark_IA
Norway_IA
Sweden_IA
Estonia_Salme_EarlyViking
Denmark_Sealand_EarlyViking
Norway_Viking
Iceland_Viking
Serbia_Viminacium
Serbia_Viminacium_Anat
Serbia_Viminacium_Roman_elite
Serbia_ViminaciumPecine
Serbia_ViminaciumPirivoj
IA and MA samples possible related to ancient Rus' population
Finland_Levanluhta
Finland_Saami_IA
VolgaOka_IA
Hungary_Conqueror_Commoner_A
Hungary_Transtisza_LAvar
UkrEMA_Saltiv_Alans
UkrEMA_Saltiv_Bulgars
UkrMA_Cuman
UkrMA_Post−Cuman_Cuman?
UkrMA_GoldenHorde_Nom
Greek_IA
Italy_Imperial
Turkey_Byzantine
Croatia_Medieval
Slovakia_LIA_LaTene
Hungary_Conqueror_Commoner
Ukraine_Chernyakhiv
Russia_Sunghir_Medieval

**Supplementary Fig. 33. Unsupervised ADMIXTURE results at K = 7 for previously published ancient samples from Rus' area and neighboring regions.**
See also Fig. 4, Supplementary Fig. 30, and Supplementary Fig. 32.

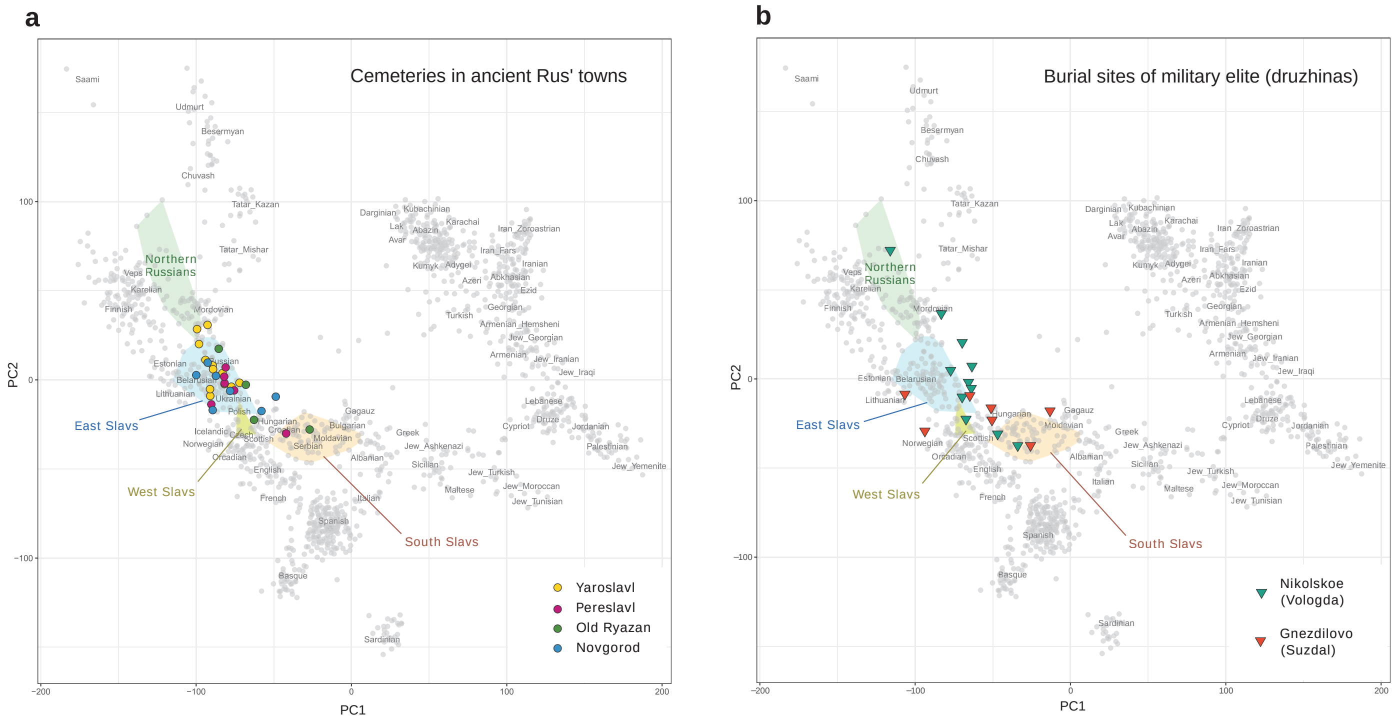
a
Cemeteries in ancient Rus' towns
Northern Russians
East Slavs
West Slavs
South Slavs
Yaroslavl
Pereslavl
Old Ryazan
Novgorod
PC1
PC2
b
Burial sites of military elite (druzhinas)
Northern Russians
East Slavs
West Slavs
South Slavs
Nikolskoe (Vologda)
Gnezdilovo (Suzdal)
PC1
PC2

**Supplementary Fig. 34. PCA analysis results of Rus' individuals from urban cemeteries (a) and barrows of presumable military elite (b) projected on present-day West-Eurasians (grey points) from the Human Origin dataset supplemented by present-day Slavs from the Simons Genome Diversity Project[159], Finnish from the 1000 Genome Project[160] and Serbians[107].**

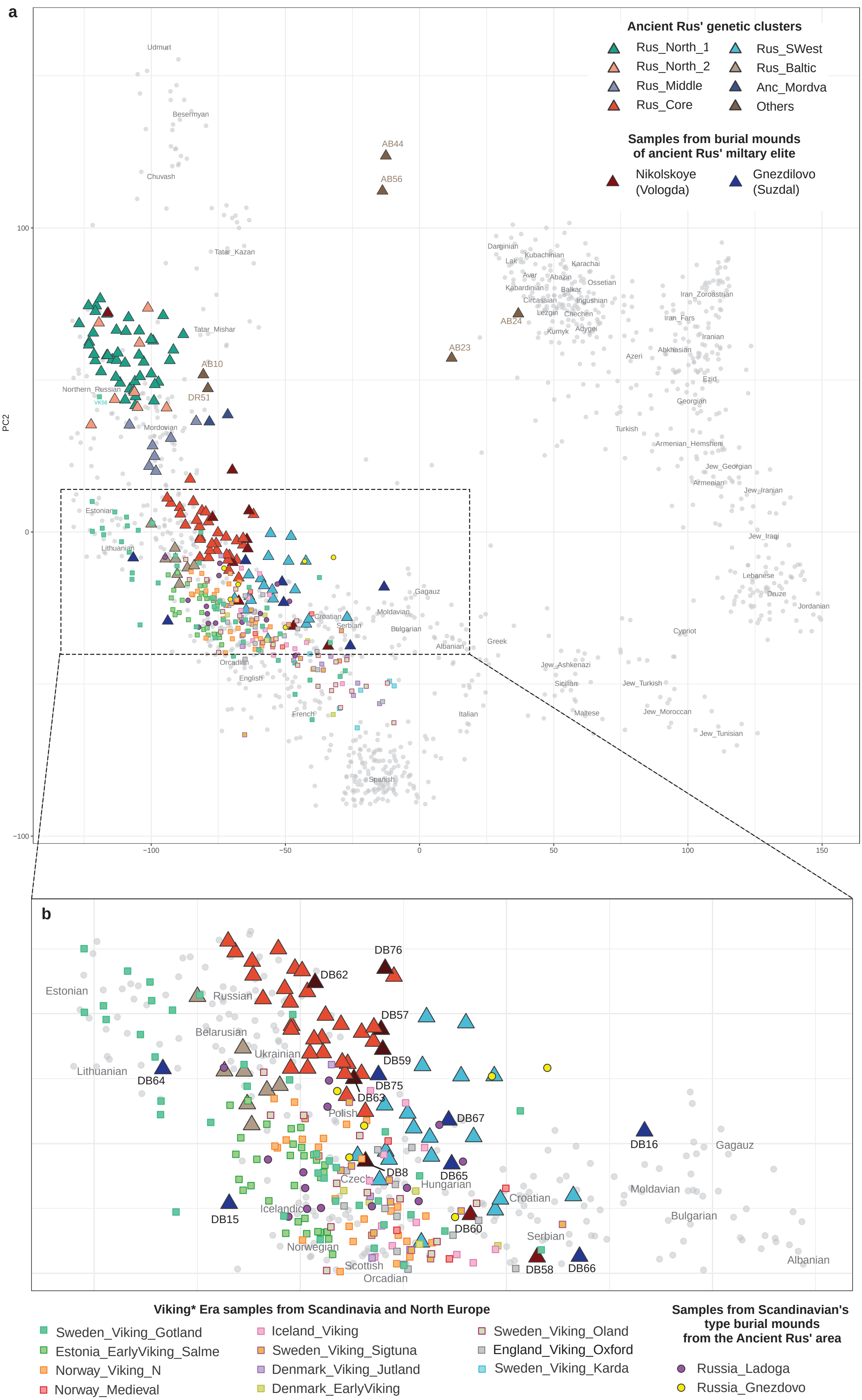
a
Ancient Rus' genetic clusters
Rus_North_1
Rus_North_2
Rus_Middle
Rus_Core
Rus_SWest
Rus_Baltic
Anc_Mordva
Others
Samples from burial mounds
of ancient Rus' miltary elite
Nikolskoye
(Vologda)
Gnezdilovo
(Suzdal)
PC2
100
0
−100
−100
−50
0
50
100
150
Udmurt
Besermyan
Chuvash
Tatar_Kazan
Tatar_Mishar
Northern_Russian
VK56
Mordovian
Estonian
Lithuanian
AB44
AB56
AB24
AB23
AB10
DR51
Darginian
Kubachinian
Lak
Karachai
Avar
Abazin
Ossetian
Kabardinian
Balkar
Circassian
Ingushian
Lezgin
Chechen
Kumyk
Adygei
Iran_Zoroastrian
Iran_Fars
Iranian
Azeri
Abkhasian
Ezid
Georgian
Turkish
Armenian_Hemsheni
Jew_Georgian
Armenian
Jew_Iranian
Jew_Iraqi
Lebanese
Druze
Jordanian
Cypriot
Gagauz
Moldavian
Bulgarian
Croatian
Serbian
Albanian
Greek
Orcadian
English
French
Italian
Spanish
Jew_Ashkenazi
Sicilian
Maltese
Jew_Turkish
Jew_Moroccan
Jew_Tunisian
b
Estonian
Russian
Belarusian
Ukrainian
Lithuanian
Polish
Czech
Hungarian
Icelandic
Norwegian
Scottish
Orcadian
Croatian
Serbian
Moldavian
Bulgarian
Gagauz
Albanian
DB76
DB62
DB57
DB59
DB75
DB63
DB64
DB67
DB16
DB8
DB65
DB15
DB60
DB58
DB66
Viking* Era samples from Scandinavia and North Europe
Sweden_Viking_Gotland
Estonia_EarlyViking_Salme
Norway_Viking_N
Norway_Medieval
Iceland_Viking
Sweden_Viking_Sigtuna
Denmark_Viking_Jutland
Denmark_EarlyViking
Sweden_Viking_Oland
England_Viking_Oxford
Sweden_Viking_Karda
Samples from Scandinavian's
type burial mounds
from the Ancient Rus' area
Russia_Ladoga
Russia_Gnezdovo

**Supplementary Fig. 35. PCA analysis results of ancient Rus’ and Viking Era individuals projected on present-day West-Eurasians (grey points) from the Human Origin dataset supplemented by present-day Slavs from the Simons Genome Diversity Project[159], Finnish from the 1000 Genome Project[160] and Serbians[107].**
Ancient Rus’ individuals (triangles) subdivided into genetic clusters: Rus_North_1, Rus_North_2, Rus_Middle, Rus_Core, Rus_Baltic, Rus_SWest, Anc_Rus and outliers. Scandinavian Vikings and Viking Era individuals from Northern Europe are presented by coloured squares. Samples from Scandinavian's type barrows from the Rus' area are represented by colored circles. VKE - Viking Era.

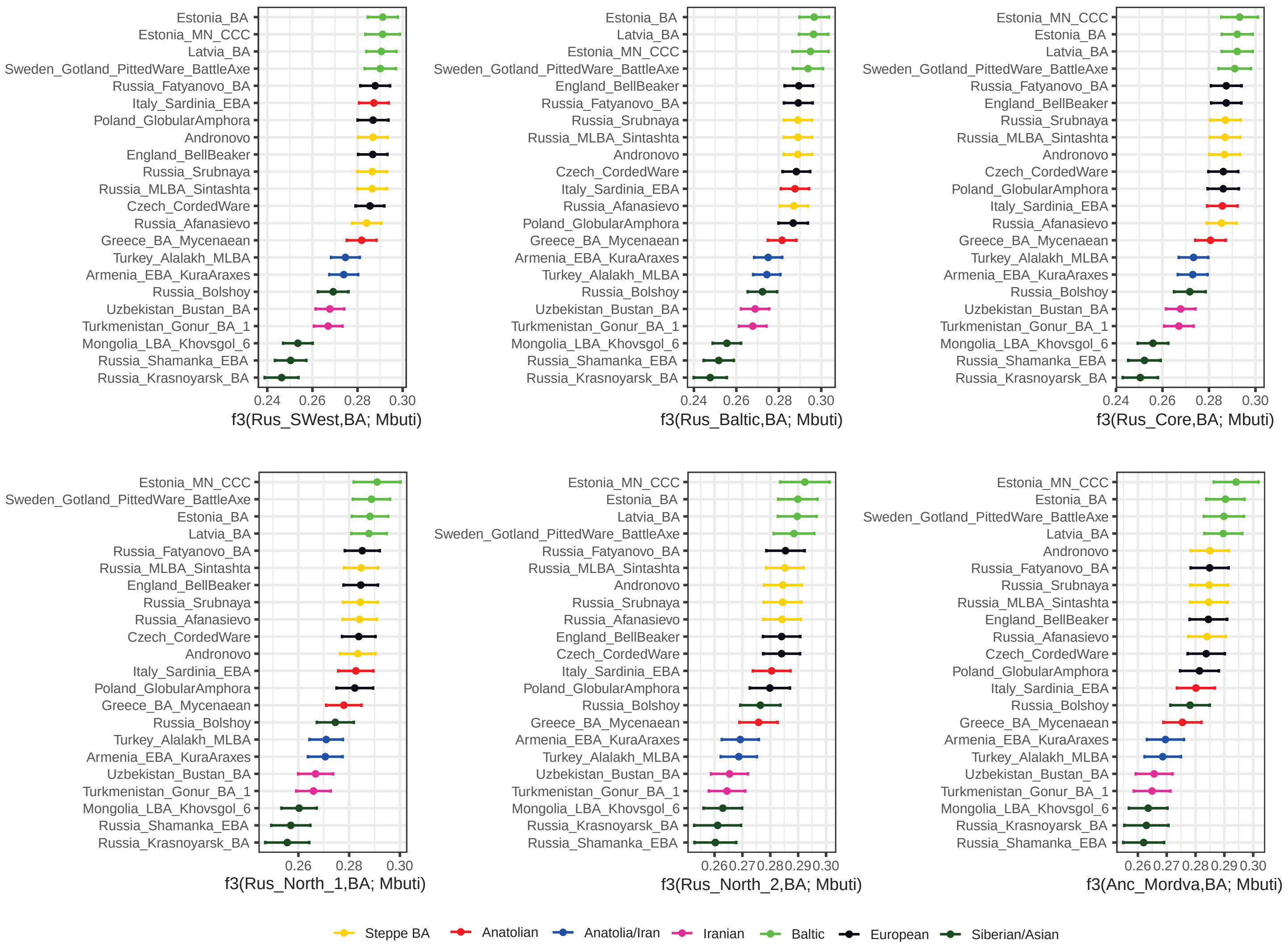
Estonia_BA
Estonia_MN_CCC
Latvia_BA
Sweden_Gotland_PittedWare_BattleAxe
Russia_Fatyanovo_BA
Italy_Sardinia_EBA
Poland_GlobularAmphora
Andronovo
England_BellBeaker
Russia_Srubnaya
Russia_MLBA_Sintashta
Czech_CordedWare
Russia_Afanasievo
Greece_BA_Mycenaean
Turkey_Alalakh_MLBA
Armenia_EBA_KuraAraxes
Russia_Bolshoy
Uzbekistan_Bustan_BA
Turkmenistan_Gonur_BA_1
Mongolia_LBA_Khovsgol_6
Russia_Shamanka_EBA
Russia_Krasnoyarsk_BA
0.24 0.26 0.28 0.30
f3(Rus_SWest,BA; Mbuti)
f3(Rus_Baltic,BA; Mbuti)
f3(Rus_Core,BA; Mbuti)
f3(Rus_North_1,BA; Mbuti)
0.26 0.27 0.28 0.29 0.30
f3(Rus_North_2,BA; Mbuti)
f3(Anc_Mordva,BA; Mbuti)
Steppe BA
Anatolian
Anatolia/Iran
Iranian
Baltic
European
Siberian/Asian

**Supplementary Fig. 36. Outgroups f3-statistic results of form *f3*(Rus’,BA; Mbuti).**

BA - BA groups, Rus’ - genetic subclusters Rus_Core, Rus_Baltic, Rus_SWest, Rus_North_1, Rus_North_2, and Anc_Mordva.

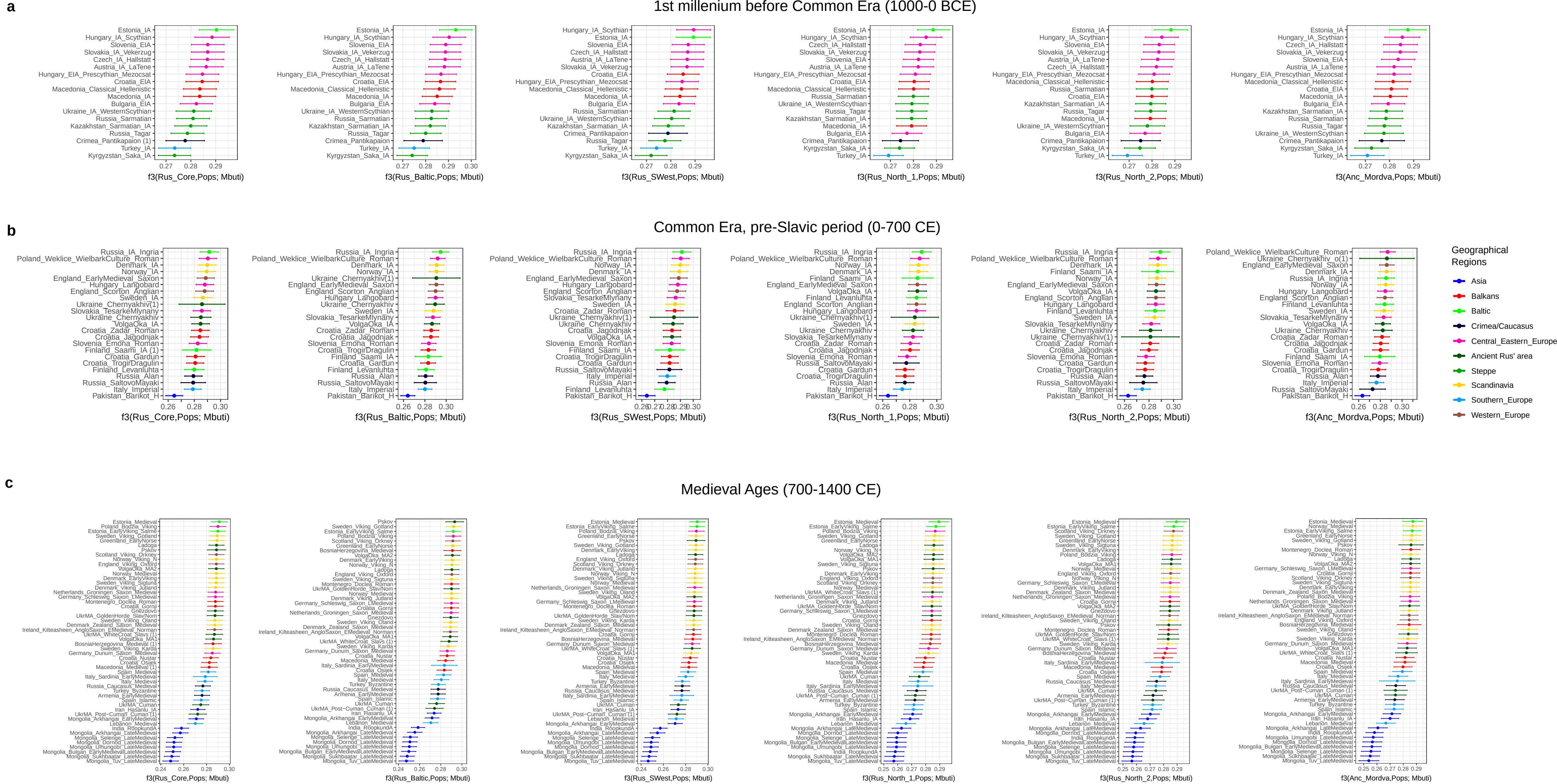
a
1st millenium before Common Era (1000-0 BCE)
f3(Rus_Core,Pops; Mbuti)
f3(Rus_Baltic,Pops; Mbuti)
f3(Rus_SWest,Pops; Mbuti)
f3(Rus_North_1,Pops; Mbuti)
f3(Rus_North_2,Pops; Mbuti)
f3(Anc_Mordva,Pops; Mbuti)
b
Common Era, pre-Slavic period (0-700 CE)
f3(Rus_Core,Pops; Mbuti)
f3(Rus_Baltic,Pops; Mbuti)
f3(Rus_SWest,Pops; Mbuti)
f3(Rus_North_1,Pops; Mbuti)
f3(Rus_North_2,Pops; Mbuti)
f3(Anc_Mordva,Pops; Mbuti)
Geographical Regions
Asia
Balkans
Baltic
Crimea/Caucasus
Central_Eastern_Europe
Ancient Rus' area
Steppe
Scandinavia
Southern_Europe
Western_Europe
c
Medieval Ages (700-1400 CE)
f3(Rus_Core,Pops; Mbuti)
f3(Rus_Baltic,Pops; Mbuti)
f3(Rus_SWest,Pops; Mbuti)
f3(Rus_North_1,Pops; Mbuti)
f3(Rus_North_2,Pops; Mbuti)
f3(Anc_Mordva,Pops; Mbuti)

**Supplementary Fig. 37. Outgroups *f*3-statistic results of form *f3*(Rus’,Pops; Mbuti).**

Pops - IA and MA groups, Rus’ - genetic subclusters Rus_Core, Rus_Baltic, Rus_SWest, Rus_North_1, Rus_North_2, and Anc_Mordva.

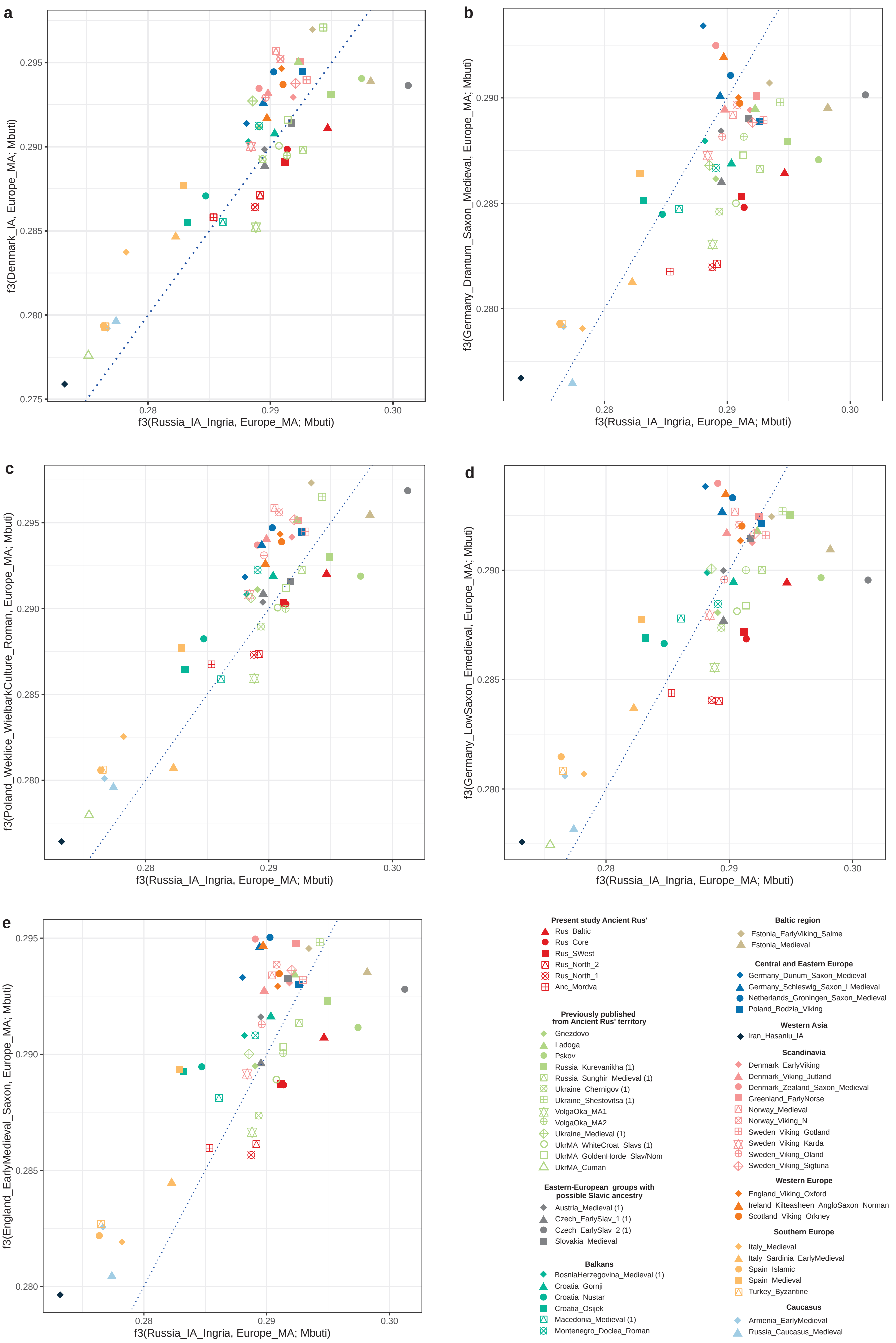

a
b
c
d
e
f3(Russia_IA_Ingria, Europe_MA; Mbuti)
f3(Denmark_IA, Europe_MA; Mbuti)
f3(Germany_Drantum_Saxon_Medieval, Europe_MA; Mbuti)
f3(Poland_Weklice_WielbarkCulture_Roman, Europe_MA; Mbuti)
f3(Germany_LowSaxon_Emedieval, Europe_MA; Mbuti)
f3(England_EarlyMedieval_Saxon, Europe_MA; Mbuti)
0.275
0.280
0.285
0.290
0.295
0.28
0.29
0.30
Present study Ancient Rus'
Rus_Baltic
Rus_Core
Rus_SWest
Rus_North_2
Rus_North_1
Anc_Mordva
Previously published from Ancient Rus' territory
Gnezdovo
Ladoga
Pskov
Russia_Kurevanikha (1)
Russia_Sunghir_Medieval (1)
Ukraine_Chernigov (1)
Ukraine_Shestovitsa (1)
VolgaOka_MA1
VolgaOka_MA2
Ukraine_Medieval (1)
UkrMA_WhiteCroat_Slavs (1)
UkrMA_GoldenHorde_Slav/Nom
UkrMA_Cuman
Eastern-European groups with possible Slavic ancestry
Austria_Medieval (1)
Czech_EarlySlav_1 (1)
Czech_EarlySlav_2 (1)
Slovakia_Medieval
Balkans
BosniaHerzegovina_Medieval (1)
Croatia_Gornji
Croatia_Nustar
Croatia_Osijek
Macedonia_Medieval (1)
Montenegro_Doclea_Roman
Baltic region
Estonia_EarlyViking_Salme
Estonia_Medieval
Central and Eastern Europe
Germany_Dunum_Saxon_Medieval
Germany_Schleswig_Saxon_LMedieval
Netherlands_Groningen_Saxon_Medieval
Poland_Bodzia_Viking
Western Asia
Iran_Hasanlu_IA
Scandinavia
Denmark_EarlyViking
Denmark_Viking_Jutland
Denmark_Zealand_Saxon_Medieval
Greenland_EarlyNorse
Norway_Medieval
Norway_Viking_N
Sweden_Viking_Gotland
Sweden_Viking_Karda
Sweden_Viking_Oland
Sweden_Viking_Sigtuna
Western Europe
England_Viking_Oxford
Ireland_Kilteasheen_AngloSaxon_Norman
Scotland_Viking_Orkney
Southern Europe
Italy_Medieval
Italy_Sardinia_EarlyMedieval
Spain_Islamic
Spain_Medieval
Turkey_Byzantine
Caucasus
Armenia_EarlyMedieval
Russia_Caucasus_Medieval

**Supplementary Fig. 38. Outgroup f3-statistic results of form *f*3(IA, MA; Mbuti) plotted against Baltic region IA group (Russia_IA_Ingria, X axis).**

European IA/MA groups (Y axis) are Denmark_IA (a), Germany_Drantum_Saxon_Medieval (b), Poland_Weklice_WielbarkCulture_Roman.SG (c), Germany_LowSaxon_Emedieval (d), England_EarlyMedieval_Saxon (e). The color and shape of the points are provided in the figure legends and correspond to tested populations. The number 1 is shown in parenthesis for a single-sample populations.

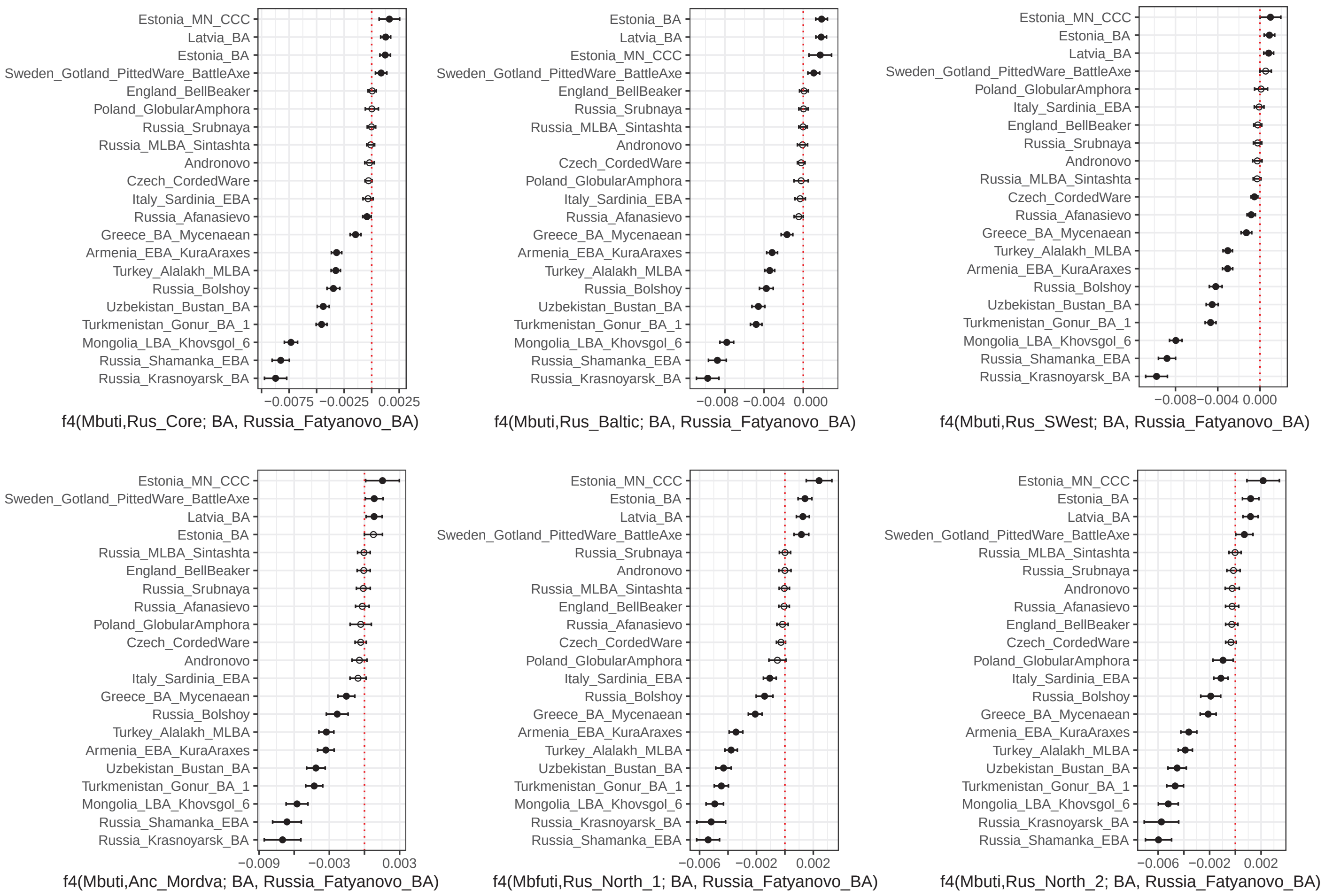
Estonia_MN_CCC
Latvia_BA
Estonia_BA
Sweden_Gotland_PittedWare_BattleAxe
England_BellBeaker
Poland_GlobularAmphora
Russia_Srubnaya
Russia_MLBA_Sintashta
Andronovo
Czech_CordedWare
Italy_Sardinia_EBA
Russia_Afanasievo
Greece_BA_Mycenaean
Armenia_EBA_KuraAraxes
Turkey_Alalakh_MLBA
Russia_Bolshoy
Uzbekistan_Bustan_BA
Turkmenistan_Gonur_BA_1
Mongolia_LBA_Khovsgol_6
Russia_Shamanka_EBA
Russia_Krasnoyarsk_BA
−0.0075
−0.0025
0.0025
f4(Mbuti,Rus_Core; BA, Russia_Fatyanovo_BA)
Estonia_BA
Latvia_BA
Estonia_MN_CCC
Sweden_Gotland_PittedWare_BattleAxe
England_BellBeaker
Russia_Srubnaya
Russia_MLBA_Sintashta
Andronovo
Czech_CordedWare
Poland_GlobularAmphora
Italy_Sardinia_EBA
Russia_Afanasievo
Greece_BA_Mycenaean
Armenia_EBA_KuraAraxes
Turkey_Alalakh_MLBA
Russia_Bolshoy
Uzbekistan_Bustan_BA
Turkmenistan_Gonur_BA_1
Mongolia_LBA_Khovsgol_6
Russia_Shamanka_EBA
Russia_Krasnoyarsk_BA
−0.008
−0.004
0.000
f4(Mbuti,Rus_Baltic; BA, Russia_Fatyanovo_BA)
Estonia_MN_CCC
Estonia_BA
Latvia_BA
Sweden_Gotland_PittedWare_BattleAxe
Poland_GlobularAmphora
Italy_Sardinia_EBA
England_BellBeaker
Russia_Srubnaya
Andronovo
Russia_MLBA_Sintashta
Czech_CordedWare
Russia_Afanasievo
Greece_BA_Mycenaean
Turkey_Alalakh_MLBA
Armenia_EBA_KuraAraxes
Russia_Bolshoy
Uzbekistan_Bustan_BA
Turkmenistan_Gonur_BA_1
Mongolia_LBA_Khovsgol_6
Russia_Shamanka_EBA
Russia_Krasnoyarsk_BA
−0.008
−0.004
0.000
f4(Mbuti,Rus_SWest; BA, Russia_Fatyanovo_BA)
Estonia_MN_CCC
Sweden_Gotland_PittedWare_BattleAxe
Latvia_BA
Estonia_BA
Russia_MLBA_Sintashta
England_BellBeaker
Russia_Srubnaya
Russia_Afanasievo
Poland_GlobularAmphora
Czech_CordedWare
Andronovo
Italy_Sardinia_EBA
Greece_BA_Mycenaean
Russia_Bolshoy
Turkey_Alalakh_MLBA
Armenia_EBA_KuraAraxes
Uzbekistan_Bustan_BA
Turkmenistan_Gonur_BA_1
Mongolia_LBA_Khovsgol_6
Russia_Shamanka_EBA
Russia_Krasnoyarsk_BA
−0.009
−0.003
0.003
f4(Mbuti,Anc_Mordva; BA, Russia_Fatyanovo_BA)
Estonia_MN_CCC
Estonia_BA
Latvia_BA
Sweden_Gotland_PittedWare_BattleAxe
Russia_Srubnaya
Andronovo
Russia_MLBA_Sintashta
England_BellBeaker
Russia_Afanasievo
Czech_CordedWare
Poland_GlobularAmphora
Italy_Sardinia_EBA
Russia_Bolshoy
Greece_BA_Mycenaean
Armenia_EBA_KuraAraxes
Turkey_Alalakh_MLBA
Uzbekistan_Bustan_BA
Turkmenistan_Gonur_BA_1
Mongolia_LBA_Khovsgol_6
Russia_Krasnoyarsk_BA
Russia_Shamanka_EBA
−0.006
−0.002
0.002
f4(Mbfuti,Rus_North_1; BA, Russia_Fatyanovo_BA)
Estonia_MN_CCC
Estonia_BA
Latvia_BA
Sweden_Gotland_PittedWare_BattleAxe
Russia_MLBA_Sintashta
Russia_Srubnaya
Andronovo
Russia_Afanasievo
England_BellBeaker
Czech_CordedWare
Poland_GlobularAmphora
Italy_Sardinia_EBA
Russia_Bolshoy
Greece_BA_Mycenaean
Armenia_EBA_KuraAraxes
Turkey_Alalakh_MLBA
Uzbekistan_Bustan_BA
Turkmenistan_Gonur_BA_1
Mongolia_LBA_Khovsgol_6
Russia_Krasnoyarsk_BA
Russia_Shamanka_EBA
−0.006
−0.002
0.002
f4(Mbuti,Rus_North_2; BA, Russia_Fatyanovo_BA)

**Supplementary Fig. 39. *f*4-statistic results of form *f4*(Mbuti,Rus’; BA, Russia_Fatyanovo_BA).**

BA - tested BA groups, Rus’ - Rus’ genetic subclusters.

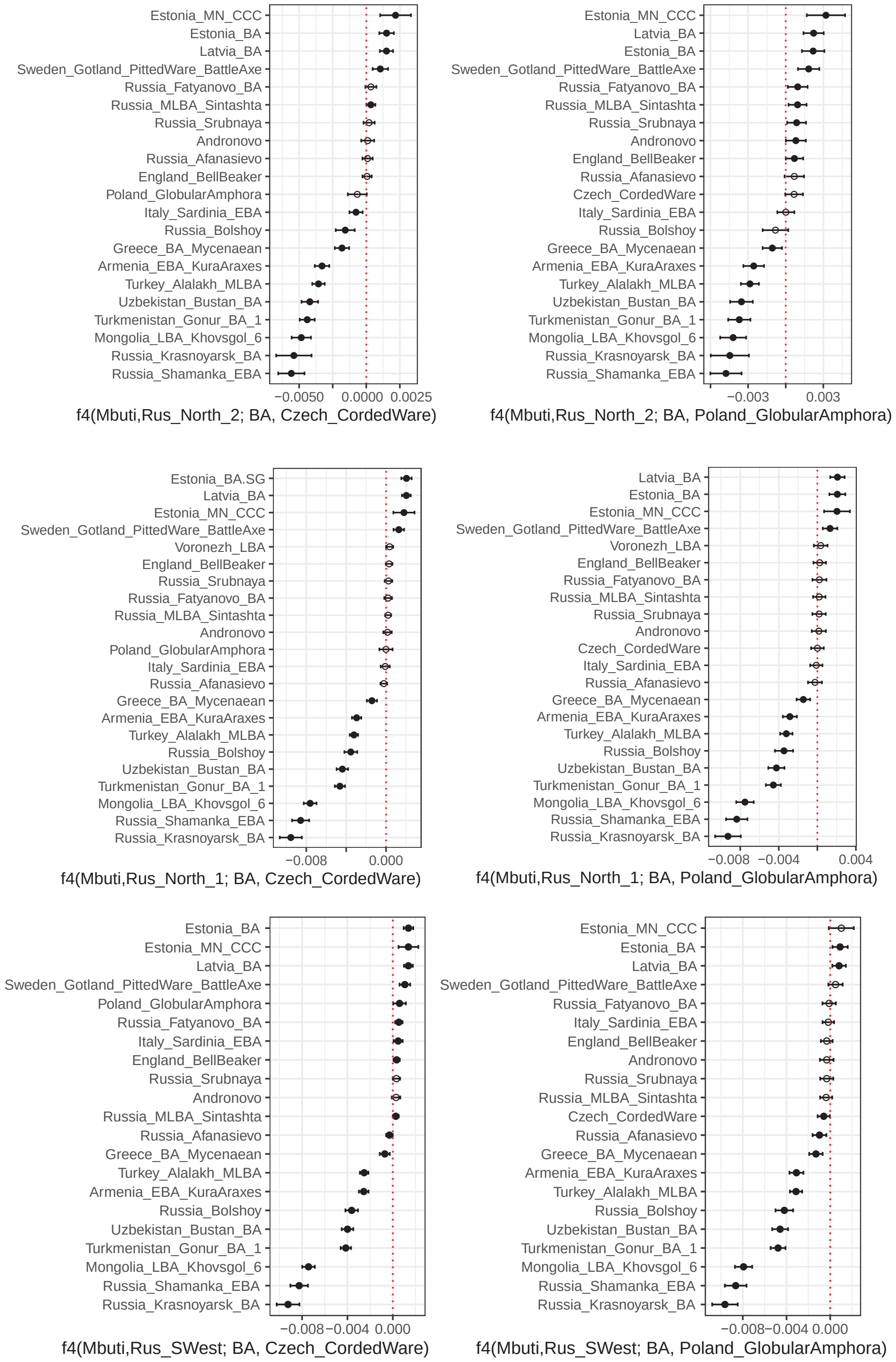

Estonia_MN_CCC
Estonia_BA
Latvia_BA
Sweden_Gotland_PittedWare_BattleAxe
Russia_Fatyanovo_BA
Russia_MLBA_Sintashta
Russia_Srubnaya
Andronovo
Russia_Afanasievo
England_BellBeaker
Poland_GlobularAmphora
Italy_Sardinia_EBA
Russia_Bolshoy
Greece_BA_Mycenaean
Armenia_EBA_KuraAraxes
Turkey_Alalakh_MLBA
Uzbekistan_Bustan_BA
Turkmenistan_Gonur_BA_1
Mongolia_LBA_Khovsgol_6
Russia_Krasnoyarsk_BA
Russia_Shamanka_EBA
−0.0050 0.0000 0.0025
f4(Mbuti,Rus_North_2; BA, Czech_CordedWare)
Estonia_MN_CCC
Latvia_BA
Estonia_BA
Sweden_Gotland_PittedWare_BattleAxe
Russia_Fatyanovo_BA
Russia_MLBA_Sintashta
Russia_Srubnaya
Andronovo
England_BellBeaker
Russia_Afanasievo
Czech_CordedWare
Italy_Sardinia_EBA
Russia_Bolshoy
Greece_BA_Mycenaean
Armenia_EBA_KuraAraxes
Turkey_Alalakh_MLBA
Uzbekistan_Bustan_BA
Turkmenistan_Gonur_BA_1
Mongolia_LBA_Khovsgol_6
Russia_Krasnoyarsk_BA
Russia_Shamanka_EBA
−0.003 0.003
f4(Mbuti,Rus_North_2; BA, Poland_GlobularAmphora)
Estonia_BA.SG
Latvia_BA
Estonia_MN_CCC
Sweden_Gotland_PittedWare_BattleAxe
Voronezh_LBA
England_BellBeaker
Russia_Srubnaya
Russia_Fatyanovo_BA
Russia_MLBA_Sintashta
Andronovo
Poland_GlobularAmphora
Italy_Sardinia_EBA
Russia_Afanasievo
Greece_BA_Mycenaean
Armenia_EBA_KuraAraxes
Turkey_Alalakh_MLBA
Russia_Bolshoy
Uzbekistan_Bustan_BA
Turkmenistan_Gonur_BA_1
Mongolia_LBA_Khovsgol_6
Russia_Shamanka_EBA
Russia_Krasnoyarsk_BA
−0.008 0.000
f4(Mbuti,Rus_North_1; BA, Czech_CordedWare)
Latvia_BA
Estonia_BA
Estonia_MN_CCC
Sweden_Gotland_PittedWare_BattleAxe
Voronezh_LBA
England_BellBeaker
Russia_Fatyanovo_BA
Russia_MLBA_Sintashta
Russia_Srubnaya
Andronovo
Czech_CordedWare
Italy_Sardinia_EBA
Russia_Afanasievo
Greece_BA_Mycenaean
Armenia_EBA_KuraAraxes
Turkey_Alalakh_MLBA
Russia_Bolshoy
Uzbekistan_Bustan_BA
Turkmenistan_Gonur_BA_1
Mongolia_LBA_Khovsgol_6
Russia_Shamanka_EBA
Russia_Krasnoyarsk_BA
−0.008 −0.004 0.004
f4(Mbuti,Rus_North_1; BA, Poland_GlobularAmphora)
Estonia_BA
Estonia_MN_CCC
Latvia_BA
Sweden_Gotland_PittedWare_BattleAxe
Poland_GlobularAmphora
Russia_Fatyanovo_BA
Italy_Sardinia_EBA
England_BellBeaker
Russia_Srubnaya
Andronovo
Russia_MLBA_Sintashta
Russia_Afanasievo
Greece_BA_Mycenaean
Turkey_Alalakh_MLBA
Armenia_EBA_KuraAraxes
Russia_Bolshoy
Uzbekistan_Bustan_BA
Turkmenistan_Gonur_BA_1
Mongolia_LBA_Khovsgol_6
Russia_Shamanka_EBA
Russia_Krasnoyarsk_BA
−0.008 −0.004 0.000
f4(Mbuti,Rus_SWest; BA, Czech_CordedWare)
Estonia_MN_CCC
Estonia_BA
Latvia_BA
Sweden_Gotland_PittedWare_BattleAxe
Russia_Fatyanovo_BA
Italy_Sardinia_EBA
England_BellBeaker
Andronovo
Russia_Srubnaya
Russia_MLBA_Sintashta
Czech_CordedWare
Russia_Afanasievo
Greece_BA_Mycenaean
Armenia_EBA_KuraAraxes
Turkey_Alalakh_MLBA
Russia_Bolshoy
Uzbekistan_Bustan_BA
Turkmenistan_Gonur_BA_1
Mongolia_LBA_Khovsgol_6
Russia_Shamanka_EBA
Russia_Krasnoyarsk_BA
−0.008 −0.004 0.000
f4(Mbuti,Rus_SWest; BA, Poland_GlobularAmphora)

**Supplementary Fig. 40. *f*4-statistic results of form *f4*(Mbuti,Rus_North_2; BA, Czech_CordedWare/Poland_GlobularAmphora).**

BA - tested BA groups, Rus’ - Rus’ genetic subclusters.

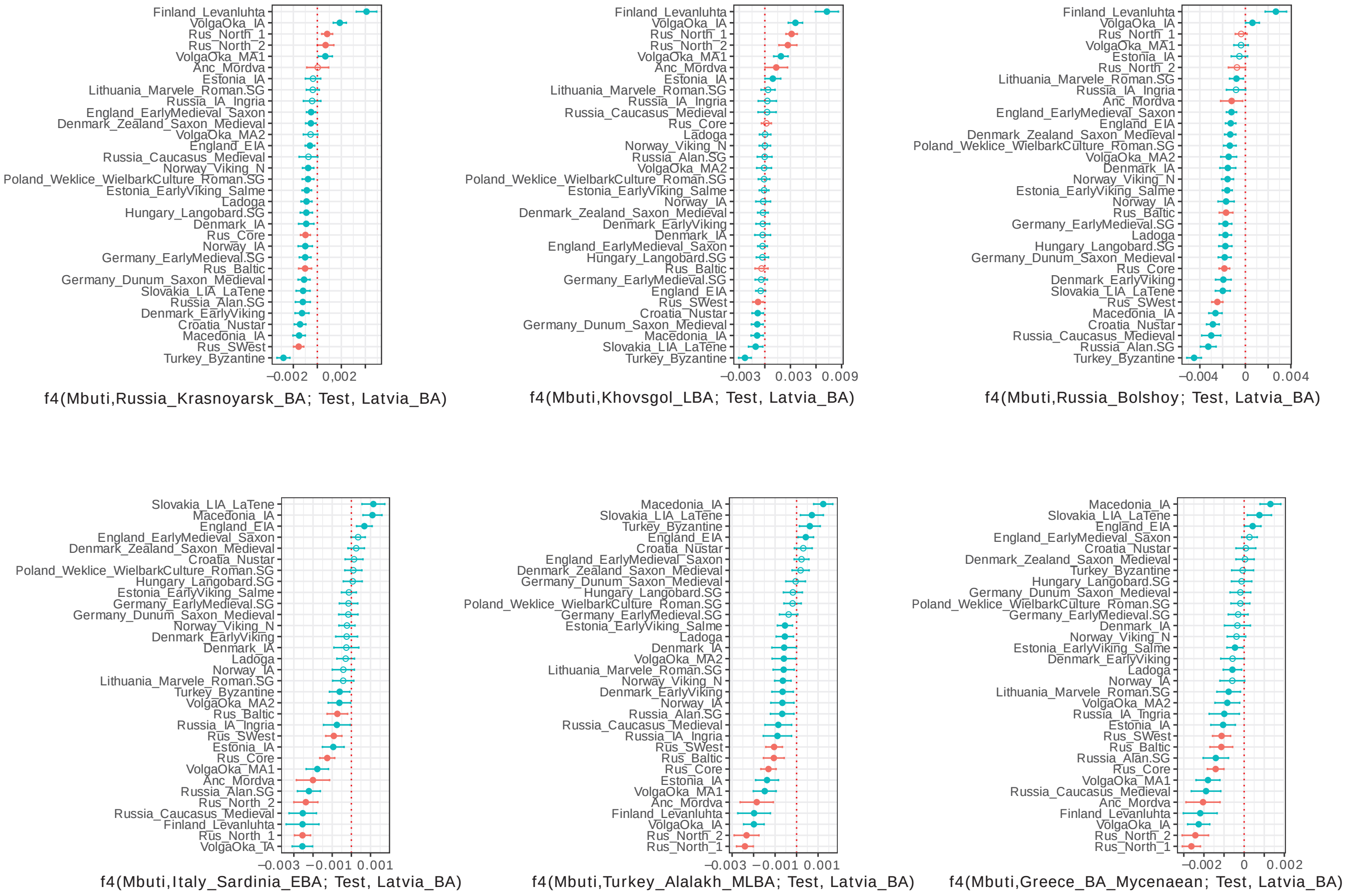
Finland_Levanluhta
VolgaOka_IA
Rus_North_1
Rus_North_2
VolgaOka_MA1
Anc_Mordva
Estonia_IA
Lithuania_Marvele_Roman.SG
Russia_IA_Ingria
England_EarlyMedieval_Saxon
Denmark_Zealand_Saxon_Medieval
VolgaOka_MA2
England_EIA
Russia_Caucasus_Medieval
Norway_Viking_N
Poland_Weklice_WielbarkCulture_Roman.SG
Estonia_EarlyViking_Salme
Ladoga
Hungary_Langobard.SG
Denmark_IA
Rus_Core
Norway_IA
Germany_EarlyMedieval.SG
Rus_Baltic
Germany_Dunum_Saxon_Medieval
Slovakia_LIA_LaTene
Russia_Alan.SG
Denmark_EarlyViking
Croatia_Nustar
Macedonia_IA
Rus_SWest
Turkey_Byzantine
−0.002
0.002
f4(Mbuti,Russia_Krasnoyarsk_BA; Test, Latvia_BA)
Finland_Levanluhta
VolgaOka_IA
Rus_North_1
Rus_North_2
VolgaOka_MA1
Anc_Mordva
Estonia_IA
Lithuania_Marvele_Roman.SG
Russia_IA_Ingria
Russia_Caucasus_Medieval
Rus_Core
Ladoga
Norway_Viking_N
Russia_Alan.SG
VolgaOka_MA2
Poland_Weklice_WielbarkCulture_Roman.SG
Estonia_EarlyViking_Salme
Norway_IA
Denmark_Zealand_Saxon_Medieval
Denmark_EarlyViking
Denmark_IA
England_EarlyMedieval_Saxon
Hungary_Langobard.SG
Rus_Baltic
Germany_EarlyMedieval.SG
England_EIA
Rus_SWest
Croatia_Nustar
Germany_Dunum_Saxon_Medieval
Macedonia_IA
Slovakia_LIA_LaTene
Turkey_Byzantine
−0.003
0.003
0.009
f4(Mbuti,Khovsgol_LBA; Test, Latvia_BA)
Finland_Levanluhta
VolgaOka_IA
Rus_North_1
VolgaOka_MA1
Estonia_IA
Rus_North_2
Lithuania_Marvele_Roman.SG
Russia_IA_Ingria
Anc_Mordva
England_EarlyMedieval_Saxon
England_EIA
Denmark_Zealand_Saxon_Medieval
Poland_Weklice_WielbarkCulture_Roman.SG
VolgaOka_MA2
Denmark_IA
Norway_Viking_N
Estonia_EarlyViking_Salme
Norway_IA
Rus_Baltic
Germany_EarlyMedieval.SG
Ladoga
Hungary_Langobard.SG
Germany_Dunum_Saxon_Medieval
Rus_Core
Denmark_EarlyViking
Slovakia_LIA_LaTene
Rus_SWest
Macedonia_IA
Croatia_Nustar
Russia_Caucasus_Medieval
Russia_Alan.SG
Turkey_Byzantine
−0.004
0
0.004
f4(Mbuti,Russia_Bolshoy; Test, Latvia_BA)
Slovakia_LIA_LaTene
Macedonia_IA
England_EIA
England_EarlyMedieval_Saxon
Denmark_Zealand_Saxon_Medieval
Croatia_Nustar
Poland_Weklice_WielbarkCulture_Roman.SG
Hungary_Langobard.SG
Estonia_EarlyViking_Salme
Germany_EarlyMedieval.SG
Germany_Dunum_Saxon_Medieval
Norway_Viking_N
Denmark_EarlyViking
Denmark_IA
Ladoga
Norway_IA
Lithuania_Marvele_Roman.SG
Turkey_Byzantine
VolgaOka_MA2
Rus_Baltic
Russia_IA_Ingria
Rus_SWest
Estonia_IA
Rus_Core
VolgaOka_MA1
Anc_Mordva
Russia_Alan.SG
Rus_North_2
Russia_Caucasus_Medieval
Finland_Levanluhta
Rus_North_1
VolgaOka_IA
−0.003
−0.001
0.001
f4(Mbuti,Italy_Sardinia_EBA; Test, Latvia_BA)
Macedonia_IA
Slovakia_LIA_LaTene
Turkey_Byzantine
England_EIA
Croatia_Nustar
England_EarlyMedieval_Saxon
Denmark_Zealand_Saxon_Medieval
Germany_Dunum_Saxon_Medieval
Hungary_Langobard.SG
Poland_Weklice_WielbarkCulture_Roman.SG
Germany_EarlyMedieval.SG
Estonia_EarlyViking_Salme
Ladoga
Denmark_IA
VolgaOka_MA2
Lithuania_Marvele_Roman.SG
Norway_Viking_N
Denmark_EarlyViking
Norway_IA
Russia_Alan.SG
Russia_Caucasus_Medieval
Russia_IA_Ingria
Rus_SWest
Rus_Baltic
Rus_Core
Estonia_IA
VolgaOka_MA1
Anc_Mordva
Finland_Levanluhta
VolgaOka_IA
Rus_North_2
Rus_North_1
−0.003
−0.001
0.001
f4(Mbuti,Turkey_Alalakh_MLBA; Test, Latvia_BA)
Macedonia_IA
Slovakia_LIA_LaTene
England_EIA
England_EarlyMedieval_Saxon
Croatia_Nustar
Denmark_Zealand_Saxon_Medieval
Turkey_Byzantine
Hungary_Langobard.SG
Germany_Dunum_Saxon_Medieval
Poland_Weklice_WielbarkCulture_Roman.SG
Germany_EarlyMedieval.SG
Denmark_IA
Norway_Viking_N
Estonia_EarlyViking_Salme
Denmark_EarlyViking
Ladoga
Norway_IA
Lithuania_Marvele_Roman.SG
VolgaOka_MA2
Russia_IA_Ingria
Estonia_IA
Rus_SWest
Rus_Baltic
Russia_Alan.SG
Rus_Core
VolgaOka_MA1
Russia_Caucasus_Medieval
Anc_Mordva
Finland_Levanluhta
VolgaOka_IA
Rus_North_2
Rus_North_1
−0.002
0
0.002
f4(Mbuti,Greece_BA_Mycenaean; Test, Latvia_BA)

**Supplementary Fig. 41. f4-statistic results of form f4 (Krasnoyarsk_BA/Greece_BA_Mycenian, Mbuti; Test, Latvia_BA).**

Test - Rus’ genetic subclusters or other ancient IA and MA groups.

Cladality test   f4(Medieval_pops, Mbuti; Major_Rus', Test)

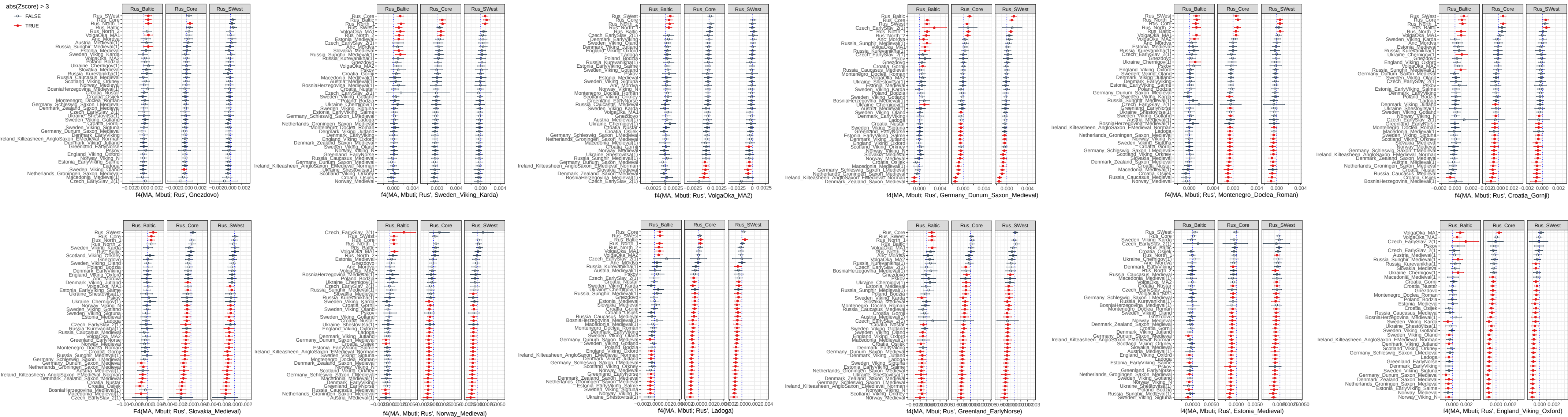

**Supplementary Fig. 42.** ***f*4-statistics of the form *f*4(Rus’, Mbuti; Ancient_pops, Ancient_pops).**

Rus’ - genetic subclusters of our Rus’ samples, Ancient_pops - previously published medieval groups.

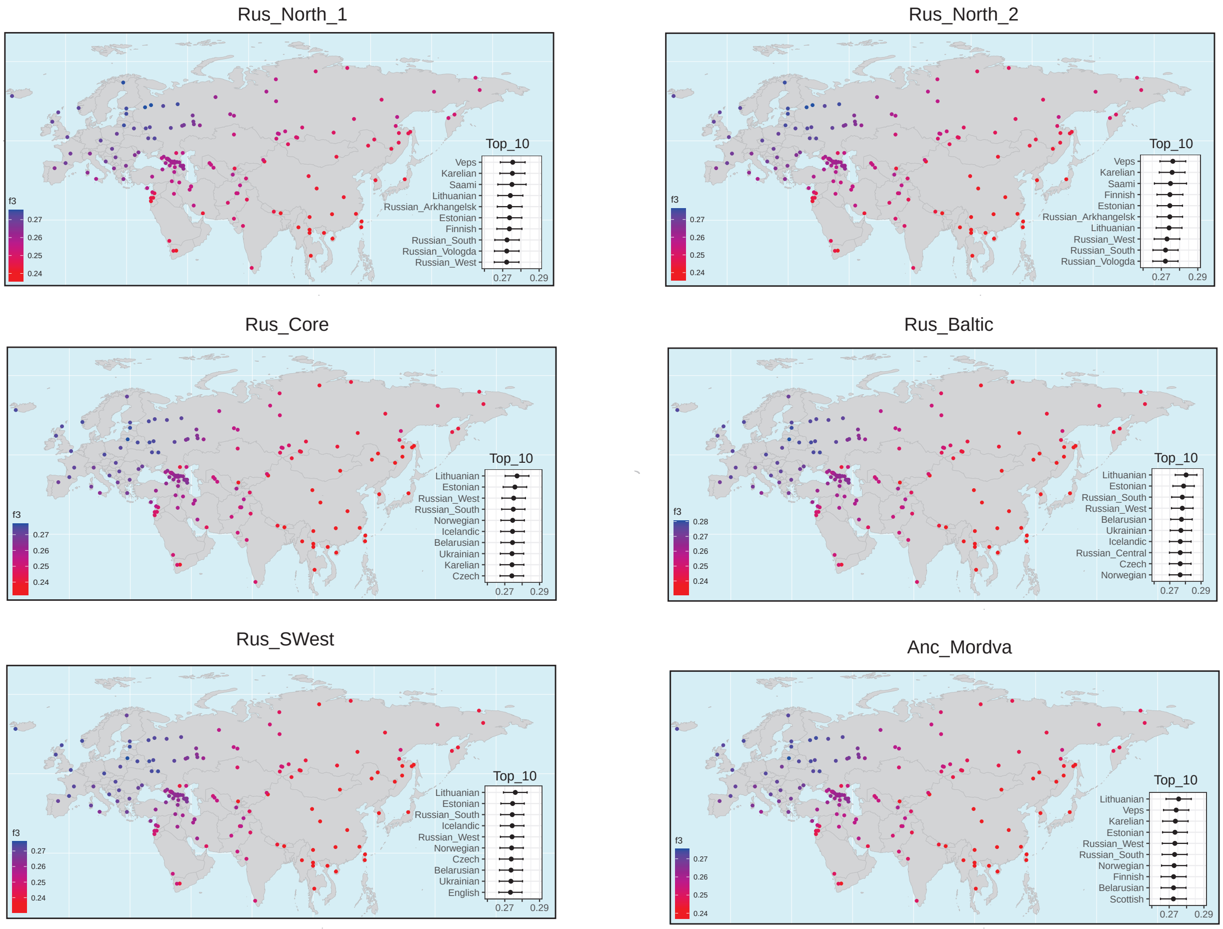

Rus_North_1
Top_10
Veps
Karelian
Saami
Lithuanian
Russian_Arkhangelsk
Estonian
Finnish
Russian_South
Russian_Vologda
Russian_West
0.27
0.29
f3
0.27
0.26
0.25
0.24
Rus_North_2
Top_10
Veps
Karelian
Saami
Finnish
Estonian
Russian_Arkhangelsk
Lithuanian
Russian_West
Russian_South
Russian_Vologda
0.27
0.29
f3
0.27
0.26
0.25
0.24
Rus_Core
Top_10
Lithuanian
Estonian
Russian_West
Russian_South
Norwegian
Icelandic
Belarusian
Ukrainian
Karelian
Czech
0.27
0.29
f3
0.27
0.26
0.25
0.24
Rus_Baltic
Top_10
Lithuanian
Estonian
Russian_South
Russian_West
Belarusian
Ukrainian
Icelandic
Russian_Central
Czech
Norwegian
0.27
0.29
f3
0.28
0.27
0.26
0.25
0.24
Rus_SWest
Top_10
Lithuanian
Estonian
Russian_South
Icelandic
Russian_West
Norwegian
Czech
Belarusian
Ukrainian
English
0.27
0.29
f3
0.27
0.26
0.25
0.24
Anc_Mordva
Top_10
Lithuanian
Veps
Karelian
Estonian
Russian_West
Russian_South
Norwegian
Finnish
Belarusian
Scottish
0.27
0.29
f3
0.27
0.26
0.25
0.24

**Supplementary Fig. 43. Outgroups *f*3-statistic results of form *f3*(Rus',Target;Mbuti).** Rus' - ancient Rus' genetic subclusters, Target - present-day populations. Blue dots indicate the highest level of allele sharing between the Rus' genetic subclusters and the target present-day population, and red dots indicate the lowest level of allele sharing. Top 10 highest f3-statistics results provided for each genetic subcluster.

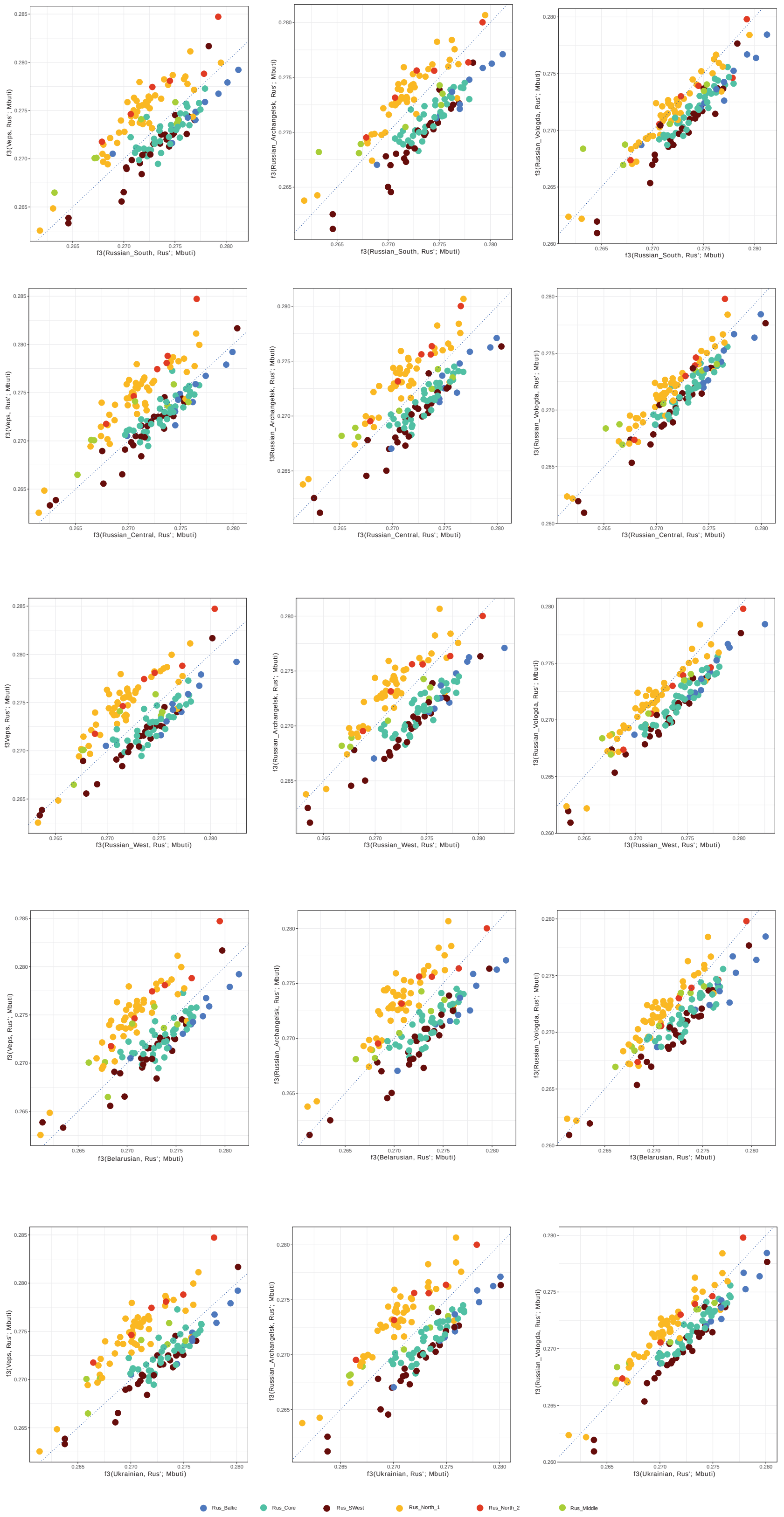
f3(Veps, Rus'; Mbuti)
f3(Russian_Archangelsk, Rus'; Mbuti)
f3(Russian_Vologda, Rus'; Mbuti)
f3(Russian_South, Rus'; Mbuti)
f3(Russian_Central, Rus'; Mbuti)
f3(Russian_West, Rus'; Mbuti)
f3(Belarusian, Rus'; Mbuti)
f3(Ukrainian, Rus'; Mbuti)
Rus_Baltic
Rus_Core
Rus_SWest
Rus_North_1
Rus_North_2
Rus_Middle

**Supplementary Fig. 44. Outgroups *f*3-statistic results of form *f3*(Anc_Rus',Modern_pops; Mbuti) plotted against each other.**

Anc_Rus' - tested Ancient Rus' individuals. Modern_pops - target present-day populations.

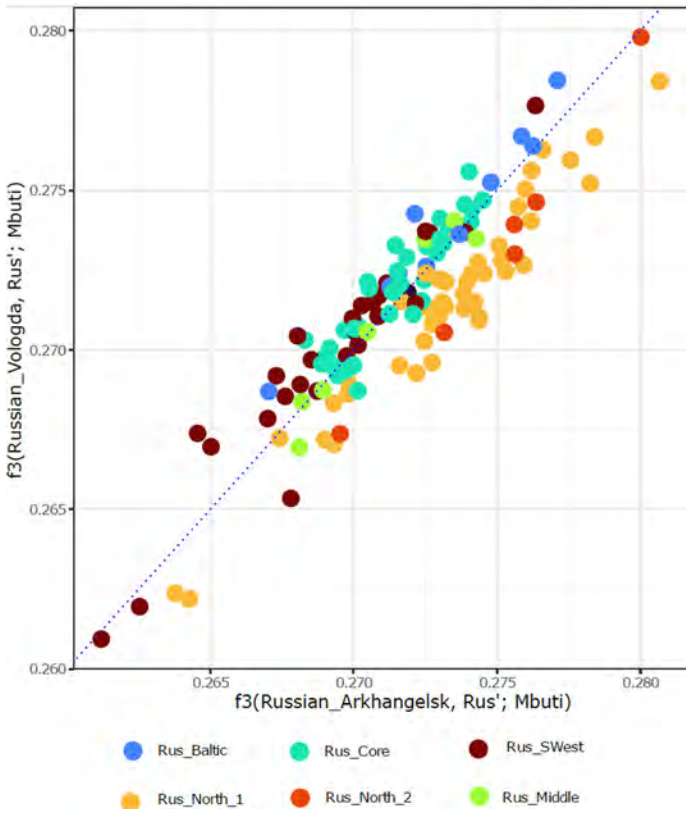


**Supplementary Fig. 45. Outgroups *f*3-statistic results of the form *f3*(Anc_Rus',Modern_pops;Mbuti) plotted against each other: present day Russians from Arkhangelsk (X axis) and present-day Russians from Vologda (Y axis).**
Anc_Rus' - tested Ancient Rus' individuals.

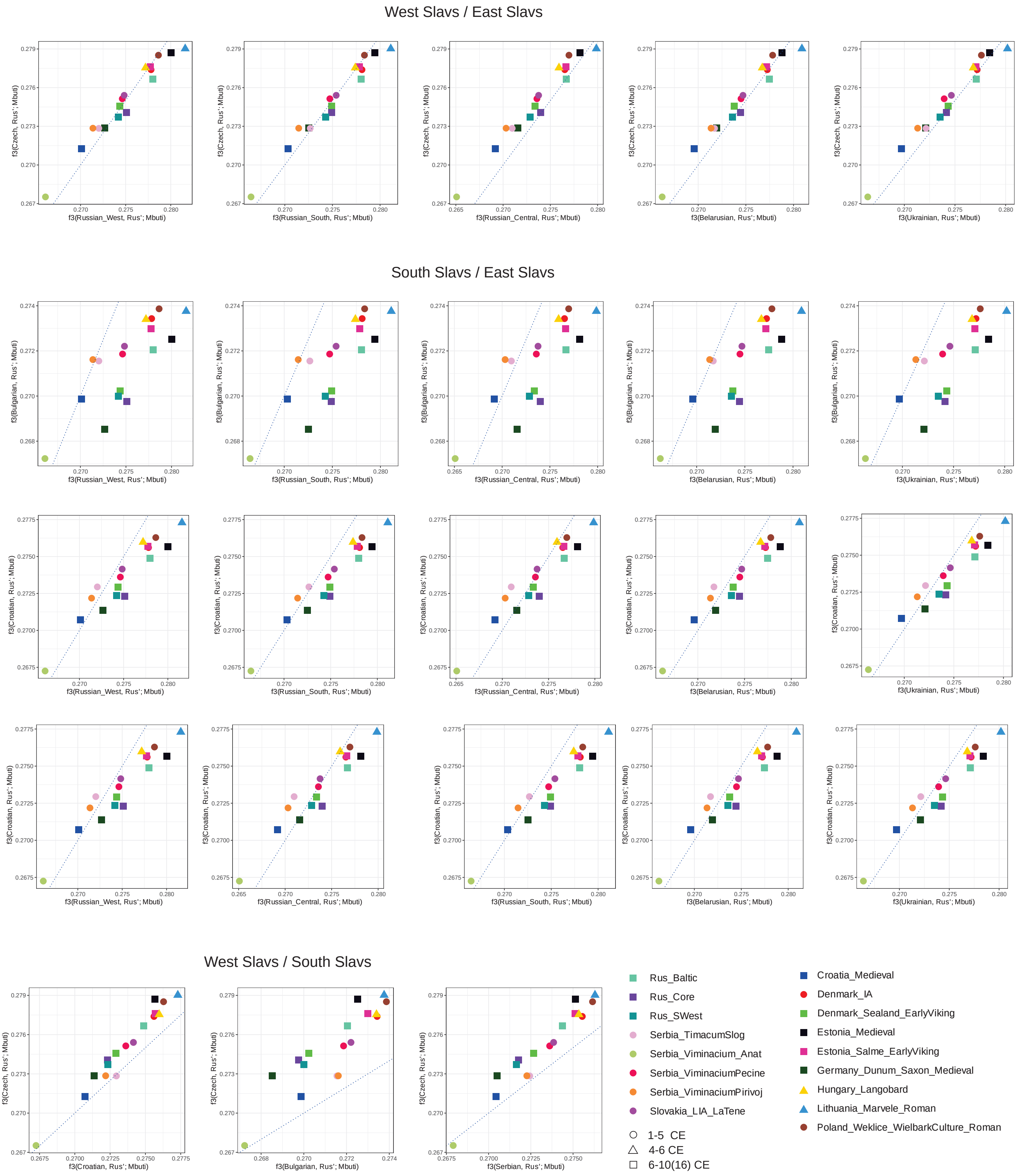
West Slavs / East Slavs
f3(Czech, Rus'; Mbuti)
f3(Russian_West, Rus'; Mbuti)
f3(Russian_South, Rus'; Mbuti)
f3(Russian_Central, Rus'; Mbuti)
f3(Belarusian, Rus'; Mbuti)
f3(Ukrainian, Rus'; Mbuti)
South Slavs / East Slavs
f3(Bulgarian, Rus'; Mbuti)
f3(Croatian, Rus'; Mbuti)
West Slavs / South Slavs
f3(Serbian, Rus'; Mbuti)
Rus_Baltic
Rus_Core
Rus_SWest
Serbia_TimacumSlog
Serbia_Viminacium_Anat
Serbia_ViminaciumPecine
Serbia_ViminaciumPirivoj
Slovakia_LIA_LaTene
1-5 CE
4-6 CE
6-10(16) CE
Croatia_Medieval
Denmark_IA
Denmark_Sealand_EarlyViking
Estonia_Medieval
Estonia_Salme_EarlyViking
Germany_Dunum_Saxon_Medieval
Hungary_Langobard
Lithuania_Marvele_Roman
Poland_Weklice_WielbarkCulture_Roman

**Supplementary Fig. 46. Outgroups *f*3-statistic results of form *f*3(Ancient_pops, Modern_pops; Mbuti) plotted against each other.**
Modern_pops – present-day East, West and South Slavs. Ancient_pops - ancient groups geographically or historically appropriated to Rus'.

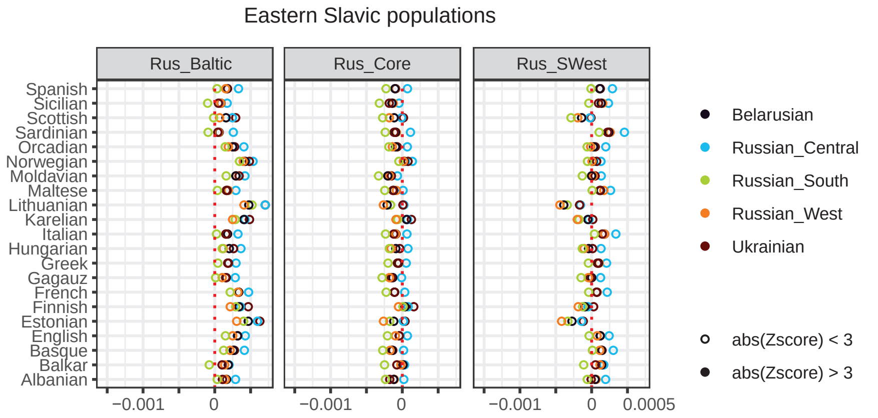
Eastern Slavic populations
Rus_Baltic
Rus_Core
Rus_SWest
Spanish
Sicilian
Scottish
Sardinian
Orcadian
Norwegian
Moldavian
Maltese
Lithuanian
Karelian
Italian
Hungarian
Greek
Gagauz
French
Finnish
Estonian
English
Basque
Balkar
Albanian
−0.001
0
0.0005
Belarusian
Russian_Central
Russian_South
Russian_West
Ukrainian
abs(Zscore) < 3
abs(Zscore) > 3

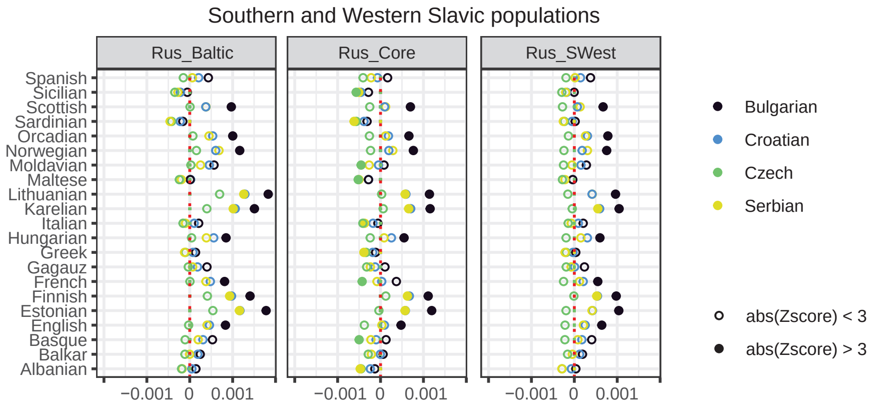
Southern and Western Slavic populations
Rus_Baltic
Rus_Core
Rus_SWest
Spanish
Sicilian
Scottish
Sardinian
Orcadian
Norwegian
Moldavian
Maltese
Lithuanian
Karelian
Italian
Hungarian
Greek
Gagauz
French
Finnish
Estonian
English
Basque
Balkar
Albanian
−0.001
0
0.001
Bulgarian
Croatian
Czech
Serbian
abs(Zscore) < 3
abs(Zscore) > 3

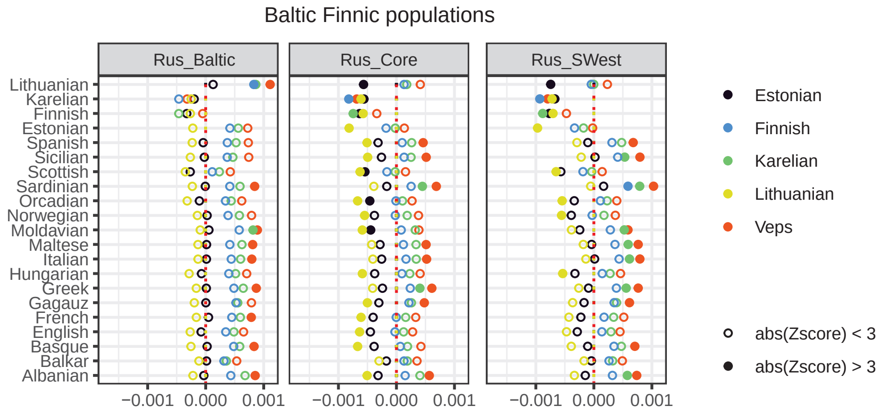
Baltic Finnic populations
Rus_Baltic
Rus_Core
Rus_SWest
Lithuanian
Karelian
Finnish
Estonian
Spanish
Sicilian
Scottish
Sardinian
Orcadian
Norwegian
Moldavian
Maltese
Italian
Hungarian
Greek
Gagauz
French
English
Basque
Balkar
Albanian
−0.001
0.000
0.001
Estonian
Finnish
Karelian
Lithuanian
Veps
abs(Zscore) < 3
abs(Zscore) > 3

**Supplementary Fig. 47. Cladality test from *f*4-statistics compared Ancient Major Rus' cluster groups to present-day European populations of the form *f*4(Mbuti, Modern_pops; Ancient_Rus',Target).**
Modern_pops are present-day European populations, and Target is a set of geographically and genetically appropriated present-day Eastern-European populations (see Supplementary Informations S2.3). Non-filled circles indicate statistically non-significant (|Z| < 3) estimates of f4.

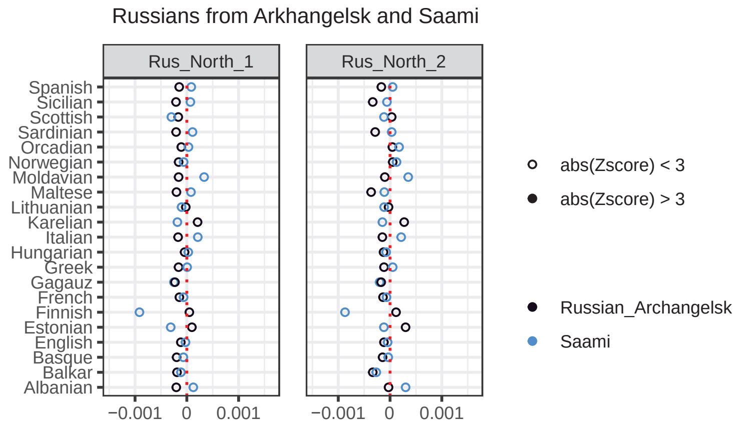
Russians from Arkhangelsk and Saami
Rus_North_1
Rus_North_2
Spanish
Sicilian
Scottish
Sardinian
Orcadian
Norwegian
Moldavian
Maltese
Lithuanian
Karelian
Italian
Hungarian
Greek
Gagauz
French
Finnish
Estonian
English
Basque
Balkar
Albanian
−0.001
0
0.001
abs(Zscore) < 3
abs(Zscore) > 3
Russian_Archangelsk
Saami

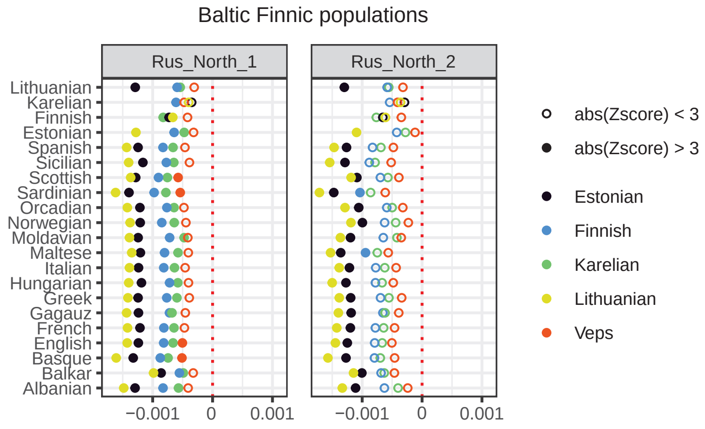
Baltic Finnic populations
Rus_North_1
Rus_North_2
Lithuanian
Karelian
Finnish
Estonian
Spanish
Sicilian
Scottish
Sardinian
Orcadian
Norwegian
Moldavian
Maltese
Italian
Hungarian
Greek
Gagauz
French
English
Basque
Balkar
Albanian
−0.001
0
0.001
abs(Zscore) < 3
abs(Zscore) > 3
Estonian
Finnish
Karelian
Lithuanian
Veps

**Supplementary Fig. 48. Cladality test of the from *f*4-statistics compared Ancient Northern Rus' cluster groups to present-day European populations of form *f4(*Mbuti, Modern_pops; Ancient_Rus',Target).**

Modern_pops are present-day European populations, and Target is a set of geographically and genetically appropriated present-day Eastern-European populations (see Supplementary Informations S2.3). Non-filled circles indicate statistically non-significant (|Z| < 3) estimates of f4.

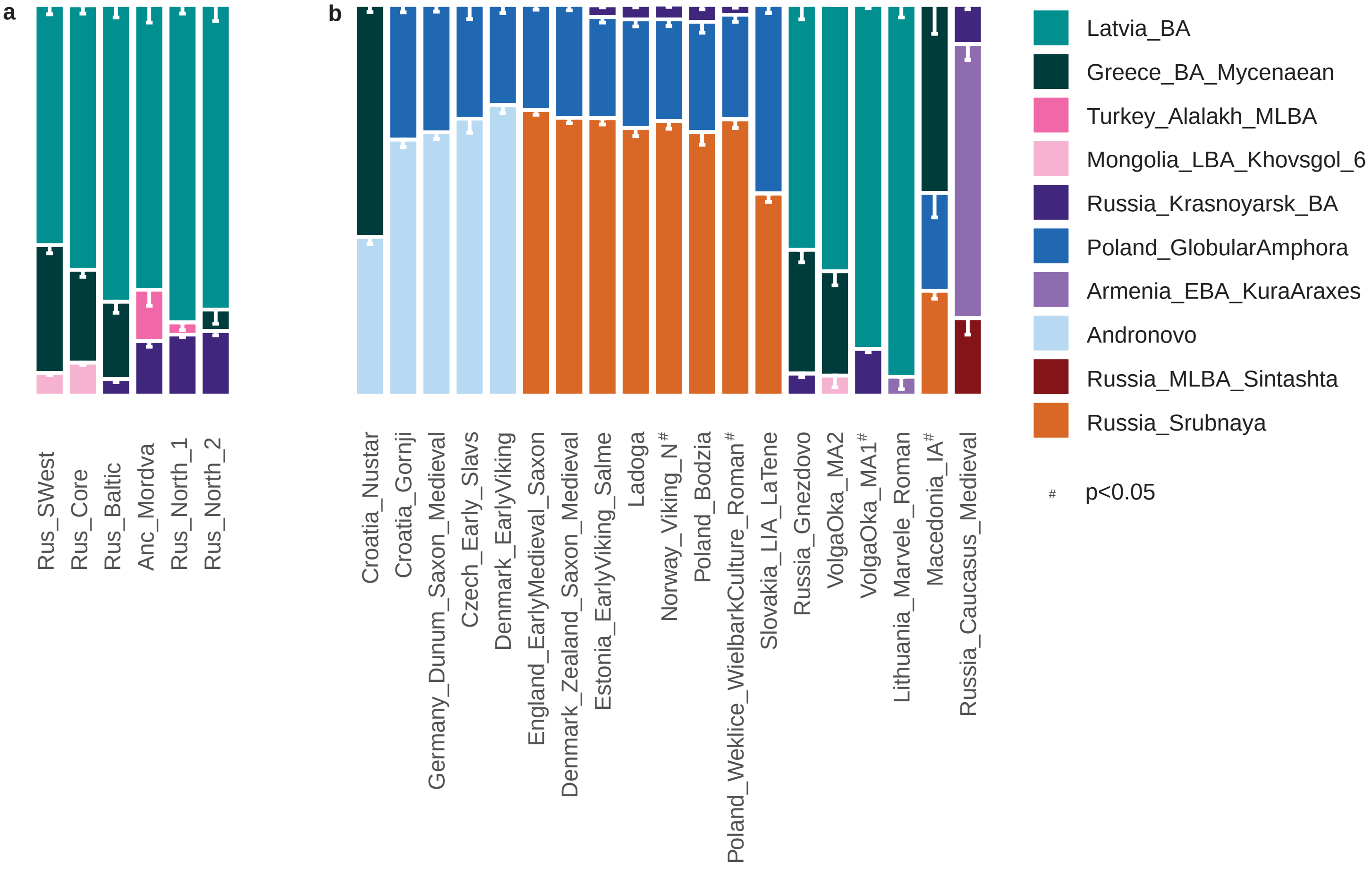
a
b
Rus_SWest
Rus_Core
Rus_Baltic
Anc_Mordva
Rus_North_1
Rus_North_2
Croatia_Nustar
Croatia_Gornji
Germany_Dunum_Saxon_Medieval
Czech_Early_Slavs
Denmark_EarlyViking
England_EarlyMedieval_Saxon
Denmark_Zealand_Saxon_Medieval
Estonia_EarlyViking_Salme
Ladoga
Norway_Viking_N#
Poland_Bodzia
Poland_Weklice_WielbarkCulture_Roman#
Slovakia_LIA_LaTene
Russia_Gnezdovo
VolgaOka_MA2
VolgaOka_MA1#
Lithuania_Marvele_Roman
Macedonia_IA#
Russia_Caucasus_Medieval
Latvia_BA
Greece_BA_Mycenaean
Turkey_Alalakh_MLBA
Mongolia_LBA_Khovsgol_6
Russia_Krasnoyarsk_BA
Poland_GlobularAmphora
Armenia_EBA_KuraAraxes
Andronovo
Russia_MLBA_Sintashta
Russia_Srubnaya
# p<0.05

**Supplementary Fig. 49. Three-source BA-based qpAdm model elucidates an hypothetical history of admixture among the genetic outliers of ancient Rus' people**
(see details in  Supplementary Informations S2.4).

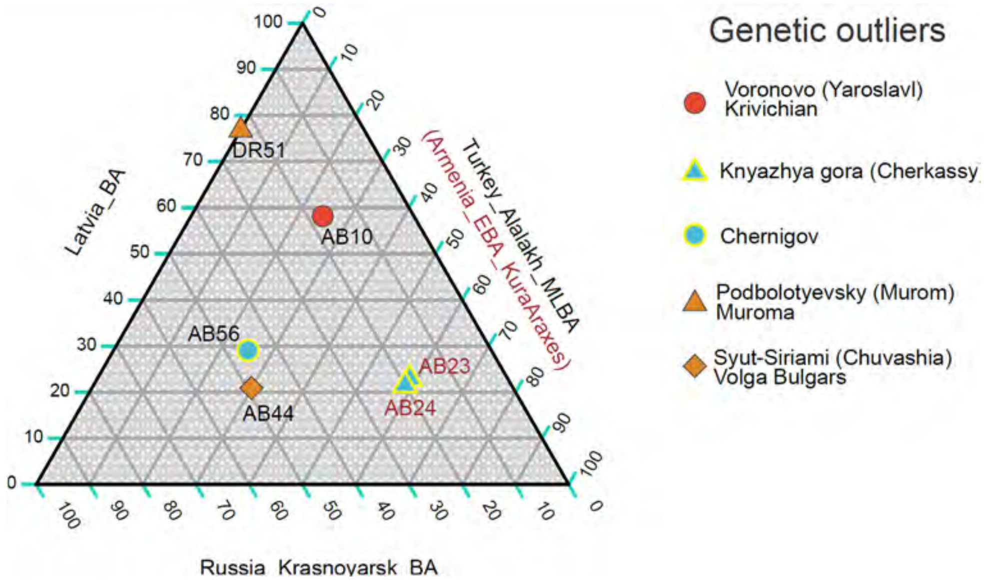


**Supplementary Fig. 50.** ***qpADM*** **models elucidate a hypothetical history of admixture for ancient European populations of the turn of the era and 1st centuries CE based on the BA sources.**

Each bar represents the admixture proportions for each group, based on the most suitable model with the highest p-value (See details in Supplementary Informations 2.4).

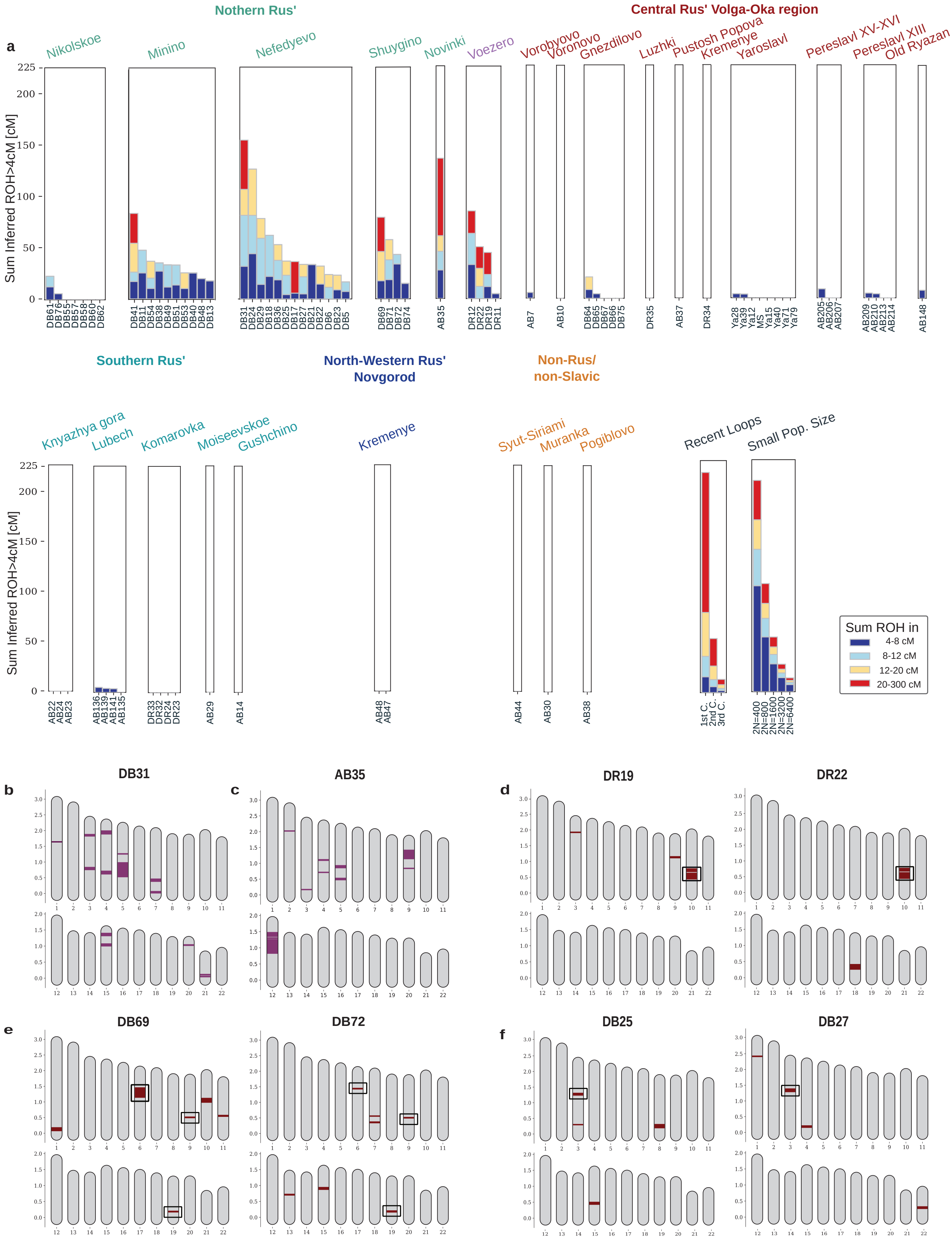

a
Nothern Rus'
Central Rus' Volga-Oka region
Nikolskoe
Minino
Nefedyevo
Shuygino
Novinki
Voezero
Vorobyovo
Voronovo
Gnezdilovo
Luzhki
Pustosh Popova
Kremenye
Yaroslavl
Pereslavl XV-XVI
Pereslavl XIII
Old Ryazan
Sum Inferred ROH>4cM [cM]
Southern Rus'
North-Western Rus' Novgorod
Non-Rus/ non-Slavic
Knyazhya gora
Lubech
Komarovka
Moiseevskoe
Gushchino
Kremenye
Syut-Siriami
Muranka
Pogiblovo
Recent Loops
Small Pop. Size
1st C.
2nd C.
3rd C.
2N=400
2N=800
2N=1600
2N=3200
2N=6400
Sum ROH in
4-8 cM
8-12 cM
12-20 cM
20-300 cM
b
DB31
c
AB35
d
DR19
DR22
e
DB69
DB72
f
DB25
DB27

**Supplementary Fig. 51. Runs of homozygosity for different geographical regions of ancient Rus' and neighboring contemporary non-Russia/non-Slavic tribes.**
(a) Distributions of inferred runs of homozygosity longer than 4 cM (ROH). (b-c) The position of ROH (purple) on the autosomes for the most homozygous individuals. (d-f) The chromosomal position of ROH (purple) on the autosomes for pairs of closely related individuals. Intersecting ROHs are framed for pairs of individuals.

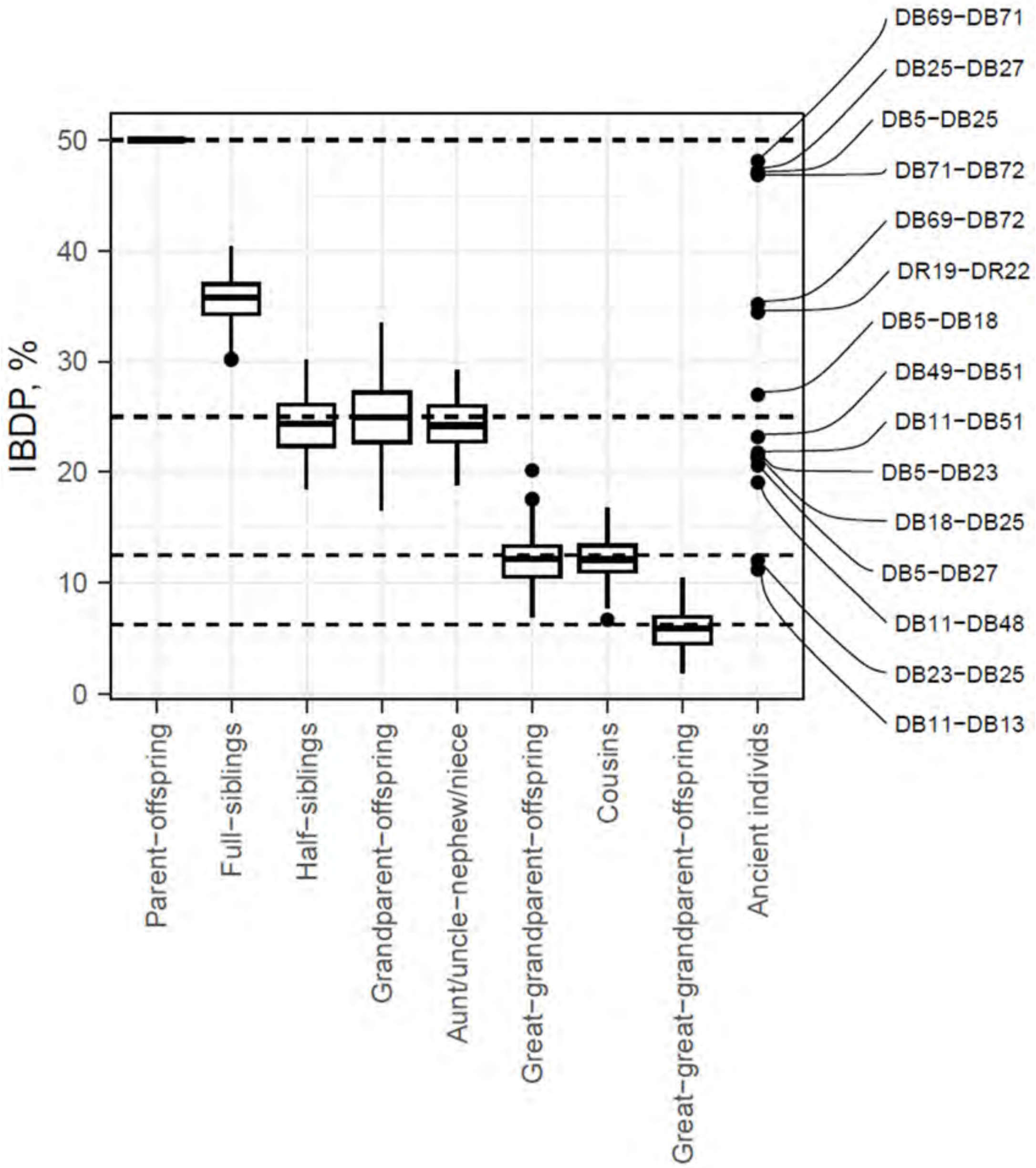


**Supplementary Fig. 52. Biological relatedness between studied individuals in comparison with simulated results.**

The simulated IBDP results for eight different degrees of relatedness (100 pairs per each degree of relatedness) are demonstrated as boxplots, while pairs of studied individuals are depicted as black filled circles. Reference levels of IBDP for parent-child (50%), grandparent-grandchild (25%), great-grandparent-great-grandchild (12.5%) and great-great-grandparent-great-great-grandchild (6.25%) relationships are shown as dashed lines. IBDP cutoff for plotting pairs of studied individuals is 10%. See also Supplementary Table 31.

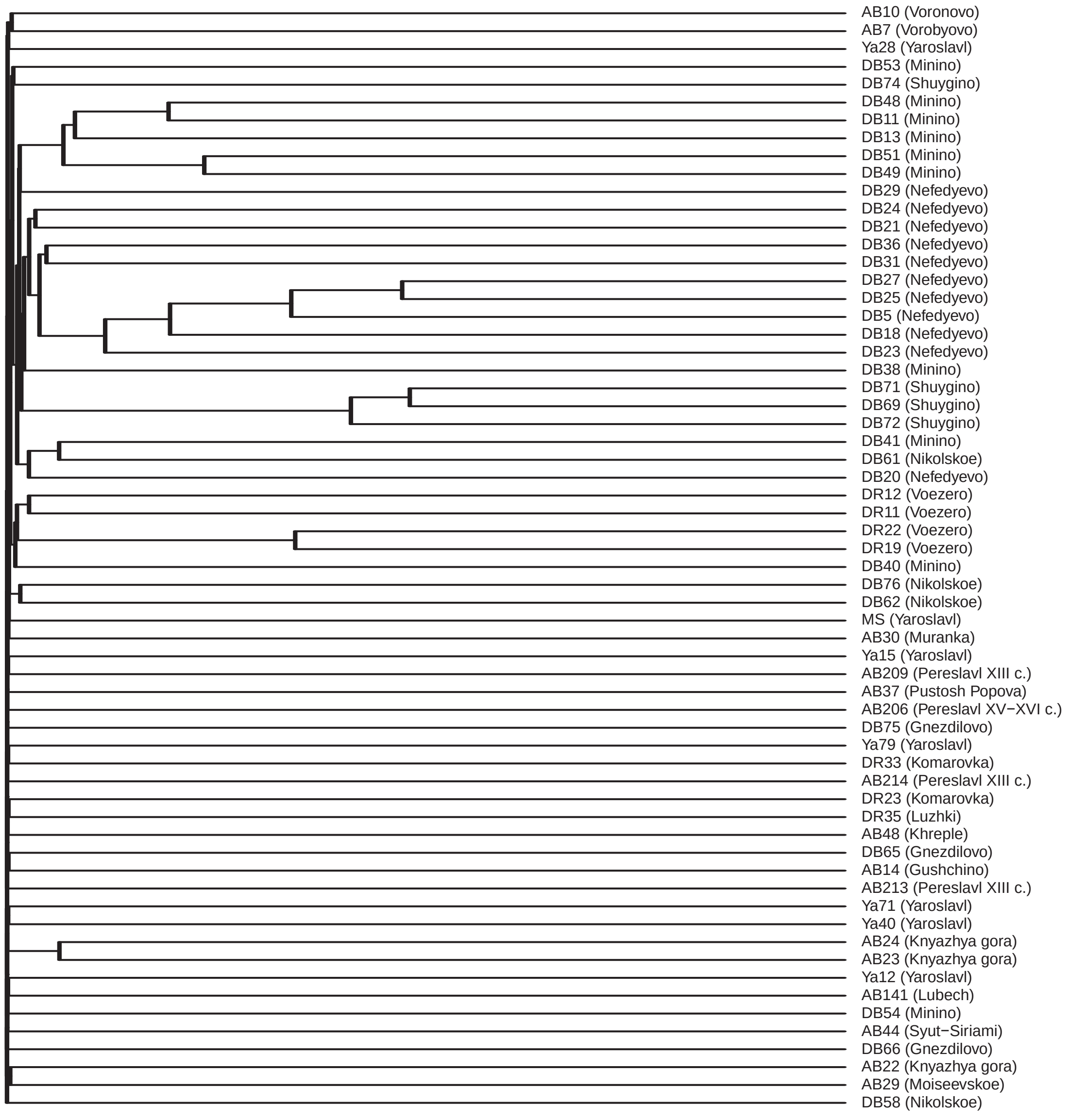
AB10 (Voronovo)
AB7 (Vorobyovo)
Ya28 (Yaroslavl)
DB53 (Minino)
DB74 (Shuygino)
DB48 (Minino)
DB11 (Minino)
DB13 (Minino)
DB51 (Minino)
DB49 (Minino)
DB29 (Nefedyevo)
DB24 (Nefedyevo)
DB21 (Nefedyevo)
DB36 (Nefedyevo)
DB31 (Nefedyevo)
DB27 (Nefedyevo)
DB25 (Nefedyevo)
DB5 (Nefedyevo)
DB18 (Nefedyevo)
DB23 (Nefedyevo)
DB38 (Minino)
DB71 (Shuygino)
DB69 (Shuygino)
DB72 (Shuygino)
DB41 (Minino)
DB61 (Nikolskoe)
DB20 (Nefedyevo)
DR12 (Voezero)
DR11 (Voezero)
DR22 (Voezero)
DR19 (Voezero)
DB40 (Minino)
DB76 (Nikolskoe)
DB62 (Nikolskoe)
MS (Yaroslavl)
AB30 (Muranka)
Ya15 (Yaroslavl)
AB209 (Pereslavl XIII c.)
AB37 (Pustosh Popova)
AB206 (Pereslavl XV−XVI c.)
DB75 (Gnezdilovo)
Ya79 (Yaroslavl)
DR33 (Komarovka)
AB214 (Pereslavl XIII c.)
DR23 (Komarovka)
DR35 (Luzhki)
AB48 (Khreple)
DB65 (Gnezdilovo)
AB14 (Gushchino)
AB213 (Pereslavl XIII c.)
Ya71 (Yaroslavl)
Ya40 (Yaroslavl)
AB24 (Knyazhya gora)
AB23 (Knyazhya gora)
Ya12 (Yaroslavl)
AB141 (Lubech)
DB54 (Minino)
AB44 (Syut−Siriami)
DB66 (Gnezdilovo)
AB22 (Knyazhya gora)
AB29 (Moiseevskoe)
DB58 (Nikolskoe)

**Supplementary Fig. 53. IBDP dendrogram.**
The length of two edges between two individuals or a group of individuals is inversely proportional to their IBDP.

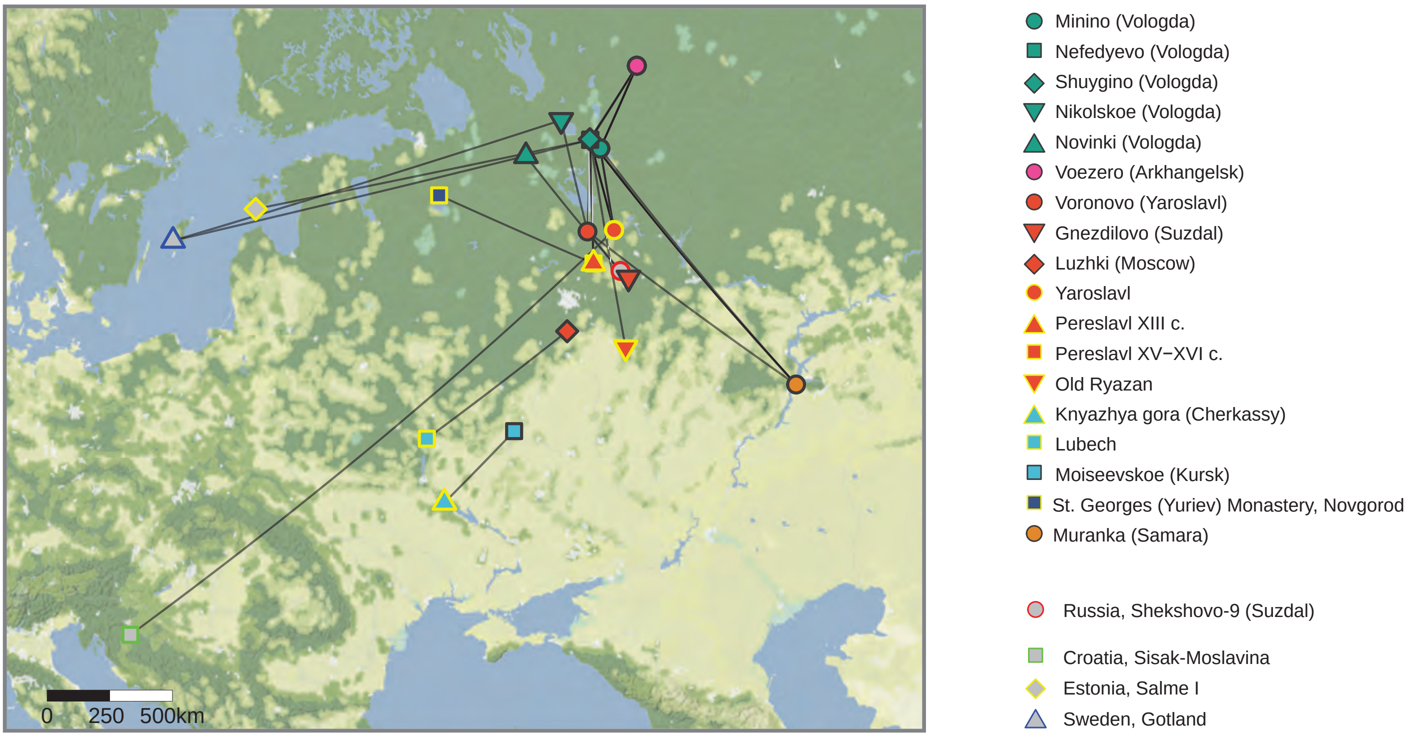
0 250 500km
Minino (Vologda)
Nefedyevo (Vologda)
Shuygino (Vologda)
Nikolskoe (Vologda)
Novinki (Vologda)
Voezero (Arkhangelsk)
Voronovo (Yaroslavl)
Gnezdilovo (Suzdal)
Luzhki (Moscow)
Yaroslavl
Pereslavl XIII c.
Pereslavl XV−XVI c.
Old Ryazan
Knyazhya gora (Cherkassy)
Lubech
Moiseevskoe (Kursk)
St. Georges (Yuriev) Monastery, Novgorod
Muranka (Samara)
Russia, Shekshovo-9 (Suzdal)
Croatia, Sisak-Moslavina
Estonia, Salme I
Sweden, Gotland

**Supplementary Fig. 54. The IBD connections among archaeological sites of medieval Rus' and with temporally appropriate Scandinavian and Balkan region groups.**

**Phylogenetic tree of haplogroup R-M420 (R1a)**

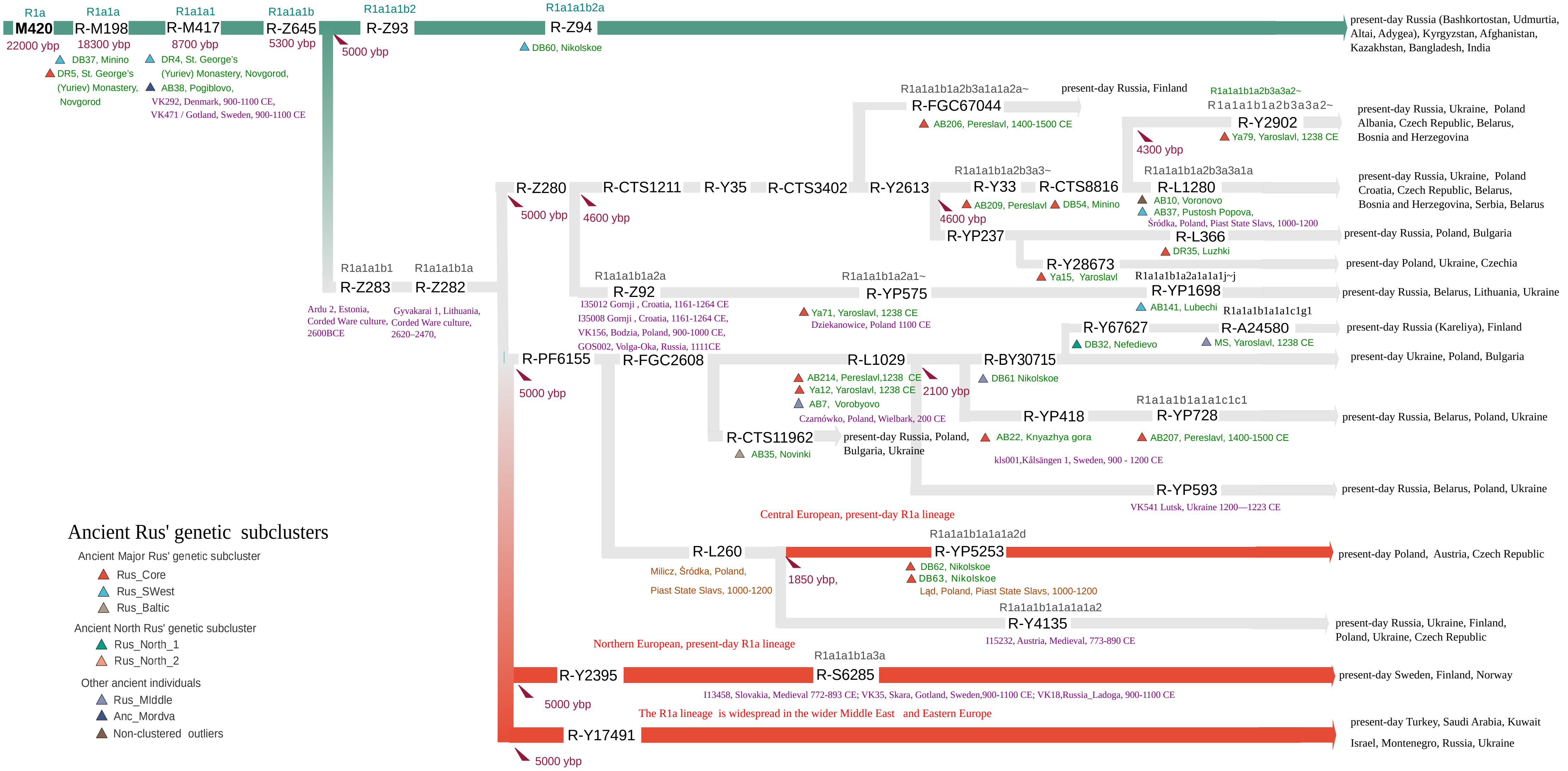

**Supplementary Fig. 55. The simplified Y-chromosome phylogenetic tree for haplogroup R1a lineages identified among newly reported ancient Rus' individuals.**

The mean age of branches is indicated according to the full database (www.yfull.com). The diagram shows two different main branches - R1a-Z93 and R1a-Z282. The R1a-Z93 lineage, which is predominant in present-day Central and South Asian populations, is indicated in green. The branch R1a-Z282 (R1a1a1b1a), which is currently predominant in Central and Eastern Europe among Slavic-related populations, is indicated in grey. The red color indicates sub-branches that are currently prevalent in Central and Northern Europe and are not found in the present-day Russian population.

Our ancient Rus' samples are indicated in green; previously published samples from the Viking Era and medieval burials in Europe and Russia are indicated in purple[97,102,161]. The brown highlight indicates samples from medieval slavic-related burials in Poland.[126] The shape and color of the symbols for the newly reported individuals correspond to those on the Fig. 3. The R1a paternal lineage, currently found in populations of Central Asia, South Asia, and West Asia, is shown in green. The R1a paternal lineage, currently found in Central and Northern European populations, is shown in gray.

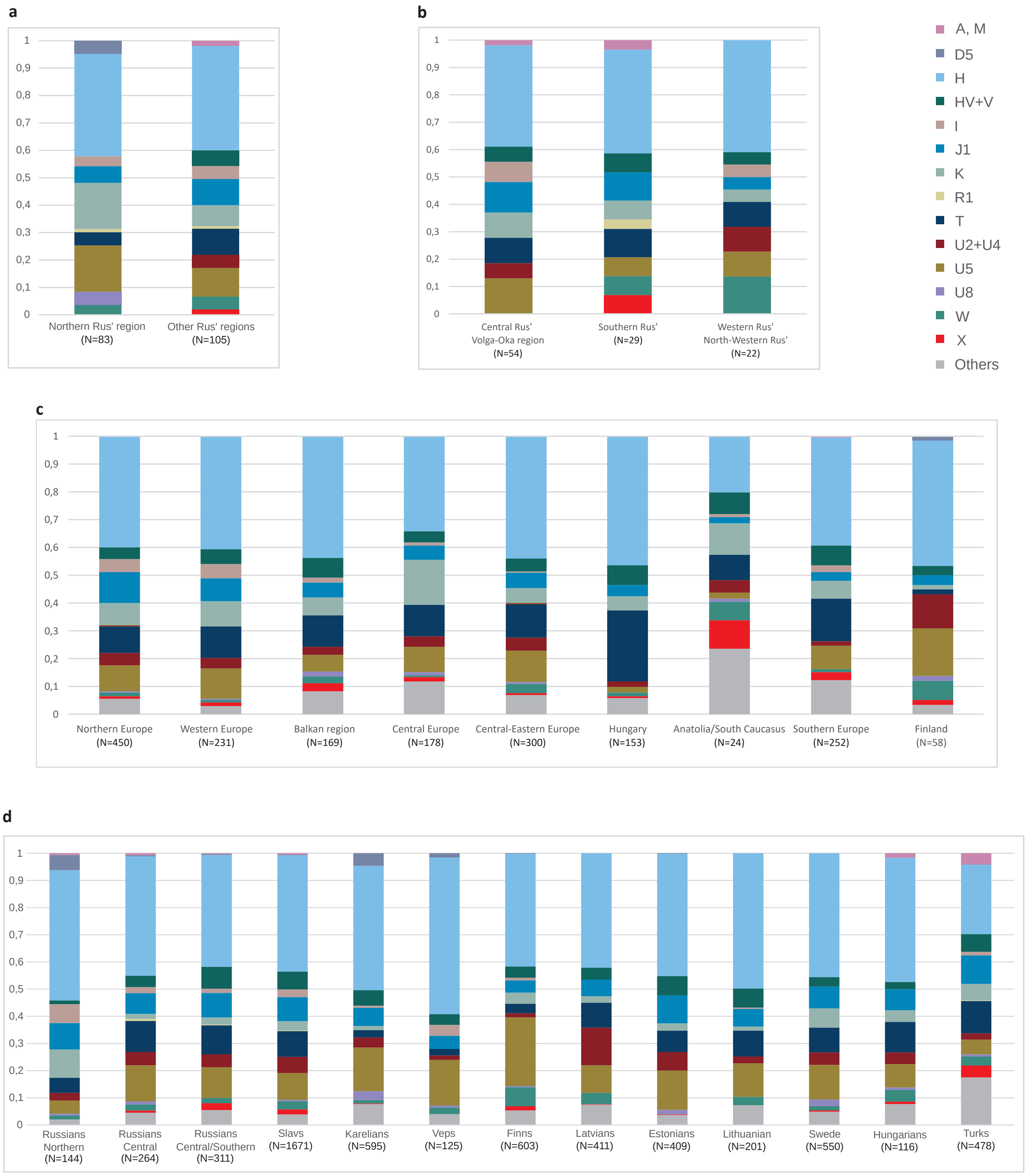
a
1
0,9
0,8
0,7
0,6
0,5
0,4
0,3
0,2
0,1
0
Northern Rus' region (N=83)
Other Rus' regions (N=105)
b
Central Rus' Volga-Oka region (N=54)
Southern Rus' (N=29)
Western Rus' North-Western Rus' (N=22)
A, M
D5
H
HV+V
I
J1
K
R1
T
U2+U4
U5
U8
W
X
Others
c
Northern Europe (N=450)
Western Europe (N=231)
Balkan region (N=169)
Central Europe (N=178)
Central-Eastern Europe (N=300)
Hungary (N=153)
Anatolia/South Caucasus (N=24)
Southern Europe (N=252)
Finland (N=58)
d
Russians Northern (N=144)
Russians Central (N=264)
Russians Central/Southern (N=311)
Slavs (N=1671)
Karelians (N=595)
Veps (N=125)
Finns (N=603)
Latvians (N=411)
Estonians (N=409)
Lithuanian (N=201)
Swede (N=550)
Hungarians (N=116)
Turks (N=478)

**Supplementary Fig. 56. Proportions of mtDNA haplogroups among tested groups.**
(a) Ancient Rus’ regions. (b) Ancient Major Rus’ genetic clusters. (c) Medieval European groups. (d) Present-day European populations, subdivided into several regional groups: Russians Central (from Tula, Vladimir, Novgorod, Pskov), Russians Northern (from Vologda and Arkhangelsk), Russians Central/Southern (from Belgorod, Orel, Stavropol, Saratov), present-day Slavic individuals (Poles, Ukrainians, Belarusians, Slovaks, Czechs), and present-day neighbouring European groups (Karelians, Veps, Finns, Latvians, Estonians, Lithuanians, Swedes, Hungarians, Turks). See Supplementary Informations S2.7.1.

a

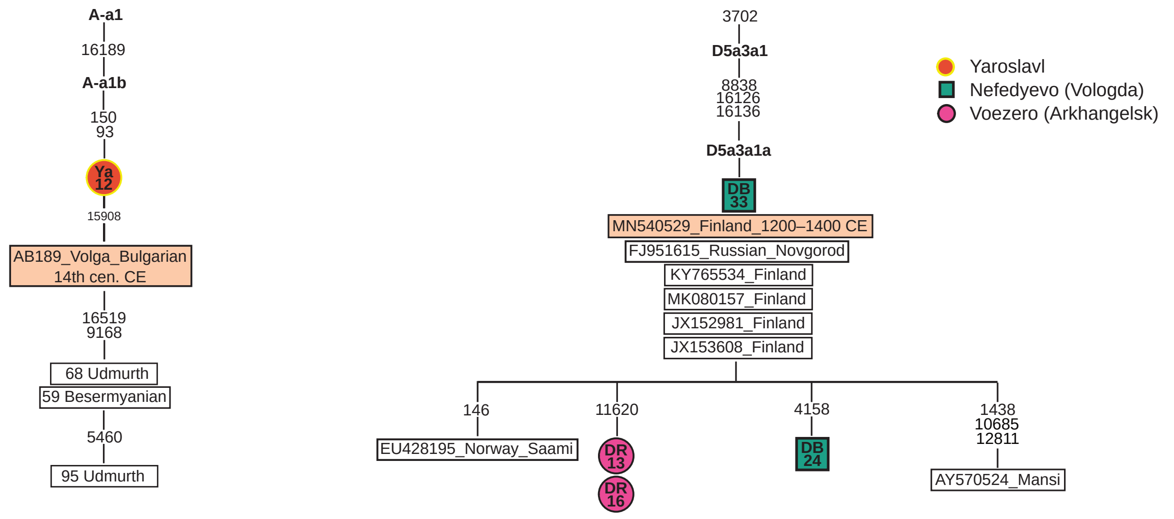

A-a1
16189
A-a1b
150
93
Ya 12
15908
AB189_Volga_Bulgarian 14th cen. CE
16519
9168
68 Udmurth
59 Besermyanian
5460
95 Udmurth
3702
D5a3a1
8838
16126
16136
D5a3a1a
DB 33
MN540529_Finland_1200–1400 CE
FJ951615_Russian_Novgorod
KY765534_Finland
MK080157_Finland
JX152981_Finland
JX153608_Finland
146
EU428195_Norway_Saami
11620
DR 13
DR 16
4158
DB 24
1438
10685
12811
AY570524_Mansi
Yaroslavl
Nefedyevo (Vologda)
Voezero (Arkhangelsk)


b

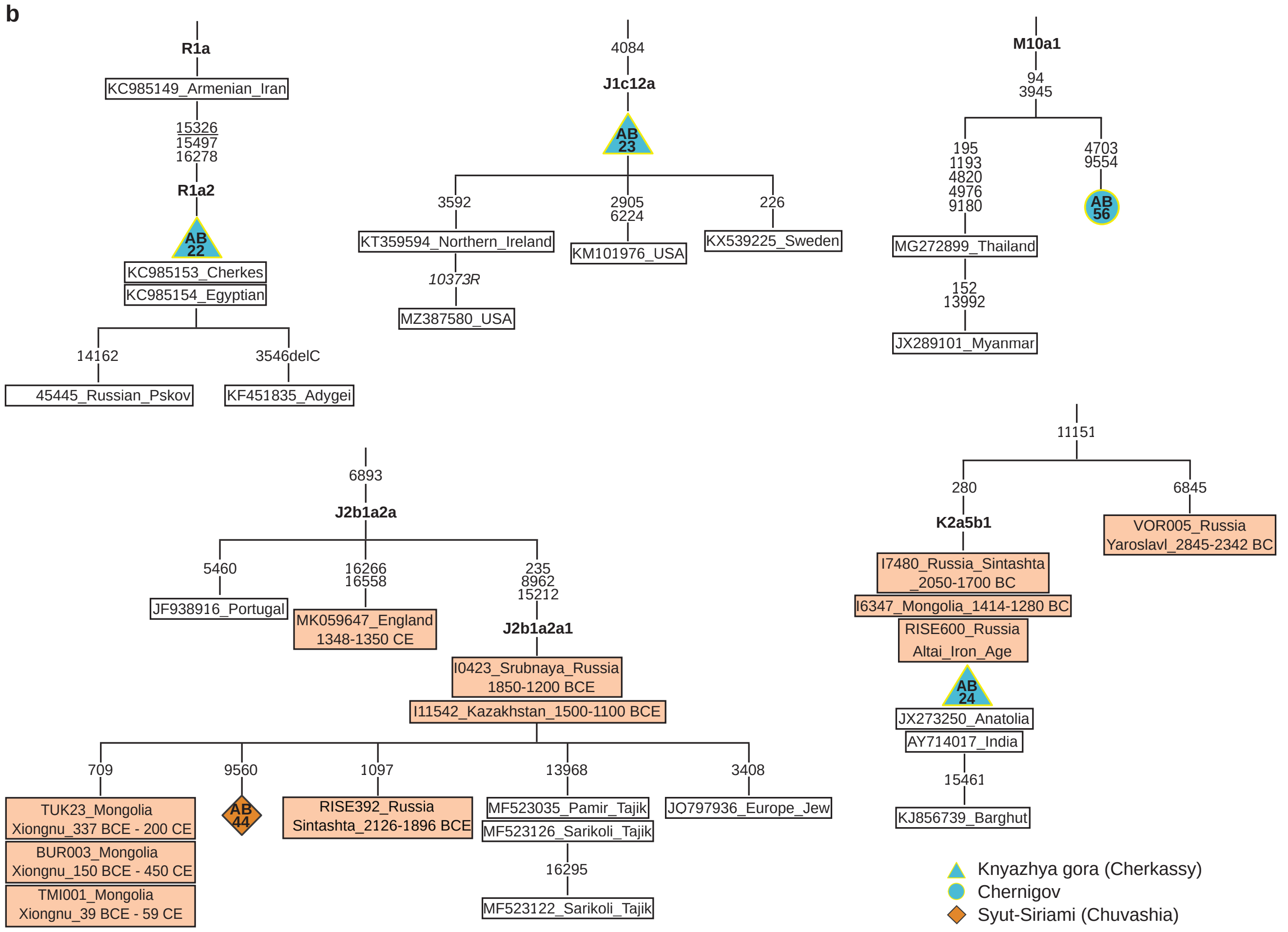

R1a
KC985149_Armenian_Iran
15326
15497
16278
R1a2
AB 22
KC985153_Cherkes
KC985154_Egyptian
14162
45445_Russian_Pskov
3546delC
KF451835_Adygei
4084
J1c12a
AB 23
3592
KT359594_Northern_Ireland
10373R
MZ387580_USA
2905
6224
KM101976_USA
226
KX539225_Sweden
M10a1
94
3945
195
1193
4820
4976
9180
MG272899_Thailand
152
13992
JX289101_Myanmar
4703
9554
AB 56
6893
J2b1a2a
5460
JF938916_Portugal
16266
16558
MK059647_England 1348-1350 CE
235
8962
15212
J2b1a2a1
I0423_Srubnaya_Russia 1850-1200 BCE
I11542_Kazakhstan_1500-1100 BCE
709
TUK23_Mongolia Xiongnu_337 BCE - 200 CE
BUR003_Mongolia Xiongnu_150 BCE - 450 CE
TMI001_Mongolia Xiongnu_39 BCE - 59 CE
9560
AB 44
1097
RISE392_Russia Sintashta_2126-1896 BCE
13968
MF523035_Pamir_Tajik
MF523126_Sarikoli_Tajik
16295
MF523122_Sarikoli_Tajik
3408
JO797936_Europe_Jew
11151
280
K2a5b1
I7480_Russia_Sintashta 2050-1700 BC
I6347_Mongolia_1414-1280 BC
RISE600_Russia Altai_Iron_Age
AB 24
JX273250_Anatolia
AY714017_India
15461
KJ856739_Barghut
6845
VOR005_Russia Yaroslavl_2845-2342 BC
Knyazhya gora (Cherkassy)
Chernigov
Syut-Siriami (Chuvashia)

**Supplementary Fig. 57. Phylogeographic analysis of whole mtDNA genome sequences for outliers and AB22.**

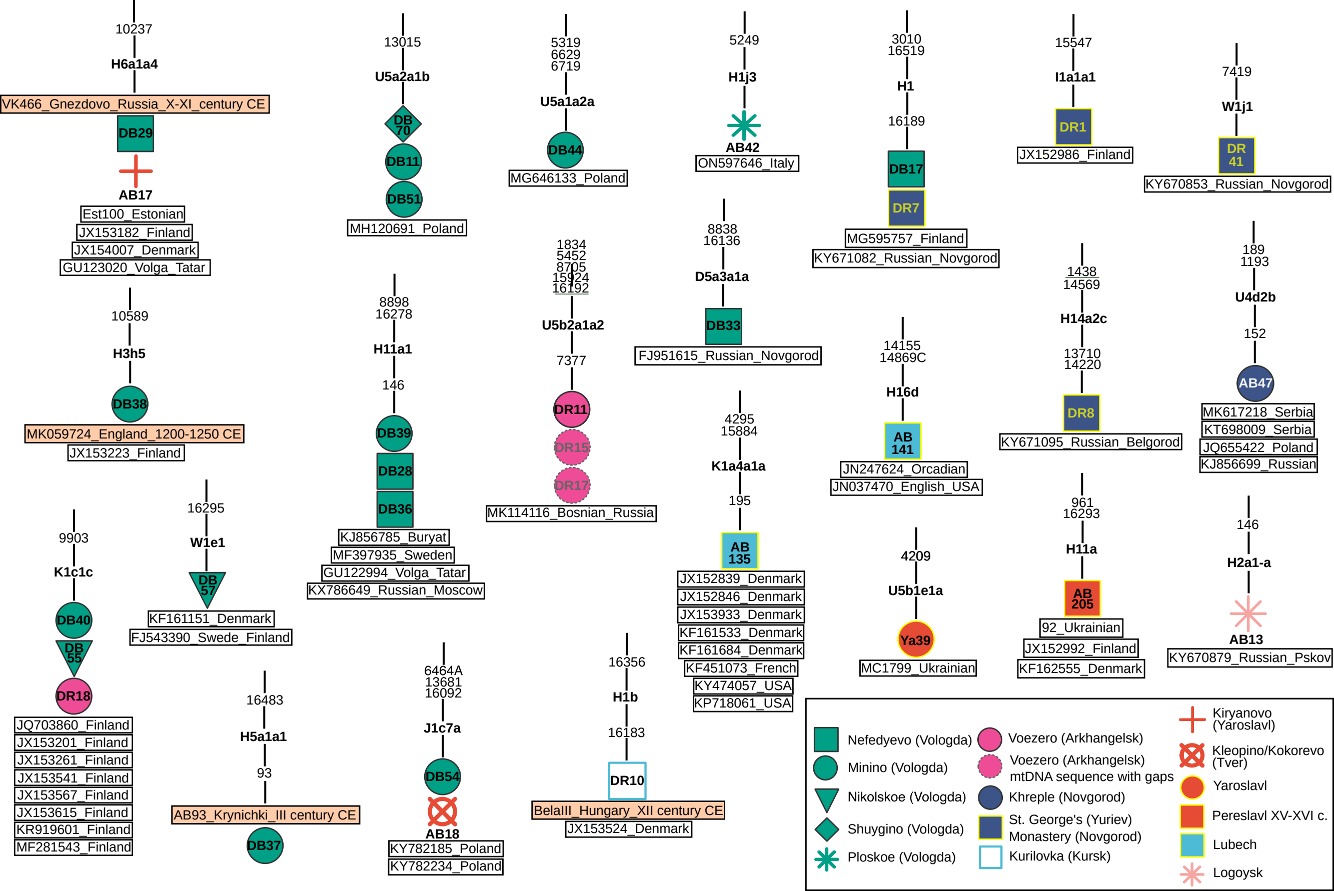
10237
H6a1a4
VK466_Gnezdovo_Russia_X-XI_century CE
DB29
AB17
Est100_Estonian
JX153182_Finland
JX154007_Denmark
GU123020_Volga_Tatar
10589
H3h5
DB38
MK059724_England_1200-1250 CE
JX153223_Finland
9903
K1c1c
DB40
DB 55
DR18
JQ703860_Finland
JX153201_Finland
JX153261_Finland
JX153541_Finland
JX153567_Finland
JX153615_Finland
KR919601_Finland
MF281543_Finland
16295
W1e1
DB 57
KF161151_Denmark
FJ543390_Swede_Finland
16483
H5a1a1
93
AB93_Krynichki_III century CE
DB37
13015
U5a2a1b
DB 70
DB11
DB51
MH120691_Poland
8898
16278
H11a1
146
DB39
DB28
DB36
KJ856785_Buryat
MF397935_Sweden
GU122994_Volga_Tatar
KX786649_Russian_Moscow
6464A
13681
16092
J1c7a
DB54
AB18
KY782185_Poland
KY782234_Poland
5319
6629
6719
U5a1a2a
DB44
MG646133_Poland
1834
5452
8705
15924
16192
U5b2a1a2
7377
DR11
DR15
DR17
MK114116_Bosnian_Russia
16356
H1b
16183
DR10
BelaIII_Hungary_XII century CE
JX153524_Denmark
5249
H1j3
AB42
ON597646_Italy
8838
16136
D5a3a1a
DB33
FJ951615_Russian_Novgorod
4295
15884
K1a4a1a
195
AB 135
JX152839_Denmark
JX152846_Denmark
JX153933_Denmark
KF161533_Denmark
KF161684_Denmark
KF451073_French
KY474057_USA
KP718061_USA
3010
16519
H1
16189
DB17
DR7
MG595757_Finland
KY671082_Russian_Novgorod
14155
14869C
H16d
AB 141
JN247624_Orcadian
JN037470_English_USA
4209
U5b1e1a
Ya39
MC1799_Ukrainian
15547
I1a1a1
DR1
JX152986_Finland
1438
14569
H14a2c
13710
14220
DR8
KY671095_Russian_Belgorod
961
16293
H11a
AB 205
92_Ukrainian
JX152992_Finland
KF162555_Denmark
7419
W1j1
DR 41
KY670853_Russian_Novgorod
189
1193
U4d2b
152
AB47
MK617218_Serbia
KT698009_Serbia
JQ655422_Poland
KJ856699_Russian
146
H2a1-a
AB13
KY670879_Russian_Pskov
Nefedyevo (Vologda)
Minino (Vologda)
Nikolskoe (Vologda)
Shuygino (Vologda)
Ploskoe (Vologda)
Voezero (Arkhangelsk)
Voezero (Arkhangelsk) mtDNA sequence with gaps
Khreple (Novgorod)
St. George's (Yuriev) Monastery (Novgorod)
Kurilovka (Kursk)
Kiryanovo (Yaroslavl)
Kleopino/Kokorevo (Tver)
Yaroslavl
Pereslavl XV-XVI c.
Lubech
Logoysk

**Supplementary Fig. 58. Complete matches of mtDNA sequences between Rus' individuals and previously published ancient and present-day samples.**

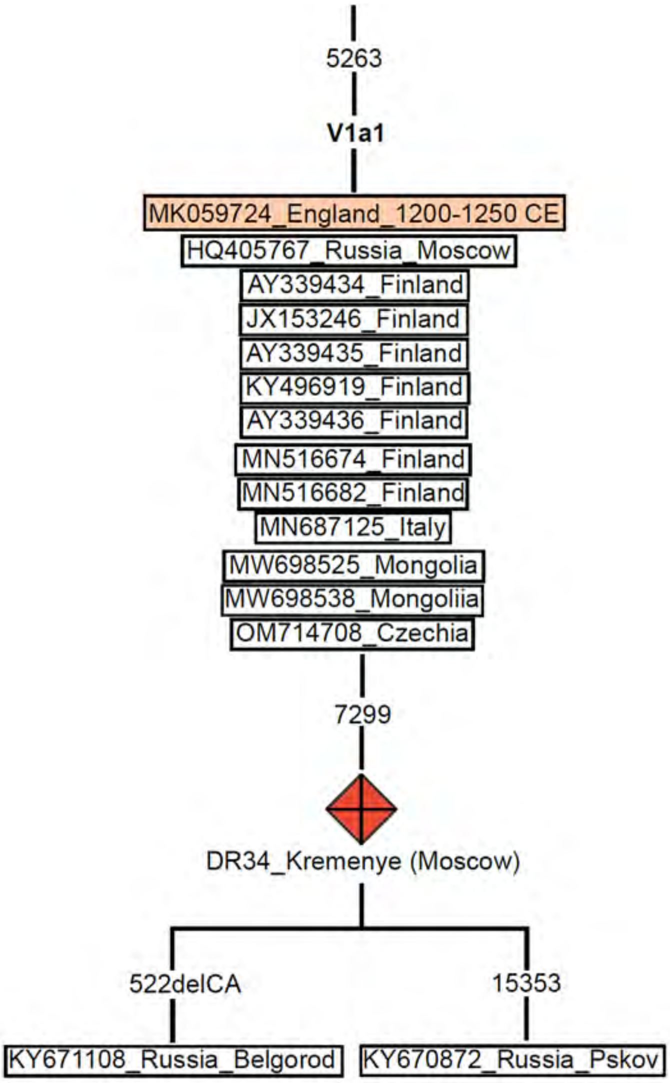


**Supplementary Fig. 59. Fragment of the phylogeographic tree constructed using the mtPhyl program for mtDNA samples of the mitochondrial branch V1a1.**[42]

DR34 is the sample from Vyatichian burial from Kremenye (Moscow), Central Rus' Volga-Oka region. The ancient individual of Viking Era highlighted in light orange. Present-day samples have an identifying number as well as geographic/ethnic information. For transitions, the position number and the substitution variant are indicated; del – deletion.

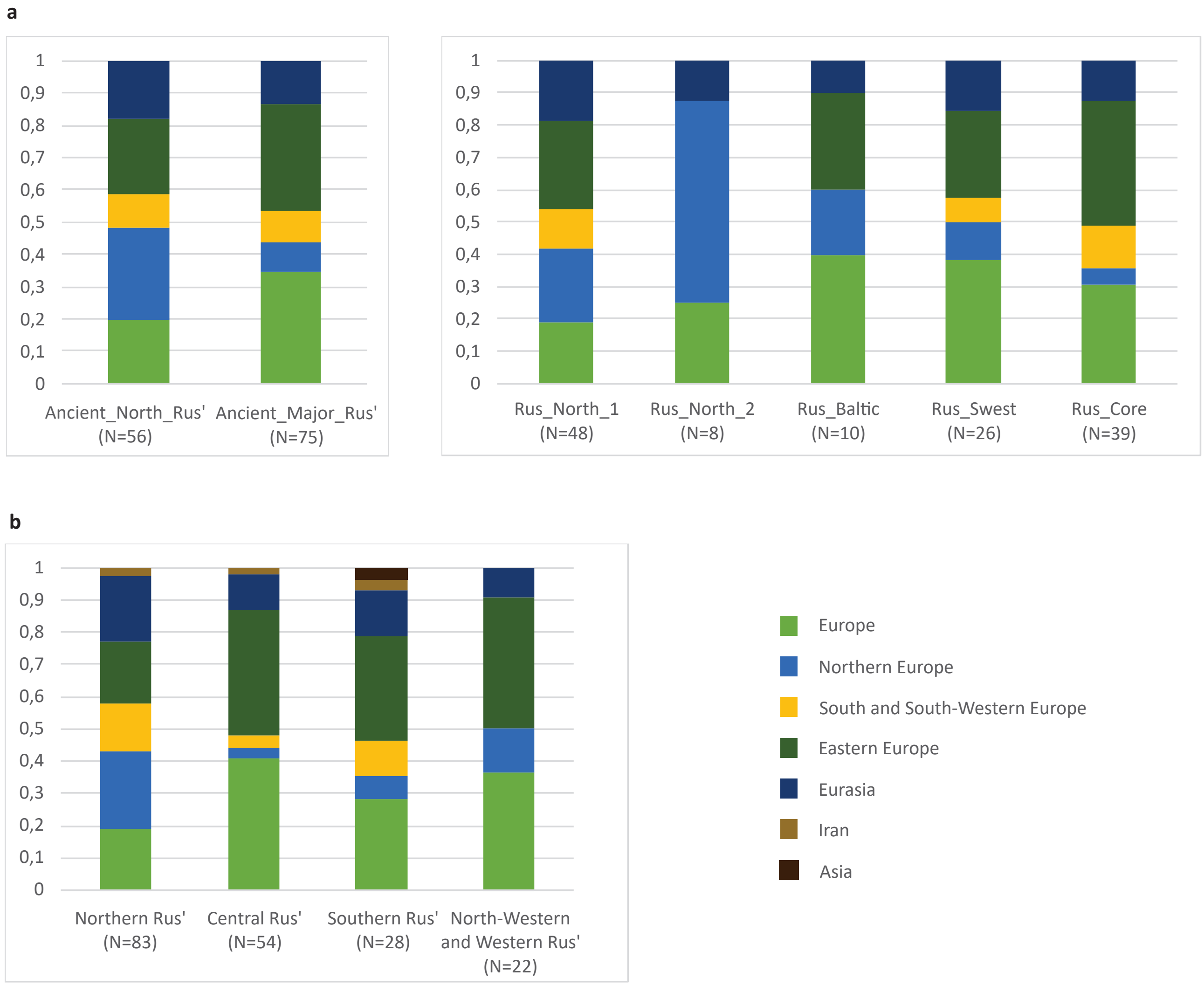

a
1
0,9
0,8
0,7
0,6
0,5
0,4
0,3
0,2
0,1
0
Ancient_North_Rus' (N=56)
Ancient_Major_Rus' (N=75)
Rus_North_1 (N=48)
Rus_North_2 (N=8)
Rus_Baltic (N=10)
Rus_Swest (N=26)
Rus_Core (N=39)
b
Northern Rus' (N=83)
Central Rus' (N=54)
Southern Rus' (N=28)
North-Western and Western Rus' (N=22)
Europe
Northern Europe
South and South-Western Europe
Eastern Europe
Eurasia
Iran
Asia

**Supplementary Fig. 60. Present-day distribution of mtDNA haplogroups identified among the ancient Rus’ genetic clusters (a) and ancient Rus’ geographical regions (b).**

Europe - the whole of Western Eurasia area. Northern Europe - Scandinavia, the British Isles, Finland, and the northern area of the European part of Russia. South and South-Western Europe - France, Spain, Italy, and Turkey. Eastern Europe - European part of Russia, Czechia, Poland, Serbia, Bulgaria, Slovakia, Ukraine, Romania, Lithuania, etc. Eurasia - the whole of Eurasia. Asia - the whole Eastern Eurasia. Iran - Iranian, Caucasian, and the Middle East regions.

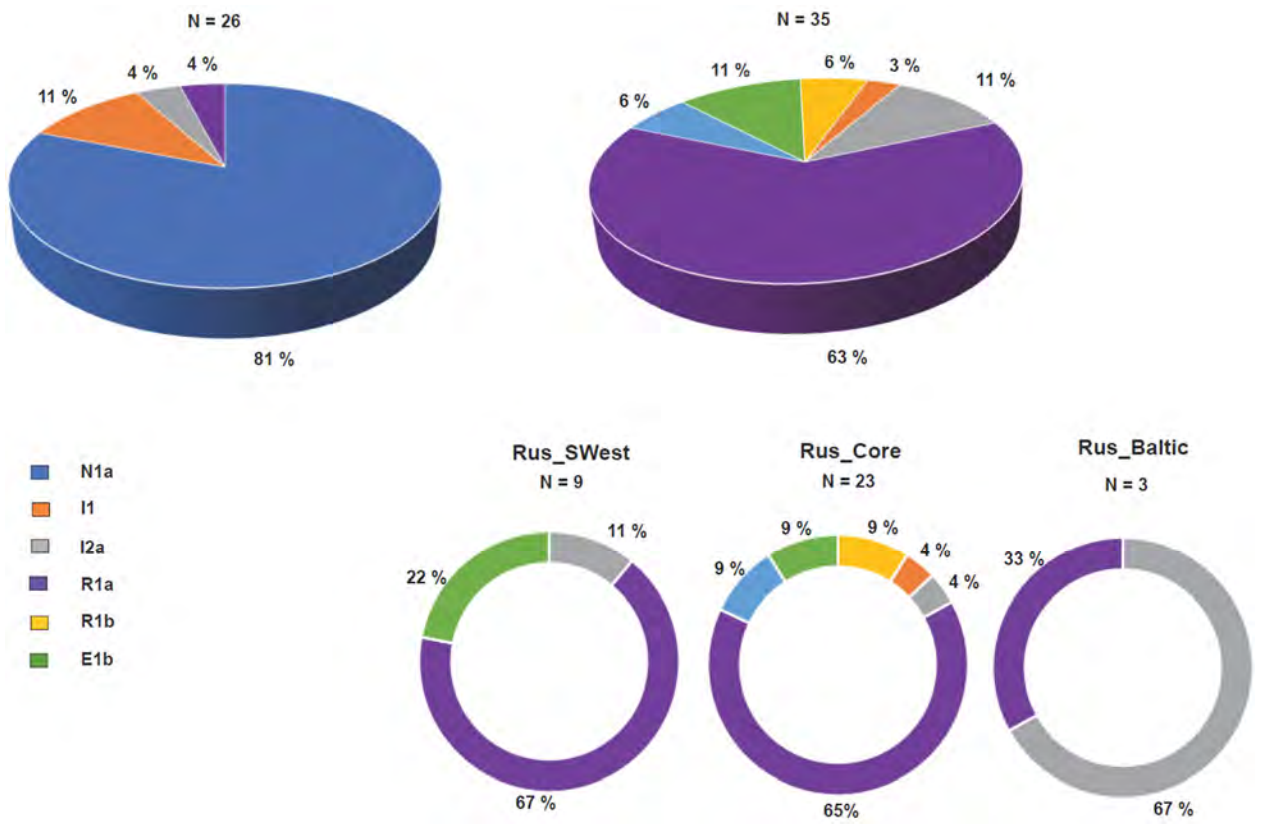


**Supplementary Fig. 61. The distribution of Y-chromosome haplogroups within the Ancient North Rus' and Ancient Major Rus' genetic clusters (top), and among the Rus_SWest, Rus_Core, and Rus_Baltic genetic subclusters (bottom).**

### Phylogenetic tree of haplogroup N1a-M231

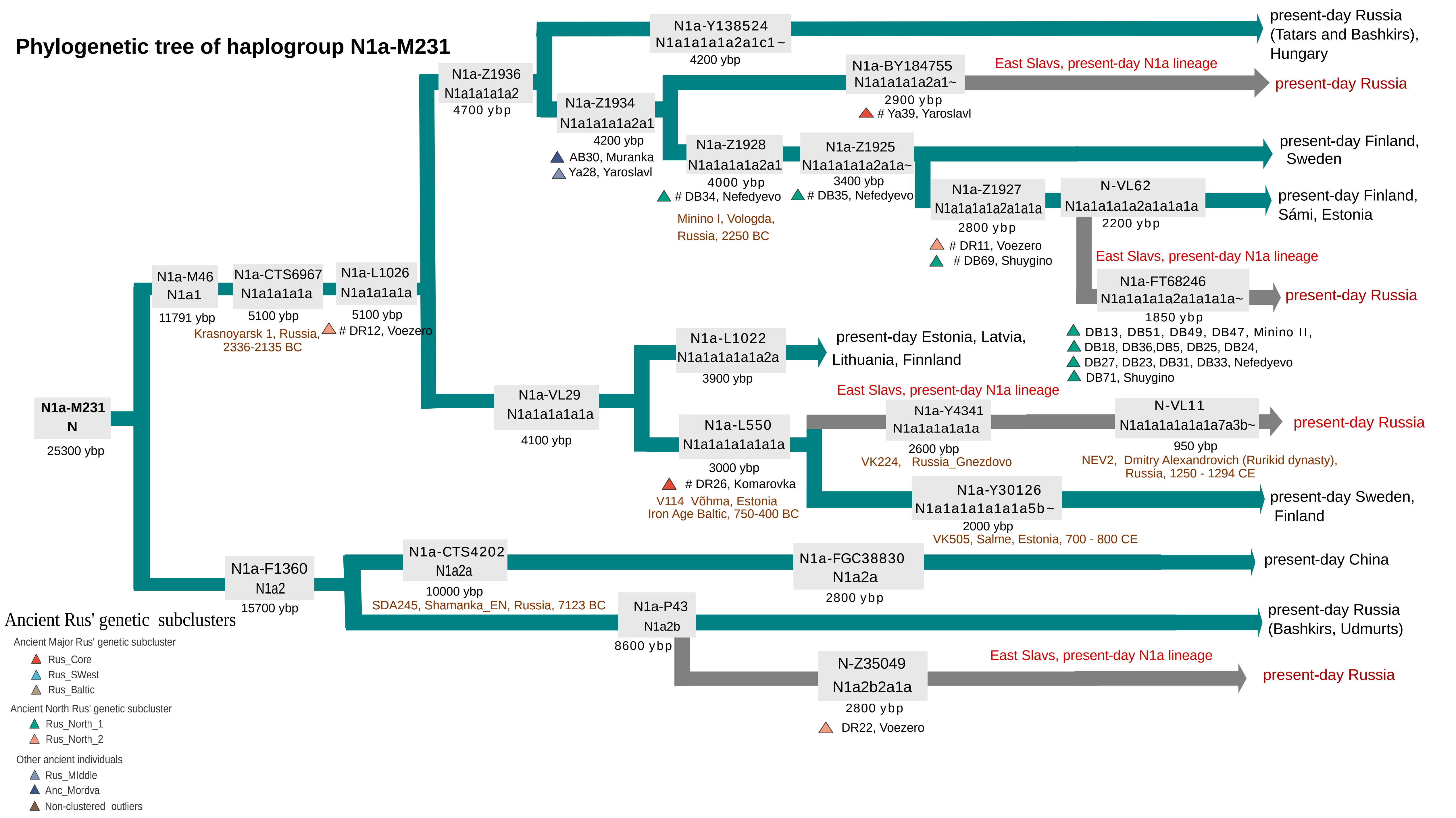

**Supplementary Fig. 62. The simplified Y-chromosome phylogenetic trees for haplogroup N1a lineages identified among newly reported ancient Rus' individuals.**
The mean age of branches is indicated according to the Yfull database (www.yfull.com). Previously published ancient samples are signed and highlighted in brown[102,136,138,140,162]. The shape and color of the symbols for the newly reported individuals correspond to the Fig. 3. The N1a paternal lineage, currently observed in East Slavic populations, is highlighted in gray.
The current study has identified three main branches of the N1a haplogroup in the samples analyzed. The first branch, designated N-Z1936, is found in all samples from the Rus_North_1 cluster. The second branch, N-VL29, is exclusively present in samples from the Ancient Major Rus' genetic cluster, and the third branch, N-P43, is identified in the Rus_North_2 genetic subcluster.
Two unique distinct haplogroups, N-FT68246 and N-Z35049, have been identified among the medieval populations of Northern Rus' (Beloozero and Voezero regions). These haplogroups are currently distributed only among the present-day Russian population. Additionally, another "East Slavic"-specific paternal lineage, N-BY184755, has been identified among the medieval Slavic-related individuals of ancient Rus', with two individuals from Yaroslavl representing this lineage.

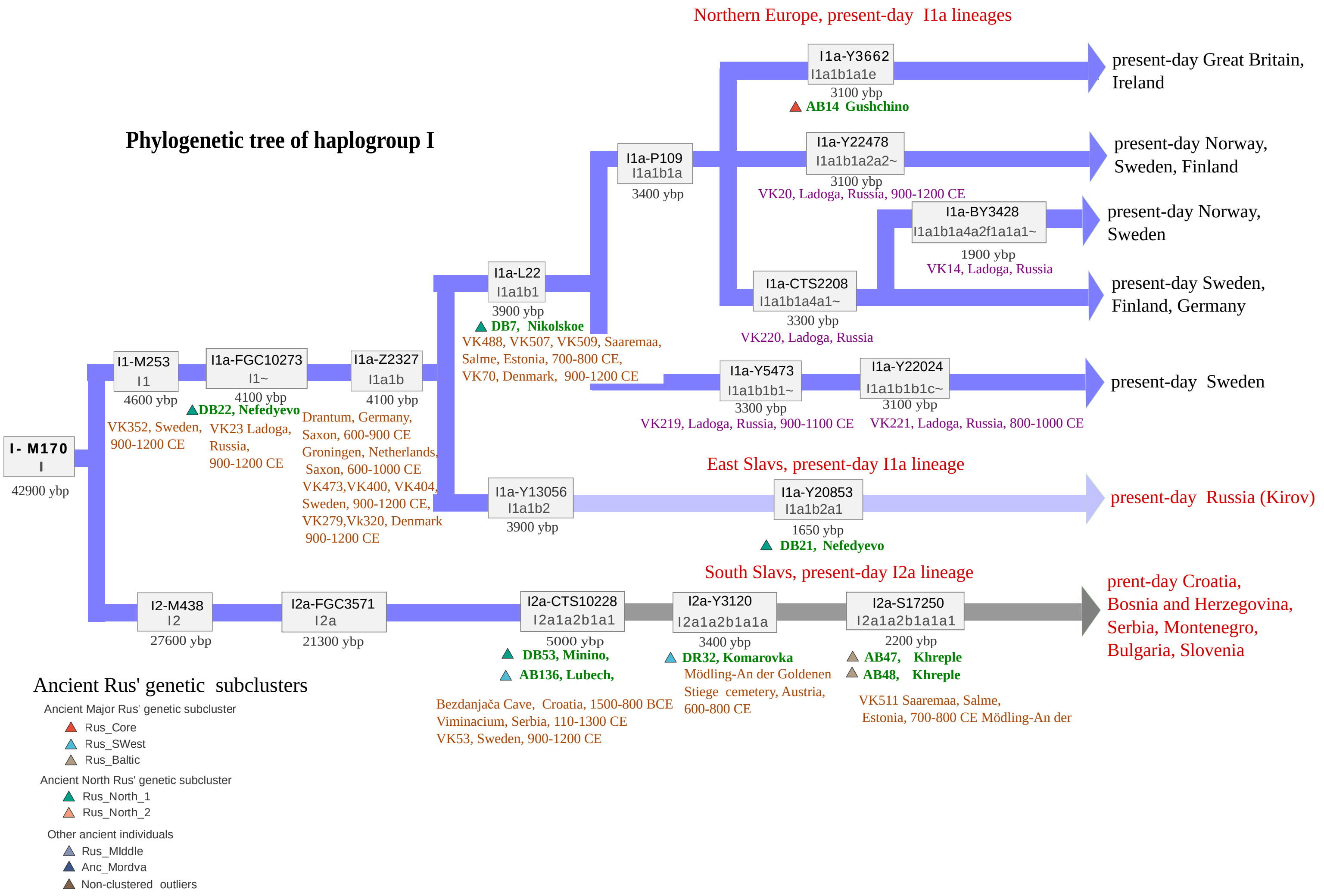

Phylogenetic tree of haplogroup I
Northern Europe, present-day I1a lineages
I- M170
I
42900 ybp
I1-M253
I1
4600 ybp
VK352, Sweden, 900-1200 CE
I1a-FGC10273
I1~
4100 ybp
DB22, Nefedyevo
VK23 Ladoga, Russia, 900-1200 CE
I1a-Z2327
I1a1b
4100 ybp
Drantum, Germany, Saxon, 600-900 CE
Groningen, Netherlands, Saxon, 600-1000 CE
VK473,VK400, VK404, Sweden, 900-1200 CE,
VK279,Vk320, Denmark 900-1200 CE
I1a-L22
I1a1b1
3900 ybp
DB7, Nikolskoe
VK488, VK507, VK509, Saaremaa, Salme, Estonia, 700-800 CE,
VK70, Denmark, 900-1200 CE
I1a-P109
I1a1b1a
3400 ybp
I1a-Y3662
I1a1b1a1e
3100 ybp
AB14 Gushchino
present-day Great Britain, Ireland
I1a-Y22478
I1a1b1a2a2~
3100 ybp
VK20, Ladoga, Russia, 900-1200 CE
present-day Norway, Sweden, Finland
I1a-BY3428
I1a1b1a4a2f1a1a1~
1900 ybp
VK14, Ladoga, Russia
present-day Norway, Sweden
I1a-CTS2208
I1a1b1a4a1~
3300 ybp
VK220, Ladoga, Russia
present-day Sweden, Finland, Germany
I1a-Y5473
I1a1b1b1~
3300 ybp
VK219, Ladoga, Russia, 900-1100 CE
I1a-Y22024
I1a1b1b1c~
3100 ybp
VK221, Ladoga, Russia, 800-1000 CE
present-day Sweden
East Slavs, present-day I1a lineage
I1a-Y13056
I1a1b2
3900 ybp
I1a-Y20853
I1a1b2a1
1650 ybp
DB21, Nefedyevo
present-day Russia (Kirov)
South Slavs, present-day I2a lineage
I2-M438
I2
27600 ybp
I2a-FGC3571
I2a
21300 ybp
I2a-CTS10228
I2a1a2b1a1
5000 ybp
DB53, Minino,
AB136, Lubech,
Bezdanjača Cave, Croatia, 1500-800 BCE
Viminacium, Serbia, 110-1300 CE
VK53, Sweden, 900-1200 CE
I2a-Y3120
I2a1a2b1a1a
3400 ybp
DR32, Komarovka
Mödling-An der Goldenen Stiege cemetery, Austria, 600-800 CE
I2a-S17250
I2a1a2b1a1a1
2200 ybp
AB47, Khreple
AB48, Khreple
VK511 Saaremaa, Salme, Estonia, 700-800 CE Mödling-An der
prent-day Croatia, Bosnia and Herzegovina, Serbia, Montenegro, Bulgaria, Slovenia
Ancient Rus' genetic subclusters
Ancient Major Rus' genetic subcluster
Rus_Core
Rus_SWest
Rus_Baltic
Ancient North Rus' genetic subcluster
Rus_North_1
Rus_North_2
Other ancient individuals
Rus_MIddle
Anc_Mordva
Non-clustered outliers

**Supplementary Fig. 63. The simplified Y-chromosome phylogenetic tree for haplogroup I lineages identified among newly reported ancient Rus' individuals.**

The mean age of branches is indicated according to the Yfull database (www.yfull.com). Newly reported ancient samples are highlighted in green. Previously published samples from the Viking Era and medieval burials in Europe are indicated in [102,107,163]. Purple marks previously published samples from Viking Era burials from the territory of Rus’. The shape and color of the symbols for the newly reported individuals correspond to those in Fig. 3. The sub-branch I-Y13056 (I1a1b2), which is currently found in East Slavic populations (mainly in Russia), is highlighted in light blue. The I2a paternal lineage now present among South Slavs is highlighted in gray.

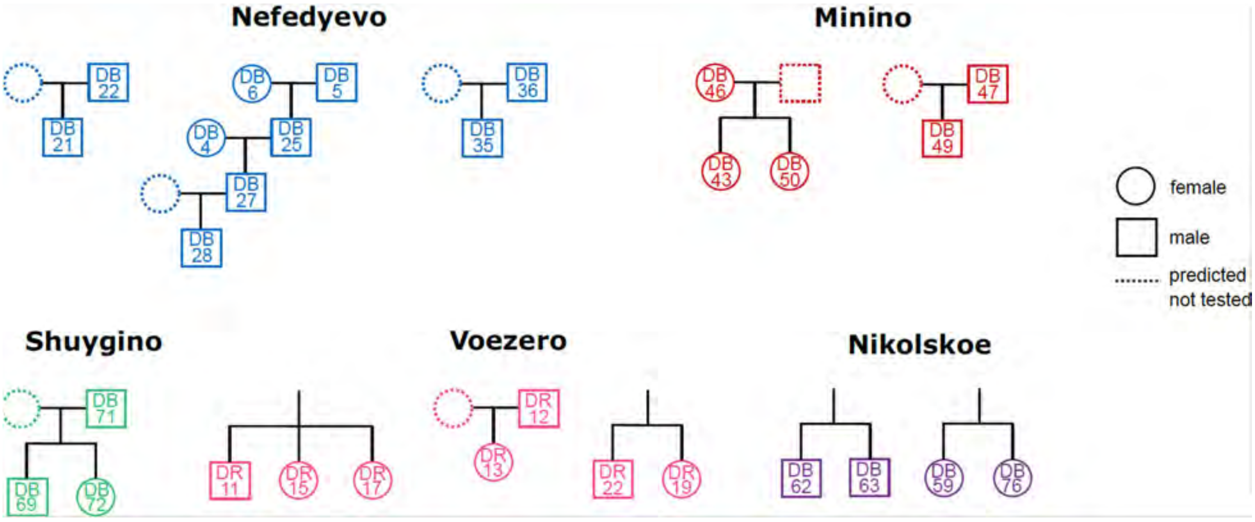


**Supplementary Fig. 64. Small pedigrees reconstructed from pairs of the first-degree relatives identified in our Rus' cohort.**

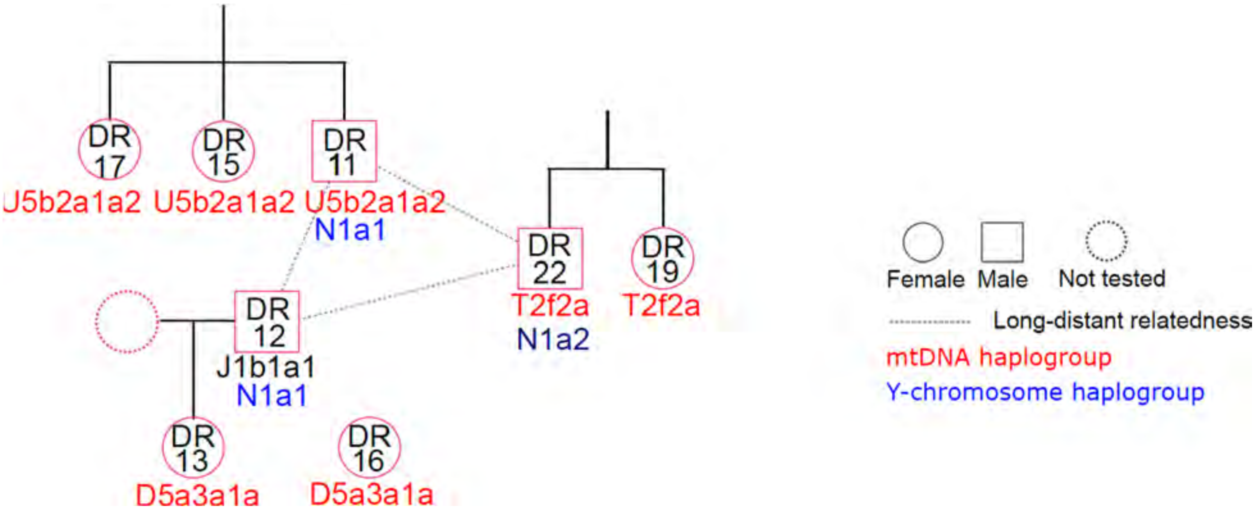


**Supplementary Fig. 65. Kinship relationships identified among individuals from the Voezero burial site in the Northern Rus' region.**

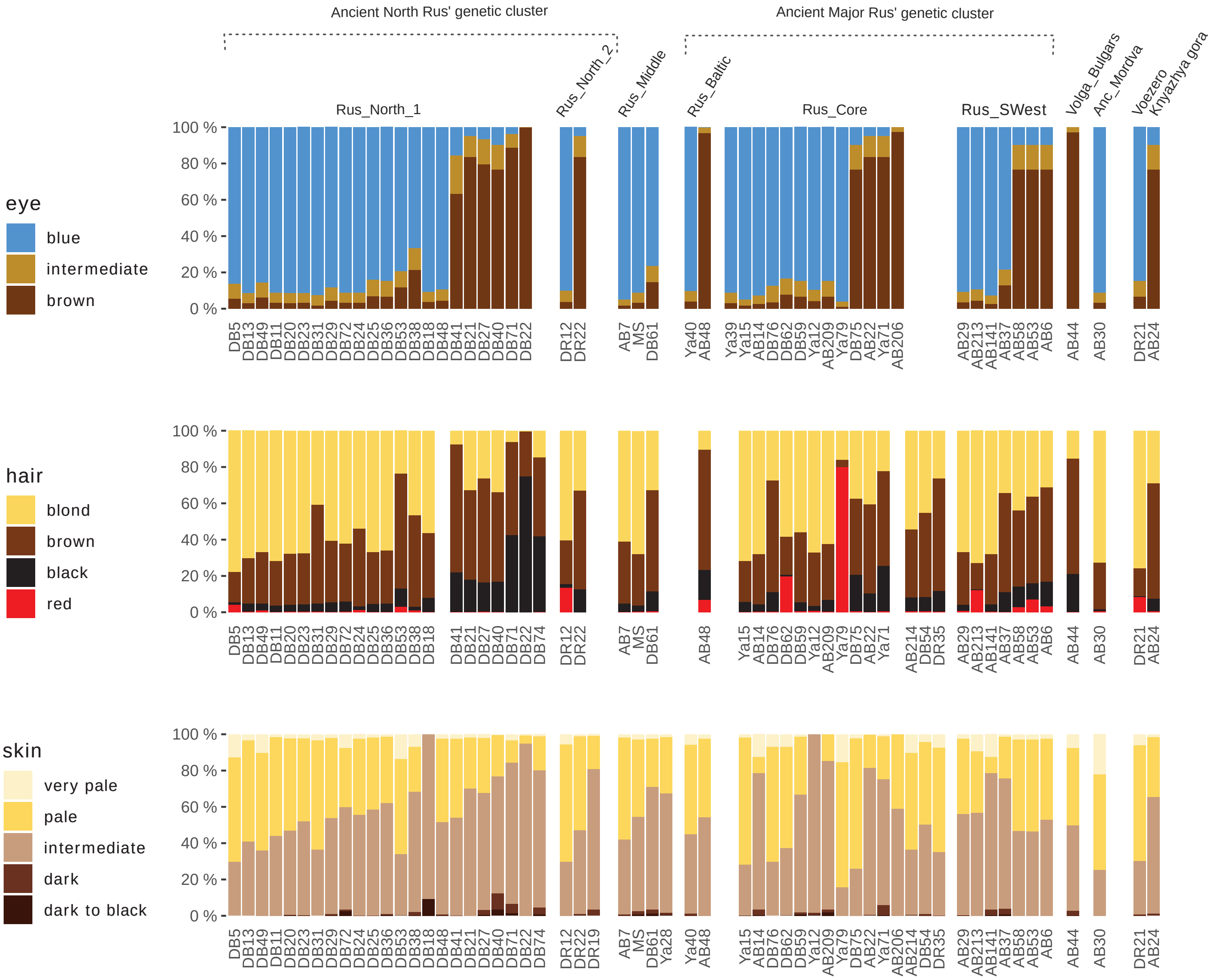

Ancient North Rus' genetic cluster
Ancient Major Rus' genetic cluster
Rus_North_1
Rus_North_2
Rus_Middle
Rus_Baltic
Rus_Core
Rus_SWest
Volga_Bulgars
Anc_Mordva
Voezero
Knyazhya gora
eye
blue
intermediate
brown
hair
blond
brown
black
red
skin
very pale
pale
intermediate
dark
dark to black
100 %
80 %
60 %
40 %
20 %
0 %

**Supplementary Figure 66. Color prediction for eyes, hair, and skin color with HIrisPlex-S tool. Prediction probability calculated** for three categories of eye color (blue, intermediate, and brown), four of hair color (black, brown, blonde, or red), and five of skin color (very pale, intermediate, dark, and dark to black). See details and losses in AUC (the Area Under the receiver operating characteristic Curve) accuracy in Supplementary Table 33.

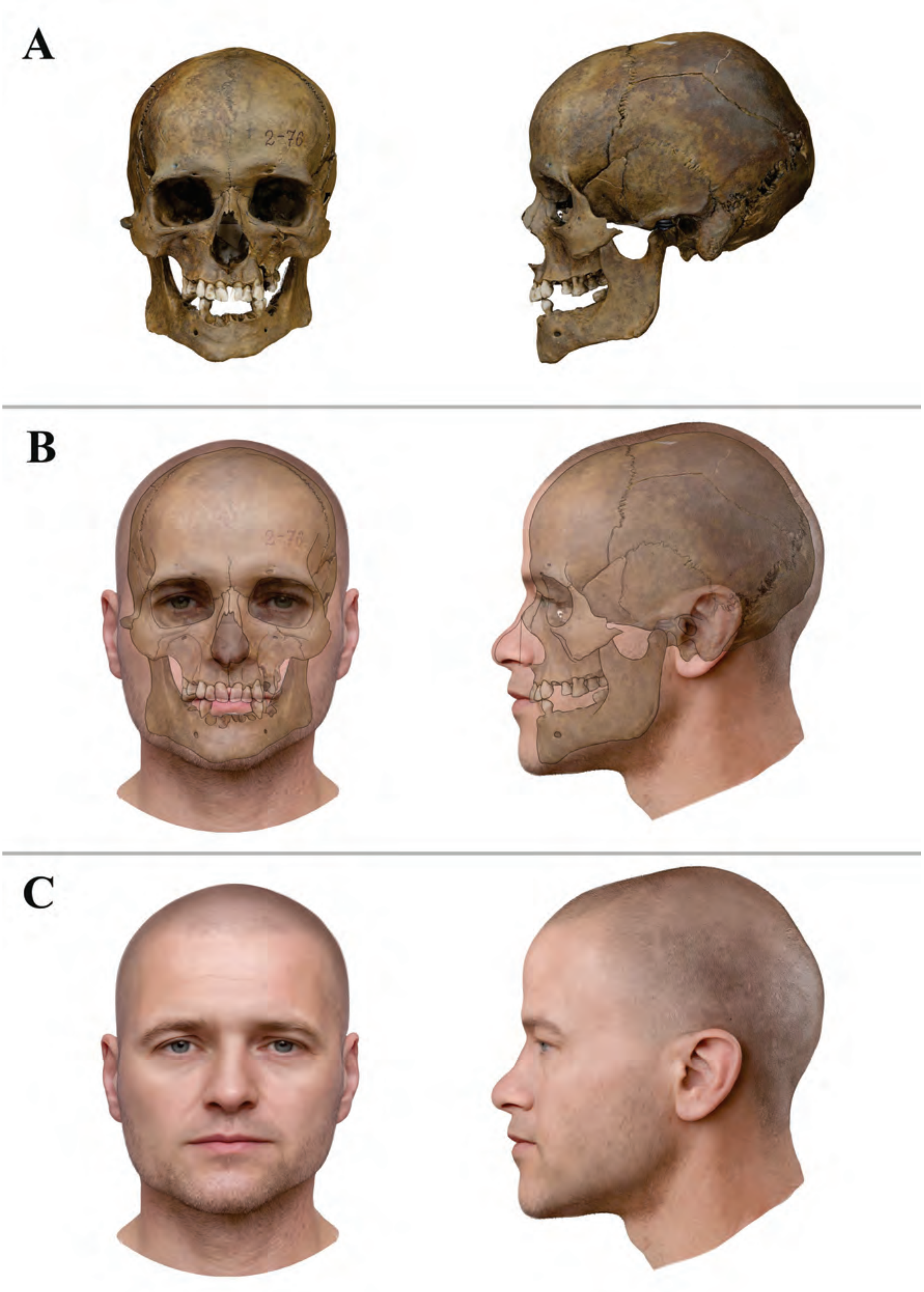
A
B
C

**Supplementary Figure 67. Stages of craniofacial reconstruction of individual MS from the Yaroslavl mass grave, 13th century.**
(a) Skull in frontal and lateral views; (b)Superimposition of the skull image on the face; (c) Completed craniofacial reconstruction.

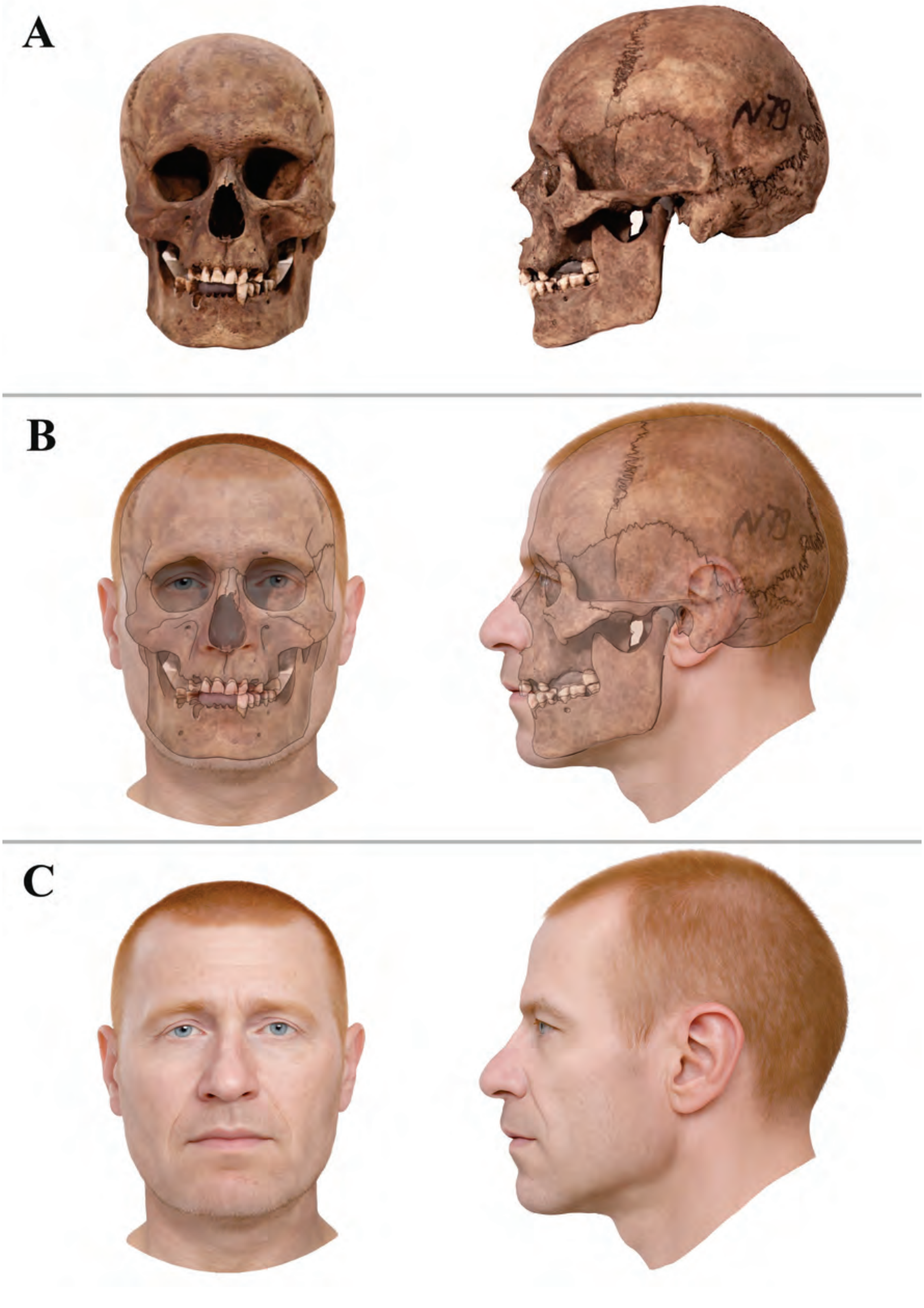
A
B
C

**Supplementary Figure 68. Stages of craniofacial reconstruction of individual Ya39 from the Yaroslavl mass grave, 13th century.**

(a) Skull in frontal and lateral views; (b)Superimposition of the skull image on the face; (c) Completed craniofacial reconstruction.

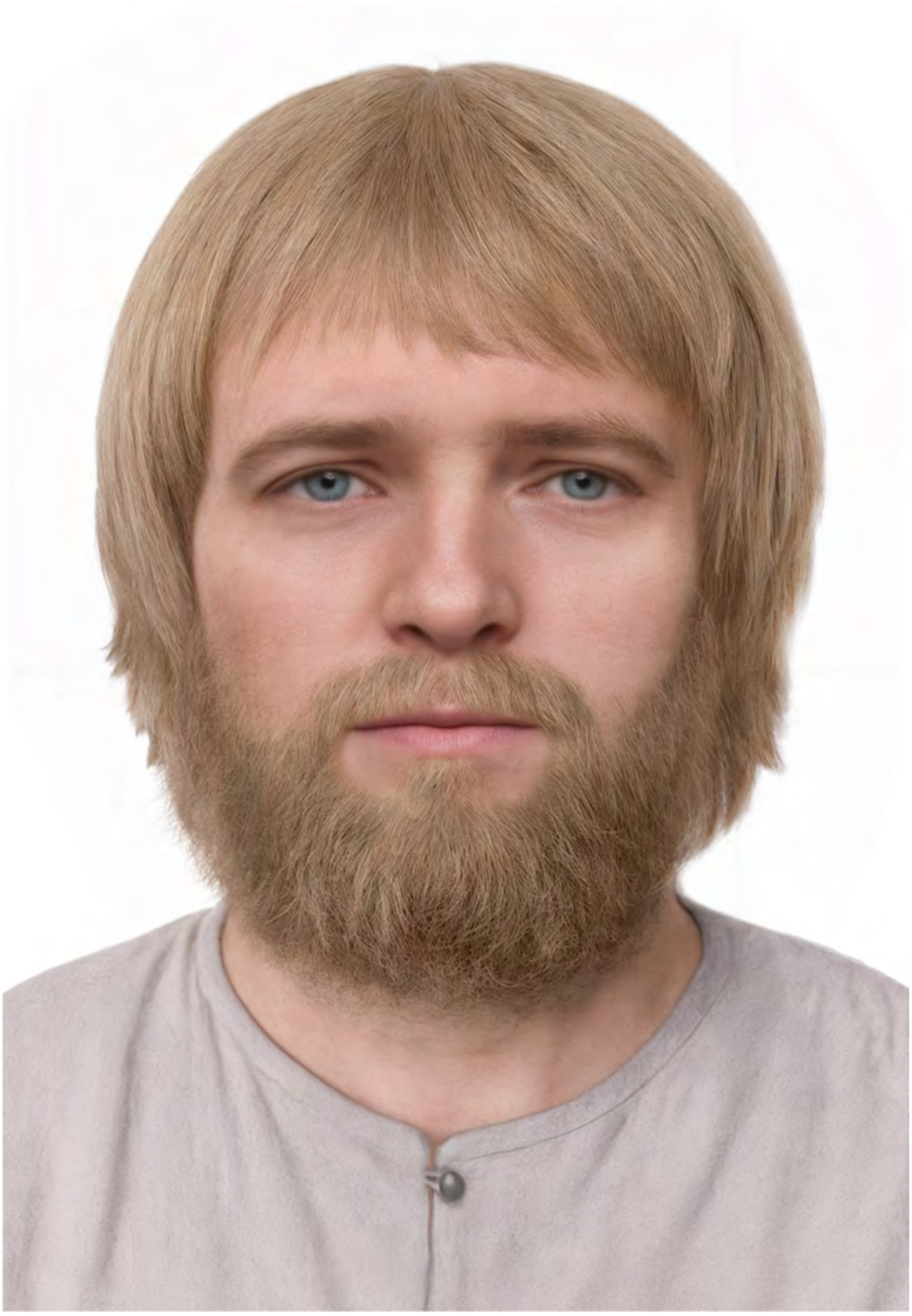

**Supplementary Figure 69. Craniofacial reconstruction with beard and hairstyle for individual MS from the Yaroslavl mass grave of the 13th century.**

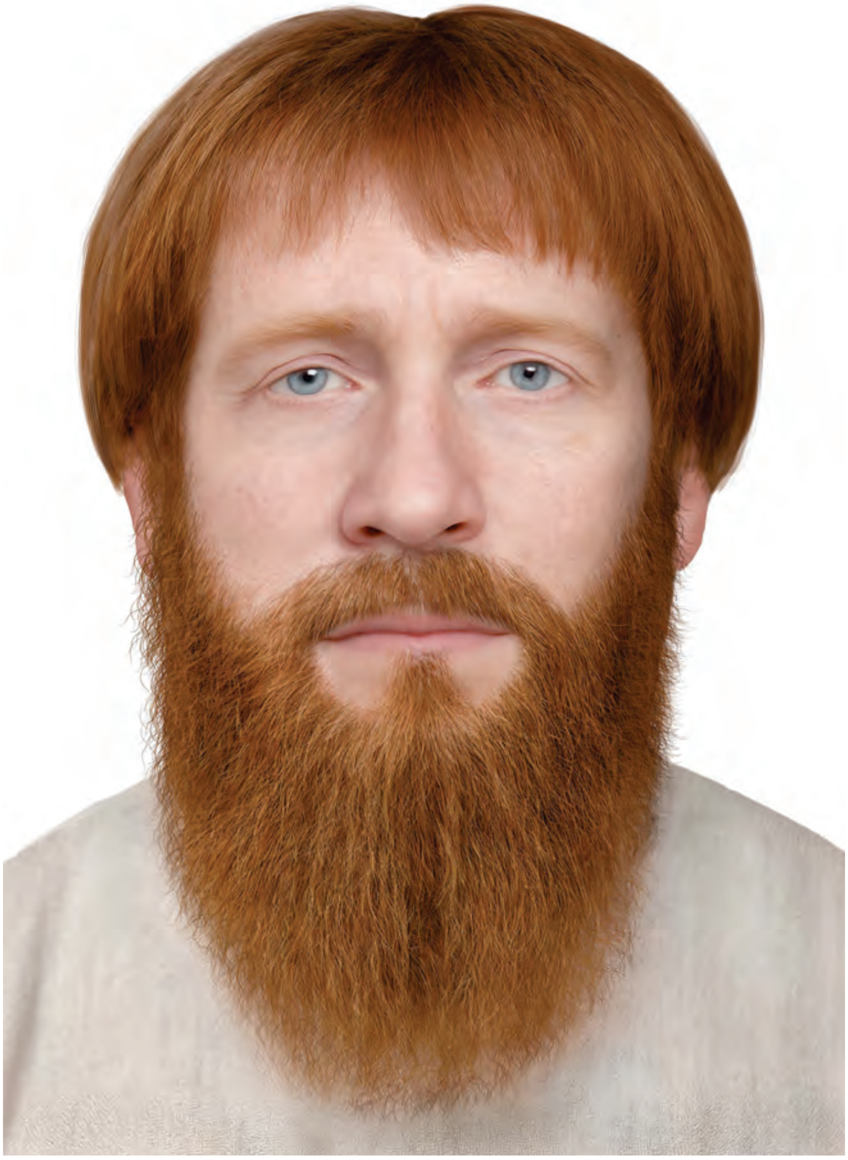

**Supplementary Figure 70. Craniofacial reconstruction with beard and hairstyle for individual Ya39 from the Yaroslavl mass grave of the 13th century.**
