## Supplementary Materials for "Genetic history of Rus’"

Tatiana V. Andreeva *et al.*

**The PDF file includes:**

Supplementary Information S1 and S2

Supplementary Figs. 1 to 70

Legends for Supplementary Tables 1 to 35

References

**Other Supplementary Material for this manuscript includes the following:**

Supplementary Tables 1 to 35

**S1. The archaeological and anthropological context of the samples	5**

**S1.1. Archaeological sites of the Northern Rus' region	9**

S1.1.1. Burial sites of Beloozero region (Vologda)	9

S1.1.1.1. The Nefedyevo burial site	9

S1.1.1.2. The Minino burial site	23

S1.1.1.3. The Shuygino burial site	33

S1.1.1.4. The Nikolskoe burial site	36

S1.1.2. Burial grounds of the Poonezhye region (Arkhangelsk)	43

S1.1.2.1. The Voezero burial site	43

S1.1.3. The Novinki burial site (Vologda)	48

S1.1.4. The Ploskoe burial site (Vologda)	49

**S1.2 Archaeological sites of other Rus’ regions	51**

**S1.2.1. Burial grounds from the Central Rus' Volga-Oka region	51**

S1.2.1.1. The Gnezdilovo burial site (Suzdal)	51

S1.2.1.2. The Luzhki burial site (Moscow)	56

S1.2.1.3. The Akatovo burial site (Moscow)	57

S1.2.1.4. The Velikoe burial site (Vladimir)	59

S1.2.1.5. The Vorobyovo burial site (Tver)	60

S1.2.1.6. The Voronovo burial site (Yaroslavl)	61

S1.2.1.7. The Zhukovo burial site (Yaroslavl)	62

S1.2.1.8. The Ziminki burial site (Murom)	63

S1.2.1.9. The Kiryanovo burial site (Yaroslavl)	64

S1.2.1.10. The Kleopino/Kokorevo burial site (Tver)	65

S1.2.1.11. The Kletnevo burial site (Vladimir)	66

S1.2.1.12. The Pustosh Popova burial site (Kostroma)	67

S1.2.1.13.The Kremenye burial site (Moscow)	68

S1.2.1.14. The Fili burial site (Moscow)	69

S1.2.1.15. Yaroslavl Kremlin (Yaroslavl) burial site	70

S1.2.1.16. Pereslavl-Zalessky burial site	73

S.1.2.1.17. Old Ryazan burial site	78

**S1.2.2. Burial grounds from the North-Western Rus' region (Novgorod)	83**

S1.2.2.1. St. George’s (Yuriev) Monastery burial site	83

S1.2.2.2. The Khreple burial site (Novgorod)	90

**S1.2.3. Burial grounds from the Western Rus' Dnieper-Dvina region	93**

S1.2.3.1. The Volochok burial site (Smolensk)	93

S1.2.3.2. The Iput burial site (Bryansk)	94

S1.2.3.3. The Logoysk burial site (Minsk)	95

S1.2.3.4. The Seltso burial site (Tver)	96

S1.2.3.5. The Gnevkovo burial site (Smolensk)	97

**S1.2.4. Burial grounds from the Southern Rus' region	98**

S1.2.4.1. The Komarovka burial site (Kursk)	98

S1.2.4.2. The Kurilovka burial site (Kursk)	103

S1.2.4.3. The Moiseevskoe burial site (Kursk)	104

S1.2.4.4. The Gushchino burial site (Chernigov)	106

S1.2.4.5. The Konotop burial site (Sumy regin)	107

S1.2.4.6. Knyazhya gora burial site (Cherkassy)	107

S1.2.4.7. Burial site in Chernigov	109

S1.2.4.8. Burial site in Lubech	110

S1.2.4.9. Burial sites in Kiev	114

**S1.3. Burial grounds in the areas adjacent to medieval Rus’	116**

S1.3.1. Burial site of the Latgalian culture	116

S1.3.1.1. The Lucinsky burial site (Ludza)	116

S1.3.2. Burial sites of Muroma culture	117

S1.3.2.1. The Podbolotyevsky burial site (Murom)	117

S1.3.3. Burial site of ancient Mordva	120

S1.3.3.1. The Pogiblovo burial site (Nizhny Novgorod)	120

S1.3.3.2. The Muranka burial site (Samara)	121

S1.3.3.3. The Korino burial site (Nizhny Novgorod)	123

S1.3.4. Burial site of Volga Bulgars	125

S1.3.4.1. The Syut-Siriami burial site (Chuvashia)	125

**S1.4. Radiocarbon dating	126**

**S2. Supplementary methods and results	127**

**S2.1. Datasets	127**

**S2.2. PCA analyses results of Rus’ individuals and previously published ancient samples	127**

S2.2.1 Ancient Rus’ towns	130

S2.2.2 Scandinavians and Social military elite (druzhina)	130

S2.2.3 Genetic outliers in Rus’ area	134

**S2.3. f3-, f4-statistics	135**

S2.3.1. Ancient populations	135

S2.3.2. Present-day populations	137

**S2.4. Admixture modeling using qpAdm	140**

**S2.5. Analysis of Runs of Homozygosity	145**

**S2.6. IBD analysis results	146**

**S2.7. Analysis of uniparental markers	150**

S2.7.1. Mitochondrial DNA diversity	150

S2.7.2. Y chromosome haplogroups diversity	156

Ancient Major Rus’ genetic cluster	156

Ancient North Rus’ genetic cluster	162

Y-chromosome lineages in individuals from Non-Rus'/non-Slavic groups	166

Y-chromosome lineages of outlier samples	166

**S2.8. Ancient Rus’ pedigrees reconstruction	167**

S2.8.1. Small pedigree identification based on READv2, IBD results, sex, age and archaeological data	167

S2.8.2. Large pedigrees reconstruction	176

**S2.9. Appearance prediction of ancient Rus' people	183**

S2.9.1. Phenotypes prediction based on genomic data	183

S2.9.2 Graphic craniofacial reconstruction of medieval Rus’ individuals	185

**References	187**

**Legends for Supplementary Tables	2**09

### S1. The archaeological and anthropological context of the samples

The origin and formation of the Rus' people remain a highly debated historical issue. According to a theory based on archaeological evidence and written sources, the various Slavic communities (often referred to as "tribes") in Eastern Europe united to form a common East Slavic Rus’ people and establish the Rus' state by the 10th century CE. This consolidation occurred within the Rus' area, which combined the territories extending from Lake Ladoga to the Middle Dnieper region under the control of princes from the Rurik dynasty^1–5^. The emergence of this state was driven by the Slavic migrations during the late first and early second millennia CE, a process that brought them into contact and interaction with other ethnic groups of the East European Plain. The archaeological data demonstrates that this process gave rise to a distinct Rus' culture, which then expanded throughout a wide region, developing local variations^6,7^. Furthermore, medieval manuscripts demonstrate the growth of the Rus' (ancient Rus') language and the adoption of the Cyrillic script^8^. The archaeogenetic investigation of ancient Rus’ people has been significantly neglected, yielding only a limited number of genetic profiles from burial sites, which offer insufficient data for developing a comprehensive view on the genetic structure and origins of the Rus' inhabitants.

The ethnic diversity of the Rus' people from the 9th to 13th century CE has yet to be explored, and the nature of its early medieval ethnic dynamics remains debated. The author of the *Primary Chronicle*—a principal source on early Rus' history—describes this land as inhabited by distinct Slavic groups (among them Polyanians, Drevlians, Severians, Radimichians, Vyatichians, Krivichians, Ilmen Slovenes, Dregovichians, Polochans, Ulichians, Dulebes, Tivercians, and others), as well as non-Slavic groups, or "inii iazytse" ("other languiges"). The chronicler identifies the Merya, Chud', Ves', and Muroma as autochthonous inhabitants of the northeastern areas that became part of the Rus' state between the 9th to 11th centuries CE^9,10^. Historians and archaeologists lack clarity on the extent of community and the origins of Slavic settlers on the East European Plain during the CE, as well as the initial area from which their expansion started. For a long time, the Slavic groups listed by the author of the *Primary Chronicle* were considered consolidated ethnic communities possessing their own distinct identity and cultural traditions, an interpretation corroborated by archaeological materials. Currently, historians debate whether these communities should be classified as polities (political and/or geographically distant communities or units), and the extent of their similarity remains a disputed issue^3,4,11^. The question of whether the Slavic communities in Rus'—specifically, two principal power centers of the northern region (Il'men and Dvina) and the southern Middle Dnieper region—shared a common origin and the degree of their interrelation remains an unresolved issue.

A key aspect in the formation of the Rus' state is the East Slavs' interaction with non-Slavic populations across the East European Plain. The circumstances regarding the migration of the Slavs to the North and Northeast, where they encountered the Fenno-Ugrian (or, more broadly defined, Uralic) ethnic groups, are currently poorly understood. To the west, Rus’ bordered Baltic tribes. In the southern regions, Slavic groups interacted with nomadic Turkic groups. However, the nature of these contacts—especially with Baltic and Finno-Ugric peoples—is poorly documented in written sources and likely varied considerably by region^12–17^.

The medieval Beloozero (Lake Beloye) groups, comprising communities from the Beloozero region and adjacent areas of contemporary Vologda and Arkhangelsk, are particularly significant for studying the interactions between the Slavs and the Fenno-Ugrian populations. These territories, located on the periphery of Rus', remained beyond the East Slavs' sphere of expansion until at least the 10th century CE. The presence of a Fenno-Ugrian population in these areas is supported by historical records, which can be validated by archaeological findings and the examination of substrate toponyms^14^. The lack of written sources makes it impossible to establish the size of the Fenno-Ugrian and Slavic populations or the precise character of their interactions. Recent archaeological data has uncovered the establishment of a new settlement network along the portages in the Beloozero region on the way to the Northern Dvina river system during the 11th–12th centuries CE. However, vast areas of the northern regions of Rus’ exhibit no evidence of permanent habitation until the 14th century CE. There is a divergence of opinion regarding the Slavic medieval colonization of these territories. Some scholars interpret it as a migration into sparsely populated areas, while others argue that Slavic served as a *lingua franca* for existing populations^14^ and has resulted in a linguistic shift within the indigenous Fenno-Ugrian groups, which has been absorbed into the Rus’ political structure, with no direct migration of Slavic communities to the northern regions. It is expected that archaeogenetics will provide independent information regarding the nature of ethnic transformations in northern areas.

A further crucial aspect of our research is the differences between urban and rural inhabitants in the context of the formation of the Rus' polity. The uniqueness of Rus' towns is frequently attributed not only to their socio-economic characteristics and institutional roles but also to the different origins of their inhabitants. The first Rus' towns are supposed to exhibit diverse ethnic composition^18–21^. Archaeological finds and written sources provide evidence of Scandinavian presence in many parts of Rus' in the 10th–11th centuries CE, particularly in early urban centers^18–20^. Therefore, genetic traces of this presence could be potentially detected.

The potential for conducting genetic research on Rus' is largely determined by the available source material. This includes three key factors: the paleoanthropological remains from excavated burial sites, their distinct archaeological and anthropological characteristics, and the historical trajectory of their excavation and study. Burial under mounds (barrows) was a widespread practice in Rus'. The remains, whether cremated (from the late 10th century) or inhumated (from circa 1000 CE), were placed in a small earthen mound or a pit beneath it. Also, a flat cremation ritual was practiced and occasionally performed in the same cemetery as barrow cremations. Other categories of sites containing medieval skeletal remains include those of 12th and 13th centuries CE urban cemeteries and mass graves related with the destruction of Rus' towns and murders of their residents by the Mongols in 1237–1238. Overall, the tradition of cremation among the East Slavs continued until the late tenth century, posing a major challenge to obtaining DNA samples from the pre-11th century era.

Large-scale excavations of medieval Rus' burials, primarily barrows, began in the mid-19th century CE. Anthropological materials obtained during the excavations of the 19th and first third of the 20th centuries are limited and often lack sufficient documentation or have not been preserved in museum collections. The human remains from the Rus', collected through excavations conducted in the late 20th and early 21st centuries, have greater significance owing to detailed documentation of burial contexts and a systematical approach to the collecting of anthropological material. They include samples from several significant historical centers of Rus’, including Novgorod (Yuriev Monastery), Old Ryazan, and Yaroslavl. The numerous, richly furnished burials in the Beloozero region (Northern Rus') are of particular interest due to their distinct cultural profiles, which reflect an interaction of Slavic and Fenno-Ugrian traditions. Genetic data from these sites are crucial, as the genetic profiles of the early medieval inhabitants of Northern Rus' region remained uncharacterized.

In the following sections, we present a comprehensive description of the anthropological material used in our whole-genome study, contextualized within both historical and anthropological frameworks. The material is organized into three sections. The first section (S1.1) focuses on samples from the Northern Rus' region. The second section (S1.2) addresses anthropological material from a broader area of Rus, predominantly derived from Slavic-related burials. The third section (S1.3) is dedicated to samples from territory neighboring Rus’, characterized by distinct cultural attributions. A comprehensive list of the samples employed in this study can be found in Supplementary Table 1.

### S1.1. Archaeological sites of the Northern Rus' region

These archaeological sites on the settlement of the northern periphery of medieval Rus' in the 11th–13th centuries CE are rural cemeteries that combine Slavic and Finnish cultural elements. They include two vast regions, Beloozero and Poonezhye, and several burials located there (Fig. 1).

#### S1.1.1. Burial sites of Beloozero region (Vologda)

The medieval Beloozero (Lake Beloye) region community includes inhabitants of areas near Lake Beloye and Lake Kubenskoye within the present-day Vologda region (Fig. 1).

##### S1.1.1.1. The Nefedyevo burial site

**Geographic Information:** The Nefedyevo I burial ground is located in the Kirillovsky district of Vologda region, on the left bank of the Porozovitsa (Itkla) river, 200 m east of the village of Nefedyevo, within the historical Beloozero region. It lies near the Volok Slavensky, which connected the Sheksna river system with the Sukhona and Northern Dvina rivers.

**Excavation History:** The site has suffered partial damage from quarrying activities. The burial ground was nearly completely excavated by the Onega-Sukhona expedition of the Institute of Archaeology of the Russian Academy of Sciences, led by N.A. Makarov, between 1983 and 1989. The northern section was designated as Nefedyevo IA and the southern section as Nefedyevo IB, with each section having separate burial numbers.

**Summary of sampled materials:** Over an area of 2400 square metres, 113 inhumation burials were discovered and excavated, with a balanced ratio of male, female, and child burials. The cultural context of the burial ground is a compound of Slavic and Finnic markers, with distinct regional "Beloozero" characteristics in burial rites, women’s costumes, and ceramics. The site dates from the early 11th to the early 13th centuries CE based on numerous finds of jewellery, coins and other grave goods^14^ (Supplementary Fig. 1).

**Burials of Nefedyevo IA**

**DB17. Burial 14**

***Grave type****:* This burial was found among a cluster of scattered bones in the arable layer at a depth of 0.25 m. The cluster contained the remains of three individuals: a woman (DB17), a man (DB18), and a child aged 4-6 years.

***Dating:*** 11th-12th centuries CE.

***Skeletal information****:* The skeleton of the woman (DB17) was oriented with her head to the east.

***Grave goods****:* 3 pendants (zoomorphic, bell-shaped, triangular), a bracelet, 5 temple rings, a button, a glass bead, a knife, and fragments of a round vessel.

***Genetic subcluster***: Rus_North_1.

**DB18. Burial 17**

***Grave type:*** The burial was found in a rectangular pit measuring 2.2 by 0.6 m. deep by about 20 cm.

***Dating:*** The first half to the middle of the 12th century CE.

***Skeletal information****:* The skeleton was placed on its back position with its head to the east with a deviation to the north. The bones of the arms were stretched along the body, the bones of the hands were on the pelvic bones. The skeleton belonged to a man aged 20-25 years.

***Grave goods****:* A bone cylinder with a round hole was found near the pelvic bones, and a knife and axe were placed on the left side, 2 stones, another bone cylinder, a silver cross, 3 glass beads and belt buckles were found near the feet.

***Genetic subcluster:*** Rus_North_1.

**DB1. Burial 19**

***Grave type****:*This burial was found among an aggregation of scattered or poorly preserved bones without anatomical order in the arable layer at a depth of 0.25 m, with the remains lying without anatomical order.

***Dating:*** The middle to the late 12th century CE.

***Skeletal information****:* The burial contained the remains of two individuals: an adult (DB1) and an infant.

***Grave goods****:* Grave goods included a bracelet, two noise-making pendants, a belt ring, metal and glass beads, a knife, and fragments of a vessel.

***Genetic subcluster***: Rus_North_1.

**DB19. Burial 20**

***Grave type****:* This burial was found in a rectangular coffin (0.37 m wide) at a depth of 0.45 m.

***Dating:*** 12th century CE.

***Skeletal information****:* The skeleton was lying on its back in an extended position with its head to the east. The skeleton belonged to a child aged 7-11 years.

***Grave goods****:* 4 temple rings (near the skull), a torc, 3 pendants (2 crosses, 1 bell-shaped), glass beads, a knife (in the abdomen area), 2 jingle bells, 2 noise-making pendants, a crutch, and a fragment of a bracelet.

***Genetic subcluster****:* Rus_North_1.

**Burials of Nefedyevo IB**

**DB20. Burial 4**

***Grave type****:* This burial was found in a rectangular grave pit (2.55x0.6 m, 0.16 m deep), with the remains of a wooden rectangular coffin (2.12x0.55 m).

***Dating:*** The second quarter to mid 12th century CE.

***Skeletal information****:* The poorly preserved skeleton was lying on its back in an extended position with its head to the east, the right hand placed on the pelvis. The skeleton belonged to a woman aged 25-35 years.

***Grave goods****:* A set of noise-making jewellery, an iron knife, a bracelet, a ring, a spindle whorl (in the pelvis), a comb (at the feet), a jingle bell, and a moulded vessel (in the southwest corner of the pit).

***Genetic subcluster****:* Rus_North_1.

**DB2. Burial 20**

***Grave type****:* This burial was found in a grave pit (2.5x0.8-1.2 m, 0.5 m deep), with remains of a wooden coffin and pieces of birch bark lying on and possibly under the skeleton.

***Coin-based dating:*** The first half of the 11th century CE.

***Skeletal information****:* The skeleton was lying on its back in an extended position with its head to the east, arms spread, the right arm slightly bent, with the hand lying at the level of the abdomen, and the left hand lying at the hip. The skeleton belonged to a woman aged 40-50 years (Supplementary Fig. 2 top right and left).

***Grave goods****:* 4 temple rings, 2 bronze and 1 iron torcs, beads (silver, stone, glass), 3 pendant coins (Samanids, Mansur b. Nuh, Bukhara, 960; Abbasids, al-Muktafi, 903/904; Marwanids, Mumahid al-Dawla, 1004/1005), horseshoe-shaped fibula, 7 jingle bells, 5 bracelets, 6 rings, a slate spinner, a coil of wire, a comb in a case, 2 iron knives, 2 moulded vessels, a dog skeleton (by the legs) (Supplementary Fig. 2 below).

***Additional information:*** Burial 20 is distinguished by the richness of the grave goods and the complexity of the burial rite, and dates from the first half of the 11th century. The burial was in a paired grave on top of a hill in the central part of the site, where no other burials were subsequently performed. Its strategic positioning within the burial ground's central area suggests a potential association with the settlement's founders. The analysis of data on strontium isotopic composition has revealed that the DB20 individual, presumed to be a "first-generation" migrant, resettled in Nefedyevo I during his adulthood^22,23^.

***Genetic subcluster:*** -

**DB21. Burial 24**

***Grave type****:* This burial was found in an oval grave pit (1.4x0.5 m) at a depth of 0.4-0.5 m.

***Dating:*** The first half to mid of the 12th century CE.

***Skeletal information****:* The skeleton, damaged by ploughing, was lying in an extended position on its back with its head to the east, left arm bent. The skeleton belonged to a child aged 4-5 years.

***Grave goods****:* 4 fang pendants (behind the skull, on the chest, and by the legs), a temporal ring, a fragment of a bronze torc, 2 crosses, beads, 2 pendants (a horse, a duck), a bone duck-shaped pendant, 2 rings, spirals, a comb (at the pelvis), a ring-shaped fibula (at the pelvis on the right), a slate spinner, 2 astragalus pendants, an iron knife, a moulded vessel (by the legs).

***Genetic subcluster****:* Rus_North_1.

**DB22. Burial 25**

***Grave type****:* This burial was found at a depth of 0.5-0.6 m from the modern surface at the contact of the arable layer and dark humus layer filling a continental hollow. The contour of the grave pit was not traced.

***Dating:*** The first half to mid of the 12th century CE.

***Skeletal information****:* The skeleton was lying in an elongated position with its head to the east. The skeleton belonged to a man aged 45-50 years.

***Grave goods****:* No grave goods were found.

***Genetic subcluster****:* Rus_North_1.

**DB23. Burial 29**

***Grave type****:* This burial ground was found within the boundaries of an oval-shaped spot (at least 2 m long) at a depth of 0.3-0.35 m, together with woody decay from a coffin (about 1.4 m long).

***Dating:*** The last third of the 11th to the turn of the 12th centuries CE.

***Skeletal information****:* The skeleton was lying in an elongated position with its head to the north-east. The poorly preserved skeletal remains belonged to a man aged 20-35 years.

***Grave goods****:* 2 nails, a ring, a knife (along the right leg), an iron object, an iron axe (near the leg), a buckle, moulded and round vessels (by the legs).

***Genetic subcluster****:* Rus_North_1.

**DB3. Burial 41**

***Grave type****:* This burial was found at a depth of 0.20-0.25 m from the modern surface in a dark humusized cultural layer, the contour of the grave pit was not traced. Wood decay from the coffin and a layer of birch bark under it were revealed.

***Dating:*** The second to third quarter of the 11th century CE.

***Skeletal information****:* The poorly preserved skeleton was lying on its back in an extended position with its head facing east, arms spread, the left arm slightly bent. The skeleton belonged to a woman aged 45-55 years.

***Grave goods****:* 7 temple rings, 2 iron torcs, 1 bronze torc, glass beads, silver pendant, 2 pendant coins (Samanids, Nasr bin Ahmed, Samarkand, 921/922; Abbasids, al-Mahdi, Madinat as-Salam, 775/776), a horseshoe-shaped fibula (on the chest), a comb in a case (on the abdomen), 2 triangular pendants, 13 jingle bells, an iron knife with a bone handle in a scabbard (by the pelvis), an iron needle, 4 bracelets, 3 rings, 2 moulded vessels (near the left hand) (Supplementary Fig. 3).

***Additional information:***

Burial 41 as distinguished by its rich array of grave goods, positioned centrally within the burial ground. This burial was identified as belonging to the earliest period, dating to the second to third quarter of the 11th century CE.

***Genetic subcluster***: Rus_North_1.

**DB24. Burial 43**

***Grave type****:* This burial was found in an oval grave pit (2.4x1.4 m, 0.2 m deep) at a depth of 0.3-0.35 m. The contour of the pit is indistinct, and the remains of a wooden coffin of rectangular shape (1.7x0.66 m) were revealed.

***Dating:*** The late 11th to first quarter of the 12th centuries CE.

***Skeletal information****:* The skeleton was lying on its back with its head to the northeast, the bone remains disturbed. The skeleton belonged to a man aged 25-35 years.

***Grave goods****:* A temporal ring, a bronze bead, a ferrule, a horseshoe-shaped fibula (on the chest), a round-shaped fire striker, a comb, a bone ornament, an iron knife (under the pelvis on the left), an iron arrowhead (at the knee on the left), a moulded vessel (by the legs).

***Genetic subcluster****:* Rus_North_1.

**DB25. Burial 45**

***Grave type****:* This burial was found in an oval grave pit (2.3x0.75 m) at a depth of 0.4-0.45 m. Woody decay from the coffin was revealed at the bottom of the pit.

***Dating:*** 12th century CE.

***Skeletal information****:* The poorly preserved skeleton was lying on its back in an extended position with its head to the northeast. The skeleton belonged to a man aged 50-55 years.

***Grave goods****:* An iron arrowhead (near the left elbow), an iron knife with a wooden handle (near the right hand), an iron axe (by the legs), a ring, a round-shaped fire striker with flint, a comb, a vessel (by the legs), dog bones (in the southwestern part of the pit).

***Genetic subcluster****:* Rus_North_1.

**DB4. Burial 46**

***Grave type****:* This burial was found in an oval grave pit (2.25x0.85 m) at a depth of 0.4-0.45 m. The 6 iron nails from the coffin were revealed.

***Dating:*** 12th to the first half of the 13th centuries CE.

***Skeletal information****:* The skeleton was lying on its back in an extended position with its head to the northeast. The skeleton belonged to a woman aged 50-60 years.

***Grave goods****:* 3 temple rings, a torc, glass beads, 4 pendants (one icon, 2 zoomorphic, 1 bell-shaped), an iron knife with a wooden handle (near the pelvis on the left), a bronze bead, a spiral bead, a moulded vessel (by the legs).

***Genetic subcluster:*** -

**DB5. Burial 47**

***Grave type****:* This burial was found in a rectangular pit with rounded corners (2.65x0.75 m) at a depth of 0.35 m. The bottom of the pit was covered with wood decay from the coffin.

***Dating:*** The turn of the 11th to 12th - the first quarter of the 12th centuries CE.

***Skeletal information****:* The skeleton was lying on its back in an extended position with its head to the northeast. The skeleton belonged to a man over 55 years old.

***Grave goods****:* An iron knife (at the pelvis on the right), a glass bead, an iron axe (at the knee on the left), an iron arrowhead, an iron fire striker, a vessel (by the legs).

***Genetic subcluster****:* Rus_North_1.

**DB6. Burial 48**

***Grave type****:* This burial was found at a depth of 0.35 m at the contact of tilth-topsoil and the mainland. Woody decay from the coffin was revealed.

***Dating:*** The turn of the 11th to 12th-the first quarter of the 12th centuries CE.

***Skeletal information****:* The poorly preserved skeleton was lying on its back in an extended position with its head to the northeast. The skeleton belonged to a woman aged 40-50 years.

***Grave goods****:* A ring, 2 temple rings, a torc, 5 pendants (4 with the image of St. George), 5 zoomorphic pendants, glass beads, a wick pipe, an iron knife (at the pelvis on the left), 7 spiral beads, a bronze bead, 2 bell-shaped pendants, a moulded vessel (by the legs), animal bones.

***Genetic subcluster****:* Rus_North_1.

**DB26. Burial 53**

***Grave type****:* Found in a rectangular grave pit with rounded corners (2.4x0.8 m) at a depth of 0.3-0.5 m. The fill contained wood decay and 3 iron nails from the coffin (2.25x0.4 m).

***Dating:*** The third quarter of the 12th century CE.

***Skeletal information****:* The skeleton was lying on its back in an extended position with its head to the east. The arms were bent and the bones of the feet were disturbed. The skeleton belonged to a woman aged 20-30 years.

***Grave goods****:* 18 temple rings, glass beads, 2 crosses (on the chest and by the feet), 2 rings, an iron knife (at the pelvis on the left), a horseshoe-shaped fibula (by the feet), a slate spinner, a conical pendant, a moulded vessel (by the feet).

***Genetic subcluster****:* Rus_Middle.

**DB27. Burial 54**

***Grave type****:* Found in a wooden coffin at a depth of 0.35-0.45 m. Woody decay and 7 iron nails from the coffin were revealed.

***Dating:*** 12th century CE.

***Skeletal information****:* The skeleton was lying on its back in an extended position with its head to the northeast. The gender of the skeleton has not been determined, the age is 45 years old.

***Additional information:*** The genetic gender is male.

***Grave goods****:* A cross worn next to the skin, glass beads, knife (by the hip), round vessel (by the feet).

***Genetic subcluster****:* Rus_North_1.

**DB28. Burial 55**

***Grave type****:* Found at a depth of 0.45-0.55 m. Woody decay, 6 nails and a coffin crutch were traced.

***Dating:*** 12th century CE.

***Skeletal information****:* The skeleton was lying on its back in an extended position with its head to the northeast, hands resting on the pelvic bones. The skeleton belonged to a man aged 45-55 years.

***Grave goods****:* A cross worn next to the skin, glass beads, a dart tip (by the abdomen), a knife (between the thigh bones), a comb (by the feet).

***Genetic subcluster****:* Rus_North_1.

**DB29. Burial 57**

***Grave type****:* Found at a depth of 0.55-0.8 m, the outline of the grave pit could not be traced. Woody decay, an iron crutch and 3 coffin nails were found.

***Dating:*** The second half of the 12th to the beginning of the 13th centuries CE.

***Skeletal information****:* The poorly preserved skeleton was lying on its back in an extended position with its head to the northeast. The right hand was resting on the pelvis and the legs were slightly tucked. The skeleton belonged to a woman over 60 years old.

***Grave goods****:* Glass beads, an iron knife (by the knee on the right), a silver ring, a round vessel (by the feet).

***Genetic subcluster****:* Rus_North_1.

**DB30. Burial 58**

***Grave type****:* Found at a depth of 0.7-0.9 m in a grave pit with indistinct boundaries (width 1.3 m). Woody decay and 2 coffin nails were revealed.

***Dating:*** The first half of the 12th century CE.

***Skeletal information****:* The skeleton was lying on its back in an extended position with its head to the northeast. The skeleton belonged to a woman aged 50-55 years.

***Grave goods****:* 3 temple rings, a bronze torc, glass beads, a duck-shaped pendant (by the pelvis on the left), a jingle bell, an iron knife (by the hip on the left), a ring, a moulded vessel (by the feet).

***Genetic subcluster****:* Rus_North_1.

**DB31. Burial 61**

***Grave type****:* Found at a depth of 0.5-0.6 m in a quadrangular grave pit with rounded corners (2.3x1.0 m).

***Dating:*** The second half of the 11th to the first quarter of the 12th centuries CE.

***Skeletal information****:* The skeleton was lying on its back in an extended position with its head to the southeast, arms slightly bent at the elbows. The skeleton belonged to a man aged 50-55 years.

***Grave goods****:* An axe with an iron wedge (by the right knee), an arrowhead, a knife (by the left hip)

***Genetic subcluster****:* Rus_North_1.

**DB32. Burial 70**

***Grave type****:* This burial was found at a depth of 0.9-1.0 m in a grave pit (1.85x0.6 m).

***Dating:*** 12th century CE.

***Skeletal information****:* The skeleton was lying on its back in an extended position with its head to the east. The skeleton belonged to a man aged 40-50 years.

***Grave goods****:* An axe (by the shoulder), an iron knife (under the pelvis).

***Genetic subcluster****:* Rus_North_1.

**DB33. Burial 73**

***Grave type****:* This burial was found at a depth of 0.45-0.55 m in a rectangular grave pit with rounded corners (2.55x0.85 m). Woody decay and 3 nails from a board coffin (2.3x0.45 m) were revealed.

***Dating:*** The second half of the 12th-the beginning of the 13th centuries CE.

***Skeletal information****:* The poorly preserved skeleton was lying on its back in an extended position with its head to the east, arms bent. The skeleton belonged to a man aged 30-35 years.

***Grave goods****:* A ring, an axe (at the right knee), a knife (between the knees), an arrowhead (by the legs).

***Genetic subcluster****:* Rus_North_1.

**DB34. Burial 75**

***Grave type****:* This burial was found at a depth of 0.3-0.4 m in a rectangular grave pit with rounded corners (2.5x0.8 m). Woody decay and 5 nails from the coffin were revealed.

***Dating:*** The middle to the second half of the 12th century CE.

***Skeletal information****:* The skeleton was lying on its back in an extended position with its head to the east, arms slightly bent. The skeleton belonged to a man aged 40-60 years (Supplementary Fig. 4).

***Grave goods****:* A dagger, an arrowhead, an iron rod, 4 buttons, a ring, a comb (at the left hip), an iron knife, an awl, a needle, 2 fire strikers with flint, an axe (by the legs). The brush, the handle of the brush with a bronze pommel

***Genetic subcluster****:* Rus_North_1.

***Additional information:*** This is a rare burial in Nefedyevo with specialised military weapons, continuing the earlier tradition of the burial rite of the military group of Nikolskoe III.

**DB35. Burial 77**

***Grave type****:* This burial was found at a depth of 0.3-0.5 m in a quadrangular grave pit with rounded corners (2.2x0.75 m). Woody decay from the coffin was traced.

***Dating:*** The second half of the 12th - the beginning of the 13th centuries CE.

***Skeletal information****:* The skeleton was lying on its back in an extended position with its head to the southeast. The skeleton belonged to a man aged 18-20 years.

***Grave goods****:* A knife, an awl, a bronze bracket, animal bones, a round vessel (by the feet).

***Genetic subcluster****:* Rus_North_1.

**DB36. Burial 78**

***Grave type****:* This burial was found at a depth of 0.3-0.4 m in a rectangular grave pit with rounded corners (2.6x0.9 m). Woody decay and 4 nails from the coffin were recorded.

***Dating:*** The second half of the 12th - the beginning of the 13th centuries CE.

***Skeletal information****:* The skeleton was lying on its back in an extended position with its head to the east. The skeleton belonged to a man aged 40-45 years.

***Grave goods****:* An axe (at the right knee), an iron knife (by the legs), bird and fish bones, a round vessel (by the legs).

***Genetic subcluster****:* Rus_North_1.

##### S1.1.1.2. The Minino burial site

**Geographic Information:** The burial ground known as Minino II is situated in the Vologda district of the Vologda region, on Lake Kubenskoye, near the mouth of the Dmitrievka river, approximately 0.5 km north of the village of Minino, within the historical Syamskaya Volost’.

**Excavation History:** Excavations conducted by the Onega-Sukhon expedition of the Institute of Archaeology of the Russian Academy of Sciences, under the direction of N.A. Makarov, and the Scientific Production Centre "Antiquities of the North", led by A.V. Suvorov, between 1997 and 2004, covered an area of over 880 square metres.

**Summary of sampled materials:** This site represents a subterranean burial ground, with no visible surface markers for the burial locations. These excavations revealed 82 mediaeval burials: 17 cremations and 65 inhumations, some of which were well-preserved while others had been partially disturbed (Supplementary Fig. 5). The cultural context of the burial ground reflects a blend of Slavic and Finnic influences, with notable regional "Beloozero" characteristics evident in burial practices, women's attire, and ceramics. The dating of the burial ground extends from the second half of the 10th to the early 13th centuries CE. This timeline is corroborated by numerous artefacts and jewellery finds, as well as radiocarbon dates obtained from coal samples from the cremation^7,24^.

**DB38. Burial 1**

***Grave type****:* This burial was found at a depth of 0.2-0.3 m in a rectangular grave pit with rounded corners (2.26x0.78 m).

***Dating:*** The first half of the 12th century CE.

***Skeletal information****:* The skeleton was lying on its back in an extended position with its head to the east. The poorly preserved skeleton belonged to a woman over 50 years old.

***Grave goods****:* 2 temple rings, glass and stone beads, a ring-bead, 3 pendants (1 round, 2 duck-shaped), a bronze buckle, a ring, a round vessel (by the legs) with fragments of bones of a teenager and a ram.

***Genetic subcluster****:* Rus_North_1.

**DB39. Burial 3**

***Grave type****:* This burial was found at a depth of 0.21-0.32 m in a rectangular grave pit with rounded corners (2.4x1.0 m). Traces of woody decay from the coffin were revealed.

***Dating:*** The turn of the 11th-12th centuries CE - the beginning of the 12th century CE.

***Skeletal information****:* The poorly preserved skeleton was lying on its back in an extended position with its head to the east. The skeleton belonged to a woman aged 25-35 years.

***Grave goods****:* 2 rings, 12 temple rings, a billonite torc, glass beads, cowrie pendants, a bronze bracelet, a bronze chain with pendants (1 duck, 2 horses), bronze spiral beads, a bone comb, an iron knife, and a moulded vessel (placed at the feet).

***Genetic subcluster****:* Rus_North_1.

**DB40. Burial 4**

***Grave type****:* This burial was found at a depth of 0.6-0.62 m in a rectangular grave pit with rounded corners (2.05x0.68 m).

***Dating:*** The last quarter of the 11th century CE.

***Skeletal information****:* The skeleton was lying on its back in an extended position with its head to the east. The skeleton belonged to a woman aged 35-45 years.

***Grave goods****:* 2 temple rings, 8 spiral beads, a ring, glass and gold-glass beads, 2 coin pendants (Henry III, 1039-1056, type 1046-1056; Conrad II, 1024-1039, or Henry III), 55 tin-lead plaques (on the hands), an iron ring (in the belt area) with cords with beads and bronze rings, a duck pendant, an iron knife in a leather sheath, a bone comb in a case (under the left leg), and a fragment of a moulded vessel (at the feet).

***Genetic subcluster****:* Rus_North_1.

**DB37. Burial 8**

***Grave type****:* This burial was found in an oval grave pit (2.25x0.67 m) at a depth of 0.55-0.6 m.

***Dating:*** 12th-beginning of the 13th centuries CE.

***Skeletal information****:* The skeleton was lying on its back in an extended position with its head to the northwest, with arms bent. The skeleton likely belonged to a young man aged 16-18 years.

***Grave goods****:* A flat stone, 2 nagels, a flint plate, and polecat bones.

***Genetic subcluster****:* Rus_SWest.

***Additional information:*** As indicated by the archaeological data, this is one of the latest burials, topographically somewhat isolated, which is quite consistent with the location of the burials of recent migrants, unrelated to the rest of the group of individuals from this cemetery.

**DB9. Burial 9**

***Grave type****:* This burial was found in an oval grave pit (2.26x0.77 m) at a depth of 0.48 m. Woody decay from the coffin was revealed.

***Dating:*** The second half of the 11th century CE.

***Skeletal information****:* The skeleton was lying on its back in an extended position, slightly turned on its side, with the head to the southeast. The skeleton belonged to a man aged 35-45 years (Supplementary Fig. 6 left).

***Grave goods****:* 2 spiral beads, an iron axe with a wedge (at the right hand), 2 rings, a leather belt with a bronze buckle, plates and a ring, 2 iron buckles, an iron ring, and pike bones (Supplementary Fig.6 right).

***Genetic subcluster****:* -

**DB10. Burial 15**

***Grave type****:* This burial was found in a sub-rectangular grave pit (1.83x0.7 m) at a depth of 0.58 m.

***Dating:*** The second half of the 10th-turn of the 10th-11th centuries CE.

***Skeletal information****:* The skeleton was lying on its back in an extended position with its head to the east. The skeleton belonged to a man aged 45-55 years (Supplementary Fig. 7 left).

***Grave goods****:* A belt with a buckle and 2 rings, leather bag, 2 fire striker flints, a fire stone, a fire striker, a bronze wick tube, an iron knife (Supplementary Fig. 7 right).

***Genetic subcluster****:* Rus_North_1.

**DB41. Burial 17**

***Grave type****:* This burial was found in a sub-rectangular grave pit (2.51x1.1 m) at a depth of 0.2 m. Evidence of wood decay from the coffin was noted.

***Dating:*** The second quarter of the 11th century CE.

***Skeletal information****:* The skeleton was lying on its back in an extended position with its head to the east. The skeleton belonged to a woman over 50 years old (Supplementary Fig. 8 top left and right).

***Grave goods****:* Tin-lead wreath, 5 temple rings, a bronze torc, glass beads, 2 coin pendants (Bavaria, Regensburg, Otto, 976-982; Swabia, Strasbourg, Henry II, 1002-1024, or Werner I, 1001-1028), a bronze horseshoe-shaped fibula (on the chest), 2 bracelets, a bone comb in a case (near the arm), an iron knife (under the right arm), a ring, a lunula, bones of a bird, a fish, large and small animals, 2 moulded vessels (at the feet) (Supplementary Fig. 8 below).

***Genetic subcluster****:* Rus_North_1.

**DB42. Burial 19**

***Grave type****:* This burial was found in a rectangular grave pit (2.32x0.87 m) at a depth of 0.54-0.59 m. Evidence of decay from the coffin was noted.

***Dating:*** The third quarter of the 11th century CE.

***Skeletal information****:* The skeleton was lying on its back in an extended position with its head to the east, with arms slightly spread apart. The skeleton belonged to a woman aged 40-49 years (Supplementary Fig. 9 left).

***Grave goods****:* 3 temple rings, glass beads, a silver ring, a bone comb in a case, an iron knife (near the right hand), a moulded vessel (near the arm), fish bones (at the legs) (Supplementary Fig. 9 right).

***Genetic subcluster****:* -

**DB11. Burial 26**

***Grave type****:* This burial was found in a grave pit (2.26x1.0 m) at a depth of 0.57-0.59 m. Evidence of decay from the coffin was noted.

***Dating:*** The middle of the 12th-the early 13th centuries CE.

***Skeletal information****:* The skeleton was lying on its back in an extended position with its head to the east. The skeleton belonged to a woman aged 25-35 years.

***Grave goods****:* 10 temple rings, 3 rings, a bronze pendant, an iron knife (under the ribs), animal bones (at the legs).

***Genetic subcluster****:* Rus_North_1.

**DB43. Burial 29**

***Grave type****:* This burial was found in a rectangular grave pit (2.09x0.7 m) at a depth of 0.51-0.57 m.

***Dating:*** The second half of the 12th-the early 13th centuries CE.

***Skeletal information****:* The skeleton was lying on its back in an extended position with its head to the east. The skeleton belonged to a child aged about 7 years old.

***Grave goods****:* 2 temple rings, glass beads, 3 lunula pendants, an iron nail, 2 iron needles, an iron knife (near the left hand).

***Genetic subcluster****:* Rus_North_1.

**DB44. Burial 30**

***Grave type****:* This burial was found in a grave pit within a spot of houmous sandy clay (0.49x0.13-0.24 m).

***Dating:*** 11th century CE.

***Skeletal information****:* The skeleton belonged to an infant under six months old, with the bones not in anatomical order.

***Grave goods****:* No grave goods were found.

***Genetic subcluster****:* Rus_Middle.

**DB45. Burial 31**

***Grave type****:* This burial was found in a burial pit within a spot of houmous sandy clay (0.49x0.13-0.24 m).

***Dating:*** 11th century CE.

***Skeletal information****:* The well-preserved skeleton was lying on its back with its head to the northeast, the left arm bent at the elbow and placed on the stomach. The skeleton belonged to an infant under six months.

***Grave goods****:* No grave goods were found.

***Genetic subcluster****:* -

**DB46. Burial 37**

***Grave type****:* This burial was found in a rectangular grave pit with rounded corners (3.14x0.82-1.02 m) at a depth of 0.66 m.

***Dating:*** 11th-12th centuries CE.

***Skeletal information****:* The skeleton was disturbed, lying with its head to the west. The skeleton belonged to a woman over 50 years old.

***Grave goods****:* A round-shaped fire striker, a bronze wick pipe, a bronze binding, a wheel-made vessel.

***Genetic subcluster****:* Rus_North_1.

**DB47. Burial 38**

***Grave type****:* This burial was found in a quadrangular grave pit (2.74x0.7-0.84 m). Evidence of wood decay from the coffin was noted.

***Dating:*** The middle of the 12th-the early 13th centuries CE.

***Skeletal information****:* The skeleton was lying in an extended position on its back with its head to the east. The skeleton belonged to a man over 50 years old (Supplementary Fig. 10 left).

***Grave goods****:* An axe (on top of the left hand), an iron knife (under the left hand), a round vessel (at the feet), a double-sided comb (at the feet) (Supplementary Fig. 10 right).

***Genetic subcluster****:* Rus_North_1.

**DB48. Burial 41**

***Grave type****:* This burial was found in a sub-rectangular grave pit (0.96x0.48 m) at a depth of 0.58 m, containing the skeletal remains of four people - one adult and three children.

***Dating:*** The second half of the 12th - the early 13th centuries CE.

***Skeletal information****:* The skeleton (DB48) was completely preserved, though the bones were not in anatomical order, with the skull in the eastern part of the pit. This skeleton belongs to a child 2-3 years old.

***Grave goods****:* No grave goods were found.

***Genetic subcluster****:* Rus_North_1.

**DB49. Burial 46**

***Grave type****:* This burial was found in a rectangular burial pit (2.29x0.8 m) at a depth of 0.57m.

***Dating:*** The middle of the 12th - the early 13th centuries CE.

***Skeletal information****:* The bone remains were not in anatomical order. The skeleton was lying with its head to the east in an extended position. The skeleton belonged to a teenager aged 15-18 years.

***Grave goods****:* An iron knife, a ring.

***Genetic subcluster****:* Rus_North_1.

**DB50. Burial 47**

***Grave type****:* This burial was found in an oval grave pit (1.59x0.75-0.8 m).

***Dating:*** The second half of the 12th-the early 13th centuries CE.

***Skeletal information****:* The well-preserved skeleton was lying with its head to the east, arms extended along the body, legs strongly bent. The skeleton belonged to a child of 4-5 years.

***Grave goods****:* A lunula pendant, a glass bead, a fragment of the fire striker (near the skull).

***Genetic subcluster****:* Rus_North_1.

**DB12. Burial 48**

***Grave type****:* This burial was found in a rectangular grave pit (2.4x0.9-0.95 m) at a depth of 0.86 m in a double burial containing the bone remains of a woman (DB12) and a man (not included in this study). Evidence of wood decay from the coffin was noted.

***Dating:*** The middle of the 12th-the early 13th centuries CE.

***Skeletal information****:* The skeleton belonged to a woman aged 45-55 years (Supplementary Fig. 11 left and central).

***Grave goods****:* 12 temple rings, glass beads, an icon pendant, a cockerel pendant, an iron knife (under the right hand), a ring, and a double-sided comb (Supplementary Fig. 11 right).

***Genetic subcluster****:* -

**DB51. Burial 54**

***Grave type****:* This burial was found in an oval grave pit (the size of the uncovered part of the pit was 1.0x1.02 m) at a depth of 0.55 m.

***Dating:*** 12th-the early 13th centuries CE.

***Skeletal information****:* A fragmented skull and a fragment of a tubular bone were revealed, with the anatomical order of the bones disrupted. The sex and age of the skeleton cannot be determined.

***Grave goods****:* A cross-shaped pendant.

***Genetic subcluster****:* Rus_North_1.

**DB13. Burial 62**

***Grave type****:* This burial was found in an oval grave pit (3.32x1.22-1.4 m), which contained the skeletal remains of three people—a woman, a child, and the man (DB13).

***Dating:*** The middle of the 12th-the early 13th centuries CE.

***Skeletal information****:* The skeletal remains were not in anatomical order. The skeleton belonged to a man aged 40-49 years.

***Grave goods****:* Glass beads, a lunula, a ring, an iron plate, a spearhead, a round vessel.

***Genetic subcluster****:* Rus_North_1.

**DB53. Burial 68**

***Grave type****:* This burial was found in an oval grave pit (1.91x0.95-0.99 m) at a depth of 0.5m.

***Dating:*** 11th century CE.

***Skeletal information****:* The well-preserved skeleton was lying on its back in an extended position with its head to the northwest. The skeleton belonged to a child of 9-10 years.

***Grave goods****:* Fish bones (along the right arm), a moulded vessel (on the right by the knee).

***Genetic subcluster****:* Rus_North_1.

**DB54. Burial 75**

***Grave type****:* This burial was found in a rectangular grave pit with rounded corners (2.7x1.0 m) at a depth of 0.6 m.

***Dating:*** The second half of the 12th - the early 13th centuries CE.

***Skeletal information****:* The skeleton was lying on its back in an extended position with its head to the northwest. The bone remains belonged to a man aged 40-49 years (Supplementary Fig. 12).

***Grave goods****:* No grave goods were found.

***Additional information:*** As indicated by the archaeological data, this is one of the latest burials, topographically somewhat isolated, which is quite consistent with the location of the burials of recent migrants, unrelated to the rest of the group of individuals from this cemetery.

***Genetic subcluster****:* Rus_Core.

##### S1.1.1.3. The Shuygino burial site

**Geographic Information:** The Shuygino burial ground is located in the Kirillovsky district of the Vologda region, on the right bank of the Itkla (Porozovitsa) river, on the western outskirts of the village of Shuygino, within the historic Beloozero region, near the Volok Slavensky, which connected the Sheksna river system with the Sukhona and Northern Dvina rivers. The site has suffered partial damage from quarrying activities.

**Excavation History:** The burial ground was completely excavated by the Onega-Sukhona expedition of the Institute of Archaeology of the Russian Academy of Sciences, led by N.A. Makarov, in 1985.

**Summary of sampled materials:** Over an area of 400 square metres, 27 inhumation burials were discovered and examined, about half of which were children's graves. The overall cultural appearance of the site is ancient Rus’, though individual elements of the burial rites are associated with the Finnic tradition, including the eastern orientation of the early group of burials. Based on the finds of jewellery and household items, the burial ground is dated to the last third of the 12th to the early 13th centuries CE.

**DB71. Burial 3**

***Grave type****:* This burial was found in a grave pit (2.05x0.75 m) at a depth of 0.2 m. Evidence of wood decay from the coffin was noted.

***Dating:*** The last third of the 12th-the early 13th centuries CE.

***Skeletal information****:* The skeleton was lying on its back in an extended position with its head to the west. The arms were bent, with one positioned on the chest and the other on the stomach. The skeleton belonged to a man aged 45-55 years.

***Grave goods****:* An iron knife situated near the pelvis on the right.

***Genetic subcluster****:* Rus_North_1.

**DB72. Burial 4**

***Grave type****:* This burial was found in a grave pit (1.45x0.75 m) at a depth of 0.15 m.

***Dating:*** The last third of the 12th-the beginning of the 13th centuries CE.

***Skeletal information****:* The skeleton was lying on its back with its head to the west. The skeleton belonged to a child approximately 5 years old±16 months.

***Grave goods****:* A lunula pendant located in the neck area and glass beads.

***Genetic subcluster****:* Rus_North_1.

**DB69. Burial 5**

***Grave type****:* Found at a depth of 0.25-0.3 m, this burial was damaged by plowing.

***Dating:*** The last third of the 12th-the early 13th centuries CE.

***Skeletal information****:* The skeleton was lying in an extended position with its head to the west. Age is undetermined due to the damage.

***Grave goods****:* No grave goods were found.

***Genetic subcluster****:* Rus_North_1.

**DB68. Burial 6**

***Grave type****:* This burial was found in a burial pit (2.35x0.65 m) at a depth of 0.35 m.

***Dating:*** The last third of the 12th-the early 13th centuries CE.

***Skeletal information****:* The skeleton was lying on its back in an extended position with its head to the west. The arms were bent with the hands raised to the shoulders. The skeleton belonged to a man aged 30-40 years.

***Grave goods****:* No grave goods were found.

***Genetic subcluster****:* Rus_North_1.

**DB70. Burial 8**

***Grave type****:* This burial was found in a grave pit (2.0x0.8 m) at a depth of 0.5 m.

***Dating:*** The last third of the 12th-the early 13th centuries CE.

***Skeletal information****:* The skeleton was lying on its back in an extended position with its head to the east. The arms were bent and placed on the stomach. The skeleton belonged to a woman aged over 50 years old.

***Grave goods****:* 4 temple rings and a slate spindle wheel.

***Genetic subcluster****:* Rus_North_1.

**DB73. Burial 12**

***Grave type****:* This is a cluster of scattered bones found in tilled soil and quarry ejecta.

***Dating:*** The last third of the 12th-the early 13th centuries CE.

***Skeletal information****:* The skeleton belonged to a woman aged 35-40 years.

***Grave goods****:* An iron knife.

***Genetic subcluster****:* Rus_North_1

**DB74. Burial 20**

***Grave type****:* Found at a depth of 0.25 m, the grave pit was not clearly traced, and the burial was heavily damaged by plowing.

***Dating:*** The last third of the 12th-the early 13th centuries CE.

***Skeletal information****:* The skeleton was lying on its back in an extended position with its head to the east. The legs were slightly bent, the arms were bent with the hands on the pelvis. The skeleton belonged to an adult woman.

***Grave goods****:* A ring, a knife near the left leg, and an iron ring.

***Genetic subcluster****:* Rus_North_1.

##### S1.1.1.4. The Nikolskoe burial site

**Geographic Information:** The Nikolskoe III burial site (Kema necropolis, burial barrows near the village of Boltinskaya) is located in the Vashkinsky district of the Vologda region, on the left bank of the Kema river, 5 km southwest of the village of Nikolskoe, within the Beloozero region. The site has suffered partial damage from quarrying activities.

**Excavation History:** Five burial barrows were excavated in 1927 by M.E. Arsakova. Between 1981 and 1985, the Onega-Sukhona expedition of the Institute of Archaeology of the Russian Academy of Sciences, under the direction of N.A. Makarov studied 37 burial barrows containing 50 burials, alongside 23 ground pit burials without barrows.

**Summary of sampled materials:** The burial ground comprised 43 burial barrows, with additional burials in grave pits unmarked on the surface. The Nikolskoe III burial ground is notable among other Beloozero necropolises for its unusual age and gender composition, featuring a predominance of male burials (51.4%). What is more, many male burials included weapons, such as a sword and 11 battle axes, and a significant number of Western European coins (82 pieces) were found in the graves, often serving as "obols of the dead" (in 26 burials). Judging by the burial rites and the assemblage of women's jewellery, the general cultural appearance of the burial ground is Rus’/East Slavic with the features of military elite necropolis This differentiates it from the culture of most other Beloozero burial grounds, which typically exhibit a blend of Slavic and Finnish cultural elements. The burial ground was in use from the second to the third quarter of the 11th century CE^24^ (Supplementary Fig. 13).

**DB77. Barrow 2. Burial 1**

***Grave type****:* This burial was discovered in the barrow (diameter 6.7 m, height 0.5 m). The burial was completely destroyed by excavation.

***Dating:*** The second-third quarter of the 11th century CE.

***Skeletal information****:* The skeleton belonged to a man aged 17-20 years.

***Grave goods****:* No grave goods were found.

***Genetic subcluster****:* -

**DB55. Barrow 8. Burial 2**

***Grave type****:* This burial was discovered in the barrow (diameter 6.6 m, height 0.6 m) in a grave pit (2.1x0.8 m, depth 0.8 m). Eight coffin nails were found.

***Coin-based dating:*** The late 1060s-1070s.

***Skeletal information****:* The skeleton was lying on its back in an extended position with its head to the west. The left arm was bent, with the hand on the pelvis. The skeleton belonged to a child aged 4-8 years.

***Grave goods****:* 7 temple rings, glass beads, a coin pendant (England, Knut, 1023-1029), 8 pendants (1 with a cross, a horse, 4 ducks, a spoon, a coin-shaped), a cross, a ring-shaped fibula (on the chest), a jingle bell, a sharp point, a bone cylinder, a bone spinning wheel, 16 pendants made of animal teeth, a claw pendant, 15 pendants made of astragalus, 17 pendants made of fruit and nut nucleuses, a cowrie shell, a wooden rod, an iron knife, a comb in a case, a conical pendant, 3 rings, a bracelet, a coin (Frisia, Egbert II, 1068-1090), half a coin (Ever, Duke Ordulf (Otto), 1059-1071 or Count Hermann, 1059-1086), a moulded vessel (at the feet on the left), a clay pysanka, and a glazed cup (at the feet on the right).

***Genetic subcluster****:* Rus_Core.

**DB56. Barrow 16. Burial 1**

***Grave type****:* This burial was found in the barrow (diameter 8 m, height 0.75 m). Ten coffin nails were found.

***Coin-based dating:*** The second-third quarter of the 11th century CE.

***Skeletal information****:* The skeleton was lying on its back in an extended position with its head to the west. The skeleton belonged to a woman aged 35-45 years.

***Grave goods****:* 4 temple rings, a coin (in the mouth) (Mainz, Archbishop Lupold, 1051-1059), a coin pendant (Regensburg, Duke Henry V, 1017-1026), beads (gold-glass, stone, bronze), a bronze ring, a comb, an iron knife (on the stomach), and a ring.

***Genetic subcluster****:* Rus_Core.

**DB57. Barrow 16. Burial 2**

***Grave type****:* This burial was found in a ground pit (2.6x1.0 m, depth 0.85 m) in the barrow (diameter 8 m, height 0.75 m). Nine coffin nails were found.

***Coin-based dating:*** The second-third quarter of the 11th century CE.

***Skeletal information****:* The skeleton was lying on its back in an extended position with its head to the west. The skeleton belonged to a man aged 25-35 years.

***Grave goods****:* 24 coins (15 denarii, minted in Eber (Duke Ordulf (Otto), 1059-1071 or Count Hermann, 1059-1086), the rest: Frisia, Dokkum, Count Egbert II, 1068-1090; Frisia, Count Bruno III, 1038-1057; Emden, Count Hermann von Calvelage, circa 1020-1051; Saxony, denarius of the 11th century CE; England, King Cnut, 1017-1023; 3 coins are unidentifiable), a fragment of a comb (on the stomach), an axe (at the right knee), and a round vessel (at the feet).

***Genetic subcluster****: -*

**DB58. Barrow 22. Burial 1**

***Grave type****:* This burial was found in a ground pit (2.6x0.9 m, depth 0.45 m) in the barrow (diameter 6 m, height 0.2 m).

***Coin-based dating:*** The second-third quarter of the 11th century CE.

***Skeletal information****:* The skeleton was lying on its back in an extended position with its head to the west. The arms were bent, with the hands on the pelvic bones. The skeleton belonged to a woman aged 20-30 years.

***Grave goods****:* A temple ring, a coin (in the mouth) (Ever, Duke Ordulf (Otto), 1059-1071 or Count Hermann, 1059-1086), glass beads, a coin pendant (Magdeburg, anonymous mint, first half of the 11th century CE), a ring, and a knife (near the pelvis on the right).

***Genetic subcluster****:* Rus_SWest.

**DB76. An extension to the Barrow 22. Burial 1**

***Grave type****:* It was found in a ground pit (1.85x0.65 m, depth 0.4 m). Only the skull remains of the skeleton.

***Dating:*** The second-third quarter of the 11th century CE.

***Skeletal information****:* The skeleton was lying with its head to the west. The skeleton belonged to a child aged 1-2 years.

***Grave goods****:* Glass beads, a coin (Germany, unknown mint), a zoomorphic pendant, an iron knife, and a moulded vessel.

**Genetic *sub*cluster**: Rus_Core.

**DB59. An extension to the Barrow 22. Burial 2**

***Grave type****:* It was found in a ground pit (1.6x0.8 m, depth 0.8 m). Eleven coffin nails were found.

***Dating:*** The second-third quarter of the 11th century CE.

***Skeletal information****:* Only the skull remains of the skeleton. The skeleton was lying with its head to the west. The skeleton belonged to a child aged 1 year±4 months.

***Grave goods****:* A glass bead, a coin pendant (undetermined), and a round vessel.

***Genetic subcluster****:* Rus_Core.

**DB7. An extension to the Barrow 22. Burial 6**

***Grave type****:* This burial was found in a grave pit (1.90x0.55 m, depth 0.45 m).

***Dating:*** The **s**econd-third quarter of the 11th century CE.

***Skeletal information****:* The skeleton was lying on its back in an extended position with its head to the west. The arms were bent, the right hand was on the stomach, and the left hand was on the chest, with the legs slightly bent at the knees. The skeleton belonged to an individual aged 20-30 years.

***Grave goods****:* No grave goods were found.

***Genetic subcluster****:* Rus_North_1

**DB8. Barrow 27. Burial 2**

***Grave type****:* This burial was found in a ground pit (3.2x1.2 m, depth 0.5 m) in the barrow (diameter 11.6 m, height 0.8 m). Twelve coffin nails were found. The pit is oriented west-east. The burial is destroyed, and the bones are mixed.

***Dating:*** The second-third quarter of the 11th century CE.

***Skeletal information****:* The skeleton belonged to a woman aged 20-25 years.

***Grave goods****:* Glass beads and a fragment of a moulded vessel.

***Genetic subcluster****:* Rus_SWest.

**DB60. Barrow 30. Burial 1**

***Grave type****:* This burial was found in a ground pit (depth 0.4 m) in the barrow (diameter 6 m, height 0.4 m).

***Dating:*** Th**e** second-third quarter of the 11th century CE.

***Skeletal information****:* The skeleton was lying on its back in an extended position with its head to the west. The skeleton belonged to a child aged 3-5 years.

***Grave goods****:* A bronze chain.

***Genetic subcluster****:* Rus_SWest.

**DB61. Barrow 36, Burial 1**

***Grave type****:* This burial was found in a ground pit (2.7x1.1 m, depth 0.7 m) in the barrow (diameter 6.5 m, height 0.4 m).

***Dating:*** The second-third quarter of the 11th century CE.

***Skeletal information****:* Only the skull remains of the skeleton. The skull belonged to a child aged 7-11 years.

***Grave goods****:* No grave goods were found.

***Genetic subcluster****:* Rus_Middle.

**DB62. Barrow 37. Burial 3**

***Grave type****:* This burial was found in a ground pit (2.0x0.9 m, depth 0.7 m) in the barrow (diameter 6.5 m, height 0.4 m). Twelve coffin nails were found.

***Dating:*** The second-third quarter of the 11th century CE.

***Skeletal information****:* The skeleton was lying with its head to the west, only the skull remains. The skeleton belonged to a child aged 3-5 years.

***Grave goods****:* A coin (in the abdominal area) (Frisia, Winsum, Count Egbert II, 1068-1090), knife (in the pelvic area), an axe-amulet (in the legs), a round vessel, and a glazed pysanka (in the legs)

**Genetic subcluster**: Rus_Core.

**DB63. An extension to the Barrow 37. Burial 4**

***Grave type****:* The burial was found in a grave pit (2.5x1.05 m, depth 0.6 m). Four coffin nails were found.

***Dating:*** The second-third quarter of the 11th century CE.

***Skeletal information****:* The poorly preserved skeleton was lying on its back in an extended position with its head to the west. The right arm was extended. The skeleton belonged to a man aged 40-50 years.

***Grave goods****:* An iron buckle and a knife in a leather sheath (in the pelvic area).

***Genetic subcluster***: Rus_Core.

#### S1.1.2. Burial grounds of the Poonezhye region (Arkhangelsk)

##### S1.1.2.1. The Voezero burial site

**Geographic Information:** The Voezero burial ground is situated in the Nyandomsky district of the Arkhangelsk region, on the western shore of Lake Spasskoye, near the southern edge of the village of Gavrilovskaya, within the historic Poonezhye region (related to Onega river). The site has been disturbed by vegetable storehouse pits.

**Excavation History:** The burial ground was investigated by the Onega-Sukhona expedition of the Institute of Archaeology of the Russian Academy of Sciences conducted in 1980-1983, during which an area of 150 square metres was excavated.

**Summary of sampled materials:** The graves are predominantly shallow, with many covered by stone coverings laid over wooden frames. The excavation revealed 34 inhumation burials, both intact and partially disturbed, with nearly half being children's burials. The burial practices and artefacts suggest an ancient Rus’ cultural context, with elements of the burial rite, such as stone coverings, and decorations indicative of Baltic-Finnic traditions. The site is dated to the second half of the 13th to the first half of the 14th centuries CE, as determined by both the artefacts and radiocarbon dates from the bone samples obtained from three burials^25^.

**DR12. Burial 8**

***Grave type****:* This burial was found on the boundary between tilth-top soil and the underlying substratum.

***Dating:*** The second half of the 13th-the first half of the 14th centuries CE.

***Skeletal information****:* The anatomical order of the bones is disrupted, with numerous bones missing. The crushed skull, oriented south, belonged to a man aged 45-55 years (Supplementary Fig. 14 left).

***Grave goods****:* No grave goods were found.

***Genetic subcluster****:* Rus_North_2.

**DR13. Burial 9**

***Grave type***: This burial was discovered in an oval grave pit (1.04x0.3 m) at a depth of 0.5 m. Traces of wood decay from the coffin were observed.

***Dating:*** The second half of the 13th-the first half of the 14th centuries CE.

***Skeletal information****:* The skeleton was lying on its back in an extended position with the head to the west. The left arm was bent with the hand resting on the pelvis. The skeleton belonged to a child aged 1-2 years (Supplementary Fig. 14 right).

***Grave goods****:* No grave goods were found

***Genetic subcluster****:* Rus_North_2.

**DR11. Burial 15**

***Grave type****:* This burial was found in an oval grave pit (1.25x0.3 m) at a depth of 0.2-0.3 m.

***Dating:*** The **s**econd half of the 13th-the first half of the 14th centuries CE.

***Skeletal information****:* The skeleton was lying on its back with the head to the west. The upper part of the skeleton was missing, with the broken skull located in the abdominal area. This burial is that of a child approximately 10 years old.

***Grave goods****:* No grave goods were found.

***Genetic subcluster****:* Rus_North_2.

**DR14. Burial 16**

***Grave type****:* This burial was found in an oval grave pit (1.1x0.3 m) at a depth of 0.1-0.25 m.

***Dating:*** The second half of the 13th-the first half of the 14th centuries CE.

***Skeletal information****:* The skeleton was lying on its back with the head oriented to the west and the arms slightly bent. This burial belongs to a child aged 4-5 years.

***Grave goods****:* No grave goods were found.

***Genetic subcluster****:* Rus_North_2.

**DR15. Burial 17**

***Grave type****:* This burial was found in an oval burial pit (0.98x0.3 m) at a depth of 0.1-0.15m. Traces of wood decay from the coffin were noted.

***Dating:*** The second half of the 13th-the first half of the 14th centuries CE.

***Skeletal information****:* The skeleton was lying on its back with the head to the west, although some bones were disordered. This burial is that of a child aged 6-12 months.

***Grave goods****:* No grave goods were found.

***Genetic subcluster****:* -

**DR16. Burial 19**

***Grave type****:* This burial was found on the boundary between tilth-top soil and the underlying substratum at a depth of 0.35-0.5 m. Traces of wood decay from the coffin were revealed.

***Dating:*** The second half of the 13th-the first half of the 14th centuries CE.

***Skeletal information****:* The skeleton was lying on its back with the head oriented to the west and the left arm bent. This burial belongs to a child aged 1-2 years.

***Grave goods****:* No grave goods were found.

***Genetic subcluster****:* -

**DR17. Burial 22**

***Grave type****:* This burial was found in an oval grave pit (0.68x0.22 m) at a depth of 0.55 m. Wood decay from the coffin was traced.

***Dating:*** The second half of the 13th-the first half of the 14th centuries CE.

***Skeletal information****:* The skeleton was lying on its back with the head oriented to the west. The arms were slightly bent, with the hands in the pelvic area. This burial is that of a newborn.

***Grave goods****:* No grave goods were found.

***Genetic subcluster****:* -

**DR18. Burial 23**

***Grave type****:* This burial was discovered in a grave pit (0.65x0.2 m) at a depth of 0.7 m.

***Dating:*** The second half of the 13th-the first half of the 14th centuries CE.

***Skeletal information****:* The skeleton was positioned on its back with a slight tilt to the right side and the head to the west. The legs were slightly bent, the right arm slightly bent, and the left arm extended. This burial belongs to a newborn.

***Grave goods****:* No grave goods were found.

***Genetic subcluster****:* Rus_North_2.

**DR19. Burial 27**

***Grave type****:* This burial was found in a grave pit of indeterminate shape at a depth of 0.6-0.65 m within a wooden block (0.77x0.17 m).

***Dating:*** The second half of the 13th-the first half of the 14th centuries CE.

***Skeletal information****:* The skeleton was lying on its back with the head oriented to the west. The anatomical arrangement of many bones was disturbed. This burial is that of a newborn.

***Grave goods****:* No grave goods were found.

***Genetic subcluster****:* Rus_North_2.

**DR20. Burial 28**

***Grave type****:* This burial was found in a grave pit of indeterminate shape at a depth of 0.6-0.65m.

***Dating:*** The **s**econd half of the 13th-the first half of the 14th centuries CE.

***Skeletal information****:* The skeleton was lying supine with the legs slightly bent, the head oriented to the west, the right arm strongly bent and resting on the chest, and the left arm extended. This burial belongs to a newborn.

***Grave goods****:* No grave goods were found.

***Genetic subcluster****:* Rus_North_2.

**DR21. Burial 29**

***Grave type****:* This burial was discovered in a grave pit of indeterminate shape at a depth of 0.6-0.65 m.

***Dating:*** The **s**econd half of the 13th-the first half of the 14th centuries CE.

***Skeletal information****:* The skeleton was lying on its back with the head oriented to the west, and the left arm bent and placed on the stomach. This burial is that of a child aged approximately 2 years old.

***Grave goods****:* No grave goods were found.

***Genetic subcluster****:* -

**DR22. Burial 31**

***Grave type****:* This burial was found at a depth of 0.5-0.7 m from the present surface.

***Dating:*** The **s**econd half of the 13th-the first half of the 14th centuries CE.

***Skeletal information****:* The skeleton was lying on its back with the head oriented to the west. This burial belongs to a child.

***Grave goods****:* No grave goods were found.

***Genetic subcluster****:* Rus_North_2.

#### S1.1.3. The Novinki burial site (Vologda)

**Geographic Information:** The burial ground is located northeast of the village of Novinka-Lebedevo, in a forested area along the road leading to the villages of Yartsevo and Timoshkino in the Babayevsky district of the Vologda region, situated on the gentle slope of the left bank of the Kolp river. The barrows were positioned on both sides of the road.

**Excavation History:** Excavations were carried out by the Vologda expedition of the Institute of Archaeology of the USSR Academy of Sciences under the supervision of A.V. Nikitin between 1964 and 1969.

**Summary of sampled materials**: All the barrows, constructed from homogeneous yellow sand, have a hemispherical shape, with diameters ranging from 6 to 9 m and heights of 0.7 to 1 m. Ashy patches were recorded at the level of the ancient ground surface. The burials were single interments, discovered in rectangular grave pits aligned along a west-east axis at a depth of 0.6–0.9 m. The deceased were found in an extended supine position, with varying hand placements. Some burials were located within wooden log structures. The grave goods included temporal rings with attached beads, rhombic shield-shaped rings, twisted bracelets, and iron knives.The burial practices and artefacts suggest ancient Rus’ cultural context. Dating: 11th-12th centuries CE^26^.

Genetic analysis was carried out on two skulls that are kept in the Museum of Anthropology, Lomonosov Moscow State University.

**AB35. Barrow 8 or 18**

***Grave type****:* The sample was from one of the two barrows, in the rectangular grave pit.

***Skeletal information****:* The skeleton was oriented westwards and belonged to a man.

***Grave goods****:* No grave goods were found.

***Museum ID:*** КО 273-9.

***Genetic subcluster****:* Rus_Baltic.

**AB36. Barrow 38**

***Grave type****:* The burial was located in a grave pit (2x0.8m and 0.3 m deep, within a barrow 5 m in diameter and 1 m in height.

***Skeletal information****:* The skeleton was in an extended supine position, with the head-oriented westwards. The right hand was placed on the chest, and the left arm was stretched along the body. The skeletal remains belonged to a man.

***Grave goods****:* Plaques, 2 wire temporal rings, an iron knife with traces of a wooden handle, a bracelet made of 3 twisted wires, a loop, and a ring with a round inset made of paste.

***Museum ID:*** КО 273-10.

***Genetic subcluster****: -*

#### S1.1.4. The Ploskoe burial site (Vologda)

**Geographic Information:** The burial ground is situated on the western side of the village of Ploskoe in the Babayevsky district of the Vologda region, on an elevated area with a height of 4-6 m.

**Excavation History:** The burial ground was investigated by the Vologda expedition of the Institute of Archaeology of the USSR Academy of Sciences under the leadership of A.V. Nikitin in 1965.

**Summary of sampled materials**: The burial ground originally contained 41 barrows, most of which were encircled by stones. The barrows have hemispherical or elongated shapes, with diameters ranging from 4 to 7 m and heights up to 1 m. The barrows predominantly contained individual burials, but some collective burials were also recorded. All burials were placed in wooden coffins within small pits on the mainland soil, at the base of the barrow. Above the burials, approximately midway through the height of the barrow, traces of "fireplaces" (ash patches) were recorded. The remains were laid out in an extended supine position, with their hands crossed on their chest. The grave goods included three-bead temporal rings, round billon pendants, barrel-shaped glass beads, bronze open-ended plate bracelets, iron axes, and fragments of pottery vessels. The burial practices and artefacts suggest ancient Rus’ cultural context.Dating: 12th-13th centuries CE^27^.

Genetic analysis was carried on a bone sample obtained from a single skull that is kept in the Museum of Anthropology, Lomonosov Moscow State University.

**AB42**

***Skeletal information****:* The burial belonged to a woman.

***Museum ID:*** КО 273-39.

***Genetic subcluster:*** -

### S1.2 Archaeological sites of other Rus’ regions

These burials include cemeteries from four geographic regions of ancient Rus’: Central Rus’ Volga-Oka region, North-Western Rus’ Novgorod, Western Rus’ Dnieper-Dvina region and Southern Rus' (Fig. 1).

#### S1.2.1. Archaeological sites of the Central Rus' Volga-Oka region

##### S1.2.1.1. The Gnezdilovo burial site (Suzdal)

**Geographic Information:** The Gnezdilovo 12 burial ground is situated in the Suzdal district of the Vladimir region, approximately 6 km southwest of Suzdal and 1.5 km northeast of the village of Gnezdilovo.

**Excavation History:** In 1851, A.S. Uvarov conducted excavations here, uncovering 28 burial barrows with both cremation and inhumation burials. Presently, the site is a flat field where geophysical exploration has identified around 80 rounded anomalies, likely corresponding to the bases of levelled burial barrows. From 2020 to 2023, the Suzdal expedition of the Institute of Archaeology of the Russian Academy of Sciences and the State Historical Museum, led by N.A. Makarov and A.M. Krasnikova, excavated an area of approximately 1200 square metres.

**Summary of sampled materials**: They examined 50 burials performed according to the inhumation rite and remnants of destroyed cremation burials. Some inhumation burials were originally beneath burial barrows, while others were barrowless. The cultural characteristics of the burial ground are distinctly Rus’/East Slavic with the features of military elite necropolis. Among the women's jewellery, items of Volga-Finnic types and some Scandinavian pieces were also found. A notable feature of the burial ground is the presence of weapons (nine battle axes) and trade inventory (scales, weights) in the men's burials. The presence of burials accompanied by equestrian equipment and weapons lends the Gnezdilovо burial ground the features of a special druzhina’s necropolis. The burial grounds of Gnezdilovo 12 offer insights into the culture of the elite and the distinctive features of the social structure of the North-East of Rus’ at the time when information about Suzdal begins to appear in chronicles. The functioning period of the cemetery spans from the 10th to the early 12th centuries CE, as determined by numerous finds of coins, jewellery, household items, and weapons. Radiocarbon dating of bone tissue samples from four burials confirms that all inhumation burials date from the turn of the 10th-11th centuries to the early 12th century CE^28,29^ (Supplementary Fig. 15).

**DB15. Burial 3**

***Grave type****:* The grave pit (2.8x1.2 m, depth 0.5 m) was located on a site surrounded by a ring-shaped groove with a diameter of 3.9 m. Three iron nails from a coffin were found in the lower part of the fill.

***Dating:*** The second half of the 11th-the first half of the 12th centuries CE.

***Skeletal information****:* The skeleton was lying on its back in an extended position with its head to the west, slightly deviated to the north. The skeleton belonged to a man aged 35-49 years. The skull was turned to the north towards the left shoulder, the right arm was extended along the body, and the left arm was bent at a right angle, lying on the chest. The skull showed traces of lifetime injuries, possibly from a chopping weapon.

***Grave goods****:* No grave goods were found.

***Genetic subcluster****:* Rus_Baltic.

**DB16. Burial 5**

***Grave type****:* This burial was found in a grave pit (2.4x1.0 m, depth 0.3 m), traces of a rectangular wooden structure were revealed in the fill of the pit.

***Dating:*** 11th century CE.

***Skeletal information****:* A poorly preserved skeleton was lying on its back in an extended position with its head to the west. The left arm was bent at the elbow, with the hand located in the pelvic area, and the legs were extended. The skeleton belonged to an adult aged 30-39 years (Supplementary Fig. 16 top left and right).

***Grave goods****:* In the fill of the pit at the bottom, a denarius pendant (Regensburg 1002-1024) and 2 blue ring-shaped beads were found, under the skull a ring-shaped temple ring, and in the neck area 18 glass beads (Supplementary Fig. 16 below).

***Genetic subcluster****:* Rus_SWest.

**DB64. Burial 16**

***Grave type****:* This burial was found in a sub-rectangular grave pit (2.5x0.9 m, depth 0.43 m), traces of a decayed coffin were found in the filling of the pit and two iron nails were revealed.

***Dating:*** 11th century CE.

***Skeletal information****:* A poorly preserved skeleton was lying on its back in an extended position, with its head to the west and arms slightly bent at the elbows. The skeleton belonged to a woman over 40 years old.

***Grave goods****:* An iron knife was found in the waist area on the right, and beads were found in the neck area: 15 glass and 1 amber (Supplementary Fig. 17 top).

***Genetic subcluster***: Rus_Baltic.

**DB65. Burial 17**

***Grave type****:* This burial was found in a sub-rectangular burial pit (2.9x1.2 m, depth 0.34 m), traces of a wooden structure (coffin) were found in the fill in the bottom part.

***Dating:*** The first half of the 11th century CE.

***Skeletal information****:* The poorly preserved skeleton was lying on its back in an extended position, the right arm extended along the body. The skeleton belonged to a woman aged 25-35 years.

***Grave goods****:* An iron knife was found by the left femur, in the area of the skull there were 6 glass beads, under the skull there were 2 temple three-bead rings, and at the feet there was a molded pot with handles. Another 8 glass beads were found in the filling of the pit Supplementary Fig. 17 below).

***Genetic subcluster****:* Rus_SWest.

**DB66. Burial 19**

***Grave type****:* This burial was discovered on the site of levelled barrow 1 with a diameter of about 7 m. It was found in a sub-rectangular grave pit (3.3x1.6 m, 0.24 m deep), oriented northwest-southeast. In the pit, traces of a sub-rectangular wooden structure made of oak and ash with cuts at the corners measuring 2.7x1.3 m were found.

***Dating:*** The turn of the 10th to 11th-the first quarter of the 11th centuries CE.

***Skeletal information****:* The skeleton was lying on its back in an extended position, with widely spread arms and the skull turned to the left. The skeleton belonged to a man aged 25-30 years (Supplementary Fig. 18 top and below right).

***Grave goods****:* A non-ferrous metal buckle with an iron prong was found under the skull on the right, a fragment of a dirham (Mansur b. Nuh or Nuh b. Mansur, 360s-370s AH (970-990)) at the shoulder on the right, an iron knife at the hip on the right, a fire striker with flint on the left, and a moulded vessel at the feet on the right. A folding scale and two weights folded in a leather bag were found near the pelvis on the right. Stirrups, a girth buckle, a bit, and a battle hatchet lay in the southwestern part of the pit (Supplementary Fig. 18 below left).

***Genetic subcluster****:* Rus_SWest.

**DB67. Burial 22/23**

***Grave type****:* The fill of the pit. The pit has been tentatively interpreted as a ditch of a levelled burial barrow.

***Dating:*** 11th-the first half of the 12th centuries CE.

***Skeletal information****:* Mixed, fragmented bone remains of two individuals—an adult, presumably a man, and a child—were found in the fill of pit 5 in excavation 2 and named burials 22 and 23. The child was taken for this study.

***Grave goods****:* Along with the bones, an iron knife, an iron nail, and a bronze plate were found in the fill of the pit.

***Genetic subcluster****:* Rus_SWest.

**DB75. Burial 24**

***Grave type****:* This burial was found on the site of a levelled barrow 3, surrounded by a ring ditch. It was found in a sub-rectangular grave pit (3.7x1.4 m, depth 0.48 m), traces of a wooden structure with cuts at the corners were found in the filling of the pit. In the lower part of the pit filling and at its bottom, nails from the coffin were found, 12 in the skull area and 9 in the leg area.

***Dating:*** The first half of the 11th century CE.

***Skeletal information****:* The skeleton was lying on the back in an extended position with its head to the west, arms extended along the body. The skeleton belonged to a man aged 35-40 years (Supplementary Fig. 19 top and below left).

***Grave goods****:* 2 iron weights were found near the right humerus, a battle axe and the remains of a bone item near the right femur, an iron knife, two bronze belt rings, and a lyre-shaped buckle near the left femur, and a silver Western European coin near the lower jaw (Supplementary Fig. 19 below right).

***Genetic subcluster****:* Rus_Core

##### S1.2.1.2. The Luzhki burial site (Moscow)

**Geographic Information:** The burial ground is located on the edge of the first above-floodplain terrace of the left bank of the Oka river, near the town of Serpukhov in the Moscow region.

**Excavation History:** The burial ground is poorly studied. Most of the collection from excavations carried out in the late 1980s-1990s has been lost. Such a composition of finds complicates the cultural interpretation of the site.

**Summary of sampled materials:** The burial site contains ground cremations from the 11th century CE and a single inhumation burial. Grave goods associated with cremation burials include monochrome ribbed cylindrical glass beads, which have analogies at Knyazhya gora, Birka, Staraya Ladoga, and the Saltovo burial ground (9th-10th centuries CE), as well as longitudinally striped triple beads (10th-11th centuries CE), buttons, and belt garniture elements in the Saltovo style, along with sułgamas and bells. The only known inhumation burial, which is older than the cremations, illustrates the cultural transformations occurring in the microregion at the beginning of the 2nd millennium CE. Dating: the first half of the 11th century CE. The radiocarbon date obtained from the collagen of the sole inhumation skeleton is 1060±20 years BP, calibrated to 981-1014 AD (IGAN-AMS 7606)^30^.

**DR35. Burial in pit 12**

***Grave type****:* The inhumation burial was located in a rectangular ground pit, traced under the cremation layer.

***Dating:*** The first half of the 11th century CE.

***Skeletal information****:* The skeleton was lying extended on its back, oriented with the head to the northwest. The burial belonged to a man aged 35-45 years.

***Grave goods****:* No grave goods were found. However, ceramic fragments with chamotte inclusions and a distinct vessel rim profile were recorded in the fill of the pit.

***Additional information:*** The nature of the rim profile suggests dating to the turn of the 1st-the early 2nd millennium CE.

***Genetic subcluster****:* Rus_Core.

##### S1.2.1.3. The Akatovo burial site (Moscow)

**Geographic Information:** The burial ground is located near the village of Akatovo, 2-3 km from the Saltykovskaya station in the Moscow region, on the high right bank of the Pehorka river.

**Excavation History:** The site was excavated by N.G. Nedoshvina in 1966-1967.

**Summary of sampled materials:** The barrow group consisted of 71 barrows, with 31 of them excavated. All barrows were of a regular hemispherical shape, most of them surrounded by ditches. The height of the barrows averaged 0.5-1 m, with a diameter of 5-6 m. Almost all burials in the burial ground were made in pits covered by barrows, and less frequently in the subsoil layer. The depth of the grave pits varied from 0.3 to 1.4 m. Most of the Akatovo barrows contained individual burials. The orientation of the burials was westward. The posture of the buried was extended supine, with bent arms and hands resting on the chest or abdomen. Most of the burials did not contain traces of wooden structures or coffins. No grave goods were found. The burial practices and artefacts suggest ancient Rus’ cultural context. Dating: the late 11th-12th centuries CE^31^.

Genetic analysis was carried out on two skulls that are kept in the Museum of Anthropology, Lomonosov Moscow State University.

**AB1. Barrow 48**

***Grave type****:* The burial was found in a pit (3.1x1.4 m, depth 1.4 m) covered by the barrow (diameter 5.7 m, height 1 m).

***Dating:*** The late 11th-12th centuries CE.

***Skeletal information****:* The skeleton was lying in an extended supine position, with its head to the west. The right hand was placed on the pelvic bones. The remains belonged to a man.

***Grave goods****:* No grave goods were found.

***Museum ID:*** КО 99-5.

***Genetic subcluster:*** -

**AB2. Barrow 34**

***Grave type****:* The burial was found in a pit (2.25x1.3 m, depth 0.3 m) covered by the barrow (diameter 5.5 m, height 1 m).

***Dating:*** The late 11th-12th centuries CE.

***Skeletal information****:* The skeleton was lying in an extended supine position, with the head oriented to the west. The right hand was placed on the pelvic bones.The remains belonged to a woman.

***Grave goods:*** Pottery fragments from two vessels were found in the barrow.

***Museum ID:*** КО 99-4.

***Genetic subcluster:*** -

##### S1.2.1.4. The Velikoe burial site (Vladimir)

**Geographic Information:** The burial ground is located in the Kasimov Сounty of the Ryazan Governorate (currently part of the Vladimir region), 2 km from the village of Parakhino, on a site bounded by the Gus river to the east, the Dukhovitsa river to the north, and the Dandur to the south.

**Excavation History:** The site was surveyed by N.F. Nefedov in 1877.

**Summary of sampled materials:** At the time of discovery, only 10 barrows were recorded. The burial practices and artefacts suggest ancient Rus’ cultural context. Dating: 11th-12th centuries CE^32^.

Genetic analysis was carried out on two skulls that are kept in the Museum of Anthropology, Lomonosov Moscow State University.

**AB4. Barrow 3. Skull 6**

***Grave type:*** The burial was found at the level of the ancient ground surface, within the barrow(diameter 8.5 m, height 1.8 m).

***Dating:*** 11th-12th centuries CE.

***Skeletal information:*** The skeleton was lying prone, in an extended position, with the head oriented northwestward. The skull was turned to the left. Charcoal and pottery fragments were found at the feet. The remains belonged to a man.

***Museum ID:*** 1208.

***Genetic subcluster:*** -

**AB5. Barrow 9. Skull 3**

***Grave type:*** The burial was found at the level of the ancient ground surface, with hands folded on the abdomen.

***Dating:*** 11th-12th centuries CE.

***Skeletal information:*** The skeleton was lying in an extended supine position, with the head oriented southwestward.

***Grave goods:*** Glass beads, temple rings, rings on the fingers, and pottery fragments.

***Museum ID:*** 1215.

***Genetic subcluster:*** -

##### S1.2.1.5. The Vorobyovo burial site (Tver)

**Geographic Information:** The burial ground is located near the village of Vorobyovo in the Kimrsky district of the Tver region, on the right bank of the Medveditsa river.

**Excavation History:** A total of 11 barrows have been excavated.

**Summary of sampled materials:** The burial ground comprises two groups of barrows, with one group containing 29 barrows and the other 49 ones. The barrows are lined with stones around their perimeter, with some stones also found within the barrows themselves. Certain barrows are surrounded by ditches up to 1.4 m wide and 0.7 m deep. The skeletal remains are in very poor condition, and the exact placement of the burials within the barrows is unclear. Several burials were interred in wooden structures. In one barrow (№4), a double burial was identified. The grave goods include bracelets, rings, amber and blue glass beads, temple rings, an iron knife with traces of a wooden handle, and ceramic vessels. The burial practices and artefacts suggest ancient Rus’ cultural context. Dating: 11th-12th centuries CE ^33^.

Genetic analysis was carried out on two skulls that are kept in the Museum of Anthropology, Lomonosov Moscow State University.

**AB7**

***Skeletal information****:* The skeletal remains belonged to a man.

***Museum ID:*** 984.

***Genetic subcluster****:* Rus_Middle.

**AB8**

***Skeletal information****:* The skeletal remains belonged to a woman.

***Museum ID:*** 981.

***Genetic subcluster****: -*

##### S1.2.1.6. The Voronovo burial site (Yaroslavl)

**Geographic Information:** The burial ground is located on the outskirts of the village of Voronovo in the Uglichsky district of the Yaroslavl region, on the bank of the Uleyma river. **Excavation History:** A total of the 21 barrows were excavated by A.I. Kelsiev in 1878. By the time of excavation, the burial barrows had been partially damaged due to construction activities.

**Summary of sampled materials:** The diameters of the barrows generally did not exceed 6 m, with the exception of one the barrow (№ 9), which measured over 8.5 m (4 sazhen) and contained two skeletons. The heights of the barrows varied between 0.35 and 2.1 m, and the burial depth of the skeletons ranged from 0.35 to 1.4 m. The burials were located in soil pits, the boundaries of which were poorly defined. The skeletons were positioned extended on their backs, with the head oriented to the east or southeast. The burial practices and artefacts suggest ancient Rus’ cultural context. Dating: 11th-12th centuries CE^34^. Genetic analysis was carried out on two skulls that are kept in the Museum of Anthropology, Lomonosov Moscow State University.

**AB9. Barrow 20**

***Grave type****:* The burial was found in a soil pit (depth 1.4 m) within the barrow (diameter 7 m, height 0.9 m).

***Dating:*** 11th-12th centuries CE.

***Skeletal information****:* The skeleton was lying extended on its back, head to the east. Based on the set of ornaments, the burial is presumed to have belonged to a woman. No anthropological identification of the skeletal remains has been conducted.

***Grave goods****:* Wire rings, green and black beads, seven-lobed temple rings, a twisted bracelet made of thick wire (on the left arm), a thick bracelet with small rings at the ends (on the right arm), a ring (on the right hand), and a triangular pendant (near the left side).

***Museum ID:*** 1076.

***Genetic subcluster****: -*

**AB10. Barrow 19**

***Grave type****:* The burial was located in a grave pit (depth 0.7 m) within the barrow (diameter and height unknown).

***Dating:*** 11th-12th centuries CE.

***Skeletal information****:* The skeleton was lying extended on its back, head to the east-southeast. The skull was turned to the left. The burial is identified as belonging to a man.

***Grave goods****:* A single ring.

***Museum ID:*** 1080.

***Genetic subcluster****:* Outliers.

##### S1.2.1.7. The Zhukovo burial site (Yaroslavl)

**Geographic Information:** The burial ground is located near the village of Zhukovo in the Uglichsky district of the Yaroslavl region.

**Excavation History:** The site was excavated by Y.A. Ushakov in 1877.

**Summary of sampled materials:** Burials were discovered in the subsoil layer within the barrow measuring 21.3 m in diameter and 0.7 m in height. The skeletons were found in an extended supine position, oriented with their heads to the northwest, and arms extended along the body. The grave goods included bronze temporal rings, an iron knife, bronze bracelets, bronze rings, iron axes, ceramic vessels, gilded glass beads, and necklaces made of glass beads.The burial practices and artefacts suggest ancient Rus’ cultural context. Dating: 11th-12th centuries CE^35^.

Genetic analysis was carried out on one skull that is kept in the Museum of Anthropology, Lomonosov Moscow State University.

**AB15**

***Skeletal information:*** The skeletal remains belonged to a woman.

***Museum ID:*** 1033.

***Genetic subcluster: -***

##### S1.2.1.8. The Ziminki burial site (Murom)

**Geographic Information:** The barrow is located in the Muromsky district of the Vladimir region, on the left bank of the Ilevna river.

**Excavation History:** The site was excavated by N.G. Kertselly in 1879 and N.F. Nefedov in 1885.

**Summary of sampled materials:** All the barrows had a hemispherical shape. The skeletal remains were found in an extended supine position, oriented with the head to the west, in rectangular-shaped ground pits. There is no information available regarding the presence or composition of grave goods. Dating: 11th-12th centuries CE^36^.

Genetic analysis was carried out on one skull that is kept in the Museum of Anthropology, Lomonosov Moscow State University.

**AB16**

***Skeletal information****:* The burial belonged to a man.

***Museum ID:*** 1194.

***Genetic subcluster***: Rus_Core.

##### S1.2.1.9. The Kiryanovo burial site (Yaroslavl)

**Geographic Information:** The burial site is situated near the village of Stromyni on the left bank of the Volga river, in the Uglich district of the Yaroslavl region.

**Excavation History:** The site was excavated by A.I. Kelsiev in 1878. A total of 39 barrows (№ 18-55) were excavated.

**Summary of sampled materials:** The burials were contained within the barrow up to 5.3 m in diameter. In some cases, the presence of stone settings around the perimeters of the barrow was recorded. The skeletons were found in an extended supine position, with heads oriented to the east. The skulls were either turned to the left or right. In the female burials, the arms were extended along the body, while in the male burials, one arm was extended, and the other was resting on the chest. The grave goods in the female burials included bronze wire temporal rings (positioned on either side of the skull), gilded glass beads (near the neck), carnelian and mosaic beads, iron knives (near the pelvic bones), iron sickles (by the left leg), bronze bracelets, both flat and triangular in cross-section (on the arms), a "bronze twisted wire hoop" (possibly a neck ring), bronze buckles (on the abdomen), bronze pendant bells attached to chains (around the neck), and ceramic vessels (at the feet). The male burials contained iron knives (between the elbow and ribs), iron sickles (by the right thigh), rings (on the right hand), and ceramic vessels (at the feet). The burial practices and artefacts suggest ancient Rus’ cultural context. Dating: 11th-12th centuries CE.

Genetic analysis was carried out on one skull that is kept in the Museum of Anthropology, Lomonosov Moscow State University.

**AB17**

***Skeletal information:*** The skeletal remains belonged to a man.

***Museum ID:*** 1063.

***Genetic subcluster:*** -

##### S1.2.1.10. The Kleopino/Kokorevo burial site (Tver)

**Geographic Information:** The barrow burial ground is located in the Staritsky district of the Tver region, 0.5 km north of the villages of Kokorevo and Klepino, on the banks of the Zhidokhovka River. There are 15 known barrows about 3-4 m high.

**Excavation History:** The excavations were carried out by L.N. Bastamov in 1879. He excavated 7 barrows.

**Summary of sampled materials:** Burials were found at the ground level of the barrow (in one instance on a clay-paved base oriented to the northeast, and in another, the upper part of the skeleton rested on a clay foundation). The skeletons were found in an extended supine position, with heads oriented to the west and southwest. Grave goods included temporal rings, pendants, silver and bronze rings, bronze bracelets, earrings, a finger ring, a leather belt with buckles (positioned around the skull of the deceased), a sabre hilt, and remnants of leather footwear. The burial practices and artefacts suggest ancient Rus’ cultural context. Dating: 11th-13th centuries CE^37,38^.

Genetic analysis was carried out on two skulls that are kept in the Museum of Anthropology, Lomonosov Moscow State University.

**AB18**

***Skeletal information:*** The skeletal remains belonged to a woman.

***Museum ID:*** 958.

***Genetic subcluster:*** -

**AB20**

***Skeletal information:*** The skeletal remains belonged to a woman.

***Museum ID:*** 965.

***Genetic subcluster:*** -

##### S1.2.1.11. The Kletnevo burial site (Vladimir)

**Geographic Information:** The burial ground is located in the Petushinsky district of the Vladimir region.

**Excavation History:** This site has not been introduced into the scientific discourse. The year and author of the excavations are unknown.

**Summary of sampled materials:** The burials were discovered at the ground level within the barrow. The skeletons were found in an extended supine position with the head oriented to the west. The accompanying grave goods included temporal rings, pendants, silver and bronze rings, bronze bracelets, earrings, and finger rings. The burial practices and artefacts suggest ancient Rus’ cultural context. Dating: 11th-13th centuries CE.

Genetic analysis was carried out on one skull that is kept in the Museum of Anthropology, Lomonosov Moscow State University.

**AB21**

***Skeletal information:*** The skeletal remains belonged to a man.

***Museum ID:*** 6700.

***Genetic subcluster:*** -

##### S1.2.1.12. The Pustosh Popova burial site (Kostroma)

**Geographic Information:** This ancient Rus’ burial ground is situated in the Krasnoselsky district of the Kostroma region, near the village of Antonovskoe, on the left bank of the Volga river, on the upper floodplain terrace.

**Excavation History:** Excavations were conducted by N.F. Nefedov, 1895.

**Summary of sampled materials:** The burial ground consisted of five barrows, each with stone perimeters around the barrows. All barrows were hemispherical in shape, ranging from 16 to 25 m in diameter and 1 to 2 m in height. All burials were single interments, located in the centre of the barrow at the subsoil level. The deceased were found in an extended supine position, with their heads oriented to the west-southwest. The left arm was placed across the chest, while the right arm was extended along the body. The grave goods included temporal rings, bell-shaped pendants, cross-shaped rhomboid pendants, twisted ribbed rings, an iron axe, and an iron knife. The burial practices and artefacts suggest ancient Rus’ cultural context. 12th-13th centuries CE^39^.

Genetic analysis was carried on a bone sample obtained from a single skull that is kept in the Museum of Anthropology, Lomonosov Moscow State University.

**AB37**

***Skeletal information****:* The specific association of skeletal remains with the burials investigated in the barrow is not determined. These remains were registered and stored in the Museum of Anthropology, Lomonosov Moscow State University under the name "Plyos". Anthropological identification of the skeletal remains has not been conducted.

***Museum ID:*** 1329.

***Genetic subcluster***: Rus_SWest.

##### S1.2.1.13. The Kremenye burial site (Moscow)

**Geographic Information:** The burial site was located in the Stupino district of the Moscow region, near the village of Kremenye, on the left bank of the Oka river, 9 km downstream from the Stupino town.

**Excavation History:** The burial ground became famous after the excavations of V.A. Gorodtsov^40,41^. Excavations at the burial ground were resumed in 2015 by Syrovatko A.S. following the discovery of synchronous ground-level cremation burials synchronous with the barrows.

**Summary of sampled materials:** The studied barrow was about 125 cm high, the diameter of the embankment within the contours of the ditch was about 11 m, the diameter of the robber's pit was about 3-3.5 m and its depth was about 1 m. Most likely this is barrow number 15 according to the passport of R.L. Rosenfeldt^42^.

On the northern side, a deep ditch was visible, and it was there that the burial was discovered. The most striking object was the ring groove encircling the central part of the barrow with a diameter (along the outer edge) of 5.5-5.9 m. Its filling was light-grey ash sand, similar to the buried soil, as well as fragments of calcined clay. In the centre of the there were two burials, a male burial, plundered by robbers, and a female burial found *in situ* in a burial pit. The remains of a woman were used in this study. The burial practices and artefacts suggest ancient Rus’ cultural context. Dating: the second quarter of the 12th century CE^43^.

**DR34. Barrow 15**

***Grave type****:* The burial is located in the centre of the barrow in a burial pit.

***Skeletal information****:* The skeleton was laying extended on its back, with the skull to the southwest, the facial part upwards, slightly turned to the right. The bones of the upper limbs were extended along the body. The skeletal remains belonged to a woman aged over 40 years.

***Grave good*s**: Temple copper rings, a copper bracelet, copper bells, carnelian beads, glass beads, copper rings.

***Genetic subcluster****:* Rus_Core.

##### S1.2.1.14. The Fili burial site (Moscow)

**Geographic Information:** The **burial site** is located on the right elevated bank of the Moscow river, within the modern city of Moscow, 700 m north of Bolshaya Filevskaya Street, on the grounds of the Filevsky park.

**Excavation History:** Excavations were conducted by V.A. Gorodtsov and B.A. Kuftin, and the materials were deposited in the Museum of Anthropology, Lomonosov Moscow State University in 1920.

**Summary of sampled materials:** All the barrows had a hemispherical shape, with a diameter of 5-8 m and a height of up to 1.5 m. Burials were situated beneath the barrow, in ground pits with a depth of 0.5-1 m. The skeletons were laid out in an extended position on their backs, oriented either to the west or southwest. Grave goods: the inventory includes advanced-type septifoil temporal rings with side loops and axe-shaped lobes, bronze twisted bracelets, bronze rings (broad-centred and lattice-patterned with triple-dot ornamentation), silver rings (pseudo-twisted and ribbed), remains of a leather pouch with a simple ring, carnelian beads (including 2 beads with white paste ornamentation), bilon pyramidal beads, spherical crystal pendants, rhomboid amber pendants, twisted glass bracelets, an iron sickle, and a ceramic pot with linear ornamentation. The burial practices and artefacts suggest ancient Rus’ cultural context. Dating: 11th-12th centuries CE^44–46^.

Genetic analysis was performed on bone samples obtained from two skulls that are kept in the Museum of Anthropology, Lomonosov Moscow State University.

**AB45**

***Skeletal information:*** The skeleton belonged to a man.

***Museum ID:*** 7219.

***Genetic subcluster: -***

**AB46**

***Skeletal information:*** The sex and age of the skeleton cannot be determined.

***Museum ID:*** 7225.

***Genetic subcluster: -***

##### S1.2.1.15. Yaroslavl Kremlin (Yaroslavl) burial site

**Geographic Information:** Yaroslavl is one of the major cities in medieval Rus', located on the right bank of the Volga River. Its foundation is connected with the reign of Yaroslav the Wise (late 10th - early 11th centuries CE), and the first mention in the *Primary Chronicle* of the city dates back to 1071. The oldest part of Yaroslavl, known as the "Rubleny Gorod" (the "Wooden Fortress") was located on a high promontory (Strelka) at the confluence of the Volga and Kotorosl rivers.

**Excavation History:** The paleoanthropological materials analysed in this study come from mass graves discovered within the Yaroslavl Kremlin during excavations conducted by the Yaroslavl expedition of the Institute of Archaeology, Russian Academy of Sciences, under the direction of A.V. Engovatova, from 2004 to 2021. The samples selected for analysis originate from two mass graves within the "Rubleny Gorod". Excavations of burials № 76 and № 110 were carried out between 2005 and 2007^44–46^.

**Summary of sampled materials:** Archaeological excavations in this area have uncovered the earliest buildings and remnants of a wooden-earthen fortification surrounded by a moat, which date back to the 11th century CE. In 1152, the city’s defences withstood a siege by the Volga Bulgars. In the early 13th century CE, under Prince Konstantin Vsevolodovich, the fortress was rebuilt, covering almost the entire Strelka. According to the *Laurentian Chronicle*^9^, a fire in 1221 initiated a new phase of construction. In the winter of 1238, the city was captured and destroyed by the Mongols.

Evidence of the city’s destruction in 1238 includes traces of a massive fire, stratigraphically dated to the first half of the 13th century CE, and mass graves found in areas that had previously been built upon. Numerous datable items were found in these Yaroslavl burials, including clothing elements and jewellery (ring-shaped temple rings, fire strikers, buckles, a stone four-pointed cross-enkolpion, glass beads, and fragments of glass bracelets), as well as household items made of wood, rope, felt, and ceramic vessels. AMS radiocarbon dating was conducted on 65 samples from nine sanitary graves of humans and animals, yielding calibrated dates in the range of 1233-1269 CE^47^.

All burials in these "mass graves" occurred simultaneously, representing the remains of those killed during the capture of the city in 1238. The anthropological series from Yaroslavl contains skeletal remains of more than 250 individuals.

The individuals selected for analysis originate from two mass graves within the "Rubleny Gorod". The samples Ya07, Ya12, Ya15, Ya28, Ya39, Ya40, Ya47, Ya71, Ya79 and Ya83 were taken from the burial № 110 (building №. 110), located near the fortifications and not far from the city gates. The individuals in building № 110 are represented mainly by scattered remains, as this sanitary burial was carried out several months after the tragic events of the capture of the city in February 1238.

One sample (MS) comes from the burial № 76, located on the slope of a filled-in moat from the 11th century CE, which had lost its defensive function by the late 12th - early 13th centuries CE and was repurposed for residential development^44,48,49^.

**Burial № 110 (Ya07, Ya12, Ya15, Ya28, Ya39, Ya40, Ya47, Ya71, Ya79, Ya83)**

***Grave type****:* The burial ground was found in a well (building № 110) that had suffered fire damage. The well is situated in close proximity to the fortification walls of the ancient city. In the central portion of the well, a number of wooden planks and logs were discovered within the well itself (Supplementary Fig. 20 top).

***Dating:*** 13th century CE.

***Skeletal information****:* In this mass grave, the most common remains were skulls of varying degrees of preservation, accompanied by fragments of frontal bones that corresponded to fragments of skulls. The sample from this burial comprises 66 individuals, representing all age groups. Of the individuals interred, 30 were male, 16 were female, 13 were children, and five were of indeterminate age and gender^44,47^ (Supplementary Fig. 20 below left and right).

The analysis of the anthropological remains revealed the presence of chopped, stabbed and incised wounds on various parts of the body, mainly in the skull area. This evidence indicates that the people died as a result of violent means^50^.

At least fourteen individuals of considerable height exhibited evidence of previous injuries.

***Grave goods****:* Clothing elements, ring-shaped temple rings, fire strikers, buckles, a stone four-pointed cross-enkolpion, glass beads, fragments of glass bracelets, rope, felt, and ceramic vessels.

***Genetic cluster:*** Rus_Cor (Ya07, Ya12, Ya15, Ya39, Ya47, YaYa71, Ya79); Rus_Middle (Ya28, Ya62); Rus_Baltic (Ya40, Ya83)**.**

**MS. Burial № 76**

***Grave type****:* This sample comes from the burial ground located on the slope of a filled-in moat from the 11th century, which was repurposed for residential development, from the excavation site "Rubleny Gorod” (2008).

***Dating:*** 13th century CE.

***Skeletal information****:* This skeleton was almost completely preserved, allowing for detailed analysis, including sex and age determinations. The remains belonged to a man aged 25-35^49^ (Supplementary Fig. 21).

***Grave goods****:* Leather boots were preserved on the lower leg bones, with a design typical of nomadic footwear^44,48^.

***Genetic subcluster:*** Rus_Middle.

##### S1.2.1.16. Pereslavl-Zalessky burial site

**Geographic Information**: Pereslavl-Zalessky (Pereslavl) is one of the cities of the Rostov-Suzdal land, located on the shore of Lake Pleshcheyevo, where the Trubezh river flows into it.

**Excavation History:** The first paleoanthropological studies of the city were started in 2012 by researchers from the Institute of Archaeology of the Russian Academy of Sciences in connection with the rescue archaeological excavations of the necropolises of the 13th century CE^51^. In addition, in 2016, during the security archaeological excavations of the Institute of Archaeology of the Russian Academy of Sciences, a mass grave was discovered in the basement of a burnt-out building, This burial is dating 13th century CE, and currently its paleoanthropological materials are the earliest in the Pereslavl^52^. This mass burial has similarities to collective burials in Yaroslavl, they were left behind after the invasion of Batu Khan in 1238 and, in fact, are a temporary slice of the genetic composition of the urban population before the Tatar-Mongol invasion^53^. For genetic analysis, samples were selected from burials uncovered during rescue archaeological excavations between 2013 and 2016 at three sites in the central fortified part of the city.

**Summary of sampled materials:** The city was founded in the mid 12th century CE by Prince Yuri Vladimirovich Dolgorukiy. In the 13th century CE it was the centre of the independent Pereslavl principality. After this time, Pereslavl-Zalessky became part of the Grand Duchy of Vladimir and had political union with the Moscow Prince. The mediaeval city comprised a fortified area, enclosed by earthen ramparts, covering 35 hectares, and an unfortified settlement (posad), the boundaries of which have not been determined^54^.

Genetic analysis was performed on bone samples obtained from nine skulls that are kept in the Museum of Anthropology, Lomonosov Moscow State University.

**Pereslavl XIII c.**

This burial ground comprises two graves located on the territory of the Kremlin. One of the graves was a complex consisting of two burials and was found in the basement of a destroyed building, while the other was a massive grave.

**Pereslavl-Zalessky, Sovetskaya Street 37. Burials from structure 22**

**AB209. Structure 22. individual 1**

***Grave type****:* The burial is located 230 cm below the modern surface, in the cellar pit (structure 22, in the northern half, on the boundary between two rooms of the building), as part of a complex consisting of two burials. It is apparent that by the time the burials were made, the building had already been destroyed, and the burials were placed in its pit.

***Dating:*** 13th century CE.

***Skeletal information****:* The skeleton was found lying on his back, with his head oriented northwest. The head was in an unnatural position. The hands were placed in the abdominal area. The temporal bone was damaged, showing evidence of a violent death. The size of the wound openings, caused by depressed fractures, measured 3x2.7 cm and 3x3 cm. These characteristics suggest the use of a combat mace (kisten), with a teardrop-shaped weight of up to 3 cm in diameter, with a smooth surface, without spikes or reinforcing ribs. The burial belonged to an elderly man.

***Grave goods****:* An iron belt buckle (in the area of the pelvic bones).

***Genetic subcluster****:* Rus_Core.

**AB210. Structure 22. individual 2**

***Grave type****:* The burial is located 230 cm below the modern surface, in the cellar pit (structure 22, in the northern half, on the boundary between two rooms of the building), as part of a complex consisting of two burials. It is apparent that by the time the burials were made, the building had already been destroyed, and the burials were placed in its pit.

***Dating:*** 13th century CE.

***Skeletal information****:* The skeleton was lying extended on its back. Preservation was incomplete. The frontal bone was damaged, and the size of the wound openings, caused by depressed fractures, measured 3x2.7 cm and 3x3 cm. These features also suggest the use of a combat mace with a teardrop-shaped weight, up to 3 cm in diameter, smooth-surfaced, without spikes or reinforcing ribs. The burial belonged to a child.

***Grave goods****:* No grave goods were found.

***Genetic subcluster****:* Rus_Core.

**Pereslavl-Zalessky, mass grave at 10a Komitetskaya Street**

**AB211. Burial 35**

***Grave type****:* The skeletal remains were found in a large burial pit, arranged in the basement of a surface structure without anatomical order. This burial was a mass grave containing the remains of at least 99 individuals, of these, 80 are adults and 19 children.

***Dating:*** 13th century CE.

***Skeletal information****:* The bones of different skeletons were intermingled. A number of bones, including 11 skulls, showed signs of trauma without evidence of healing.

***Grave goods****:* Various household items and jewellery, including fragments of glass bracelets, were found in the fill of the pit.

***Genetic subcluster****:* Rus_Baltic.

**AB212. Burial 18**

***Grave type****:* The mass grave.

***Dating:*** 13th century CE.

***Skeletal information****:* The skeleton belonged to a man.

***Additional information:*** Radiocarbon date: (UGAMS-61311) – 1041-1213 calBC (95.4%)^54^.

***Genetic subcluster****:* Rus_Core.

**AB 213. Burial 30**

***Grave type****:* The mass grave.

***Dating:*** 13th century CE.

***Skeletal information****:*The skeletal remains belonged to a man.

***Genetic subcluster****:* Rus_SWest.

**AB214. Burial 52**

***Grave type****:* The mass grave.

***Dating:*** 13th century CE.

***Skeletal information****:* The skeletal remains belonged to a man.

***Additional information:*** Radiocarbon date: (UGAMS-61312) – 1052-1263 calBC (95.4%)^54^. ***Genetic subcluster****:* Rus_Core.

**Pereslavl XV-XVI c.**

**Pereslavl-Zalessky, Komitetskaya Street, 22. Burials from Excavation Site III**

**AB205. Burial 68**

***Grave type****:* The burials were located at a depth of 80-120 cm below the surface level, with the grave pits cut into the cultural layer but not clearly traceable.

***Dating:*** 15th-16th centuries CE.

***Skeletal information****:* The skeleton was found lying extended on its backs, with its heads oriented towards the north-northwest. Its hands were positioned on the chests. The skeleton belonged to a man.

***Genetic subcluster****:* Rus_Core.

**AB206. Burial 214**

***Grave type****:* The burials were located at a depth of 80-120 cm below the surface level, with the grave pits cut into the cultural layer but not clearly traceable.

***Dating:*** 15th-16th centuries CE.

***Skeletal information****:* The skeleton was found lying extended on its backs, with its heads oriented towards the north-northwest. Its hands were positioned on the chests. The skeleton belonged to a man.

***Genetic subcluster****:* Rus_Core.

**AB207. Burial 122**

***Grave type****:* The burials were located at a depth of 80-120 cm below the surface level, with the grave pits cut into the cultural layer but not clearly traceable.

***Dating:*** 15th-16th centuries CE.

***Skeletal information****:* The skeleton was found lying extended on its backs, with its heads oriented towards the north-northwest. Its hands were positioned on the chests. The skeleton belonged to a man.

***Genetic subcluster****:* Rus_Core.

##### S.1.2.1.17. Old Ryazan burial site

**Geographic Information:** Old Ryazan was the capital of the Ryazan principality until it was destroyed by the Mongols in 1237. It is located on the right (eastern) bank of the middle course of the Oka river.

**Excavation History:** The first excavations at the site of Old Ryazan were started by A.V. Selivanov in 1888. However, large-scale excavations in Old Ryazan began in 1926 under the direction of V.A. Gorodtsov and then excavations of Old Ryazan were resumed only in 1946.

**Summary of sampled materials:** The city's history spans a little over three centuries: from its emergence in the first third of the 11th century CE as a small princely fortress on the eastern borders of Rus' to its peak in the mid-12th to early 13th centuries CE, followed by a century of decline after its destruction by the Mongol forces of Batu Khan in the winter of 1237. The ancient settlement of Old Ryazan consisted of two parts: the southern settlement and the northern one. The seven samples used in this study correspond to two distinct periods in the city’s history, which differ not only chronologically but also culturally. Burials from excavations 13 and 17, located in the southern settlement of Old Ryazan, belong to the early urban necropolis of the 11th to early 12th centuries CE. This extensive necropolis lay beyond the walls of the Northern Settlement, the oldest part of the city. During this period, Ryazan was a small fortress on the eastern edge of the ancient Rus' state, formed with the active involvement of princely authority. This is reflected in the diverse composition of its population. Based on an analysis of cultural traditions, the early settlers originated from the Middle and Upper Dnieper regions, as well as the Dvina basin.

The burials in the Northern Settlement belong to a typical urban Christian burial ground from the 12th to 13th centuries CE, a time of Ryazan’s prosperity. In this period Old Ryazan was the capital of an independent principality and is known to have several cemeteries that existed within the settlement in this time (both in the oldest northern section and in the southern part). These cemeteries were likely associated with separate parish churches established on a territorial basis. The city’s rapid growth during this period was undoubtedly linked to an influx of new inhabitants, which significantly changed the demographic composition of the city. Thus, the second period in Ryazan’s history, like the earlier one, is characterised by a diverse population with a significant number of newcomers.

However, in the Northern Settlement, a certain degree of continuity from earlier times persisted, especially in the preservation of the city's layout and the estate plots. The burials in the second (later) group can be considered a continuation of the first, with the burial ground in the Northern Settlement serving as the necropolis for the descendants of those who had been buried in the early necropolis a century earlier^55^.

Different burial rites were observed in various parts of the Old Ryazan complex. Samples were taken for analysis both from the barrows in the Southern Settlement (excavations 13 and 17) and from the collective grave associated with the 1238 invasion of Batu Khan in the eastern part of the Northern Settlement^56–58^.

Genetic analysis was performed on bone samples obtained from seven skulls that are kept in the Museum of Anthropology, Lomonosov Moscow State University.

**AB142. Excavation 13. Burial 96a**

***Grave type****:* A double burial was discovered in a subrectangular soil pit, within a destroyed barrow, in the coastal part of the Southern Settlement.

***Dating:*** The second half of the 11th- the first half of the 12th centuries CE.

***Skeletal information****:* The skeletons were laid out extended on their backs, with their arms along their bodies and their heads to the west.

***Grave goods****:* An iron belt buckle from the male burial.

***Museum ID:*** KO 355-1.

***Additional information:*** expedition of the IA of the USSR Academy of Sciences, head V.P. Darkevich, excavations by Khatuntsev P.B. in 1977-1978.

***Genetic subcluster****:* Rus_Core.

**AB143. Excavation 13. Burial 101**

***Grave type****:* The burial was discovered in a shallow subrectangular soil pit, within a destroyed barrow, in the coastal part of the Southern Settlement.

***Dating:*** The second half of the 11th to the first half of the 12th centuries CE.

***Skeletal information****:* The skeleton was lying extended on its back, with arms along the body, and head facing west. The skeletal remains belonged to a woman.

***Grave goods****:* No grave goods were found.

***Museum ID:*** KO 355-2.

***Additional information:*** Expedition of the IA of the USSR Academy of Sciences, head V.P. Darkevich, excavations by Khatuntsev P.B. in 1977-1978.

***Genetic subcluster****:* Rus_SWest.

**AB144. Excavation 17. Burial 24**

***Grave type****:* The burial was discovered in a shallow subrectangular soil pit, within a destroyed burial in the barrow, on the territory of the Southern Settlement.

***Dating:*** The second half of the 11th to the first half of the 12th centuries CE.

***Skeletal information****:* The skeleton was lying extended on its back, with arms along the body and head facing west. Beneath the lower jaw and skull were remains of gold-threaded braid with a zigzag pattern of Byzantine origin The skeleton belonged to a man.

***Grave goods****:* No grave goods were found.

***Museum ID:*** KO 355-3.

***Additional information:*** Expedition of the IA of the USSR Academy of Sciences, head V.P. Darkevich, excavations by Khatuntsev P.B. in 1977-1978.

***Genetic subcluster****:* -

**AB146. Excavation 17. Burial 26**

***Grave type****:* The burial was discovered in a rectangular pit.

***Dating:*** The second half of the 11th to the first half of the 12th centuries CE.

***Skeletal information****:* The skeleton was lying extended on its back, with arms along the body and head facing west. The skeletal remains belonged to a man.

***Grave goods****:* No grave goods were found.

***Museum ID:*** KO 355-6.

***Additional information:*** Expedition of the IA of the USSR Academy of Sciences, head V.P. Darkevich, excavations by Khatuntsev P.B. in 1977-1978.

***Genetic subcluster****:* Rus_Core.

**AB147**

***Grave type****:* The burial was found in a Christian burial ground near the Spassky Cathedral within the city during excavations led by Selivanov in 1888. The burial was in a rectangular soil pit, within a wooden coffin.

***Dating:*** The second half of the 11th to the first third of the 13th centuries CE.

***Skeletal information****:* The remains of the skeleton belonged presumably to a woman.

***Museum ID:*** 6736.

***Additional information:*** Excavations by Selivanov A.V. in 1888.

***Genetic subcluster****: -*

**AB148. The Excavation 7. Burial 8. Trench 7**

***Grave type****:* The burial was found in a four-sided "ossuary pit" in the eastern part of the Northern Settlement, along with 14 other burials. The excavation diary by V.A. Gorodtsov notes the remains of wood discovered in the pit (possibly coffin remains). According to Gorodtsov's observations, this was a mass grave associated with the events of December 1237, when the capture and destruction of the capital of the Ryazan principality by Khan Batu.

***Dating:*** December, 1237.

***Skeletal information****: -*

***Grave goods****:* 3 spherical bells with a slit, used as buttons were found in the burial.

***Museum ID:*** 8420.

***Additional information:*** Excavations by Gorodtsov V. A. in 1926.

***Genetic subcluster****:* Rus_Swest.

**AB149. The Excavation 2a. Layer 3. Burial 45**

***Grave type****:* Located in the eastern part of the Northern Settlement, in close proximity to 12th-13th centuries CE dwellings. The burial was found in a shallow soil pit.

***Dating:*** 12th -the early 13th centuries CE.

***Skeletal information****:* The skeleton was lying extended on its back, with the head facing west or southwest. The hands were placed on the chest. The skeleton belonged to a man. No special features or pathologies were identified.

***Grave goods****:* No grave goods were found.

***Museum ID:*** 10255.

***Additional information:*** The Staroyazanskaya expedition, excavations by Mongate A.A. in 1946-1950.

***Genetic subcluster****: -*

#### S1.2.2. Archaeological sites of the North-Western Rus' region (Novgorod)

##### S1.2.2.1. St. George’s (Yuriev) Monastery burial site

**Geographic Information:** St. George's (Yuriev) Monastery is situated in Veliky Novgorod, one of the oldest cities in Rus', located in the northwestern part of the European plain, approximately 6 kilometers from Lake Il'men. The monastery is located on the southern periphery of the city, on the banks of the Volkhov River.

**Excavation History:** The necropolis within the St. George's Cathedral of the monastery was unearthed in the 1930s by M.K. Karger. However, the skeletal remains were not preserved. Between 2015 and 2023, the Novgorod Archaeological and Architectural Expedition, led by Vladimir V. Sedov, discovered and excavated the necropolis areas surrounding the cathedral. Excavations took place on the northern and southern sides of the cathedral, with the eastern side being partially explored and the western side remaining unexcavated.

**Summary of sampled materials:** The St. George's Cathedral was founded in 1119 and completed around 1130 by order of Prince Vsevolod Mstislavich. Initially, the cathedral was probably the burial site of princes. However, after the expulsion of Prince Vsevolod in 1136, the monastery came under the jurisdiction of the Novgorod [Veche (popular assembly)](https://www.prlib.ru/en/section/683143). From the end of the 12th to the middle of the 15th centuries CE, the cathedral became the resting place of the service princes of Novgorod. Additionally, two boyars from the Miroshkinichi clan were buried here in the late 12th and early 13th centuries CE.

In total, around one hundred burials were identified around the cathedral, including burials in stone and plinth (brick-like) sarcophagi. These composite sarcophagi, consisting of six slabs, date back to the pre-Mongol period, specifically to the 12th and early 13th centuries CE. Most of these sarcophagi were constructed in this period, although their upper chronological limit may extend to the mid-13th century CE.

A total of eight stone sarcophagi and one plinth sarcophagus have been identified. The burials in these sarcophagi may be linked to boyar families, as female interments are also present. An added complexity is that, in some cases, later family members or more distant relatives (perhaps dating to the 14th century CE) were reburied by opening the lids of earlier sarcophagi. Among these, there are also sarcophagi belonging to monks (e.g., sarcophagus 1), who were reburied with some rationale alongside earlier burials, potentially based on their positions within the monastic community (such as abbot, steward, etc.) (Supplementary Fig. 22).

Individuals from six sarcophagi №№72-77 were taken into the study.

In addition to sarcophagus burials, the area around the cathedral revealed burials in wooden coffins placed under large stone slabs. In some cases, these may be considered family plots beneath several slabs. These burials and slabs are dated to the 14th-15th centuries CE^60–63^.

**Burial 72. Sarcophagus 7. Individual 3-4 (DR8, DR9)**

A white stone sarcophagus along the southern wall of St. George's Cathedral contained the remains of four individuals, but only Individual 3 (DR9) and Individual 4 (DR8) were used in this study. A datable artefact - earrings - allowed the female burial within the sarcophagus to be assigned to the second half of the 13th to the first half of the 14th centuries CE^59–62^ (Supplementary Fig. 23).

**DR8. Individual 4**

***Grave type****:* Sarcophagus

***Skeletal information****:* Disarticulated bones of a previously interned child aged 10-12 years shifted within the sarcophagus.

***Genetic subcluster****:* Rus_Baltic.

**DR9 . Individual 3**

***Grave type****:* Sarcophagus.

***Dating:*** The second half of the 13th - the first half of the 14th centuries CE.

***Skeletal information****:* This skull of the individual 3 was found at the feet of Individual 1 and belonged to a woman aged 20-25 years.

***Grave goods****:* Earrings.

***Genetic subcluster****:* Rus_Baltic.

**Burial 73. Sarcophagus 8. Individual 1-3 (DR49, DR6, DR7)**

Upon removal of the sarcophagus lid, the burial, wrapped in birch bark, was revealed.

The sequence of burials was as follows: Individual 4 was buried first (was not used in this study), followed by the woman (DR6) and child (DR7), and finally Individual 1 (DR49) was placed on top (Supplementary Fig. 24). Dating: 12th-the middle of the 13th centuries CE.

**DR6. Individual 2**

***Grave type****:* Sarcophagus

***Skeletal information****:* The skeleton was positioned along the northern wall and belonged to a woman aged 20-30 years.

***Additional information:*** The skeleton had an infant in its lap, although this child was not assigned a separate number.

***Genetic subcluster****: -*

**DR7. Individual 3**

***Grave type****:* Sarcophagus

***Skeletal information****:* The skeleton belonged to a child aged 8-9 years, located in the southern side of the eastern part of the sarcophagus.

***Genetic subcluster****:* Rus_SWest.

**DR49. Individual 1**

***Grave type****:* Sarcophagus

***Skeletal information****:* Skeleton was lying on its back, hands folded at a slight angle with palms resting on the lower abdomen and was displaced south of the sarcophagus axis. The skull rested on its occipital part, and soft tissue remnants were preserved in the torso area. Notable injuries include a healed fracture of the left ulna and a nasal fracture. The skeleton belonged to a man aged 50-60 years old.

***Grave goods****:* The deceased was wrapped in birch bark both above and below the body.

***Additional information:*** On the lower level of the same sarcophagus, three more individuals were discovered.

***Genetic subcluster****: -*

**Burial 74 (burial in a pit). Individual 2 (DR41)**

This burial contained the skeletons of four individuals: Individual 1 - a man over 35 years, Individual 2 - a newborn, Individual 3 - a man over 55 years old and Individual 4 - an infant, approximately one year old, located at the feet. Only Individual 2 (DR41) was used in this study (Supplementary Fig. 25).

Between the burials of Individual 1 and Individuals 3 and 4, the bottom of a coffin was identified, indicating that Individuals 1 and 2 were buried together in one coffin, placed on top of Individuals 3 and 4, who were in another coffin. The burials contained no artefacts, and only the lower parts of the skeletons were preserved below the pelvis. Dating: 14th–15th centuries CE.

**DR41. Individual 2**

***Grave type****:* Wooden coffin.

***Dating:*** 14th-15th centuries CE.

***Skeletal information****: The* s*keleton belonged* to a newborn.

***Grave goods****:* No grave goods were found.

***Additional information:*** The skeleton is located together with a man in one coffin, which was located on top of another coffin.

***Genetic subcluster****:* Rus_Core.

**Burial 75. Sarcophagus 9. Individual 3-4 (DR48, DR38)**

This sarcophagus contained the remains of several people. In the western part under the lid, birch bark wrapping was preserved, while in the eastern part, the remains of Individual 1 and below the birch bark, which separated the upper individual from the lower ones, the burial of Individual 2. Below, the remains of two more individuals were found: Individual 3, near the northern wall, and Individual 4, near the southern wall. Two more artefacts were found in the lower level: a bronze bell and a textile ribbon. It appears that Individuals 3 and 4 were the first to be buried, after which their remains were moved to accommodate Individual 2, and finally, Individual 1 was placed on top. Dating: 12th-the middle of the 13th centuries. Only Individual 3 (DR48) and Individual 4 (DR38) were used in this study (Supplementary Fig. 26).

**DR38. Individual 4**

***Grave type****:* Sarcophagus.

***Skeletal information****:* The skeleton belonged to a man aged 50-60 years.

**DR48. Individual 3**

***Grave type****:* Sarcophagus.

***Skeletal information****:* The skeleton belonged to a man aged 35-45 years.

***Genetic subcluster****: -*

**Burial 76. Sarcophagus 10. Individual 3 and Individual 4 (DR4, DR5)**

The slabs covering this sarcophagus were shattered, and the remains inside were disturbed. In the western part of the upper layer, four skulls were found. The lower layer contained the remains of four individuals. Only Individual 3 (DR4) and Individual 4 (DR5) were used in this study. Dating: 12th-the middle of the 13th centuries CE (Supplementary Fig. 27).

**DR4. Individual 3**

***Grave type****:* Sarcophagus.

***Skeletal information****:* The skeleton belonged to a man over 60 years old.

***Genetic subcluster****:* Rus_SWest.

**DR5. Individual 4**

***Grave type****:*Sarcophagus.

***Skeletal information****:* The skeleton belonged to a man aged 35-49 years, lying on his back. His right arm was slightly bent, resting on the pelvis, while only the humerus of the left arm remained. To place Individual 4, some remains were displaced northwards, and others to the south. Under the body of Individual 4, traces of organic decomposition were observed. His legs were extended and covered with birch bark.

***Additional information:*** The lower level contained the skeletons of three people.

***Genetic subcluster****:* Rus_Core.

**Burial 77. Sarcophagus 11. Individual 1-3 (DR1, DR2, DR3)**

This sarcophagus contained the remains of several people. The upper skeleton belonged to Individual 1 (DR1), At its feet, the remains of Individual 2 (DR2) On the lower level, bones were found in the eastern part of the sarcophagus and in the southwest corner, the remains of Individual 3 (DR3). Dating: 12th-the middle of the 13th centuries CE (Supplementary Fig. 28).

**DR1. Individual 1**

***Grave type****:* Sarcophagus.

***Skeletal information****:* The skeleton belonged to a woman aged over 50 years old.

***Additional information:*** The upper skeleton was covered with birch bark.

***Genetic subcluster:*** Rus_Core.

**DR2. Individual 2**

***Grave type****:* Sarcophagus.

***Skeletal information****:* The skeleton belonged to a child aged 4-5 years.

***Genetic subcluster: -***

**DR3. Individual 3**

***Grave type****:* Sarcophagus.

***Skeletal information****:* The skeleton belonged to a man over 50 years old.

***Genetic subcluster: -***

##### S1.2.2.2. The Khreple burial site (Novgorod)

**Geographic Information:** The burial ground is located on the right bank of the Khrepelka river (a tributary of the Luga river), surrounded by the structures of the old village of Khreple in the Leningrad region (present-day Novgorod region).

**Excavation History:** The site was investigated by A.V. Artsikhovsky in 1929.

**Summary of sampled materials:** All the barrows were encircled by stones. The barrows were hemispherical in shape, ranging in diameter from 1.6 m (questionable measurement) to 13 m, and from 0.42 to 1.69 m in height. The burial ground contained both individual and paired primary burials, as well as secondary burials in the barrow. All burials were found in rectangular grave pits, laid on their backs with their heads oriented westward. The burial practices and artefacts suggest ancient Rus’ cultural context dating: 11th-12th centuries CE, based on the finds of coins in the barrow № 20, which included Arab dirhams attributed to Emir Ismail, son of Ahmed (895 CE) and Emir Nasr, son of Ahmed, minted in Samarkand in 923 CE. Western European denarii from the 11th century, minted at the Lower Lorraine Abbey of Stablo, were also found in the barrow № 12. The general cultural appearance of the burial ground is distinctly Rus’/East Slavic^63^.

Genetic analysis was carried out on bone samples obtained from three skulls, which are kept in the Museum of Anthropology, Lomonosov Moscow State University.

**AB47. Barrow 7**

***Grave type****:* The burial was discovered under the barrow, encircled by a stone ring, in a rectangular grave pit.

***Dating:*** 11th-12th centuries CE.

***Skeletal information****:* The skeleton was in an extended supine position, with the head oriented to the west. The remains belonged to a man aged 30-39 years. There were signs of a fatal injury on the skull, likely caused by a strike with a bladed weapon with a broad cutting edge to the occipital region. The wound measured approximately 70 mm in length, with no signs of healing. An additional trauma was observed on the lower third of the ulnar bone, featuring an oval cut (40x9 mm) with smooth edges and no signs of inflammation or healing. This injury may have occurred at the same time as the skull wound, suggesting an attack.

***Grave goods****:* No grave goods were found.

***Museum ID:*** 7850.

***Genetic subcluster****:* Rus_Baltic.

**barrow 7. Paired Burial**

***Grave type****:* This paired burial was discovered under the barrow, surrounded by a stone ring, in a rectangular grave pit and had two Individuals.

***Dating:*** 11th-12th centuries CE.

***Skeletal information****:* Both skeletons were lying in an extended supine position with their heads oriented westward.

***Grave goods****:* No grave goods were found.

**AB48. Individual 1**

***Skeletal information****:* The skeletal remains belonged to a man.

***Museum ID:*** 8321.

***Genetic subcluster****:* Rus_Baltic.

**AB49. Individual 2**

***Skeletal information****:* The skeletal remains belonged to a man.

***Museum ID:*** 8318.

***Genetic subcluster****:* Rus_Core.

#### S1.2.3. Archaeological sites from the Western Rus' Dnieper-Dvina region

##### S1.2.3.1. The Volochok burial site (Smolensk)

**Geographic Information:** The burial ground is located near the village of Aleksino, Dorogobuzhsky district, Smolensk region.

**Excavation History:** The materials were first published by A.A. Spitsyn in 1892. Eight burial barrows have been excavated to date.

**Summary of sampled materials:** The burials were predominantly placed in the subsoil layer, less frequently in a soil pit or within the barrow itself. Single inhumations are most common, though there are two instances of double burials (man and woman, man and child). All skeletal remains are oriented with the head towards the west. The body position is supine (lying on the back) with the skull either facing straight upwards or turned to the right (occasionally to the left). Traces of coffins have only been documented in one burial barrow. In the majority of burials, evidence of bedding (wooden remains) was recorded. Pots or iron-banded wooden buckets were placed at the feet of the deceased. The funerary inventory of male burials includes iron knives, buckles, and bronze buttons. The grave goods found in female burials comprise temple rings, necklaces, bracelets, rings, carnelian, crystal, and silver beads, bead necklaces, wire and plate bracelets with open ends, lunulae (crescent-shaped pendants), and round plaques. The specific association of bone samples selected for analysis to individual burials within the burial ground could not be established. The burial practices and grave goods suggest ancient Rus’ cultural context. Dating: 11th-12th centuries CE^64^.

Genetic analysis was carried out on a bone sample obtained from a single skull that is kept in the Museum of Anthropology, Lomonosov Moscow State University.

**AB6**

***Skeletal information:*** The skeletal remains belonged to a woman.

***Museum ID:*** 1371.

***Genetic subcluster****:* Rus_SWest.

##### S1.2.3.2. The Iput burial site (Bryansk)

**Geographic Information:** The burial ground is located on the floodplain terrace of the Iput river, in the Surazhsky district, Bryansk region.

**Excavation History:** The excavations were carried out by P. M. Eremenko in 1891 and 1894; a total of 160 burial barrows were examined, comprising 15 burial barrow groups located in different places along the Iput river. In total, 61 male and 65 female burials were found in the burial barrows during Eremenko’s excavations, and 37 burials remain unidentified.

**Summary of sampled materials:** Most of the barrows had a hemispherical shape, while some were elongated, with diameters ranging from 25 to 38 m and heights of up to 1 to 3 m, often featuring well-defined ditches around them. All burials were performed at the level of the ancient ground surface, at the base of the barrow, on an ash layer up to 8-12 cm thick and 21-35 m in diameter. Predominantly, single inhumations were observed, though double burials were less commonly documented. The skeletal remains were found in an extended supine position, oriented with the head to the west, and arms laid along the body. In the male burials, rectangular and round iron belt buckles, rings, whetstones, iron bits with round rings on the ends and curved lighters were found. Female burials contained seven-lobed temple rings, barrel-shaped beads of gold and glass, fourteen-sided square-shaped carnelian beads, neck rings, bronze rings with smaller rings suspended from them, and bracelet-shaped temple rings with S-shaped spirals at the ends. The general cultural appearance of the burial ground is distinctly Rus’/East Slavic. The burial ground is dated to the 11th century CE^65^. Genetic analysis was carried out on a bone sample obtained from a single skull that is kept in the Museum of Anthropology, Lomonosov Moscow State University.

**AB53**

***Skeletal information****:* The bone remains belonged to a man.

***Museum ID:*** 2100.

***Genetic subcluster****:* Rus_SWest.

##### S1.2.3.3. The Logoysk burial site (Minsk)

**Geographic Information:** The burial ground of Logoysk, also known as Greben, dating to the Rus’ period, is situated near the village of Vidogoshche, in proximity to the contemporary town of Logoysk in the Minsk region.

**Excavation History:** The site was excavated by Count K.P. Tyshkevich in 1866.

**Summary of sampled materials:** All barrows had a hemispherical shape, with a diameter ranging from 15 to 30 m. The burials were found at the ground level, in an extended supine position, oriented from northwest to southwest. The grave goods included glass beads, buckles, bells, bracelets, and wire rings. The burial practices and grave goods suggest ancient Rus’ cultural context. /Dating: 11th-12th centuries CE^66,67^.

Genetic analysis was carried out on a bone sample obtained from a single skull that is kept in the Museum of Anthropology, Lomonosov Moscow State University.

**AB13. Barrow 11, Skull 7**

***Grave type:*** The barrow had a diameter of 23 m. The burial was found at the ground level, in an extended supine position, with the head oriented southwestward.

***Dating:*** 11th-12th centuries CE.

***Genetic subcluster****: -*

##### S1.2.3.4. The Seltso burial site (Tver)

**Geographic Information:** The burial ground is located in the basin of the Western Dvina river, on the right bank of the Ushitsa river, a tributary of the Velesa river, in the Zapadnodvinsky district of the Tver region (previously part of Belsky uyezd, Smolensk Governorate).

**Excavation History:** The burial ground was excavated by K.A. Gorbachev in 1886, who investigated a total of 20 barrows in the area.

**Summary of sampled materials:** All the barrows in the burial ground had a hemispherical shape, reaching up to 5 m in height. The burials were single interments situated in the central part of the barrow’s base, at the subsoil level. The deceased were found in an extended supine position, with their heads oriented to the east. The arms were positioned along the body, and the skull was usually lying straight, though occasionally turned to the left side. The grave goods included lobed temporal rings (of the "Seltso" type), temporal rings with tied ends, twisted torcs, beads, glass bead strings, round pendants with crosses, metal beads, and ceramic vessels. The precise association of the skeletal remains with specific burials investigated in the burial ground has not been established. The burial practices and grave goods suggest ancient Rus’ cultural context. Dating: 11th-12th centuries CE^68–70^.

Genetic analysis was carried out on two skulls that are kept in the Museum of Anthropology, Lomonosov Moscow State University.

**AB40**

***Skeletal information:*** The sex and age of the skeleton cannot be determined.

***Museum ID:*** 1423.

***Genetic subcluster: -***

**AB41**

***Skeletal information:***

***Additional information:*** Museum № 1431.

***Genetic subcluster: -***

***Skeletal information:*** The hands were positioned near the head. The sex and age of the skeleton is undetermined.

***Museum ID:*** 1915.

***Genetic subcluster: -***

##### S1.2.3.5. The Gnevkovo burial site (Smolensk)

**Geographic Information:** The burial ground was located 0.7 km southwest of the village of Gnevkovo, Shumyachsky district, Smolensk region, on the right bank of the Ostra river. **Excavation History:** There are records of 93 barrows in this area. In 1960-1961, V.V. Sedov excavated 35 of these barrows.

**Summary of sampled materials:** All the kurgans had a hemispherical shape, with diameters ranging from 2 to 11 m and heights from 0.15 to 2.25 m. The burials were predominantly single (occasionally paired), found primarily in soil pits, less frequently in the subsoil layer. The deceased were placed in extended supine positions, with their heads oriented west-northwest. The male burials lacked grave goods, whereas the female burials were accompanied by ring-shaped temporal rings, bronze bracelets, and glass beads. The burial practices and grave goods suggest ancient Rus’ cultural context. Dating: 12th-13th centuries CE^71^.

Genetic analysis was carried out on two skulls that are kept in the Museum of Anthropology, Lomonosov Moscow State University.

**AB11. Barrow 18**

***Grave type****:* The skeleton was discovered in a rectangular grave pit.

***Dating:*** 12th-13th centuries CE.

***Skeletal information****:* The skeleton was oriented with the head to the west. The body was positioned extended on its back. The skeletal remains belonged to a man.

***Museum ID:*** 958.

***Genetic subcluster:*** -

**AB12. Barrow 19. Burial 9**

***Grave type****:* Two burials are known to have been made in grave pits beneath the barrow. One, located in the central part, belonged to an adolescent. The second, found in the southern part of the barrow, belonged to an adult man.

***Dating:*** 12th-13th centuries CE.

***Skeletal information****:* Both bodies were lying extended on their backs, with their heads to the northwest and arms positioned along the body.

***Additional Information:*** The association of skeletal remains with a specific burial within the barrow has not been established.

***Museum ID:*** 11457.

***Genetic subcluster:*** -

#### S1.2.4. Burial grounds from the Southern Rus' region

##### S1.2.4.1. The Komarovka burial site (Kursk)

**Geographic Information:** The necropolis is located at the Komarovka settlement, which sits on the left bank of the Snagost river, a left tributary of the Seym river. This site is located 1.3 km southwest of the village of Komarovka in the Kursk region. The settlement is situated on the northeastern edge of a steep terraced slope on a subrectangular platform measuring 35x30 m. The platform rises 10-11 m above the floodplain. On the southern side, the settlement was fortified by two ramparts, 0.1-0.5 m in height, and two ditches, 0.4-0.6 m deep.

**Excavation History:** In 2019, a team led by N.A. Birkina from the State Historical Museum (SHM) investigated 13 inhumation burials, which cut through the cultural layer. Cultural layers from both the Roman culture and Rus’ period (12th-13th centuries CE) have been identified at the site.

**Summary of sampled materials:** Some of these burials were damaged or destroyed by plowing. Few grave goods were found, including a fragment of gold-embroidered textile (Burial 12) and appliqué plaques (Burial 10). The general cultural appearance of the burial ground is distinctly Rus’/East Slavic (rural necropolis). Dating: Second half of the 12th- 14th centuries CE^72^.

**DR23. Burial 2**

**Grave type:** The burial is oriented southwest–northeast. The grave pit was not identified, and the dimensions of the burial remain unknown.

***Dating:*** The second half of the 12th-beginning of 14th centuries CE.

***Skeletal information:*** The skeleton was lying in an extended supine position, with the head facing northwest. A stripe of wood decay measuring 0.5x0.1 m and up to 5 cm thick was observed along the northern side of the skeleton. The remains probably belonged to a woman aged 45-55 years.

***Genetic subcluster:*** Rus_Swest.

**DR24. Burial 4**

***Grave type:*** A ground burial oriented southwest–northeast. The grave pit was not identified, and the dimensions of the burial are unknown.

***Dating:*** The second half of the 12th-14th centuries CE.

***Skeletal information:*** The skeleton was lying in an extended supine position. The remains belonged to a man over 50 years old.

***Grave goods:*** No grave goods were found.

***Genetic subcluster:*** Rus_Core.

**DR25. Burial 5A**

***Grave type:*** A ground burial oriented southwest–northeast. The grave pit was not identified, and the dimensions of the burial remain unknown.

***Dating:*** The second half of the 12th-14th centuries CE.

***Skeletal information:*** The skeleton was well-preserved and lying in an extended supine position. The remains belonged to a man over 50 years old.

***Grave goods:*** No grave goods were found.

***Genetic subcluster:*** Rus_SWest.

**DR26. Burial 5B**

***Grave type:*** A ground burial oriented west–east. The grave pit was not identified, and the dimensions of the burial are unknown

***Dating:*** The second half of the 12th-14th centuries CE.

***Skeletal information:*** The skeleton was in a well-preserved state, lying in an extended supine position, with the skull facing upwards. The remains belonged to a man over 60 years old.

***Grave goods:*** No grave goods were found.

***Genetic subcluster:*** Rus_Core.

**DR27. Burial 6**

***Grave type:*** A ground burial oriented along a west-east axis. The grave pit could not be identified within the stratigraphic layer, and the dimensions of the burial remain undetermined.

***Dating:*** *The s*econd half of the 12th-14th centuries CE.

***Skeletal information:*** The skeleton was in a good state of preservation and was lying in an extended supine position. The remains belonged to a man aged 40-49 years.

***Grave goods:*** No grave goods were found.

***Genetic subcluster:*** Rus_Core.

**DR28. Burial 7**

***Grave type:*** A ground burial oriented along a west-east axis with a slight deviation to the north. The skull was shifted 15 cm to the north. The grave pit could not be identified, and the dimensions of the burial remain undetermined.

***Dating:*** The second half of the 12th-14th centuries CE.

***Skeletal information:*** The skeleton was in a good state of preservation and was lying in an extended supine position. The remains belonged to a man aged 30-39 years.

***Grave goods:*** No grave goods were found.

***Genetic subcluster:*** Rus_Core.

**DR29. Burial 8**

***Grave type:*** A ground burial oriented along a northwest-southeast axis.

***Dating:*** The second half of the 12th-14th centuries CE.

***Skeletal information:*** The skeleton was in a good state of preservation and was lying in an extended supine position. The remains belonged to a man aged over 40 years.

***Grave goods:*** No grave goods were found.

***Additional information:*** Partially overlain by Burial 7.

***Genetic subcluster: -***

**DR31. Burial 10**

***Grave type:*** A ground burial oriented along a west-east axis. The grave pit could not be identified, and the dimensions of the burial remain undetermined.

***Dating:*** The second half of the 12th-14th centuries CE.

***Skeletal information:*** The skeleton was in a good state of preservation and was lying in an extended supine position. The remains belonged to a woman aged over 60 years old.

***Grave goods:*** 2 round appliqué plaques (located in the skull area near the left clavicle) and an ellipsoidal button (positioned between the right humerus and the ribs).

***Genetic subcluster:*** Rus_SWest.

**DR32. Burial 11**

***Grave type:*** A ground burial oriented along a west-east axis. The grave pit could not be identified, and the dimensions of the burial remain undetermined.

***Dating:*** The second half of the 12th-14th centuries CE.

***Skeletal information:*** The skeleton was in a good state of preservation and was found with the lower jaw separated from the skull and positioned on the right side.The remains belonged to a man aged over 50 years old.

***Grave goods:*** No grave goods were present.

***Genetic subcluster:*** Rus_Core.

**DR33. Burial 12**

***Grave type:*** A ground burial oriented along a west-east axis, with slight deviations to the north. The grave pit could not be identified, and the dimensions of the burial remain undetermined.

***Dating:*** The second half of the 12th-14th centuries CE.

***Skeletal information:*** The skeleton was poorly preserved, represented only by a fragmented skull and isolated bones. The remains belonged to a woman aged 35-45 years.

***Grave goods:*** A fragment of textile with gold thread was found on the chest.

***Genetic subcluster:*** Rus_Core.

##### S1.2.4.2. The Kurilovka burial site (Kursk)

**Geographic Information:** The Kurilovka 2 site is located in the Sudzha district of the Kursk region, in the northern part of the Dnieper forest-steppe on the left bank, near the confluence of the Sudzha and Psel rivers. Several archaeological layers associated with Proto-Slavic and Early Slavic cultures have been identified at this site. The most recent of these layers corresponds to the Volyntsevo culture. A detailed description of the Kurilovka 2 site has been provided

**Summary of sampled materials:** Dwelling 13, from excavation 3 of the Kurilovka 2 settlement, where human remains were discovered, belongs to the "Sakhnovo-Volyntsevo" settlement of the Volyntsevo culture, dating from the late 7th to the first half or middle of the 8th centuries CE. The anthropological remains found in the dwelling consisted of fragments of skull bones.

In part, we have previously reported the archeogenetic data for this individual^73^. The genetic pool of the Proto-Slavic and early Slavic populations remains largely unexplored, owing to the prevalence of cremation burial practices in Slavic cultures prior to the 10th century CE. Consequently, any skeletal remains dating from early Slavic sites are of significant scientific value. The genetic data of individual DR10, belonging to the Volyntsevo culture (7th-8th centuries CE), is included in this study to investigate the genetic continuity and ancestry of the population in the territory of ancient Rus'. Representatives of the Volyntsevo culture formed the autochthonous population of the Dnieper Forest-Steppe, which preceded the early eastern Slavic groups that contributed to the formation of ancient Rus. Archaeologically, the Volyntsevo culture exhibits close cultural connections with the Prague-Korchak and Luka-Radkovets traditions, which have been widely regarded as foundational to the material culture of the early East Slavs. The DR10 individual offers a representative sample for examining the ancestral component of the gene pool in the Middle Dnieper region, which subsequently formed part of Rus'.

**DR10. Burial from the dwelling 13**

***Skeletal information:*** Only the skull remains of the skeleton belonged to a child aged 3-4 years.

***Genetic subcluster: -***

##### S1.2.4.3. The Moiseevskoe burial site (Kursk)

**Geographic Information:** The Moiseevskoe gorodishche (ancient settlement) is located on one of the spurs of the high right bank of the Svapa river and on the left bank of the small, partially dried-up Iput River, in the Dmitrovsky district of the Kursk region. The settlement has a narrow and elongated shape, stretching from north to south, and consists of three distinct platforms.

**Excavation History:** Excavations at the settlement were conducted in 1955 by E.A. Alikhova.

**Summary of sampled materials:** It was determined that the earliest horizon of the cultural layer dates back to the Early Scythian period, while subsequent occupation of the site occurred only during the Rus’ period. Within the third platform, which is protected by ramparts on all sides, the detinets (inner fortified area) was situated. In this part of the settlement, burials were discovered in ground burial pits, at depths ranging from 0.3 to 0.7 m. The bodies were laid in an extended supine position with their heads oriented to the west–southwest. Frequently, skeletal remains were displaced and scattered. The basis for the site's dating, as noted by E.A. Alikhova, is the presence of a temporal ring with a spiral-shaped end, which has close parallels to items found at the Gochevsky burial ground in the Kursk region. The settlement is dated to the 11th-12th centuries CE^74^.

Genetic analysis was performed on bone samples obtained from two skulls that are kept in the Museum of Anthropology, Moscow Lomonosov State University.

**Excavation V. Paired Burial 2**

This is a double burial in a grave pit with a depth of 0.3 m. The buried individuals were laid in an extended supine position, with their heads oriented to the west-southwest. The bones of both individuals were shifted and intermixed, and the skulls were turned to the side.

**AB28. Individual 1**

***Dating:*** The 11th-12th centuries CE.

***Skeletal information:*** The remains belonged to a woman.

***Grave goods:*** An iron knife.

***Museum ID:*** 10048В.

***Additional information:*** The expedition of the IIMC of the USSR Academy of Sciences and the Kursk Local Museum, excavations by E.A. Alikhova in 1955.

***Genetic subcluster: -***

**AB29. Individual 2**

***Dating:*** The 11th-12th centuries.

***Skeletal information:*** The remains belonged to a man.

***Grave goods:*** An iron knife.

***Museum ID:*** 10048A.

***Additional information:*** The expedition of the IIMC of the USSR Academy of Sciences and the Kursk Local Museum, excavations by E.A. Alikhova in 1955.

***Genetic subcluster:*** Rus_SWest.

##### S1.2.4.4. The Gushchino burial site (Chernigov)

**Geographic Information:** The burial barrow complex is located near the modern village of Gushchino in the Chernigovsky district of Chernigov region, on the high bank of the Belous river (a tributary of the Desna river), 5 km from the city of Chernigov.

**Excavation History:** Excavations were conducted by D.Ya. Samokvasov in 1872.

**Summary of sampled materials:** Burials were found in hemispherical barrows, within grave pits that ranged from 0.71 to 2.13 m deep. Coffin remains and iron nails were often recorded. The skeletons were laid in an extended position on their backs, oriented with their heads to the west. The skulls faced straight forward, and the hands were resting on the abdomen. Grave goods: the burial inventory includes iron knives, earrings, beads, bronze pendants, rings, buckles, and other pendants. The burial practices and artefacts indicate ancient Rus’ cultural context with the features of military elite necropolis. Dating: 10th-11th centuries CE^75,76^.

Genetic analysis was carried out on a bone sample obtained from a skull that is kept in the Museum of Anthropology, Lomonosov Moscow State University.

**AB14**

***Skeletal information:*** Skeletal remains belonged to a man.

***Museum ID:*** 2083.

***Genetic subcluster:*** Rus_Core.

##### S1.2.4.5. The Konotop burial site (Sumy regin)

**Geographic Information:** The ancient burial ground of Konotop, currently located in the Sumy region, was situated within the confines of a fortress dating from the Middle Ages and the early Modern period. This site is situated on the marshy left bank of the Ezuch river.

**Excavation History:** The excavations were carried out by T.V. Kibalchich in 1878 on the territory of the now non-existence Ponomarev estate. The burial was found on the territory of the settlement.

**Summary of sampled materials:** The documentation of the excavations is incomplete, but it is known that the burial was found in a rectangular grave pit, with the remains laid on their backs and oriented along a west-east axis^77^.

Genetic analysis was carried out on a bone sample obtained from a skull that is kept in the Museum of Anthropology, Lomonosov Moscow State University.

**AB58**

***Dating:*** 11th-13th century CE.

***Skeletal information:*** Skeletal remains belonged to a man. Features: enamel hypoplasia.

***Grave goods:*** No grave goods were found.

***Museum ID:*** 2074.

***Genetic subcluster:*** Rus_SWest.

##### S1.2.4.6. Knyazhya gora burial site (Cherkassy)

**Geographic Information:** The ground necropolis (burial site) is located at the Knyazhya gora settlement, situated on a long and narrow ridge along the right bank of the Dnieper river, at the mouth of the Ros river (which historically flowed into the Dnieper), 7.5 km south of the town of Kanev (Cherkassy region).

**Excavation History:** During excavations conducted in 1891 by N.F. Belyashevsky, five burials were discovered on the outskirts of the site.

**Summary of sampled materials:** The Knyazhya gora settlement is identified with the *Primary Chronicle* city of Roden, where Yaropolk Svyatoslavich took refuge from his brother Vladimir in 980 CE. Five burials were found in ground grave pits at a depth of 1-1.35 m from the modern surface. The deceased were positioned in an extended supine posture, with their heads facing west, and with slight deviations to the south or north. Their hands were folded on the chest. Grave goods: no burial goods were found. Dating: 10th-12th centuries CE^78^.

Genetic analysis was carried out on three skulls that are kept in the Museum of Anthropology, Lomonosov Moscow State University.

**AB22**

***Skeletal information****:* Skeleton belonged to a man. Features and Pathologies: A cut trauma to the left parietal bone, showing signs of healing. The cut was 53 mm long.

***Museum ID:*** 8112.

***Genetic subcluster****:* Rus_Core.

**AB23**

***Skeletal information****:* Skeleton belonged to a man.

***Museum ID:*** 7335.

***Genetic subcluster****:* Outlayer.

**AB24**

***Skeletal information****:* Skeleton belonged to a woman.

***Museum ID:*** 8863.

***Genetic subcluster****:* Outlayer.

##### S1.2.4.7. Burial site in Chernigov

**Geographic Information:** Chernigov was one of the largest cities of ancient Rus' and the capital of the Chernigov principality, the first mention of Chernigov dates back to 907 CE. The city is located in the Dnieper Lowland, on the right bank of the Desna river, at its confluence with the Strizhen river. The burial ground is located near the Church of Boris and Gleb.

**Excavation History:** The bone remains were transferred for storage by the N.D. Dolgorukov estate in 1898, south of the Church of Boris and Gleb.

**Summary of sampled materials:** The burial ground was located at the Cathedral Square of the ancient Chernigov's citadel, on a high terrace of the right bank of the Desna river. The skulls submitted for analysis were found in a mass grave with disordered bone remains. No grave goods were found. Dating: probably 11th-13th centuries CE^77^.

Genetic analysis was carried out on two skulls that are kept in the Museum of Anthropology, Lomonosov Moscow State University.

**AB56**

***Skeletal information:*** The burial belonged to a man. A chop wound was found on the left parietal bone, 30 mm in length, and a trauma to the frontal bone on the left side, 25 mm in length, with no signs of healing.

***Museum ID:*** 2067.

***Genetic subcluster:*** outlayer.

**AB57**

***Skeletal information:*** The burial belonged to a woman.

***Museum ID:*** 2062.

***Genetic subcluster: -***

##### S1.2.4.8. Burial site in Lubech

**Geographic Information:** Lubech, one of the oldest urban centres in the Southern Rus’, was situated on the left bank of the Dnieper river.

**Excavation History:** The city was excavated from 1957 to 1960 by the Chernigov expedition of the Institute of Archaeology of the USSR Academy of Sciences, led by B.A. Rybakov. The excavations uncovered most of the citadel area, including burials.

**Summary of sampled materials:** The site, consisting of two fortified areas with a total area of approximately 10 hectares, is situated in the Zamkovaya gora locality in present-day Lubech. Lubech is first mentioned in the *Primary Chronicle* in 882 CE in the account of Oleg's campaign to Kiev. Initially, it was under the rule of Kiev princes, but came under the control of the Chernigov princes (the Olgovichi and Davydovichi) in the 11th century CE, remaining their dominion until the 13th century CE. In 1097, Lubech became the site of the Lubech Congress of Princes, where a new order of inheritance among the Rurikid princes was established, giving each prince the right to inherit only their father's lands. In the 14th-16th centuries CE, Lubech was part of the Grand Duchy of Lithuania, retaining its significance as a major princely fortress in the Kiev Voivodeship. By the mid 17th century CE, Lubech had become the centre of the Cossack Chernigov regiment. The burial grounds, discovered in the southeastern part of the settlement, contained graves from different periods of the settlement's history - 11th-12th and 17th centuries CE. The 11th-12th centuries CE burials in excavation 12 were found in rectangular grave pits, 0.5-1.15 m deep, with the heads oriented westward, occasionally with slight deviations to the south or north. The skeletons were laid flat on their backs with arms crossed over the chest or abdomen. No grave goods were found. According to B.A. Rybakov, the dating of the burials was based on their partial destruction by structures dated to the 12th-13th centuries CE. The 17th century CE burials in excavations 7 and 11 were found in rectangular grave pits, 0.3-0.5 m deep, often within wooden coffins. The deceased were laid on their backs with hands folded on the chest. A distinctive feature of these burials was the presence of stones and bricks under the head of the deceased. No grave goods were found^79–81^.

Genetic analysis was carried out on seven skulls that are kept in the Museum of Anthropology, Lomonosov Moscow State University.

**AB141. Excavation 12. Burial 38**

***Grave type:*** The burial was found in a rectangular grave pit, oriented southwest-northeast, with a depth of 0.8 m from the modern surface.

***Dating:*** 11th-12th centuries CE.

***Skeletal information:*** The skeleton was lying flat on the back. The right arm was bent at the elbow, resting on the abdomen, while the left arm’s radius bone was not preserved. The burial belonged to a man.

***Museum ID:*** 10914.

***Genetic subcluster:*** Rus_SWest.

**AB134. Excavation 11. Burial 4**

***Grave type:*** The burial was found in a rectangular wooden coffin (thin strips of rotted wood forming a rectangular shape in the grave) within a rectangular grave pit, dug into the natural soil to a depth of 0.3-0.5 m.

***Dating:*** 17th century CE.

***Skeletal information:*** The skeleton was lying on its back with arms folded on the chest. The burial belonged to an adult man.

***Museum ID:*** КО 1-5.

***Additional information:*** Bricks and stones were found under the head and feet.

***Genetic subcluster:*** Rus_SWest.

**AB135. Excavation 7. Burial 4**

***Grave type:*** The burial was found in a rectangular grave pit.

***Dating:*** 17th century CE.

***Skeletal information:*** The skeleton was lying flat on the back, with the head oriented southwest. The arms were folded on the abdomen. The burial belonged to a woman.

***Museum ID:*** KO 1-6.

***Genetic subcluster:*** Rus_SWest.

**AB136. Excavation 11. Burial 6**

***Grave type:*** this burial was found in a rectangular wooden coffin, within a rectangular grave pit dug into the natural soil, 0.3-0.5 m deep.

***Dating:*** 17th century CE.

***Skeletal information:*** The skeleton was lying on their back with arms folded on the chest.The burial belonged to a man.

***Museum ID:*** KO 1-8.

***Additional information:*** Bricks and stones were found under the head and feet.

***Genetic subcluster:*** Rus_SWest.

**AB137. Excavation 11. Adjacent to Burial 6**

***Gave type:*** The burial was found in a rectangular wooden coffin (thin strips of rotted wood, forming a rectangular shape in the grave) within a rectangular grave pit, dug into the natural soil to a depth of 0.3-0.5 m.

***Dating:*** 17th century CE.

***Skeletal information:*** The skeleton was lying on its back with arms folded on the chest. The burial belonged to a man.

***Museum ID:*** KO 1-9.

***Additional information:*** Bricks and stones were found under the head and feet.

***Genetic subcluster: -***

**AB138. Excavation 11. Burial 29**

***Grave type:*** The burial was found in a rectangular wooden coffin (thin strips of rotted wood, forming a rectangular shape in the grave) within a rectangular grave pit, dug into the natural soil to a depth of 0.3-0.5 m.

***Dating:*** 17th century CE.

***Skeletal information:*** The skeleton was lying on its back with arms folded on the chest. The burial belonged to a man.

***Museum ID:*** 10909.

***Additional information:*** Bricks and stones were found under the head and feet.

***Genetic subcluster: -***

**AB139. Excavation 11. Burial 31**

***Grave type:*** The burial was found in a rectangular wooden coffin (thin strips of rotted wood, forming a rectangular shape in the grave) within a rectangular grave pit, dug into the natural soil to a depth of 0.3-0.5 m.

***Dating:*** 17th century CE.

***Skeletal information:*** The skeleton was lying on the back with arms folded on the chest. The burial belonged to an adolescent.

***Museum ID:*** 10910.

***Genetic subcluster:*** Rus_SWest.

##### S1.2.4.9. Burial sites in Kiev

Kiev, one of the oldest and greatest cities in the region historically referred to as Rus’, functioned as the capital of the first East Slavic political entity. Kiev is situated along the banks of the Dnieper river, downstream from the confluence with its left tributary, the Desna. Two cemeteries within the Kiev citadel on the right bank of the Dnieper river were examined. One is the city's ancient cemetery, while the other is a burial site adjacent to the Desyatinnaya Сhurch.

***Burial site of the ancient cemetery***

**Geographic Information:** The burial was located within the boundaries of the city's citadel, near the present-day Trubetskoy estate, on the right bank of the Dnieper river.

**Excavation History:** According to reports by V.B. Antonovich and T.V. Kibalchich Kibalchich who excavated the Kiev ancient cemetery in 1870, both seated burials and cremations were also found in the necropolis.

**Summary of sampled materials:** All the burials were conducted in grave pits measuring 1.5-2 m in length and 2-3.8 m in depth from the surface. These pits were situated in levelled hemispherical barrows, surrounded by ring-shaped trenches. The skeletons were laid extended on their backs, with heads oriented from north to west. Grave goods included "silver earrings of the Kiev type and pottery associated with the grand princely period of Kiev." The burial ground is dated to the 9th-10th centuries CE^77,82,83^.

Genetic analysis was carried out on a bone sample obtained from a skull that is kept in the Museum of Anthropology, Lomonosov Moscow State University.

**AB54**

***Skeletal information:*** The skeleton belonged to a man.

***Museum ID:*** 2115.

***Genetic subcluster:*** -

***Burial site at the Desyatinnaya Church***

**Geographic Information:** The burial ground is located close to the Desyatinnaya Church (Church of the Tithes) within the citadel of Kiev on the right bank of the Dnieper river.

**Excavation History:** Archaeological excavations of the remnants of the church and surrounding burial grounds were conducted by V.B. Antonovich and T.V. Kibalchich who excavated the Kiev ancient cemetery in 1878.

**Summary of sampled materials:** It is the oldest stone Christian temple in the ancient Rus', founded by Vladimir Sviatoslavich in 989 CE on the site of an earlier necropolis. All the burials were discovered in rectangular grave pits, 0.5 m deep, near the foundation of the Desyatinnaya Church, in wooden coffins fastened with iron clamps. The skeletons were laid out extended on their backs, with their heads oriented westward, and less frequently, to the northwest. The stratigraphic position of the graves, located below the foundation elements and the wooden substructures of the church, allows the dating of these burials to a period preceding^83^.

Genetic analysis was carried out on a sample obtained from a skull that is kept in the Museum of Anthropology, Lomonosov Moscow State University.

**AB55. Burial 7**

***Skeletal information:*** The skeleton belonged to a man.

***Museum ID:*** 9418.

***Genetic subcluster:*** -

### S1.3. Burial grounds in the areas adjacent to medieval Rus’

#### S1.3.1. Burial site of the Latgalian culture

##### S1.3.1.1. The Lucinsky burial site (Ludza)

**Geographic Information:** The burial ground is located on the southern shore of the Great Ludza lake near Ludza, Eastern Latvia's oldest town.

**Excavation History:** Excavations of the burial ground were conducted by G. Romanov in 1890-1891 and by V.I. Sizov in 1891.

**Summary of sampled materials:** A total of 338 burials were uncovered. The burials were made in earthen grave pits. Wooden planks were often found at the bottom of the graves, serving as bedding. The skeletons, covered with birch bark or boards, lay extended on their backs. Occasionally, remnants of wooden coffins, without traces of iron nails, were found along the sides of the skeletons. The skulls were typically turned to the right. Wooden headrests were found under the skulls in the graves of women and children. Grave goods: horseshoe-shaped fibulae, bronze trapezoidal pendants, crescent-shaped pendants with extended "horns" and loops for attaching additional decorations, chest chains, headbands with rectangular casings and spiral beads, twisted torcs, bronze bracelets with flaring ends, belt tips, fittings, and buckles with casings. The site is attributed to Latgalian culture^84^.

Genetic analysis was carried out on a bone sample obtained from a single skull that is kept in the Museum of Anthropology, Lomonosov Moscow State University.

**AB27**

***Skeletal information:*** The burial belonged to a man.

***Museum ID:*** 2612.

***Genetic subcluster:*** -

#### S1.3.2. Burial sites of Muroma culture

##### S1.3.2.1. The Podbolotyevsky burial site (Murom)

**Geographic Information:** This burial ground is located on the southwestern outskirts of the village of Verbovsky in the Muromsky district of the Vladimir region. It sits on the main terrace of the left bank of the Ilevna river (a left tributary of the Oka river) at the head of the ravine of the Babiy Bereznyak area. The terrace is bordered by the Ilevna river to the west and southwest, and by a damned stream to the south, which flows into the river.

**Excavation History:** The burial ground was discovered in 1910 during the construction of the Murom-Melenki highway. Information about the site was reported to P.S. Uvarova, who ensured the preservation of the finds and invited V.A. Gorodtsov, who led the first rescue excavations. V.V. Golmsten and D.N. von Eding was also involved. A total of 260 burials

were investigated in the early 20th century. Subsequent excavations were carried out by the Volga Archaeological expedition of the IA RAS from 2012 to 2014 due to the construction of a bypass road around town Murom. Over three years, 200 burials were explored over an area of 2.531 square metres. The Podbolotyevsky burial ground is one of the most well-known burial grounds of the chronicled Fenno-Ugrian people of Murom and for many years was considered a reference burial ground for studying the Murom people.

**Summary of sampled materials:** Excavations covered an area of 2.154 square metre, with trenches 22-25 m long and 2 m wide. Most burials were by inhumation, while 14% followed the cremation rite. The funeral rites of the female attire suggest that the burial ground belonged to the chronicler Fenno-Ugrian people of the Muroma. The women's burials of the Podbolotyevsky burial ground were distinguished by a complex set of metal costume ornaments, which included ethnocultural markers: spiral head harnesses, bracelet-shaped temporal rings of the so-called "Murom type", temporal moonlight pendants, frontal corollas. Women's jewelry included noisy pendants, hryvnia, necklaces made of noisy pendants or glass beads, beads, permeated and spiraled. A striking detail of the Murom women's costume was a side belt with delicate rustling buckles. The burial ground is dated from the late 7th to the 11th centuries CE. The dating is supported by numerous metal costume accessories and grave goods, as well as radiocarbon dates from organic samples^85–87^.

For the genetic study, only samples dated from the 9th-11th centuries CE were taken

**DR51. Excavation 4. Burial 105**

***Grave type:*** a grave pit (2x0.7 m) at a depth of 0.2 m from the bedrock level.

***Dating:*** The late 10th - the early 11th centuries CE.

***Skeletal information:*** The skeleton was lying on its back in an extended position, with the hands near the abdomen and the head facing north. The burial belonged to a woman around 20 years old.

***Grave goods:*** 4 bracelet-shaped temporal rings with a shield and hook lock, a Glazov-type necklace, a cluster of glass beads, small beads, and remains of tin beads, traces of 6 coin-like pendants made of tin alloy, a round fibula belonging to the circle of Scandinavian antiquities, an openwork lunula pendant with jingling pendants on a chain, 2 narrow massive bracelets and 1 plate bracelet, traces of 5 tin alloy rings, an iron knife with a wooden handle, an iron awl, remains of tin alloy, 2 clay vessels, a clay spindle whorl.

***Genetic subcluster:*** Muroma.

**DR52. Excavation 4. Burial 202**

***Grave type:*** a grave pit (2 x 0.9 m) at a depth of 0.1 m from the bedrock level.

***Dating:*** 9th century CE.

***Skeletal information:*** The burial site has been significantly disturbed, leaving only the skull and some fragments of the rib cage in place. The skeleton's original position is reconstructed as extended on its back, with the head facing north. The burial belonged to a woman aged approximately 35-45 years.

***Grave goods:*** 2 head braids, 69 plate trapezoidal pendants, a fragment of a narrow forehead band, fragments of plate fittings from a forehead band, 6 bead-shaped pendants with claw-shaped attachments, an openwork rectangular pendant with bottle-shaped attachments, a horseshoe-shaped spiral-ended fibula, 3 spiral rings, 3 wide-centered "mustached" rings and one silver ring with a shield decorated with a pitted design, a fitting from a yoke-shaped pendant, fragments of bracelet-shaped temporal rings, a crescent-shaped pendant, a fragment of a hair decoration with a bottle-shaped attachment, plate fittings from a side belt, 12 claw-shaped pendants, 6 bottle-shaped pendants.

***Genetic subcluster: -***

#### S1.3.3. Burial site of ancient Mordva

##### S1.3.3.1. The Pogiblovo burial site (Nizhny Novgorod)

**Geographic Information:** The Pogiblovo (Malinovka) burial ground is located 1 km northwest of the Arzamas-Temnikov postal route, on the gentle slope of an elevated plateau near the Irzha river, in the vicinity of Pogiblovo village, Ardatovsky district, Nizhny Novgorod region. The natural boundaries of the burial ground include the steep left bank of the Irzha river to the southeast and the steep hillside to the southwest. The total area of the burial ground is approximately 4.800 square meters.

**Excavation History:** The burial ground was first surveyed by the Anthropological Complex expedition led by A.E. Alikhova and M.V. Voevodsky in 1926. In 1927, excavations were carried out by E.I. Goryunova and K.I. Churazova, covering a total area of 175 square meters. **Summary of sampled materials:** A total of 18 burials were discovered, 9 of which had been disturbed. Bone preservation was poor in most cases, with some remains entirely absent. The burials were located in oval or circular pits, with traces of wooden bedding, bast, or birch bark. Male burials were extended on the back, with arms placed alongside the body. Female burials were found on the left side in a flexed position, with legs bent and hands brought to the face. Female skeletons were generally located at depths of 60-70 cm, male skeletons at approximately 1.4 m, and children's remains at depths of 40-50 cm. The orientation of the burials was inconsistent. The burial ground is associated with the Mordva, as indicated by E.N. Goryunova's work. O.V. Zelentsova noted that a male burial (№ 9) in trench IV, identified by the presence of a burial girdle, was dated to the 12th-13th centuries CE. However, the cultural affiliation of the burial ground remains tied to the Mordva. The overall dating of the burial ground is from the 12th-13th centuries CE^88^.

Genetic analysis was carried out on a bone sample obtained from a single skull that is kept in the Museum of Anthropology, Lomonosov Moscow State University.

**AB38.**

***Grave type:*** In excavation IV, a single burial (№ 10) was found in the northwestern part of the burial pit, at a depth of 0.6 m.

***Dating:*** 12th-13th centuries CE.

***Skeletal information:*** The individual was lying extended on the back, with the head oriented to the northwest and the arms slightly bent at the elbows.

***Grave goods:*** No grave goods were found.

***Museum ID:*** 8474.

***Genetic subcluster:*** Mordva.

##### S1.3.3.2. The Muranka burial site (Samara)

**Geographic Information:** The archaeological complex near the village of Muranka is located in the middle reaches of the Usa river, in the Shigonsky district of the Samara region, 3 km south of the village. The burial ground and settlement are situated on opposite banks of the river: the burial ground on the right bank and the settlement on the left.

**Excavation History:** The complex was discovered in the late 19th century, and excavations were conducted by V.N. Polivanov in 1891, 1892, 1893, and 1900. A total of 445 burials were uncovered by him. In 1950, excavations were resumed by the Kuibyshev Archaeological expedition, led by A.E. Alikhova. During the excavations of areas I and III, 38 medieval Mordva burials were uncovered. In area VI, a Muslim cemetery was revealed, separate from the Mordva burials. According to D. Stashenkov, who studied the territory of the burial ground and settlement in 2010, the Muransky settlement reached its peak between 1320 and 1350 CE, with its decline linked to the period of the "Great Troubles" in the Golden Horde during the 1360s CE.

**Summary of sampled materials:** All Mordva burials were conducted in rectangular pits. Some burials were found in wooden coffins or log structures, constructed without the use of iron nails. The skeletal remains in the male burials were laid out in a supine position, with the head oriented to the south or southeast. The female skeletons were found lying on their right side in a slightly flexed position, with bent legs and arms, and the hands near the face. In the upper layers of the burial fill, fragments of broken ceramic vessels and individual animal bones were found. In one case, the shell of a bird’s egg was discovered. The funerary inventory predominantly originates from female burials. Items include silver and bronze syulgamas (a type of ornament), small wire rings with open ends, earrings from the Golden Horde period shaped like question marks, glass and stone beads, amuletic pendants made from animal bones and teeth, nakosniki (braid ornaments), silver rings with large shields, as well as iron axes, knives, and fire strikers. The site is dated to the 14th century CE^89–91^.

Genetic analysis was carried out on three skulls that are kept in the Museum of Anthropology, Lomonosov Moscow State University.

**AB30.**

***Skeletal information:*** The sex and age of the skeletal remains cannot be determined.

***Museum ID:*** 1719.

***Additional information:*** The excavation by V.N. Polivanov in 1950.

***Genetic subcluster:*** Mordva.

**AB31**

***Skeletal information:*** The skeletal remains belonged to a woman.

***Museum ID:*** 1731.

***Additional information:*** The excavation by V.N. Polivanov in 1950.

***Genetic subcluster: -***

**AB32**

***Skeletal information:*** The skeletal remains belonged to a woman.

***Museum ID:*** 1724.

***Additional information:*** The excavation by V.N. Polivanov in 1950.

***Genetic subcluster: -***

##### S1.3.3.3. The Korino burial site (Nizhny Novgorod)

**Geographic Information:** The burial ground is located near the large Erzya village of Korino, 4 km west of the Shatki railway station and 1 km northwest of the village of Korino. **Excavation History:** The site was first discovered and investigated in 1913 by local landowner Bogodurov, who excavated two barrows (№ 12 and № 14). In 1926, the burial ground was surveyed by a reconnaissance unit of the Anthropological Complex expedition, led by E.A. Alikhova and M.V. Voeyvodsky, under the supervision of E.I. Goryunova. The burial complex consists of a group of barrows (barrows) and inhumation burials. The barrow group was located in the northern part of the burial ground, on a small elevated area on a promontory facing a seasonal stream known as "Vonyachka", situated 50 m from the road connecting the Villages of Korino and Khirino. In total, 15 barrows are known. In 1926, ten of the barrows were excavated. The barrow burials of the site are dated by silver coins bearing the names of Moscow tsars from the 16th and 17th centuries CE. In the southern part of the burial ground, on the flatter side, are earlier inhumation burials dating to the 12th-13th centuries CE. The burial ground belongs to the ancient Mordva.

**Summary of sampled materials:** The barrows had a "round-domed" shape, with diameters ranging from 5 to 6 m and heights between 0.5 and 1.5 m. All burials followed the rite of inhumation at the level of the ancient ground surface, with the bodies laid on their backs, arms either extended alongside the body or placed on the pelvic bones. Both individual and paired burials have been recorded. The orientation of the burials is to the northwest. Burials in coffins, on bedding, or in birch bark (lub) are also known. Along with the deceased, animal bones (cattle, calves, small livestock, and less frequently wild animals) were found. The date of these barrows is from the 16th-17th centuries CE^92^.

Genetic analysis was carried out on a bone sample obtained from a single skull that is kept in the Museum of Anthropology, Lomonosov Moscow State University.

**AB25. Burial 10a.**

***Grave type:*** a double barrow. Under the northern barrow, a paired burial was found. However, only the disordered tibiae and femora of the second burial remain. E.I. Goryunova suggested that these bones were accidentally disturbed during earth removal for the barrow construction and subsequently reburied.

***Skeletal information:*** The primary burial (AB25) consists of the skeletal remains of a man. The body was laid out on its back at the base of the moon, with the head to the northwest. The skull was slightly tilted back and to the left, with the legs extended. The body was covered with a layer of birch bark.

***Museum ID:*** 34.

***Additional information:*** The excavation by E.I. Goryunova in 1926.

***Genetic subcluster:*** *-*

#### S1.3.4. Burial site of Volga Bulgars

##### S1.3.4.1. The Syut-Siriami burial site (Chuvashia)

**Geographic Information:** The burial ground is situated on the plateau of the Syut-Siriami fortified settlement, measuring 525 by 525 m. The site is bordered to the west by the Tayabinka river and to the southwest by the Syut-Siriami tributary, located in the Yalchinsky district of the Chuvash Republic (Chuvashia), near the village of Bolshaya Tayaba.

**Summary of sampled materials:** The settlement was fortified with two defensive ramparts.The graves were positioned in rectangular pits oriented from west to east. The depth of these burial pits varied from 28 to 80 cm. The skeletons were laid out on their backs, with a slight turn to the right side, their heads facing west, and their faces turned southward, towards Mecca. The right arm was extended along the body, while the left arm was slightly bent at the elbow and rested on the waist or pelvic bones. No burial goods were found. Cultural affiliation: Volga Bulgars. Dating: 10th-13th centuries CE^93,94^.

Genetic analysis was carried out on three skulls that are kept in the Museum of Anthropology, Lomonosov Moscow State University.

**AB3**

***Skeletal information:*** Skeletal remains belonged to a woman.

***Museum ID:*** 10593.

***Additional information:*** The expedition of the IIMC, the excavation by A.P. Smirnov in 1948.

***Genetic subcluster: -***

**AB43**

***Skeletal information:*** Skeletal remains belonged to a woman.

***Museum ID:*** 10590.

***Additional information:*** The expedition of the IIMC, the excavation by A.P. Smirnov in 1948.

***Genetic subcluster: -***

**AB44**

***Skeletal information:*** Skeletal remains belonged to a woman.

***Museum ID:*** 10591.

***Additional information:*** The expedition of the IIMC, the excavation by A.P. Smirnov in 1948.

***Genetic subcluster:*** Volga Bulgars.

### S1.4. Radiocarbon dating

Radiocarbon dating was performed on 12 bone and tooth specimens from four archaeological sites: three burial grounds in Northern Rus’ located in the Beloozero region (Minino, Nefedyevo, Nikolskoe) and the Gnezdilovo burial ground in the Central Rus’ Volga-Oka region (Supplementary table 5). The majority of the obtained dates fall within the investigated historical ancient Rus’ period. Nonetheless, the radiocarbon dates were consistently older than the age estimates derived from archaeological methods. The systematic bias towards older dates is likely due to the freshwater reservoir effect, which has not yet been fully understood. Potential impacts from Miyake events—a rapid increase in carbon-14 (^14^C) identified in 993-94 CE^95,96^ —may also contribute to dating uncertainties, a phenomenon observed at other medieval sites in the Rus’ region^97^.

Despite the dating challenges mentioned above, we consider dating based on archaeological materials in our study. It should be noted that all of the burials for which radiocarbon dating was obtained contained artefacts, the chronology of which is well developed based on dendrochronology and closed complexes with coins.

In addition, radiocarbon analysis of human remains from a mass grave in Yaroslavl was performed previously in the laboratory CAIS (The Center for Applied Isotope Studies, University of Georgia. USA) and confirmed the archaeological age of the Yaroslavl burials that coincides with the period of the Mongol invasion in 1238^98^.

### S2. Supplementary methods and results

#### S2.1. Datasets

We merged our ancient DNA (aDNA) data with previously published datasets of 1085 ancient and 3049 present-day individuals reported by the Reich Lab in the Allen Ancient DNA Resource v54.1 (Supplementary Table 6).

#### S2.2. PCA analyses results of Rus’ individuals and previously published ancient samples

When we projected our ancient Rus’ samples on the first two principal components of the present-day Eurasian populations, we found our samples clustered predominantly with present-day European groups. Most of the ancient Rus’ individuals are clustered into two distant genetic clusters, that we termed Ancient North Rus and Ancient Major Rus (Fig. 2, Supplementary Fig. 29), further subdivided into several genetic groups (subclusters) named Rus_Core, Rus_Baltic, Rus_SWest, Rus_North_1, and Rus_North_2 (Fig. 3A). Remarkably, in comparison to other ancient Rus' samples, the Rus_Baltic genetic cluster has a higher level of the European HG ancestry, whereas the Rus_SWest subcluster is characterised by a higher ancestry component of European Neolithic farmers linked to ancient Anatolian populations (Fig. 4).

To better assess the genetic makeup of Rus’ population in the context of other ancient groups, we projected our Rus’ samples, along with previously published ancient individuals dated from the Iron Age (IA) to the late Medieval Ages (MA) (Supplementary Fig. 30 [Supplementary_Fig_30. PCA_MA_IA.pdf](https://drive.google.com/file/d/1dSzyEKjEb17sjjxBTpr6_fQIHAocEZwJ/view?usp=drive_link)) from the East European Plain and adjacent regions of Eurasia, onto present-day West Eurasian populations. We used numerous ancient groups from the early IA to the late Medieval period for this analysis to get a comprehensive overview (Supplementary table 6).

We utilized unsupervised ADMIXTURE analysis results for further evaluation of Ancient North Rus and Ancient Major Rus genetic clusters (Supplementary Fig. 31, 32, 33). Among the genetic subclusters of **Ancient Major Rus** (Supplementary Fig. 30), our Rus’ samples of the **Rus_Core** group separate from the most previously published medieval groups. However, several medieval individuals from the burial site Shekshovo 9 (VolgaOka_MA2), individuals from supposedly "Viking" (Scandinavian-type) burials from Gnezdovo, Ladoga (Staraya Ladoga), and Pskov, as well as single individuals from the Slavic-related cemetery from Slovakia and Croatia (Croatia_Gornji) intersect with the periphery of the **Rus_Core** subcluster (Supplementary Fig. 30). On the PCA plot, previously published samples from the Gnezdovo^100^ overlapped with individuals from the burial site Nikolskoe (individuals DB56 and DB60). Another Gnezdovo sample overlapped on the plot with sample DR24 (Komarovka, Kursk). One sample from the Pskov^100^ located together with DR5 (St. George's (Yuriev), Monastery, Novgorod), Ya07 (Yaroslavl), AB214 (Pereslavl), and DR26 (Komarovka, Kursk). One individual from Ladoga^100^ is placed on the PCA plot next to the sample DR32 (Komarovka, Kursk).

In the PCA plot, three previously published medieval samples fall into the **Rus_Baltic** genetic subcluster. Among them, one individual from Ladoga is located together with an individual AB35 from the burial site Novinki (Vologda). An individual from Shestovitsa (Chernigov) is closer to individual AB211 (Pereslavl). And one individual from the Early Slavic cemetery in Central Europe (Czech_EarlySlavs) is also positioned on the PCA among **Rus_Baltic** genetic subclusters. Remarkably, other individuals from the Ladoga^100^ positioned with groups of northern European (Scandinavian) origin (Scandinavia_Viking group, see Supplementary Information S2.2.1), along with individuals from the Bodzia site (Poland_Bodzia_VKE) and individuals from Kurevanikha (Russia_Kurevanikha_VKE) (Supplementary Fig. 30). The placement may suggest a shared northern European ancestry among these people, apart from Rus’ genetic groups.

Notably, among the individuals of IA and MA, on the PCA plot, Rus_North_1 and Rus_North_2 subjects exhibit affinity only with individuals from Bolshoye Davydovskoye 2 (VolgaOka_IA)^97^, a burial site representing an IA culture (3rd–4th centuries CE), and two medieval individuals from the Shekshovo 9 (VolgaOka_MA), a burial site associated with a large medieval settlement (10th–12th centuries CE) (Supplementary Fig. 30). Both burial sites are located in the northeastern part of Central Rus’ Volga-Oka region. They exhibit Volga-Fennic cultural appearance (Bolshoye Davydovskoye 2) and the blend of Slavic and Finnish elements (Shekshovo 9) cultural context partially overlapped with Uralic-speaking populations. These findings may indicate a relationship between our people from the Rus_North_1 and Rus_North_2 genetic subclusters and some Fenno-Ugrian (Uralic) populations.

In summary, our findings indicate a partial continuity of the Ancient Major Rus’ genetic cluster with early Slavic individuals from the non-Rus’ regions of Eastern Europe. Furthermore, these early Slavic individuals exhibit genetic differences when compared to other major populations in Europe.

#### S2.2.1 Ancient Rus’ towns

Medieval towns served as the centers of power and economic activity, with diverse social groups, including military elite, merchants, craftsmen and clergy. The concept of the Rus’ towns as the melting pots that integrated individuals from various regions and formed urban communes with diverse ethnic structures is based on historical and archaeological evidence and has gained widespread recognition^2,99–101^.

At the same time, a substantial segment of the urban populations may be associated with the local communes linked to the territory where the ancient town was founded. To test this hypothesis, we performed PCA analysis for samples of urban burials of the medieval Rus’ cities: Yaroslavl, Pereslavl-Zalessky (Pereslavl), Novgorod, and Old Ryazan (Supplementary Fig. 34а). We found no correlation between the geographical location of the city and the genetic structure of its population. Furthermore, we confirmed a high level of genetic heterogeneity among the medieval inhabitants of ancient Rus' cities. Notably, the genetic diversity of the ancient urban population encompasses all ancient Rus’ and present-day East Slavic groups.

#### S2.2.2 Scandinavians and Social military elite (druzhina)

It is widely accepted that the princely retinue and military elite of ancient Rus' were predominantly composed of Varangians, who were likely recruited as warriors and may have had Scandinavian or other Northwestern ancestry^104^. The "Varangian question," which pertains to the role of Scandinavian Vikings in this historical context, has long been a subject of considerable scholarly interest and debate. Research in this area addresses several critical issues related to the early history of Rus', particularly the contribution of Scandinavians to the formation of the ancient Rus' state. Historical chronicles frequently reference the Varangians, who served as essential warriors and mercenaries for rulers such as Vladimir the Great and Yaroslav the Wise during military campaigns and internal governance. Furthermore, archaeological findings from significant sites along trade routes—including Ladoga near Novgorod, Gnezdovo near Smolensk, and Shestovitsa near Chernigov—have uncovered numerous warrior burials dating to the 9th and 10th centuries CE. The graves are believed to be of Scandinavian origin, as indicated by the presence of artefacts characteristic of Scandinavian culture found within them^105–108^.

To investigate the presumable contribution of individuals of Scandinavian descent to the population structure of ancient Rus', we conducted a comparative analysis of our Rus’ samples with previously published genome sequencing data of the European individuals of the Viking Era dated to the 8th-11th centuries CE that also include the samples on the territory of Rus’^102^. Here, we designated all these individuals as Viking Era samples (VKE). Among them, we defined genetically homogeneous groups originating from Scandinavia as "Scandinavian Vikings.". This group is characterized by genetic similarity to early (pre-medieval) Scandinavians and is associated with distinct Viking’s artefacts, including weaponry and boat burials. In our subsequent analysis we assumed that VKE samples collected from Rus’ territory may not represent this specific "Scandinavian Viking" group^102^.

We projected our newly Rus’ samples onto a PCA plot, along with VKE samples from various archaeological sites in different territories, including the historically described Viking areas in Scandinavia, as well as the regions of Eastern Europe and Rus' (Supplementary Table 7). The “Scandinavian Vikings” group include the following: Denmark_IA, Norway_IA, Sweden_IA, Estonia_Salme_EarlyViking, Denmark_Sealand_EarlyViking, Norway_Viking and Iceland_Viking. Viking Era individuals also included samples of MA groups from from the ancient Rus' area: Russia_Pskov, Russia_Ladoga, Russia_Gnezdovo, Ukraine_Shestovitsa, Ukraine_Chernigov and Russia_Kurevanikha (Supplementary Fig. 30). Based on PCA results we have identified that some VKE individuals overlap with ancient Rus' samples (Supplementary Fig. 35). On the PCA plot, there is only one VKE sample from Scandinavia positioned within the **Ancient North Rus' cluster** (VK56, see also Supplementary Informations S2.6), but 68 out of 445 analyzed VKE samples show genetic affinities to the newly sequenced ancient Rus' individuals of the **Ancient Major Rus'** genetic cluster.

It should be mentioned that there was an uneven distribution of these VKE samples among subclusters within the **Ancient Major Rus'** genetic cluster (Supplementary Fig. 35). The majority of previously published VKE samples are positioned on the PCA within the **Rus_SWest** subcluster (n=44), whereas only 13 and 11 VKE samples, respectively, are located within the other two subclusters, **Rus_Baltic** and **Rus_Core**.

Notably, among the 68 previously published VKE individuals genetically linked to our ancient Rus’ samples, 14 were excavated within the Rus’ territory at well-known sites such as Gnezdovo and Ladoga (Staraya Ladoga). These sites have yielded Scandinavian artefacts, confirming a permanent Scandinavian presence. However, for these 14 individuals, neither the archaeological context of their burials nor previous genetic data indicates their Scandinavian origin^102^.

It is noteworthy that genetic affinity with the Ancient Major Rus' genetic cluster has been observed for non-Rus’ VKE individuals who were buried in key European trade centers or along major commercial routes during the Viking and early medieval periods, such as the island of Gotland (Sweden)^2,102,103^. Previously published data indicate that these VKE people are of non-Scandinavian and non-native origin, which has been confirmed by genetic analysis, as well as isotope analysis and archaeological items^102^.

Additionally, we analysed Y-chromosome haplogroups among the 44 male VKE samples that exhibited genetic affinity to Ancient Major Rus’ genetic cluster (Supplementary Table 7). We revealed that predominant Y-chromosome lineages among them (30% of these VKE individuals) belong to the haplogroup R1a branches that are prevalent in both ancient and present-day Slavic populations (Supplementary Table 7, Supplementary Fig. 55). It is quite probable that these individuals are of Slavic rather than Scandinavian descent. Significantly, the same R1a Y-chromosome lineage also represent the majority haplogroup (90%) in males from the two studied military burial sites, Gnezdilovo and Nikolskoe, of аncient Rus’ (Supplementary Table 4).

Based on archeological data, these two burial sites from our study—Nikolskoe and Gnezdilovo —indicate the presence of people connected to the military elite ("Druzhina" in the *Primary Chronicle*). In addition to prestigious decorations and coins, the presence of artefacts like notable weapons and horse equipment in those burials indicates that the people buried during this time were people of considerable status, perhaps affiliated with the princely retinue (see for archaeological and anthropological information of that burials sites Supplementary Informations S1.1.1.4 and S1.2.1.1, Supplementary Fig. 18. The Gnezdilovo 12 burial site. Burial 19 (DB66); Supplementary Fig. 19. The Gnezdilovo 12 burial site. Burial 24). Even though the military elite's archaeological sites being located in two distinct regions of ancient Rus’—Nikolskoe in Northern Rus’ and Gnezdilovo in the Volga-Oka region—the individuals interred at these sites displayed genetic heterogeneity characteristic of the ancient Rus’ population, similar to the inhabitants of Rus’ cities (Supplementary Fig. 34b), rather than “Scandinavian Vikings”. Notably, a significant number of these individuals from military elite burials, 7 out of 18, were found to be members of the Rus_SWest genetic subcluster.

Our data suggest that the Scandinavian contribution to the military elite at Gnezdilovo and Nikolskoe was likely negligible. The genetic evidence indicates that this elite was primarily composed of individuals of local or broader Eastern European origin, underscoring the complex, multicomponent nature of the Rus' military elite's formation, which cannot be attributed solely to Scandinavian influence. These findings challenge the hypothesis of a predominantly Scandinavian Viking ancestry for the individuals buried at Gnezdilovo and Nikolskoe. The observed genetic overlaps of genetic profiles of Northern Europe and ancient Rus’ samples can be explained by shared ancestral genetic origins and the high mobility of various population groups of the Viking, facilitating gene flow across Eastern and North-Western Europe^103–105^. It is important to note that our sample set from the territory of ancient Rus' included individuals interred according to ancient Rus' or East Slavic burial practices, and did not encompass specimens from graves characterized by Scandinavian burial traditions.

#### S2.2.3 Genetic outliers in Rus’ area

In the burials attributed to the East Slavs in the Southern Rus' towns, we revealed three genetic outlier samples. On the PCA plot, two of them (AB23 and AB24) from the necropolis at Knyazhya gora hillfort exhibit genetic affinity to Caucasian populations (Fig. 2, 4). In the male AB23 we identified Y-chromosome haplogroup J2a1 (J-Y12379), which is common among both ancient and present-day men of Caucasian ancestry. The mitochondrial lineage of the female AB24 corresponds to the Central and East Asian mitochondrial branch K2a5b (See Supplementary Informations S2.7.1). In contrast, other Rus' samples in our dataset are predominantly carriers of European mitochondrial lineages (Supplementary Table 4, Supplementary Fig. 60). The third outlier individual, AB56, from the burial near the Boris and Gleb Church in Chernigov, has a genetic profile similar to the Volga Bulgar individual (AB44). Both of them have a notable level of genetic ancestry related to ancient East Asian populations (Fig. 4). Consistent with this, AB56 is a carrier of East Asian mitochondrial haplogroup M10a1 (See Supplementary Informations S2.7.1). Despite these striking genetic differences, all three outliers were buried according to the East Slavic/Rus' burial practice, suggesting integration of these non-Slavic individuals into the local Rus' society. Notably, with respect to the deep ancestral roots of Rus' and non-Rus' populations, we also noted that the mitochondrial lineage, mtDNA haplogroup R1a2, of AB22 individual from Knyazhya gora is linked to Central Asian populations (See Supplementary Informations S2.7.1), whereas autosomal marker analysis places it on the PCA plot within European genetic variation, pointing to some ancient migration preceding the Rus' state formation and subsequent assimilation within the Slavic-related population.

### S2.3. *f*3-, *f*4-statistics

#### S2.3.1. Ancient populations

To measure allele sharing between samples from ancient Rus’ and other ancient populations, we calculated outgroup *f*3-statistics of the form *f*3 (Ancient_pops, Ancient_Rus’; Mbuti) for both **Ancient Major Rus'** and **Ancient North Rus’** genetic clusters (Supplementary Table 8), and further for each Ancient Rus’ genetic subcluster (Rus_Core, Rus_Baltic, Rus_SWest, Rus_North_1, Rus_North_2). The highest *f*3-statistic estimates indicate a higher level of common alleles between two tested populations. We performed outgroup *f*3-statistics calculations for several chronological periods from the early BA to the MA. For analysis, IA and MA we classified samples into three chronological subgroups: the 1st millennium BCE (1000-0 BCE), the first centuries of the Common Era, pre-Slavic period (0-700 CE), and MA after the Slavic expansion in the Eastern Europe (700-1400 CE) (Supplementary Table 9, 10, 11, Fig. 5, 6). Outgroup *f*3-statistics of the form *f*3(Ancient_IA, Rus; Mbuti) were also calculated for individual samples of every **Ancient Major Rus’** genetic subclusters (Rus_Core, Rus_Baltic and Rus_SWest), Ancient North Rus’ genetic subclusters (Rus_North_1 and Rus_North_2), and ancient Mordva group (Supplementary Fig. 36, 37, 38, Supplementary Table 12).

Next, to measure the genetic affinity of our Ancient Rus’ subcluster to preceding ancient populations we calculated *f*4-statistics of the form *f*4 (Ancient_Rus’, Mbuti; Ancient_pops, Ancient_pops) (Supplementary Table 13, 14).

All tested Rus' genetic clusters and subclusters had Baltic region populations among their highest *f*3- and *f*4-estimates during all tested chronological periods: 1st millennium BCE, 1st millennium CE before Slavic expansion in Eastern Europe (0-700 CE) and after 7th century CE. Despite the geographic proximity of the BA Fatyanovo group to ancient Rus’, Fatyanovo samples did not share a notably higher number of alleles with any of the Ancient Rus’ genetic subclusters that we examined (Supplementary Fig. 39). We detected higher affinity to BA Steppe-related groups for Rus_North_2 group (Russia_MLBA_Sintashta, Russia_Srubnaya, Andronovo) and to Central European BA populations (Italy_Sardinia_EBA and England_BellBeaker) for Rus_SWest subcluster in comparison to other ancient Rus’ groups Supplementary Table 13, Supplementary Fig. 40).

In addition, to measure the relative affinity of Ancient Rus' genetic subclusters to BA groups with ancient Siberian and European Neolithic farmer ancestries, we calculated *f*4-statistics in the form of *f*4 (Krasnoyarsk_BA/Greece_BA_Mycenian, Mbuti; Test, Latvia_BA), where Test is Rus’ genetic subclusters or other ancient groups dated from the Iron Age to the Middle Ages (Supplementary Table 15). We found significant gene flow between both Rus_North_1/Rus_North_2 groups and Siberian BA groups (Supplementary Fig. 41).

We performed cladality tests in the form *f*4 (Medieval_pops, Mbuti; Major_Rus', Test) for all Rus’ genetic subclusters to measure their allele sharing with other medieval populations. We would expect no statistically non-zero estimations of *f*4 (Z score > |3|) if our Rus’ genetic cluster was cladal with a tested MA group (Test). The results revealed notable differences between three Rus’ genetic subclusters (Rus_Core, Rus_Baltic, and Rus_SWest) in their genetic affinity to other groups (Supplementary Fig. 42).

To test a hypothesis on the admixture origin (as observed in the PCA analysis) of the Rus_Middle genetic group out of the Slavic-related and Northern Rus’ populations, we performed admixture *f*3-statistics of the form *f*3 (Ancient_Major_Rus/VolgaOka_MA, Ancient_North_Rus; Rus_Middle). We included the VolgaOka_MA groups in the analysis, along with our Slavic-related groups. All tests, with the exception of the earliest VolgaOka_MA1, show significantly negative admixture-*f*3 values, indicating the potentially mixed Slavic and Fenno-Ugrian ancestry of Rus_Middle genetic group (Supplementary Table 16).

#### S2.3.2. Present-day populations

To measure allele sharing between samples from ancient Rus’ and present-day populations (Target), we calculated outgroup *f*3-statistics of the form *f*3 (Ancient_Rus’, Target; Mbuti) for Ancient Rus’ genetic subclusters and for individual samples (Supplementary Table 18). Chalmny-Varre samples (Murmansk region, 17th-19th centuries CE^106^ were added to the present-day set. To avoid bias related to specific ancient DNA substitutions during the analyses of a combined set of 173 present-day Eurasian and North African populations and tested ancient groups, we performed two rounds of the outgroup *f*3-statistics analysis - using all SNPs and using only transversions (Supplementary Table 17, 18, 19, 20). Both approaches produced similar results; therefore, further tests were performed on the whole set of SNPs (Supplementary Fig. 43).

We also performed comparative *f*3-statistics analysis of the Ancient Rus’ genetic subclusters with the present-day East Slavic and Northern Russians groups (Supplementary Table 19, Supplementary Fig. 44), as well as with the all present-day Slavs (East, West, and South Slavs). A similar assay was performed to test allele sharing between present-day northern Russians from the Vologda and Arkhangelsk regions (Supplementary Fig. 45).

The f3-statistics results indicate different allele sharing patterns with present-day groups for both main Ancient Rus' genetic clusters (Supplementary Table 18, Supplementary Fig. 43, 44). All subclusters of the Ancient Major Rus' showed high levels of shared genetic drift with present-day Lithuanians, probably due to high percentages of common ancestral alleles between the ancient Central-East genetic substratum, represented by Lithuanians and most Eastern-European populations^107^. In contrast, all subclusters of the Ancient North Rus' share more alleles with present-day Veps, Karelians, Saami, and Northern Russians from the Arkhangelsk region (Supplementary Table 18, Supplementary Fig. 43, 44). Notably, we observed high levels of allele sharing between Ancient Northern Rus' individuals with Russians from Arkhangelsk rather than Russians from Vologda (Supplementary Fig. 45).

Next, in order to compare the genetic affinity of present-day East, West, and South Slavs to our Rus’ subclusters and other ancient populations (Ancient_pops) dated to the 1st millennium CE, we performed an analysis of the outgroup *f*3-statistics in the form *f*3(Ancient_pops, Modern_pops; Mbuti) for ancient groups geographically or historically appropriated to Rus’. We revealed distinct differences in allele sharing with present-day Eastern-Europe populations between Ancient_Major_Rus’ subclusters and other tested IA and MA groups (Supplementary Table 21, Supplementary Fig. 46).

For further *f*4-test with present-day populations, we filtered out transitions. To test cladality with present-day groups, we calculated *f*4-statistics of the form *f*4 (Mbuti, Modern_pops; Ancient_Rus', Target), where Modern_pops are present-day Eurasian and North African populations, and Target is a set of geographically and genetically appropriate present-day East European populations (Supplementary Fig. 47, Supplementary Fig. 48). For Ancient North Rus’ genetic subclusters, we also test cladality with present-day populations with Ancient Siberian ancestries (Supplementary Table 22). The option *inbreed* was set to YES in all *f*4-statistics calculations. We would expect zero, or no statistically different from zero, estimation for *f*4 if the tested ancient Rus' group was cladal with the present-day Target population.

By calculating *f*4-statistics, we found that Ancient Major Rus' individuals of the Rus_Core, Rus_Baltic, and Rus_SWest subclusters are cladal with present-day East Slavs (Belarusians, Ukrainians, and Russians) and not cladal with West Slavs (Czech), South Slavs (Bulgarian, Croatian, and Serbian) or Baltic and Baltic Finnic populations (Estonian, Finnish, Karelian, and Veps) when we compare them to a set of present-day non-Slavic European populations (Z score < |3| in all tests) (Supplementary Table 22, Supplementary Fig. 47). Both Rus_North_1 and Rus_North_2 are cladal with present-day Russians from Arkhangelsk and Saamis (Supplementary Table 22, Supplementary Fig. 48). In addition, when testing Baltic Finnic groups (Estonian, Finnish, Karelian, Estonian, Veps), the Rus_North_2 subcluster is cladal with present-day Veps and Karelian populations. Notably, the Rus_North_1 subcluster is cladal with the Veps when compared to all European populations, except for the English, Scottish, Basque, and Sardinian groups, and it shows greater affinity to Karelian and Finnish populations (Supplementary Fig. 48).

Overall, our results indicate genetic continuity between the Slavic-related Ancoent_Major_Rus’ genetic clusters and present-day East Slavic groups. Ancient_North_Rus’ genetic cluster showed notable affinity to several present-day Fenno-Ugrian (Uralic) groups from the Northern Eastern European regions, however, we found no direct genetic continuity between medieval groups from the Northern Rus’ region and exact present-day northern Russian populations.

### S2.4. Admixture modeling using qpAdm

To trace the population history of the ancient Rus' individuals through different time periods, we performed qpAdm admixture modeling. We used *qpAdm* software from ADMIXTOOLS v. 7.0.1^108^ to model the ancestry of our ancient Rus’ samples. We selected the "allsnps: YES" option and ran the analyses on the “1240k dataset.” We excluded individuals with fewer than 20000 SNPs from the *qpAdm* and ran several separate *qpAdm* tests. As testing every possible combination of sources is challenged due to the large number of potential source populations, we ran the analysis just with combinations of one, two, or three sources. We considered the *qpAdm* models to be acceptable if they had p>0.05 and the value of the admixture proportion was above zero.

In the first step, to assess the main ancestries present in Ancient Rus’ genetic subclusters, we performed a distal *qpAdm* analysis and used the earliest ancestral sources of Eurasian hunter-gatherers and Neolithic populations (Russia_DevilsCave_N, Iran_GanjDareh_N, Russia_Tyumen_HG, Luxembourg_Loschbour, Germany_EN_LBK, Russia_Karelia_HG, Israel_Natufian). Russia_Krasnoyarsk_BA was also included in the distal qpAdm sources list because no other ancient sources of ancient Siberian ancestry were available. We used the following outgroup populations in distal modeling: Ethiopia_4500BP, ONG.SG, Mixe.DG, Ami.DG, Papuan.DG.

The best distal models for all tested Ancient Rus’ subclusters comprised approximately 60-80% ancestries from Neolithic farmers (Iran_GanjDareh_N or Germany_EN_LBK) and 20-30% of European Eastern hunter-gatherers ancestry (Russia_Karelia_HG or Russia_Tyumen_HG) (Supplementary Table 23). Eastern Asian (Russia_DevilsCave_N) and/or ancient Siberian (Russia_Krasnoyarsk_BA) ancestry gradiently increases from 2.5-3% in the Rus_SWest subcluster up to 14-18% among Ancient Rus’ North genetic subclusters, in line with the similar tendency of increasing the ancient Siberian components that we revealed in the ADMIXTURE analysis (Fig. 4).

Next, we run a second (proximal) round of *qpAdm* with more relevant sources. To date, there is only one report on the genomic data of the populations inhabiting the Rus'-related area during the period spanning the first millennium BCE to the first millennium CE^97^.

All local populations of the forest-steppe and forest zones of Eastern Europe practiced cremation, leaving no anthropological material available for genetic analysis. Consequently, all possible models of origin for Rus' people based on post-BA groups will be influenced only by populations from neighboring territories for which anthropological material is accessible for analysis, rather than local. To mitigate this potential bias, we refrained from conducting simulations involving groups from the period of widespread cremation in the East European Plain. Instead, we focused our modeling on BA, for which material is available from both the ancient Rus’ area and adjacent regions. The point of our admixture modeling is not to find the exact admixture sources of the tested ancient Rus' groups, which is challenging currently since we don't have any real ancestry sources. Rather, we assess the basic ancestral substratum of previously untested medieval North Rus' and Slavic-related populations. The outgroups used in analysis were Ethiopia_4500BP, Russia_DevilsCave_N, Russia_Karelia_HG, Iran_GanjDareh_N, Russia_Tyumen_HG, and Germany_EN_LBK). We selected 22 ancient BA groups as potential proximal sources for modeling the ancestries of our Ancient Rus’ genetic subclusters^109,110^:

1. Ancient Anatolian and Iranian ancestries: Italy_Sardinia_EBA, Armenia_EBA_KuraAraxes, Turkey_Alalakh_MLBA, Greece_BA_Mycenaean, Turkmenistan_Gonur_BA_1, Uzbekistan_Bustan_BA

2. Ancient Siberian and Eastern Asian ancestries: Russia_Krasnoyarsk_BA.SG, Russia_Bolshoy, Mongolia_LBA_Khovsgol_6

3. Steppe-related Eurasian ancestries: Russia_Afanasievo, Andronovo, Russia_MLBA_Sintashta, Russia_Shamanka_EBA.SG, Russia_Srubnaya, Voronezh_LBA

4. European ancestries: Poland_GlobularAmphora, Czech_CordedWare, England_BellBeaker, Russia_Fatyanovo_BA

5. Baltic-related ancestries: Sweden_Gotland_PittedWare_BattleAxe, Estonia_MN_CCC, Estonia_BA.SG, Latvia_BA

In all tests, to reject the models, we used p<0.05 as a cutoff. All component proportions must fall between 0 and 1, and the standard error must not be greater than the percentage value for fitting models. Out of all the fitting models, we selected the model with the minimal number of sources and the highest p-value for each of our tested Rus' and other medieval groups. As numerous three-way models were found for every tested Ancient Rus’ genetic subcluster (Supplementary Table 24) we performed the rotation strategy to select the best sources. To increase the resolution during the modeling, we tested several models by removing some populations (Estonia_BA.SG, Latvia_BA, Russia_Fatyanovo_BA, Greece_BA_Mycenaean, Russia_Krasnoyarsk_BA.SG, and Russia_Bolshoy) from the sources and placing them into outgroups (Supplementary Table 25). In the first stage of proximal modeling, we identified several models that cannot be rejected with two or three sources out of the 22 tested groups. All of them are based predominantly on Baltic-related sources (60-82%). By rotating strategy, we showed that Latvia_BA is the better Baltic region source for Rus_North_1, Rus_SWest and Anc_Mordva genetic groups when compared to the geographically related Estonia_BA (Supplementary Table 25 a, b). Ancient Anatolian-related ancestry could be represented by Greece_BA_Mycenaean in most tested groups and accounts up to 32% in the Rus_SWest subcluster, but is minimal among Ancient North Rus’ groups (3-6%). Remarkably, Rus_Core subcluster gives a perfectly nonrejected result when Greece_BA_Mycenaean is included in the admixture model, among other BA sources related to ancient Anatolian ancestries (Supplementary Table 22f, Supplementary Fig. 49a). Notable, the best fitted models for Rus_North_2 and Anc_Mordva groups include Turkey_Alalakh_MLBA, while other subclusters modelled with Greece_BA_Mycenaean. This result points to differences in genesis of these groups, in particular, Rus_North_1 and Rus_North_2, which are similar according to the PCA. Northern Rus’ region inhabitants The disparities observed in the genetic composition of local groups in the Northern Rus’ region align with historical evidence suggesting that the population in the North resided in small, migratory groups that traversed extensive territories and participated in fur hunting^2^, while the Northern regions served as a crossroad for both western (from Novgorod and Ladoga) and southern (Volga region) migration flows^21^. The genetic origins of these two groups varied slightly. In particular, the genetic composition of the group in the Poonozhye region (Rus_North_2) include contributions from not just from Anatolian-related ancestors (Greece_BA_Mycenaean), as Rus_North_1 do, but also from a population carrying an ancient Iranian or Caucasian component represented by Turkey_Alalakh_MLBA in the qpAdm model.

The third ancestral source of ancient Rus’ genetic subclusters is ancient Siberian/Eastern Asian ancestry, which is notable (15-20%) in the Ancient North Rus’ and low among the Ancient Major Rus’ (4-8%) (Supplementary Fig. 49a). Because of the small variability in the proportions of related sources, we use the model with Latvia_BA, Greece_BA_Mycenaean, and Russia_Krasnoyarsk_BA to describe all tested Rus' subclusters with unified sources (Fig. 7).

Additionally, we conducted 2- or 3-way qpAdm modeling at the individual level for genetic outliers identified in our ancient Rus’ dataset, revealing variations in the potential BA ancestry composition of these samples (Supplementary Fig. 50). The best-fitting models for all outliers indicate that their origins can be traced from two or three ancestral sources. The two sources align with those identified in the ancient Rus’ genetic subclusters: Baltic-related ancestry (Latvia_BA) and Ancient Siberian ancestry (Russia_Krasnoyarsk_BA). The third source is related to Ancient Anatolian/Iranian ancestries, which differ from those represented among the ancient Rus’ genetic subclusters. The best non-rejected models for Yaroslavl region individual from Voronovo (AB10), from Chernigov (AB56) and Mordova individual (AB44) from the Syut-Siriami burial ground (Chuvashia) include 25-30% of Turkey_Alalakh_MLBA source. Both Kyazhya Gora individuals (AB22 and AB23) can be modeled with 56% ancestry from Armenia_EBA_KuraAraxes as the third ancestral source.

We also tested a set of previously published medieval populations to compare their presumably genetic ancestry with Rus’ genetic subclusters. Given the extensive computational requirements for modeling each group from all 22 ancestral populations, we restricted these tests to the 11 most common in different model sources (Latvia_BA, Greece_BA_Mycenaean, Turkey_Alalakh_MLBA, Mongolia_LBA_Khovsgol_6, Russia_Krasnoyarsk_BA.SG, Poland_GlobularAmphora, Armenia_EBA_KuraAraxes, Andronovo, Russia_MLBA_Sintashta, Russia_Srubnaya, Russia_Fatyanovo_BA) (Supplementary Table 26, Supplementary Fig. 49). The results show a difference in ancestral structure of ancient Rus' genetic subclusters compared to medieval Scandinavian groups and groups from Central and Southern Europe.

### S2.5. Analysis of Runs of Homozygosity

Different types of parental relatedness can be detected by distributions of inferred runs of homozygosity. In cases when the sum of all ROHs more than 20cM of one individual is greater than 50 cM then parents of this individual are likely to be first- or second-degree cousins or closer relatives. To infer runs of homozygosity, we used hapROH software^111^. In our study, 8 samples exhibit ROHs >20 cM (Supplementary table 27). Moreover, they are all from the Northern Rus’: 3 samples are from Voezero (DR12, DR19, DR22), 2 individuals are from Nefedyevo (DB17, DB31), and one sample is from Minino (DB41), Shuygino (DB69) and Novinki (AB35). The most homozygous individuals, DB31 (Nefedyevo), AB35 (Novinki) (Supplementary Fig. 51b,c[)](https://drive.google.com/file/d/19TXJLrqcF4rmREcvgRDyFH1KXsjIisRK/view), and some of those that were mentioned above have the length distribution of ROHs indicating that their parents are likely to be first or second cousins. DR19 and DR22 (Supplementary Fig. 51d) were siblings of closely related parents, and both died in early childhood. DB69 exhibits long ROHs that suggest that his father DB71 (Shuygino), identified by the READv2 and IBD methods, and his mother were related. Additionally, DB69 and DB72 (Shuygino) were predicted to have a sibling relationship and have intersecting parts of ROHs (Supplementary Fig. 51e). DB25 and DB27 from Nefedyevo, predicted as relatives of the first degree and have intersecting parts of ROHs on the third chromosome (Supplementary Fig. 51f). ROHs of 4-8cM indicate background relatedness that dates back several generations ago. The majority of Northern burials have a large number of ROH less than 20cM in length, which may indicate a small population size or a consequence of founder effects in tight-knit groups.

Individuals from Central and Southern Rus' are characterized by the absence of long ROHs (except for DB64 from Gnezdilovo, which exhibits ROHs of 12-20 cM length).

In medieval Rus' 11th–13th centuries CE, there existed prohibitions on closely related marriages, reflecting both church laws and secular regulations. For instance, the written Orthodox canon law, such as the Efremov Kormchaya (12th–13th centuries CE), generally prohibited marriage within the seventh degree of consanguinity^112,113^. Despite the gradual process of Christianization of the Northern Rus’ during the 11th–13th centuries CE, and the introduction of religious prohibitions against marriages between close relatives in this region^114^, we identified individuals whose parents were close relatives, which is indicated by long ROHs. In contrast, individuals from other ancient Rus’ regions are generally lacking long ROHs. Middle levels of inbreeding represented by shorter ROHs may indicate that there were small, isolated communities in Rus’ North populations in Minino, Nefedyevo, and Shuygino.

### S2.6. IBD analysis results

Based on the identification of haplotype blocks of certain lengths that are shared between individuals (see *Material and Methods,* Supplementary table 28), we show numerous shared IBD blocks among individuals within the Northern Rus' region, indicating close relatedness of most tested individuals that formed the family clan (see details in Supplementary Note S2.8.2, Supplementary Fig. 52, 53).

Next, we observe IBD sharing between several individuals from geographically distant Rus' regions (Supplementary Fig. 54). A total of 13 individuals from the Beloozero region presented IBD connections (17-26 cM) with other sites on the Northern Rus', both Beloozero and Poonezhye regions, implying genetic links between Northern Rus’ populations from different locations. Twelve Beloozero samples shared long, yet singular, IBD fragments (up to >33 cM) with individuals from Yaroslavl, Pereslavl, and Old Ryazan (Central Rus' Volga-Oka region) These findings suggest a potential common ancestry and 6 to 7 degree relatedness^111^. These data further demonstrate the certain genetic links between Fenno-Urgian and Slavic-related groups. Distant relatedness is proposed by shared IBD segments >17 cM between Novinki (Vologda) and Gnezdilovo individuals. We also found that the close relatives from the Nefedyevo burial site (DB5, DB17, DB18, DB25, DB27) and from the Minino (DB38), along with the outlier (AB10) from the Voronovo (Yaroslavl), share IBD fragments (the longest > 29 cM) with the ancient Mordva individual (AB30) of Fenno-Ugrian cultural appearance from the Muranka burial site. All these individuals display the notable ancient Siberian component (Fig. 4), and the data demonstrate the genetic exchange between two distant Fenno-Ugrian (Uralic) groups.

Regarding the samples from the Rus’ regions with Slavic-related burials, IBD connections were revealed between geographically close pairs of individuals from Southern Rus’ (AB22 (individual from Knyazhya gora) and AB29 (Moiseevskoe), as well as between pairs of individuals from geographically distant urban cemeteries: DR5 (Novgorod) with AB213 (Pereslavl), and AB136 (Lubech) with DR35 (Luzhki).

Although we identified shared IBD segments between individuals from different regions of Rus’, these segments are represented solely by single shared fragments. Such patterns of IBD sharing suggest the existence of distant genetic connections and indicate recent common ancestry or interactions that may have occurred centuries prior^111^. This period predates the consolidation of East Slavic groups and the formation of the Rus’ state, providing evidence of genetic and cultural interaction within the Rus’ region during the earlier periods.

Finally, by screening IBD sharing of all pairs containing one sample from our dataset and other previously published medieval individuals, we found four shared IBD fragments (17–22 cM) between our Rus' individuals and a set of 438 previously published European medieval samples (Supplementary Table, 29 Supplementary Fig. 54), potentially suggesting shared ancestors. One shared long IBD fragment (>26 cM) was revealed in Rus’ area between the outlier AB10 (Voronovo) and an individual from the Suzdal region of Rus’ (SHE003, Shekshovo)^97^. Both samples (AB10 and SHE003) are from the Central Rus’ region but projected on the PCA plot closer to samples of Ancient North Rus’ genetic clusters (Supplementary Fig. 30). Remarkable that SHE003 represents a burial site associated with multi-component culture (Volga Finnic and Rus’ "Slavic") and was also proposed in a previous study as a possible outlier from genetic clusters that were tested there^97^.

Only one of the shared IBD fragments was found between Rus’ individual and Scandinavian Viking sample VK505^102^, who was from a Viking boat burial Salme I, Estonia Another VKE individual (VK56 from the largest and most important harbour and commerce site on the western coast of the island of Gotland) shared IBD fragments of > 20cM and was therefore related to two individuals from the Northern Rus’ area DB61 (Nikolskoe) and DB36 (Nefedyevo). This is the single individual from Scandinavia (Gotland) who is positioned on the PCA plot among the samples of the Rus_North genetic subclusters (Supplementary Fig. 35, and therefore could be proposed to be a migrant to the Gotland Island from the Northern Rus’ area.

A shared single IBD fragment > 16cM was found between the Yaroslavl individual (Ya71) and the early Medieval individual from Croatia (I26748). Notably, this male sample from the Balkan Peninsula had Anatolia Roman/Eastern Europe ancestry and belongs to the Y-chromosome haplogroup R1a1a1b1a1a1a^107^, which is distributed now predominantly among present-day East Slavs (See Supplementary Infromations S2.8.2, Supplementary Fig. 55).

Our IBD tests were restricted to previously published medieval samples from Europe. This choice was primarily motivated by the preferential connections of Ancient Rus' with European populations, as well as the genomic data we obtained (including PCA and ADMIXTURE), which indicate similarities in the genetic profiles of our samples with representatives of predominantly European populations. However, we cannot exclude the potential genetic links of some of our samples with populations from the Caucasus and Eastern Eurasia (in particular, for samples AB23, AB24 (Knyazhya gora), AB56 (Chernigov), and AB44 (Syut-Siriami**)** with notable Caucasian and Asiatic genetic profiles), although we propose that such links are rarer in our whole ancient Rus' dataset.

### S2.7. Analysis of uniparental markers

#### S2.7.1. Mitochondrial DNA diversity

Exploring the mitochondrial DNA (mtDNA) variabilities and phylogeny^115^ (MTree v. 1.02.22369, https://www.yfull.com/) we determined mitochondrial haplogroups for all sequenced individuals (Supplementary Table 4).

In our set of the samples from Rus’ territory (N=188), we found 124 different mtDNA deep haplogroups, falling into ten main Western Eurasian haplogroups (H, U, I, J1, K, T, R1, W, HV+V, and X) and three Eastern Eurasian haplogroups (A, D5, and M) (Supplementary Fig. 56). Haplogroups H and U are common and prevalent in both Northern and other Rus’ regional groups, whereas haplogroups R1, W, V, HV, and X are rare. Mitochondrial lineages H, I, J1, K, R1, T, U5, W are found both in Northern and other Rus’ region groups with the remarkable differences in mtDNA haplotype frequencies between these groups (Supplementary Fig. 56a,b). Haplogroups K1, D5, and U8 are significantly more prevalent in the Northern Rus' region compared to other areas of Rus', where K1 is relatively rare and D5 and U8 are absent. MtDNA haplogroup diversity in Northern Rus’ is substantially lower than in other Rus’ regions.

Next, we analysed the mtDNA sequences that we determined for the individuals not included in the Rus’ cohort (presumably non-Slavic medieval groups: Mordva, Muroma, Volga Bulgars, Latgalians (N=11). All but two of our tested individuals also exhibit mitochondrial haplogroups of European lineages, specifically H, V, J1, and K1. Only two mitochondrial sequences of Volga Bulgarians from Syut-Siriami belong to the Eastern Eurasian clades A-a1 and J2b1a2, the latter of which is widely distributed among various Eurasian populations (see Supplementary Table 4, Supplementary Fig. 57).

Further, we determined the occurrence and frequencies of all main mtDNA lineages that we found in ancient Rus’ (H, U, I, J1, K, T, R1, W, HV+V, X, A, D5, and M) across different European regions from the beginning of the Common Era to the end of the 14th century CE. For this purpose, we used data from the AADR database^109^. The total sample set (N=1815) was divided into the following geographical regions: Northern Europe, Western Europe, Central Europe, Balkan region, Southern Europe, Central-Eastern Europe, and Anatolia/South Caucasus. Frequencies for these mtDNA lineages were also calculated separately for ancient samples from the territories Hungary and Finland, as their populations during this period were characterized by high heterogeneity or were distinct from other populations, respectively. The ancient Rus' samples we studied were also divided geographically into Northern Rus’ region and all others, the latter encompassing geographic Central Rus’, Southern Rus’, and Western/North-Western Rus’ regions (Supplementary Fig. 56a,c, Fig. 2).

The distribution of mtDNA lineage frequencies in Northern Rus' region differs significantly from all other studied ancient populations, including those from the territory of Finland, whose Fenno-Ugrian population might have been expected to show some similarities with Northern Rus’ groups. Notably, unique mtDNA haplogroups absent in other analysed ancient populations of the studied period were identified in Northern Rus' region. These lineages constitute 15% of all mtDNA haplogroups identified in the Northern Rus’ and include K1a1b5, D5a3a1a, H1j3, I3d2, K1c1c, U5a1f, and W3a1t, highlighting a unique maternal genetic signature of this region (Supplementary Table 4).

Overall, the frequency distribution of main mitochondrial lineages in the rest of Rus' is similar to that found in ancient populations of central and northern European areas and slightly different from populations of Southern Europe (Supplementary Fig. 56a,c). The vast majority of mtDNA haplogroups identified in these non-Northern Rus’ regions were also present in other contemporary European populations. This finding indicates a common ancestral mitochondrial DNA pool of ancient Rus’ and ancient European populations. However, as in the northern regions of Rus', we also revealed here several unique mtDNA lineages not found in other European populations of the time. These include R1a2, U4a2e2, V37a1, W1j1, and W3b5a haplogroups. It is noteworthy that all these mtDNA lineages, with the exception of W3b5a, have been identified in urban cemeteries and in genetic outliers. These findings point to a presumably migrant influx having contributed to the local Rus' population structure (R1a2, V37a1, W3b5a) or, in contrast, reflect autochthonous origin in Rus’ area of several maternal lineages (U4a2e2, W1j1) (see Supplementary Table 4 for distributions of related haplogroups among ancient and present-day populations).

We also compared the distribution of mtDNA haplogroups frequencies of our ancient Rus’ samples with a large set (N=5303) of present-day individuals subdivided into several regional groups: Russians Central (from Tula, Vladimir, Novgorod, Pskov), Russians Northern (from Vologda and Arkhangelsk), Russians Central/Southern (from Belgorod, Orel, Stavropol, Saratov), other present-day Slavic individuals (Poles, Ukrainians, Belarusians, Slovaks, Czechs), and present-day neighbouring European groups (Karelians, Veps, Finns, Latvians, Estonians, Lithuanians, Swedes, Hungarians, Turks)^116–122^. The results indicated that distribution of mtDNA haplogroups among the ancient Rus’ population with the exception of the northern Rus’ region is similar to that of present-day Russians and other Slavs. Notable, that the mtDNA haplogroup composition of ancient Northern Rus’ populations differ from the present-day Northern Russian groups (Supplementary Fig. 56a,d). This further emphasizes the specific genetic profile of Northern Rus’ region.

In total, the analysis of mitochondrial haplogroups aligns with the findings from genomic data, revealing distinct genetic differences between two geographically distant regions of ancient Rus’: the main Rus’ territory and its northern periphery. This suggests at least a partially different genetic origin for the two groups that contributed to the formation of the ancient Rus’ state.

Next, we focused on analyzing the complete mtDNA sequences and performed phylogeographic analysis to trace maternal lineages and to assess the affinity of our samples with both ancient and modern individuals. We searched for matching haplotypes in ancient Rus’ and previously published complete mitochondrial genome sequences^109^ (Supplementary Fig. 58). Identical mtDNA sequences suggest a likely relationship of individuals by maternal lineages, as well as the continuity of populations over time and provide additional information to autosomal data.

A number of identical mtDNA sequences were found among samples from burials of Northern Rus'. The complete mtDNA sequence match was also found between two second-degree relatives from Komarovka, Southern Rus’ region. Most of our samples' identical mtDNA sequences were found in people from the same or nearby burial sites, and they most likely belonged to family members (Supplementary Fig. 58). Identical whole mtDNA sequences of haplogroup K1c1c were found in individuals from the geographically distant Poonezhye and the Beloozero regions of Northern Rus’. Interestingly, two individuals from Nefedyevo shared the same mtDNA sequence of K1c1c haplogroup, but have heteroplasmy at position 12810 (Supplementary Fig. 58). In addition, we found that some present-day Finland individuals shared the same mtDNA sequences, demonstrating their maternal lineage link to the subjects from medieval Northern Rus’.

Several pairs of individuals from medieval geographically distant burial sites exhibited identical whole mtDNA sequences belonging to haplogroups H1-16189, H6a1a4, and J1c7a. These findings may indicate a widespread distribution of certain mitochondrial lineages or the long-distance migration of ancient individuals (Supplementary Fig. 58). The maternal lineage connecting Northern Rus’ with Slavic-related groups is represented by haplogroup J1c7a. Identical mtDNA sequences of this haplogroup were found in individual DB54 from Minino (Beloozero region, Northern Rus’) and in individual AB18 from Kleopino/Kokorevo (Tver region, Central Rus’) (see Supplementary Table 4). It is noteworthy that the archaeological context suggests that individual DB54 from Minino was likely a migrant to the Beloozero region (see Supplementary Note S1.1.2). Furthermore, his genetic profile corresponds to the Rus_Core genetic subcluster and he had a Slavic-related Y-chromosome haplogroup R1a (R-CTS8816) (see Supplementary Fig. 55).

Identical mtDNA sequences of haplogroup H6a1a4 were identified in a woman, DB29 from Nefedyevo (Beloozero region, Northern Rus’) and in a man AB17 (Kiryanovo), from the Yaroslavl region (Central Rus’). The complete matching mtDNA sequence was also  detected in sample VK466 from Gnezdovo, dating to the 10th–11th centuries СE^102^ (Supplementary Fig. 58). It is noteworthy that an exact match for this H6a1a4 mtDNA sequence occurs in present-day individuals from Denmark, Finland, Estonia, and the Volga Tatars, revealing its wide distribution across Northern and Eastern Europe (Supplementary Fig. 58).

Surprisingly, we also observed a complete match of mtDNA sequences belonging to haplogroup H5a1a between individual DB37 from Minino (Beloozero region, Northern Rus’), that showed the Slavic genetic profile, and a sample from a Chernyakhov culture burial (3rd century CE) at the Krinichki site in the Middle Subnistria region^123,124^. This finding provides, for the first time, direct genetic evidence of links between the Chernyakhov culture and the early Slavs (Supplementary Fig. 58).

Our study has also identified a significant genetic link: an individual DR10 whose remains were found in a dwelling attributed to the Volintsevo culture from the Kurilovka 2 site (Kursk region), dating back to the late 7th - early/middle 8th centuries CE, shares the same mtDNA sequence as King Béla III of Hungary (12th century CE). This mitochondrial lineage was passed to the king through his mother, Eufrosinya Mstislavovna, a daughter of the ancient Rus' prince Mstislav. The origin of King Béla III's mother is connected to the environment of Novgorod nobility. Consequently, this result provides direct evidence for the connection between Early Slavic and North-Western European maternal gene pools^73^ (Supplementary Fig. 58).

The female individual DR34 from the barrow of the Kremenye archaeological site (Moscow region) (12th century CE), belongs to the V1a1 mitochondrial haplogroup^43^. Comparative analysis of the mtDNA reveals a difference at one nucleotide position between DR34 and the Viking individual VK144 with a known ancestry from the Danish Viking clan, which was found in a mass grave of Viking warriors in Oxford dating to 1002^102^. Currently, mtDNA DR34 shows the closest affinity to two individuals from modern-day Belgorod and Pskov in Russia. All three genomes share a unique single nucleotide variant, A7299G, which is absent in other sequenced individuals. Overall, this data provides a case of a maternal genetic link between some ancient Rus' and present-day Russian populations^43^ (Suppementary Fig. 59).

In total, we revealed twenty three complete mtDNA sequence matches between present-day humans and ancient Rus’ samples (Supplementary Fig. 58). All the haplotypes, with the exception of one (H1j3), have been identified among contemporary inhabitants of Northern Europe, as well as in present-day Slavic groups (Supplementary Fig. 58). It should be noted that the analysis of sequences similar to those found in ancient Rus’ also indicated that the maternal lineages identified in our sample of ancient Rus’ are predominantly distributed in populations of East and Northeast Europe, are rare in West Europe, and are virtually absent in East Asia, with the exception of mitochondrial lineages that are widely distributed across the Eurasia (Supplementary Table 4, Supplementary Fig. 60).

#### S2.7.2. Y chromosome haplogroups diversity

Y-chromosome haplogroups were successfully determined for 69 male samples (Supplementary Table 4). These haplogroups represent seven major paternal lineages common in Western Eurasia and include N1a, R1a, R1b, I1a, I2a, J2a, and E1b. The two main Rus’ genetic clusters exhibited divergent Y-chromosome profiles: R1a was the most frequent (63%) in the Ancient Major Rus' cluster, contrasting with the dominance of N1a (81%) in the Ancient North Rus' genetic cluster (Supplementary Fig. 61).

##### S2.7.2.1 Ancient Major Rus’ genetic cluster

Within the **Ancient Major Rus'** genetic cluster, six different Y-chromosome lineages were identified: R1a, N1a, I2a, I1a E1b, and R1b (Supplementary Table 4). The predominant haplogroup found in the Ancient Major Rus' genetic cluster was R1a-M420, accounting for 63% of individuals, followed by E1b and I2a, each at 11%. The prevalence of haplogroups N1a and R1b was 6%, whereas I1a constituted about 3% (Supplementary Fig. 61).

#### R1a

Within the **Ancient Major Rus'** genetic cluster, haplogroup **R1a-M420 (R1a)** was identified in 22 male individuals. All of them belong to haplogroups within two sub-branches: **R1a-Z282** (R1a1a1b1a) and **R1a-Z94** (R1a1a1b2a) (see Supplementary Table 4), Supplementary Fig. 55). By far, the most common paternal lineage in the **Ancient Major** Rus**'** genetic cluster is **R1a-Z282** (R1a1a1b1a). Lineage **R1a-Z282** shows the highest frequencies in Slavic-speaking groups from Central and Eastern Europe, reaching frequencies of ~50% in present-day East Slavic populations (up to 47% among Russians, up to 44% among Ukrainians, and up to 50% among Belarusians)^125^. The R1a-Z282 haplogroupe is further subdivided into four major branches: **R1a-Z280**, **R-PF6155**, **R1a-Y2395**, and **R-Y17491**, and two of them have been identified in samples from ancient Rus' (see Supplementary Fig. 55).

The paternal haplogroup **R1a-Z280**, which is widespread among modern East Slavic populations^125^ was identified in eight Rus’ individuals (see Supplementary Fig. 55). This lineage is well-represented in ancient males from the Volga-Oka region of Central Rus’. The carriers were primarily from the urban populations of Yaroslavl (individuals Ya15, Ya71, Ya79) and Pereslavl (AB206, AB209). A derived subclade, **R1a-L1280**, was found in sample AB37 from the barrow of the Pustosh Popova in the Volga-Oka region. The R1a-L1280 haplogroup is presently common in Poland, Russia, eastern Germany, Serbia, Bosnia and Herzegovina, and Slovakia (<https://www.yfull.com/tree/R-L1280/>; <https://www.familytreedna.com/>). Notably, this subclade has been previously reported in an individual from a medieval Slavic burial site in Śródka, Poland, dating to the Piast dynasty period (1000–1200 CE)^126^. Further downstream branches of R1a-Z280 were also observed. Haplogroup **R-YP1698** was identified in individual AB141 from Lubech, while individual DR25 from the Luzhki burial site (Moscow) belonged to the **R1a-L366** subclade (see Supplementary Fig. 55). The R1a-L366 lineage is predominantly found in modern Eastern European populations, including Russians, Poles, and Bulgarians.

The Slavic R1a-Z280 lineage was found for DB54 from the Northern Rus’ Minino burial site. Our whole genome analysis showed Slavic-related genetic profile of the individual DB54 (Ancient Major Rus' cluster). The subclade **R-CTS8816** is presently found in various Slavic populations, primarily in Russia (FamilyTreeDNA, https://www.familytreedna.com/). This suggests that Y-chromosomal lineage R-CTS8816 was likely introduced into the Northern Rus' area during the Slavic expansion in the 10th–12th centuries CE.

The second most prominent lineage in the **Ancient Major Rus' cluster** was **R1a-PF6155**, identified in eight individuals from urban centers including Yaroslavl and Pereslavl (Supplementary Fig. 55). This haplogroup is prevalent in Central and Eastern Europe, constituting over 20% of paternal lineages in modern Western Slavic and 11–15% in Eastern Slavic populations, with a lesser presence in Balkan groups of Slavic heritage^125^. Several subclades of R1a-PF6155 were identified. The **R-L1029** branch and its derivatives were observed in six individuals from Yaroslavl, Pereslavl, the Nikolskoe site, and the Knyazhya gora burial. Furthermore, individual AB35 (Novinki, Vologda region) was assigned to subclade **R-CTS11962**, which is also widespread in Central and Eastern Europe (https://www.yfull.com/tree/R-CTS11962/). A notable West Slavic branch, **R1a-L260**, was found in two individuals from the Nikolskoe site (DB62 and DB63). Both were further resolved to the derived haplogroup **R1a-YP5253**. This specific subclade has been previously reported in a medieval Slavic-related burial from Ląd, Poland (1000–1200 CE), dated to the Piast dynasty period^126^.

The second sub-branch of R1a found among ancient Rus’ samples, R1a-Z94, is a subbranch of its ancestral haplogroup **R1a-Z93** has been identified in a single individual, DB60, a child who was buried at the Nikolskoye burial site in the Beloozero region of Northern Rus’. (Supplementary Table 4, Supplementary Fig. 55. The R1a-Z93 lineage is associated with the ancient populations of the Western Eurasian steppes^110,125,127^, with subbranches currently prevalent in Central and South Asia, as well as among present-day populations in the Volga-Ural and Caucasus regions^128^. Our results indicate a steppe-associated origin for the paternal lineage of the individual from Nikolskoe (DB60). Notably, he possessed the mtDNA haplogroup H8c, which has already been identified among the Scythians and various steppe populations^110,129^. This combined evidence suggests that the individual's parents likely had nomadic origins or ancestral ties to steppe populations that migrated into the Northern Rus' territory during the formative period of the Rus' state.

#### I2a

The Y-chromosome lineage **I2a-M423** was identified within the **Ancient Major Rus'** genetic cluster in four samples, including the male individual from the rural necropolis of Komarovka (DR32) and individuals AB136 from Lubech and Khreple (AB47, AB48). All of them were classified into the clade I2a-CTS10228 and its downstream branches I2a-Y3120 and I2a-S17250 (Supplementary Fig. 63). The Y-chromosome haplogroup I2a-M423, indicative of Paleolithic European heritage, is prevalent in present-day Southeastern European populations, especially in Bosnia and Herzegovina (about 50%), Croatia, and Serbia (roughly 40%)^130^. The present distribution is predominantly associated with the substantial demographic growth of Slavic-speaking populations during the Migration Period, which significantly impacted the genetic landscape of Eastern and Southeastern Europe^130,131^. As mentioned above, haplogroup I2a-CTS10228 is the most common Y-chromosome DNA haplogroup among present-day South Slavs (over 50% among Croats and Serbians 50)^132^. The I2a-CTS10228 subclade has been recognized in previously published data from medieval Europe, specifically in an individual from the Viking Era in Sweden and another from the Bezdanjača Cave (Croatia) and Viminatium burial site in Serbia (110–1300 CE)^102,107^.

The Y-chromosomal haplogroup I2a and its subclades, found in samples within the **Ancient Major Rus'** genetic cluster provide evidence that individuals with a genetic relationship to South Slavic populations may have been present within the territory of medieval Rus’.

#### E1b

The paternal Y-chromosome lineage E1b-V13 and its derived subclade E1b-Y3183 have been identified in four individuals from the Rus_SWest and Rus_Core genetic subclusters: AB213 from Pereslavl, DB66 from Gnezdilovo, DR24 from Komarovka, and Ya07 from Yaroslavl (Supplementary Table 4). This Y-chromosome lineage is not present in samples of the Ancient North Rus' genetic cluster (Supplementary Fig. 61). The E1b-V13 lineage, a subclade of E1b-M78 has been prevalent among the people of the Balkan Peninsula populations since the IA^133^ and reaches frequencies of approximately 30% in some present-day populations of the region^134^. E1b-V13 (E1b1b1a1b) haplogroup is hypothesized to have migrated from the Middle East and Western Asia to the Balkan region during the early-to-mid BA (~2500–2000 BCE)^134^, suggesting that the ancestors of the examined ancient Rus’ males may have originated from the Balkan region.

#### R1b

Only two individuals (AB205 and DB75) from our samples belong to the R1b haplogroup, which was represented by subclade R1b1a1b1a1(R1b-L52) and its derived branch R1b1a1b1a1a2c1a4b2c1a1b1b1(R1b-Y17690), respectively (Supplementary Fig. 61, Supplementary Table 4). Both samples are from Central Rus’ Volga-Oka region and belong to the Rus_Core genetic subcluster. The Y-chromosome haplogroup R1b is the predominant Y-chromosome lineage in Western Europe, with the highest prevalence in Atlantic European populations^135^. Currently, its derived subclade R1b-L52, which were identified in the AB205 male individual from Pereslavl is common in Northern and Western Europe (according to the Yfull database, <https://www.yfull.com/tree/R-L52/>).

The male individual DB75 from the Gnezdilovo burial site, archaeologically associated with the Rus’ military elite (druzhina), was assigned to Y-chromosome haplogroup R1b-Y17690. This lineage is a subclade of R1b-A2072, which is predominantly found in present-day Northwestern Europe, particularly in the British Isles (Scotland, England) and among individuals from the United States likely of British Isles descent FamilyTreeDNA (<https://discover.familytreedna.com/y-dna/R-A2072/tree>). Notably, its derived haplogroup R1b-Y17690 has, to date, only been identified in present-day Russian populations. The estimated time of its emergence, approximately the 9th century CE (according to *FamilyTreeDNA*), coincides with the formation of the Rus'. This data suggests that this derived R1b-Y17690 branch may have been originated from Northwestern clade in medevial Rus’ and subsequently become a distinctive paternal line among Russians.

#### N1a

In contrast to the Ancient North Rus’ genetic cluster, which is predominantly defined by haplogroup N1a, the Ancient Major Rus' genetic cluster (Rus_Core) includes only two male individuals belonging to Y-chromosome haplogroup N1a (Supplementary Fig. 62). The first, individual Ya39 from Yaroslavl, was assigned to the unique **N1a-BY184755** lineage, which is, to date, only reported in present-day Russian populations (YFull, <https://www.yfull.com/tree/N-BY184755/>; [familytreedna.com/y-dna/N-BY184755](http://familytreedna.com/y-dna/N-BY184755)). The second, individual DR26, which was buried at a rural settlement Komarovka (Kursk region) had the Y-chromosomal haplogroup **N1a-L550** (N1a1a1a1a1), a sub-clade of the main N-VL29 branch, which is common in present-day populations from the the Baltic region and Northern Europe (Estonian, Latvians, Lithuanian, southwestern Finnish)^139,140^. This haplogroup has also been detected in several VKE samples from Sweden and Estonia, as well as in one individual from the Gnezdovo burial site within Rus’ area^102^. Notable, Y-chromosome lineage of the subject from Komarovka positioned upstream the **N1a-VL11**, branch, which includes a presumable (but yet to be proven) member of the Rurik dynasty, namely the son of Prince Alexander Nevsky^136^ (Supplementary Fig. 62). The derived markers downstream of N1a-L550 in the Komarovka person could not be ascertained due to insufficient genome coverage; hence, an unambiguous conclusion about the possible paternal link between putative members of Rurik dynasty and this ancient individual from Komarovka have yet to be made.

#### I1a

One individual from the necropolis of the military elite, located in the Gushino burial site, has been identified as belonging to the haplogroup **I1a** (AB14) (Supplementary Fig. 63). Individual AB14 belongs to the haplogroup **I1a-Y3662**, which is presently distributed throughout Great Britain and Ireland (https://www.yfull.com/arch-4.03/tree/I-Y3662/)

##### S2.7.2.2 Ancient North Rus’ genetic cluster

The results demonstrate a restricted diversity of paternal lineages among people from the **Ancient North Rus'** genetic cluster. Of the 26 male individuals in the **Ancient North Rus'** cluster, 21 were classified as belonging to the Y-chromosome haplogroup N1a-M231. The remaining individuals were distributed among three other Y-chromosome haplogroups: haplogroup I1~ (1 individual), I1a (2 individuals), I2a (1 individual) and R1a (1 individual). (Supplementary Table 4, Supplementary Fig. 61). Haplogroup N1a-M231 is a widespread Y-chromosome clade across Northern Eurasia, frequently observed in present-day Uralic- and Altaic- speaking populations^137^. With the exception of one individual (DR22 from Voezero), all N1a-M231 carriers in our dataset belonged to its N1a-M46 (also known as N-Tat) subclade (Supplementary Table 4, Supplementary Fig. 62).

According to the phylogenetic tree, most of these N1a-M46 samples fall within downstream branches of the clade defined by marker N1a-CTS6967 (N1a1a1a1a) (Supplementary Fig. 62). The earliest known sample of this subclade was a BA individual from Western Siberia (kra001, Russia_Krasnoyarsk_BA)^138^, the genetic profile of which is represented by the ancient Siberian genetic component. Furthermore, all twenty one samples belong to a specific subclade, N1a-Z1936. This clade is now found at high frequency almost exclusively in Uralic-speaking populations, being most prevalent among Finnic-speaking groups around the Baltic Sea. It has high frequencies in Saami (46.1%) and Finnish (42.2%) populations, with substantial frequencies also in adjacent eastern populations such as Karelians (20.1%) and Northern Russians from the Vologda and Arkhangelsk regions (~20%)^139^. Interestingly, an individual from Minino I (2250 BCE), a representative of the Mesolithic hunter-gatherer culture in northeastern Europe^140,141^ also belongs to the same Y-chromosome lineage (Supplementary Fig. 62). The data indicate the continued presence of the N1a-Z1936 subclade with ancient Siberian ancestry in this region from the Mesolithic period to present.

The presence of the N1a-Z1936 lineage in Northern Rus' area likely originated from ancient migrations emanating from Western Siberia and the Volga-Ural regions, with estimates placing its arrival in North-Eastern Europe around 3,500 years ago. It was previously shown that the migration of populations carrying haplogroup N1a can be traced into Fennoscandia and Northwestern Russia, and correlated with the introduction of an ancient Siberian genetic component into the gene pool of northeastern European populations^106,142^. The ancient Siberian component present in the genomes of representatives of the Ancient Northern Rus’ genetic cluster is probably related to these migration events.

A unique paternal lineage was revealed under haplogroup N1a, specifically with terminal marker N1a-FT68246 (N1a1a1a1a2a1a), a subgroup of N1a-VL62. This lineage was prevalent among the medieval inhabitants of the settlements of Beloozero, having been precisely recognized in 14 of 29 male individuals: four from Minino, nine from Nefedyevo, and one individual from the Shuygino burial site (Supplementary Fig. 62). The parental branch, N1a-VL62, is widespread among contemporary Fenno-Ugrian groups in Finland (Finns and Sami) and Estonia. Interestingly, however, that the derived subclad N1a-FT68244 is now exclusive to present-day Russians from the Sverdlovsk, Voronezh, and Moscow regions (https://www.yfull.com/tree/N-VL62, <https://www.yfull.com/tree/N-FT68244>, 2024). Based on an estimated time of appearance around the 5th century CE (<https://www.yfull.com/tree/N-FT68244>, 2024) and a geographical distribution pattern, N1a-FT68246 (N1a1a1a1a2a1a) subclade likely originated and spread among the Beloozero population, forming a distinct Russian-specific branch of the N1a lineage.

It should be mentioned that because of low coverage of some of the examined samples from the Minino, Nefedyevo, and Shuygino burial sites, we were unable to thoroughly identify their Y-haplogroups.  For instance, sample DB34 (Nefedyevo) and sample DB35 (Nefedyevo) were found to belong to common Y-chromosome haplogroup N1a-Z1928 and N1a-Z1925, respectively. We can, however, further conclude with a high degree of certainty that both of these samples belong to the N1a-FT68246 clade based on the findings of the parental lineage kinship study. Similarly, for sample DB69 from the Shuygino, the haplogroup N1a-Z1925 was identified. Given that this individual was a first-degree male relative of DB71 with haplogroup N1a-FT68246, it is quite likely that DB69 also belongs to the N1a-FT68246 haplogroup (see Supplementary Informations S2.8).

In addition, our analysis identified a single carrier (DR22) of the haplogroup N1a2 and, specifically, N1a2-Z35049 (N1a2b2a1a) subbranch in the Northern Rus’ Voezero burial site of the Poonezhye region.  This lineage has diverged from the ancestral clade N1a-F1360 (N1a2), which is primarily distributed among the East Asian and Uralic populations, but the specific N1a-Z35049 haplogroup subbranch is presently exclusive to Russians (Supplementary Fig. 62).

The occurrence of N1a2-Z35049 and N1a-FT68246 subclades in the Ancient North Rus' leads us to infer that these subbranches arose from Fenno-Ugrian (Uralic) speaking populations concurrent with the migration and settlement of East Slavs in the northern lands. Subsequently, these paternal lineages expanded and became integrated exclusively into the gene pool of present-day Russian populations.

The remaining lineages within the Ancient North Rus' genetic cluster exhibited the following frequencies: I1~: 4%, I1a: 7.69 %, I2a: 4%, R1a: 4% (Supplementary Fig. 61[)](https://drive.google.com/file/d/1thnmI_dcCPGC2KE4qwsJ_BMP88Ox0IKT/view?usp=drive_link).

The Y-chromosomes of two males from the Nefedyevo burial site, DB21 and DB22, were classified within haplogroup I1. DB21 belonged to the sub-clade I1a-Y20853 (I1a1b2a1~). Due to the low genomic coverage, only the root haplogroup I1-FGC10273 (I1~) was identified for DB22. Given that these individuals are first-degree relatives—most likely a father and son—they are presumed to belong to the same sub-clade (see Supplementary Informations S2.8). On the phylogenetic tree, the I1a-Y20853 clade is positioned within the I1a lineage of present-day East Slavs and was identified among Russians from the Kirov region (Supplementary Fig. 63).

Notably, both R1a and I1a lineages appeared among the individuals of the Ancient North Rus' cluster during the later stage of the Beloozero community’s existence, specifically in the 12th and 13th centuries CE. This finding implies that during that time a number of new people moved into the northern region of ancient Rus'.

Within the **Ancient North Rus'** genetic cluster, the Y-chromosome haplogroup I2a was found in a single individual, DB53 from Minino. This individual was further characterized as belonging to the **I2a-CTS10228** subclade (Supplementary Fig. 63). This specific subclade is widespread among modern South Slavic populations, reaching its highest frequency—over 50%—in the regions of Dalmatia (Croatia) and Bosnia and Herzegovina^132^.

In the **Ancient North Rus'** cluster, the Y-chromosome haplogroup R1a was identified in only one individual. This individual, DB32 (Nefedyevo), belonged to the subclade R-Y67627. Notably, DB32 is the single male individual from this burial site for whom no familial connections to others have been established (see Supplementary Fig. 55; Supplementary Information S2.8, Fig. 9). Currently, haplogroup R-Y67627 and its derived subclades are found in present-day populations from Russia and Finland.

##### S2.7.2.3 Y-chromosome lineages in individuals from Non-Rus'/non-Slavic groups

*Ancient Mordva burial sites*

Genetic analysis of two individuals from the Pogiblovo (AB38) and Muranka (AB30) burial sites, associated with the ancient Mordva culture, revealed two distinct paternal haplogroups (Supplementary Table 4). The individual AB38 (Pogiblovo) was identified as belonging to the haplogroup R1a-M417 (R1a1a1) Supplementary Fig. 55). Unfortunately, limited genomic coverage prevented the identification of a more derived subclade, hindering the precise determination of the origin of this patrilineage. However, it is worth noting that this macro-haplogroup is among the dominant lineages in the present-day Mordva people with a frequency of approximately 26.5%^143^.

The individual AB30 (Muranka) was assigned to the subclade N1a-Z1934 (N1a1a1a1a2a) of haplogroup N1a-Z1936 (Supplementary Fig. 62). This lineage is widespread among modern Fenno-Ugrian populations of the Volga-Uralic and the Baltic regions, reaching an average frequency of up to 25% in populations east of the Urals and in the Volga-Ural region^139,144^. Notably, its frequency in the modern Mordvin population is significantly lower, at approximately 5%^139,143^.

##### S2.7.2.4 Y-chromosome lineages of outlier samples

#### R1a

Individual AB10, a genetic outlier from the Voronovo, belongs to the **R1a-L1280,** a derived subclade of haplogroup R1a-Z280 (Supplementary Fig. 55). The same haplogroup was identified in individual AB37 from the Pustosh Popova burial site (Kostroma), which is part of the Rus_SWest genetic subcluster. Both individuals are from barrows of Central Rus’ Volga-Oka region. The R1a-L1280 haplogroup is widely spread over present-day individuals from Poland, Russia, East Germany, Serbia, Bosnia & Herzegovina, and Slovakia (https://www.familytreedna.com/).

#### J2a

We found a single sample in our dataset that belonged to Y-chromosome haplogroup J2a1 (J2a-Y12379) (Supplementary Table 4). This is the male AB23 from the Knyazhya Gora cemetery of the Southern Rus' region, a genetic outlier with genetic affinities to Caucasian populations. This J2a1 subclade is one of the most ancient Y-chromosome lineages found in the Caucasian region. It was found in a Mesolithic hunter-gatherer from the Kotias Klde karst grotto in Western Georgia, who lived approximately 10,000 years ago^145^. The J2a-Y12379 clade is predominantly found in the present-day populations of the North and Central Caucasus regions (<https://www.yfull.com/tree/J-Y12379/>. The identification of Y-chromosomal lineage (J2a-Y12379), presumably of Caucasian origin in our sample, suggests potential long-distance migration of individuals in ancient Rus' area.

### S2.8. Ancient Rus’ pedigrees reconstruction

#### S2.8.1. Small pedigree identification based on READv2, IBD results, sex, age and archaeological data

Numerous first-, second-, and third-degree relatives were found among tested ancient Rus’ individuals, predominantly in the Northern Rus’ region, by using READv2 software (Supplementary Table 30). We used these data in combination with IBD analysis results along with mtDNA and Y-chromosome haplogroup data to reconstruct ancient Rus’ pedigrees (Supplementary Table 31, Supplementary Table 4). The reconstruction also included data on the individuals' sex and age.

In total, based on READv2 estimation, we found 19 first-degree relative pairs in our cohort (Supplementary Table 30, Supplementary Fig. 64). Among them, we identified seven parent-offspring pairs and 2 sibling pairs, all within Northern Rus’ region. We combined READv2 results with data on individual sex and age, which resulted in reconstruction of small pedigrees or family trios among the individuals from Beloozero region (Nefedyevo, Minino, Shuygion), as well as among individuals from Poonezhye region (Voezero). Additionally, one presumably 1st degree relative pair was found in two St. George’s (Yuriev) Monastery sarcophagi of Novgorod (DR38 and DR4).

##### 2.8.1.1 First-degree relatedness

**DB21** and **DB22 (Nefedyevo)**: son and father

Parent-offspring relatedness has been established by READv2. DB21 is a child estimated to be 4-5 years old, and DB22 is an adult 45-50 years old. Both samples belong to the male and share a common Y-chromosome haplogroups and mtDNA sequences. Therefore, we can assume that DB22 was the father of DB21.

**DB5, DB6** and **DB25 (Nefedyevo)**: father, mother and son

DB5-DB25 and DB6-DB25 pairs were shown to have parent-offspring first-degree relatedness, whereas DB5 and DB6 are unrelated. It was found that DB5 and DB25 shared the common Y-chromosome haplogroup, and that DB6 and DB25 had identical mtDNA sequences. This evidence suggests that DB5 and DB6 are patents of DB25.

**DB25** and **DB27 (Nefedyevo)**: father and son

Parent-offspring first-degree relatedness was identified among DB25 and DB27 Both of them are adult males and have the same Y-chromosome haplogroup and different mtDNA sequences; therefore, they must be father and son. Since we have shown that the father of DB25 is DB5, we assume DB27 is a son of DB25.

**DB4** and **DB27 (Nefedyevo)**: mother and son

First-degree relatedness was identified between DB4 and DB27. Presumably, DB4 and DB27 have identical mtDNA sequences. However, due to the low coverage of mtDNA DB4, we were unable to confirm this. We can propose that DB4 is the mother of DB27.

Although DB27 is a son of DB25, the results show that DB4 and DB25 are not genetically related, yet their burials are located in close proximity, and they are also the probable parents of DB27. Consequently, it can be hypothesized that DB4, DB25, and DB27 form a familial unit where DB4 and DB25 are parents of DB27.

**DB27** and **DB28 (Nefedyevo)**: father and son

First-degree relatedness was identified among DB27 and DB28. Both of them are adult males and have the same Y-chromosome haplogroup and different mtDNA sequences; therefore, they must be a father and a son. Since we have shown that the father of DB27 is DB25, we assume DB28 is a son of DB27.

**DB35** and **DB36 (Nefedyevo)**: son and father

Parent-offspring first-degree relatedness was identified using READv2. Both individuals are males. DB36 is an adult (40-45 years old), and DB35 is a young man (18-20 years old). They share an Y-chromosome haplogroup and have different mitochondrial genomes. Although the kinship between DB35 and DB36 was predicted to be parent/offspring, it was impossible to identify which of them is a father and which is a son.

**DB46, DB43** and **DB50 (Minino)**: presumably mother and daughters or sisters

First-degree relationship of the DB43-DB50 and DB46-DB50 pairs was identified using READv2. From their ages and shared mtDNA haplotype, we infer that they might be sisters or a mother (DB46, over 50) and two daughters (DB43 and DB50, 4-5 and 7 years old). It should be mentioned that the precise relationships between these individuals are difficult to ascertain due to a wide range of their archaeological dates based on archaeological context.

**DB47** and **DB49 (Minino)**: father and son

First-degree relationship was identified using READv2. The assumption that these individuals are father and son is based on the fact that their Y chromosomes belong to the same haplogroup, with mitochondrial sequences differing. The discrepancy in age further substantiates this hypothesis: DB47 is over 50 years old, while DB49 is between 15 and 18 years old.

**DB59** and **DB76 (Nikolskoe)**: sisters

Siblings first-degree relatedness was identified using READv2. Both individuals are female children (1 year ± 4 months and 1-2 years old) and have identical mitogenomes, therefore they could be sisters.

**DB62** and **DB63 (Nikolskoe)**: brothers

Siblings first-degree relatedness was identified using READv2. Both individuals are males, 3-5 and 40-50 years old. They share mtDNA haplotype and Y-chromosome lineage, therefore they could be brothers.

**DB71** and **DB69**, **DB72 (Shuygino)**: father, son and daughter

Parent-offspring first-degree relatedness of DB71-DB69 and DB71-DB72 pairs and siblings first-degree relatedness of DB69-DB72 pair was identified using READv2. DB69 and DB72 are children with the same mtDNA haplotype. DB71 is a 45-55 year old male with a different mitochondrial genome, he must be their father.

**DR19** and **DR22 (Voezero)**: brother and sister

Siblings first-degree relatedness was identified using READv2. Both male and female individuals are children and share the same mtDNA haplotype, therefore they could be brother and sister.

**DR11, DR15** and **DR17 (Voezero)**: brother and sisters

First-degree relationship of the DR11-DR15 and DR11-DR17 pairs was identified using READv2. All of them are children (from newborn to 10 years old) and share the same mtDNA sequence; therefore, we assume that these three individuals are siblings, namely one brother and two sisters.

**DR12** and **DR13 (Voezero)**: father and daughter

First-degree relationship was identified using READv2. The samples are characterised by different mtDNA haplotypes, different sex and a large age difference (45-55 and 1-2 years old). Archaeological data indicates the burial sites of DR12 and DR13 are located in close proximity. This data allowed us to assume that they could be father and daughter.

**DR4** and **DR38** **(Novgorod):** father and son or brothers

First-degree relatedness was identified by READv2, but with a small number of overlapped SNPs (751 SNPs). The two individuals are advanced in age. Due to the limited coverage of individual DR38, it is not possible to verify the identity of their mitochondrial haplotypes. However, based on single reads that cover some positions where substitutions have occurred in the mitogenome of DR4, it can be deduced that these individuals are not maternally related. Presumably, they could be father and son or brothers.

##### 2.8.1.2 Second-degree relatedness

We identified ten pairs of second-degree relatives in our sample using the READv2 (Supplementary Table 30). We found the majority of second-degree relative pairs in the Nefedyevo sites of Northern Rus’. We identified one pair of second-degree relatives in Komarovka, Southern Rus' region (DR24 and DR32), and another pair in Pereslavl, Central Rus’ Volga-Oka region (AB209 and AB210).

**DB18** and **DB25 (Nefedyevo)**: fraternal nephew and uncle

Preliminary analysis of the READv2 results indicates that DB18 and DB25 are second-degree relatives. The presence of distinct mitochondrial genomes suggests that they are likely to be paternal relatives, as evidenced by their shared Y-chromosome haplogroup. Both individuals are elderly males. Given DB18's relationship to DB25, as well as his parents, DB5 and DB6, it can be deduced that DB18 is a fraternal nephew of DB25.

**DB5, DB6,** and **DB18** **(Nefedyevo)**: paternal grandparents and grandson

DB6 and DB18 have divergent mitochondrial sequences, and DB5 and DB18 share a Y-chromosome haplogroup, indicating paternal relatedness. The coincidence of the mtDNA haplotypes of DB5 and DB18 can be explained by the prevalence of the lineage in the population or by the distant relationship of the parents of DB18 (as confirmed by the ROН profile). Given that DB18 is also a second-degree relative of the son of DB5 and DB6 of DB25, its position in the pedigree is paternal grandson.

**DB5, DB6,** and **DB27 (Nefedyevo)**: paternal grandparents and grandson

DB27 is the son of DB25, who is the son of DB5 and DB6. Consequently, DB27 is the grandson of DB5 and DB6, a conclusion that is substantiated by the calculated READv2 second-degree kinship.

**DB25** and **DB28** **(Nefedyevo)**: paternal grandfather and grandson

DB28 is the son of DB27, who is in turn the son of DB25. Consequently, DB27 is the grandson of DB25, a fact that is corroborated by the calculated READv2 second-degree relationship.

**DB33** and **DB34** **(Nefedyevo)**

DB33 and DB34 are second-degree relatives, based on the READ data, with different mitochondrial haplogroups. This indicates that their relationship is not maternal. Both are adult males aged 30-35 years and 40-60 years, and they have the same Y-chromosome haplogroup. It is possible that they are half-siblings through their father, either as uncle and nephew or as grandfather and grandchild.

**DB34** and **DB36** **(Nefedyevo)**: paternal grandfather and grandson

DB34 and DB36 are second-degree relatives, based on the READ data. They have different mitochondrial haplogroups, indicating that their relationship is not through the maternal line. However, they share the same Y-chromosome haplogroup, which suggests a potential paternal relationship. This could imply a relationship such as uncle and nephew or grandfather and grandson. According to archaeological evidence, DB34 was buried earlier than DB63. Therefore, it is likely that DB34 is the grandfather and DB36 is the grandson.

**DB23** and **DB5** **(Nefedyevo)**: fraternal uncle and nephew

The DB23 individual is a second-degree relative of DB5, based on the results of the READ and IBD analyses. Since DB23's archaeological evidence suggests an earlier date than DB5's, and they share the same Y-chromosome haplogroup, it can be inferred that DB23 is either DB5's uncle or grandparent.

**DB11** and **DB43** **(Minino)**

The two subjects are both female, with DB11 being a 25-35 years old and DB43 a seven-year-old child.We conclude that they are not maternally related due to the differences in their mitochondrial genomes. They could be paternal aunt and niece. However, due to the low genome coverage of DB43, we were unable to confirm this.

**DB11** and **DB51 (Minino)**

We have established that individuals of different sexes share a single mitochondrial haplotype, indicating that they are maternal relatives. The precise morphological sex and age of DB51 remain unknown; however, genetic analysis has determined that it is male. Based on the age of DB11, the mtDNA data, and the results of IBD fragments analysis (second degree of relatedness) it is probable that DB51 is a maternal nephew of DB11.

**DB49** and **DB51 (Minino)**: grandson and grandfather or fraternal nephew and uncle

Both subjects are male. DB49 is a 15-18 year-old adolescent. The IBD results revealed a second-degree relatedness between the two individuals. Since their mitochondrial genomes are different, but they share the same Y-chromosome haplogroup, it suggests a paternal relationship between them. Therefore, they might be uncle and nephew or grandfather and grandson.

**DR24 and DR32 (Komarovka):** maternal half-brothers or uncle and sororal nephew

DR24 and DR32 exhibit an identical complete sequence of mitochondrial DNA, belonging to the mtDNA haplogroup H1a1. This finding suggests a relationship by maternal lineage. Both individuals were buried at an age exceeding 50 years and possess different Y-chromosome haplogroups. Consequently, it is plausible that they were maternal half-brothers, or alternatively, that one of them was an uncle and the other a sororal nephew.

**AB209** and **AB210 (Pereslavl)**: grandson and grandfather or fraternal nephew and uncle

AB209 and AB210, both from Pereslavl, exhibit different mtDNA haplogroups and a shared Y-chromosome haplogroup R1a1a1b. Further analysis was conducted on AB209, while insufficient coverage precluded analysis of AB210. The AB210 remains were attributed to an elderly individual, while AB209 was linked to a child. Consequently, it can be hypothesized that AB210 is the paternal grandfather of AB209, although the possibility of AB210 being an uncle cannot be discounted.

##### 2.8.1.3 Third-degree relatedness

Finally, three third-degree relative pairs were identified: one was from Nefedyevo (DB17 and DB6), one was from Shuygino (DB69 and DB73) of the Northern Rus’ (Beloozero) region, and one pair was from the Gnezdilovo site of the Central Rus' Volga-Oka region (DB66 and DB67).

#### S2.8.2. Large pedigrees reconstruction

We applied the obtained IBD fragment analysis results (Supplementary Fig. 52, 53) for validation of the first- and second-degree relationships identified by READv2 prior to reconstruction of genealogical connections between studied individuals and representing them as a pedigree. More distant relationship connections (third- and fourth-degree) were identified solely based on the IBDP results using the reference points obtained from the simulated IBD distribution. The analysis of shared IBD fragments revealed additional biological relatedness up to the 6th degree between individuals within and between the reconstructed pedigrees from the dataset under consideration. This data was used to construct the five pedigrees, designated further as Minino, Nefedyevo, Shuygino, and Nikolskoe (Fig. 9, Supplementary Fig. 64) pedigrees. Familial ties between individuals from neighbouring settlements in the Beloozero region were also identified. In total, there was at least a 75% of tested individuals from the Northern Rus' were genetically connected to one another.

***Nefedyevo pedigree***

Based on READv2 relatedness tests, we identified a four-generation pedigree in the Nefedyevo site, along with two parent-offspring pairs (Supplementary Fig. 64). Archaeological evidence suggests that aged woman DB2 could be a principal female ancestor of individuals in the Nefedyevo site and all the pedigree, representing the earliest known burial at this location. This is also confirmed by data from isotopic analysis, which revealed that the woman DB2 was a "first-generation" migrant resettled to Nefedyevo I in an adulthood^22^. We could not identify the male principal ancestor ("founder") of the Nefedyevo family group; nevertheless, it can be postulated that he was a carrier of the Y-chromosome haplogroup N-FT68246, as most tested males from both early and late burials shared this lineage (Fig. 9). Due to the low quality of her genomic data, only the mtDNA genome for this sample was reconstructed, and a complete match of the maternal lineage J1c1b1a5a was identified with an aged man, DB31, buried not far from DB2 (Supplementary Fig. 1). The close relationship of these two individuals (DB2 and DB31) was suggested by their anthropological features, since both of them had the same rare bone pathology - a narrowing of the frontal suture of the skull^23^. Interestingly, DB31 had the highest sum of ROHs among all tested Northern Rus’ individuals, indicating close relatedness of his parents. Based on IBD analysis results, DB31 had a fourth- to fifth-degree degree relationship with the **DB5** and **DB18** individuals and a more distant relationship with at least eight Nefedyevo individuals (DB36, DB29, DB49, DB21, DB23, **DB25**, **DB27**, and individual DB11 from Minino (Supplementary Table 31), including individuals of reconstructed small pedigrees (Supplementary Fig. 63). The earliest representative of this family was **DB23**, a presumed uncle or grandfather of female **DB5**. According to long shared IBD fragments, **DB18** had fourth-degree relatedness with **DB27** (Komarovka) and was more distantly related to DB23, DB31, DB49 (burial 46, Minino II), and DB36 (Supplementary Table 31). Therefore, the small pedigree from the male DB5 was supplemented with DB18 and DB23. Next, a pair of the second-degree relatives, DB33 and DB34, were also included in the pedigree, as DB34 was the second-degree relative (most likely grandfather) of DB36, who's connected with DB18 and other members of his family with fourth- to sixth-degree relationships.

Despite the low coverage and the number of SNPs (~210 000), DB6, a female from a family pair DB5-DB6, had numerous long (up to 61 cM) IBD fragments with individual DB17 from Nefedyevo and 1-2 fragments of > 32 cM common with DB21, DB22, and DB36. Thus, all three small families from Nefedyevo form one large pedigree (Fig. 9).

Next, several fourth- to sixth- degree relatedness pairs were identified for DB24, DB29, and DB20 with other individuals from Nefedyevo (Supplementary Table 31). In addition, DB19 and DB30 share a common haplotype of the K1a1b5 mitochondrial haplogroup identical to Nefedyevo family members DB27 and DB35.

Notably, all but two males from the Nefedyevo family belong to the same Y-chromosome lineage, N1a1 (N-FT68246). Only DB21 and DB22 had the Y-chromosome of the I2 haplogroup, which can indicate the arrival of new men to the Beloozero region at the turn of the 12th century, who were assimilated by the local family group.

In total, in the Nefedyevo I site we have reconstructed family ties across seven generations of a single genetic lineage. Twenty-one out of the twenty-six tested individuals have been found to be genetically related (Fig. 9).

***Minino pedigree***

DB41 and DB46 are among the earliest individuals identified in the Minino pedigree, both are female and date from the late 11th century to the 12th century CE (Fig. 9a). Female DB46 is a member of a familial group that includes DB50 and DB43, all three of whom are presumed to be first-degree relatives. Additionally, READv2 analysis indicates that DB43 has a second-degree relatedness with DB11 (see Supplementary Note S2.8.1). Unfortunately, the genome coverage for all three first-degree female samples is low, making them unsuitable for IBD testing. Nonetheless, IBD analysis suggests that DB11, presumably the niece of DB43, also exhibits second-degree relatedness with DB51 and DB48 (Supplementary Table 31). Both of these burials are associated with later archaeological contexts than DB11. Considering mitochondrial haplogroup data, it is plausible that DB48 and DB51 could be the niece and nephew of DB11, respectively. However, it cannot be excluded that they might also be the grandchildren of DB11. Further, DB11 was found to be a third-degree relative of male DB13, who shared a mitochondrial sequence with numerous individuals not only from the Minino but also from the Nefedyevo burial site (Supplementary Table 31). Overall, these seven individuals represent one Minino four-generation family tree related to their earliest DB41 female. Significantly, DB51, the member of this family, had DB49 as a second-degree relative, who, in turn, was the son of the DB47. Relatedness of DB51 with DB49, DB13, and DB48 from Minino was confirmed by numerous shared long IBD fragments (>32 cM) (Supplementary Table 28), and two individuals, DB12 and DB42, shared the same maternal mitochondrial lineages as some members of the Minino family (U8a1a1b1 and K1a1b5). In total, at least twelve tested individuals from the Minino burial site were members of one family group (Fig. 9), and all tested male members shared an Y-chromosome lineage N1a (N-FT68246) as Nefedyevo pedigree members do.

Remarkably, the second earliest individual from the Minino burial site, DB41, exhibited no close familial relationships but demonstrated long shared IBD fragments (>32 cM) with samples DB47 and DB40, both from Minino burials, which suggests a distant relatedness (Supplementary Table 31).

DB41, an elderly woman interred in the second quarter of the 11th century, the earliest tested individual in our Minino cohort, she could potentially be linked to relatives from all four burial sites in the Beloozero region, namely Minino, Nefedyevo, Shuygino, and Nikolskoe. In addition to her connections to Minino samples (DB49, DB40), she appears to have a distant relationship with several individuals from Nefedyevo (DB20, DB21, DB22, DB29, DB36, and DB22), Shuygino (DB69, DB72), and even Nikolskoe (DB61) Supplementary Table 31).

Sixteen out of eighteen male individuals from Nefedyevo and Minino pedigrees, as well as both tested male individuals from the Shuygino, belong to the same branch of the Y-chromosome haplogroup N1a1 characterized by terminal markers N1a1a1a1a2a1a1a (N-FT68246). This data indicates a strong patrilineal pattern among the pedigrees from all three burial sites. However, it is noteworthy that six individuals carrying the N1a haplogroup exhibit low genomic coverage, limiting the resolution of their terminal branches.

In contrast to the common paternal Y-chromosome lineage observed among individuals from the Minino, Nefedyevo, and Shuygino sites, genetic heterogeneity in maternal mitochondrial lineages was identified among members of these pedigrees. Although various mtDNA haplogroups were detected in the tested samples, several of these haplogroups (U5a2a1b, H1a, K1a1b5, U8a1a1b1, and H11a1) were shared by individuals from different Northern Rus’ burial sites. This finding further substantiates the relationships among individuals from burials with mixed Slavic and Finnish cultural contexts within the Beloozero region.

***Shuygino pedigree***

One small family was identified among tested individuals from the Shuygino burial site (Northern Rus’, Beloozero region), which combined the father (DB71), his son (DB69), and his daughter (DB72). Two females (DB73 and DB74) are maternal lineage relatives of DB69 and DB72 and share common mtDNA haplotype H1+16189+152. Another identical mtDNA lineage of haplogroup U5a2a1b was revealed between female DB70 from Shuygino and individuals from Minino burials, DB11 and DB51. Long shared IBD fragments (>32 cM) were detected between Shuygino individuals and several samples from Nefedyevo, as well as single individuals from Minino and Nikolskoe (Supplementary Table 31, Supplementary Table 28). Therefore, we can assume that all tested individuals from Shuygino represent one family group, related to the Nefedyevo family and some other individuals from the Belloozero region. This corresponds to the archaeological evidence from these burial sites^23^.

***Nikolskoe pedigree***

Two pairs of first-degree siblings (DB62 and DB63, and DB59 and DB76) have been identified in Nikolskoe (Fig. 9). All of them share a common mtDNA haplotype, H18. DB76 was also found to be a fourth-degree relative of DB55 from the same site, and distant relatedness could be proposed for DB62 with DB55 and DB36, as they shared IBD fragments of > 25 cM (Supplementary Table 28).

DB61 from Nikolskoe was a distant relative of DB41 from the Minino site and also had relatedness with at least two individuals from Nefedyevo (Supplementary Table 31). Notably, DB61, in contrast to all other Nikolskoe site samples, belongs to the Rus_Middle group and most likely represents a direct descendant of intermarriage between Finnish- and Slavic-related individuals.

Based on shared IBD fragments longer than 25cM, long-distance relatedness with individuals from other Northern Rus’ burial sites (Nefedyevo and Minino) could also be proposed for DB62 and DB76 from Nikolskoe (Supplementary Table 31).

***Voezero pedigree***

We found three groups of first-degree relatives within the Northern Rus’ burial site Voezero (Supplementary Fig. 65). A first-degree relationship was identified between pairs of individuals DR11 and DR15, and DR11 and DR17. But the evidence for kinship between DR15 and DR17 was weak, possibly due to insufficient coverage and overlapping SNPs. Nevertheless, all three were children and exhibited identical mitochondrial genomes (with the exception of a few gaps in DR15 and DR17 mitochondrial sequences); we then considered them as siblings.

Another pair of individuals from the Voezero site, DR12 and DR13, are a man and a child (girl) of first-degree relatedness. They possess different mtDNA haplotypes, with the mitogenome of DR13 being identical to that of another individual from the Voezero sample, DR16.

Siblings DR19 and DR22 are brother and sister, respectively (Supplementary Table 30, 31, Supplementary Fig. 65). Intriguingly, both of them exhibit the highest amount of ROH when compared to the other representatives of the Voezero group (Fig. 9c); therefore, they were siblings of closely related parents, and both died in early childhood.

Based on IBD analysis, all three small families had individuals with long shared IBD fragments (Supplementary Table 31, Supplementary Table 28), assuming that all tested individuals from the Voezero burial site are relatives.

Our kinship analysis of five burial sites in medieval Northern Rus' revealed that 66% of the individuals under study were genetically related. We identified 42 related individuals across four Beloozero region burial sites and 9 relatives at Voezero (Arkhangelsk region). Notably, burial grounds of Nefedyevo and Minino II showed strong familial clustering. A particularly illustrative example is a four-generation family plot in Nefedyevo's southern sector (Fig. 9c, Supplementary Fig. 1). Similarly, the Nikolskoe burial grounds yielded evidence of the interment of individuals with close familial ties within the same barrows. These genetically confirmed patterns demonstrate kinship as a primary organizer of medieval Northern Rus' necropolises, offering new insights into their social structure.

Despite the substantial sample size from Central Rus’, only a minimal number of kinship pairs were identified. Among them, for example, an old male and a juvenile, identified genetically as second-degree relatives, were interred together in a combined grave in the ancient Rus' town of Pereslavl. In Komarovka, a rural settlement in ancient Rus', it was discovered one pair of relatives shared mitochondrial DNA, but they had distinct Y-chromosome haplogroups, confirming maternal kinship. Finally, in the Gnezdilovo burial site, which includes a number of military elite (Druzhina) graves, a pair of third-degree relatives were identified as probably having a common paternal ancestor.

Thus, within the ancient Rus' area, our results show regional differences in demographic group formation. While urban and rural areas in central Rus’ were more diverse, settlements in Northern Rus’ were made up of tiny, closely connected groups. Perhaps the patrilineal kinship at the Gnezdilovo burial site, which is connected to the military elite, reflects the tradition of the formation of such military settlements.

### S2.9. Appearance prediction of ancient Rus' people

#### S2.9.1. Phenotypes prediction based on genomic data

In Russian bylinas (oral epic poems) and folk tales, the ancient Rus' heroes are predominantly depicted as tall bogatyrs (epic knights) with light hair and blue or bright eyes. Female characters are described as fair-skinned and possessing long, light-brown braids^146^. Notably, this folkloric of ancient Rus' people aligns remarkably well with descriptions found in medieval chronicles. Indeed, numerous medieval chroniclers—such as Byzantine sources like Leo the Deacon (10th century CE), Western European accounts like those of Adam of Bremen (11th century CE), and Arab writers — described the inhabitants of ancient Rus' (East Slavs) as tall, fair-skinned, light-haired (blond or redheaded), and light-eyed with blue eyes being the most frequently mentioned^147–149^. Certain Arab authors have observed geographical variations among Slavic types; for instance, the Arab geographer al-Idrisi (12th century CE) remarked that southern Slavs possess a darker complexion and hair^150^.

According to the chronicles, the northern and northeastern peripheries of ancient Rus’were inhabited by several non-Slavic peoples, which were referred to in the *Primary Chronicle* as the Chud', Merya, and Ves'. In Russian folklore narratives (Legends), as well as in Norwegian folklore traditions (referred to as Saga), the Chud’ people - an ancient population inhabiting northwestern Russia and the Baltic-Finnic region - are frequently mentioned^151^. In Russian folkloric sources, the Chud’ are often described as "Chud’ white-eyed"^152^. This distinctive characteristic finds a parallel in Norwegian sagas, where the Chud’ people (likely referring to the same or a related people group) are also noted for their "white eyes"^153^. Such a description may indicate unusually light iris pigmentation, possibly reflecting a unique phenotypic characteristic in this group of people.

We examined the G allele frequency of the genetic marker rs12913832 within the HERC2 gene in our sampling, which is associated with blue eyes^154^. We genotyped 50 samples for this marker using whole genomic sequencing data, along with target sequencing 24 HIrisPlex markers (n=10) and pyrosequencing (n=23), the assay that we developed for highly degraded DNA (see Materials and Methods) (Supplementary Table 32). Taken together, we estimated that the frequency of allele G was 0.807 in the Ancient Rus’ population (Supplementary Table 33). We analyzed frequencies of the G allele according to the geographical areas in Ancient Rus’ and revealed that the frequencies in Northern Rus' (0.826) and Central Rus' Volga-Oka region (0.9), approximately equal to the set of Viking Era samples^102^ and slightly less than among medieval Finns (0.93)^155^, but are comparable for those in present-day Russians from Yaroslavl (0.83), and Arkhangelsk (0.79) as well as for present-day population from Poland (0.82)^156,157^. We also noted a slight decrease of G allele frequency in Southern Rus' region (0.75), but due to the small sampling this result is not clear.

Next, using HIrisPlex-S web tool, we obtained prediction probabilities for 59 individuals by merging genotype data for markers of HIrisPlex-S system obtained from whole genome sequencing, target sequencing, and pyrosequencing (Supplementary Table 34, Supplementary Fig. 66). According to our results, Ancient Rus’ inhabitants mostly had blue eyes with maximum percentages in the Northern Rus' and Central Rus' Volga-Oka regions (73% and 77%, respectively). Blond, dark blond, and light brown were likely the most common hair colors among the ancient Rus' population. Red- and black-haired individuals were singular as in Northern Rus’ region, as in Southern Rus’ region. Noteworthy that in the HIrisPlex-S system “dark blond” and “light brown” overlap in colors, and is similar to the Russian term “rusyj”^158^

The skin color of all studied Rus’ individuals ranged from pale to intermediate (Supplementary Table 34, Supplementary Table 35 Supplementary Fig. 66).

Consequently, it can be inferred that the genetic phenotyping of our sample aligns with medieval sources that describe the physical appearance of individuals in ancient Rus’.

#### S2.9.2 Graphic craniofacial reconstruction of medieval Rus’ individuals

Based on the skulls of two individuals from the Yaroslavl mass grave from the 13th century CE (individual MS and individual Ya79), graphic craniofacial reconstructions were made in frontal and lateral projections (Supplementary Fig. 67, 68).

Eye, hair and skin pigmentation were determined based on genomic data (see Supplementary Fig. 66, Supplementary Table 34).

For individual MS, the estimated probability of having blue eyes is 0.911, while the hair color is blond (p=0.68). The estimated probability of having intermediate colour skin is 0.521 and pale skin is 0.425) (Supplementary Fig. 69).

For individual Ya79, the estimated probability of having brown eyes is 0.912, and the highest p-value is 0.800 for red hair colour. The probabilities of having pale skin is 0.688. Based on these results, a phenotype of blue eyes, red hair, and pale skin was predicted for individual Ya79 (see Supplementary Fig. 70).

7.	*Archaeology of the Northern Rus’ village of the 10th–13th centuries: mediaeval settlements and burial grounds on Lake Kubenskoye. V. I - Mediaeval settlements and burial grounds (Археология севернорусской деревни X–XIII веков: средневековые поселения и могильники на Кубенском озере. Т. I - средневековые поселения и могильники*. (Nauka (Наука), Moscow, 2007).

8.	Franklin S. *Writing, Society, and Culture in Early Rus, c. 950-1300*. (Cambridge University Press, Cambridge, 2002).

9.	Nestor. *The Russian Primary Chronicle: Laurentian Text*. (Mediaeval Academy of America, Cambridge, 1953).

10.	Lihachev, D. S. *Tale of Bygone Years (Повесть временных лет)*. vol. 1 (Publishing House of the USSR Academy of Sciences (Изд-во Академии наук СССР), Moscow, Saint Petersburg, 1997).

11.	Lukin, P. V. East Slavic ‘Tribes’ and their Princes: Constructing History in Ancient Rus’ (Восточнославянские «племена» и их князья: конструирование истории в древней Руси). in 83–89 (Indrik (Индрик), Moscow, 2010).

12.	Ryabinin E.A. *Finno-Ugric tribes as part of Ancient Rus: On the history of Slavic-Finnic ethnocultural relations (Финно-угорские племена в составе Древней Руси: К истории славяно-финских этнокультурных связей)*. (Publishing House of Saint Petersburg University (Publishing House of St. Petersburg University), Saint Petersburg, 1997).

13.	Leontiev, A.E. *The Archaelogy of the Merya: The early histiry of North-Eastern Russia (Towards the Prehistory of North-Eastern Rus’)*. (Moscow, 1996).

14.	Makarov N.A. *Colonization of the northern outskirts of Ancient Rus in the 11th–13th centuries. Based on materials from archaeological sites on the Belozerye and Poonezhye portages (Колонизация северных окраин Древней Руси в XI–XIII вв. По материалам археологических памятников на волоках Белозерья и Поонежья)*. (Research Center ‘Scriptorium’ (Научный центр ‘Скрипториум’), Moscow, 1997).

15.	Eremeev I.I. *Antiquities of the Polotsk land in the historical study of the East Baltic region: essays on medieval archeology and history of the Pskov-Belarusian Dvina (Древности Полоцкой земли в историческом изучении Восточно-Балтийского региона: (очерки средневековой археологии и истории Псковско-Белорусского Подвинья)*. (Dmitry Bulanin (Дмитрий Буланин), Saint Petersburg, 2015).

16.	Sedov, V. V. *The Slavs in Roman Times: The 13th International Congress of Slavists (Славяне в Римское Время: XIII Международный Съезд Славистов)*. (IA RAS (ИA РАН), Moscow, 2003).

17.	Lindstedt, J. S. & Salmela, E. Migrations and language shifts as components of the Slavic spread. in *New Perspectives on the Early Slavs and the Rise of Slavic Contact and Migrations* 275–300 (Universitätsverlag winter Heidelberg, Heidelberg, 2019).

18.	Jansson, I. Skandinavien, Baltikum och Rus’ under vikingatiden. in *Det 22. nordiske historikermøte. Rapport I: Norden og Baltikum* 5–25 (Oslo, Oslo, 1994).

19.	Lind J. H. Problems of Ethnicity in the Interpretation of Written Sources on Early Rus’. in *The Slavicization of the Russian North. Mechanisms and Chronology* 246–258 (Helsinki University Press, Helsinki, 2006).

20.	Nosov, E. N. Novgorod land: Northern Priilmenye and Povolkhovye (Новгородская земля: Северное Приильменье и Поволховье). in *Rus’ in the IX-X centuries: an archaeological panorama (Русь в IX-X вв.: археологическая панорама)* 92–121 (Antiquities of the North (Древности Севера), Moscow, Vologda, 2012).

21.	*Rus’ in the IX–X centuries: an archaeological panorama (Русь в IX–X веках: археологическая панорама)*. (Antiquities of the North (Древности Севера), Moscow, Vologda, 2012).

22.	Dobrovolskaya, M. V., Makarov, N. A. & Kiseleva, D. V. Experience in Estimating Individual Mobility of People from the Cemeteries of the Medieval Rus Population Using Data of Isotopic Analysis of Tooth Enamel (Опыт оценки индивидуальной мобильности людей из могильников Северо-Восточной Руси по данным изотопного анализа эмали зубов). *KSIA КСИА* 433–452 (2024) doi:10.25681/IARAS.0130-2620.277.433-451.

23.	Makarov, N. A., Zakharov, S. D. & Buzhilova, A. P. *Medieval settlement on Lake Beloe (Средневековое расселение на Белом озере)*. (Languages of Russian culture (Языки Русской культуры), Moscow, 2001).

24.	Makarov N.A. *Population of the Russian North in the 11th–13th centuries. Based on materials from burial grounds in the eastern Prionezhye (Население русского Севера в XI–XIII вв. По материалам могильников восточного Прионежья)*. (Moscow, 1990).

25.	Makarov N.A. *Colonization of the northern perifery of Ancient Rus in the 11th–13th centuries. Based on materials from archaeological sites on the Belozerye and Poonezhye portages (Колонизация северных окраин Древней Руси в XI–XIII вв. По материалам археологических памятников на волоках Белозерья и Поонежья)*. (Research Center ‘Scriptorium’ (Научный центр ‘Скрипториум’), Moscow, 1997).

26.	Nikitin, A. V. *Report on the excavations of the Vologda expedition of the burial mounds near the village of Novinki in 1969 (Отчет о раскопках Вологодской экспедиции курганов у д. Новинки в 1969 г.)*. (1969).

27.	Nikitin, A. V. *Archaeological report on the excavations at the village Krestzy (Vologda region, Ustyuzhensky District) and the mounds at the village Ploskoe (Novgorod region) (Археологический отчет о раскопках у д. Крестцы (Вологодская область Устюженский район) и курганов у д. Плоское (Новгородская область)*. (1965).

28.	Makarov, N. A., Krasnikova, A. M. & Ugulava, N. D. The first results of the excavations Gnezdilovo Burial ground near Suzdal (Первые результаты раскопок могильника Гнездилово под Суздалем). *Archaeol. Vladimir-Suzdal Land Археология Владимиро-Суздальской Земли* 7–20 (2021).

29.	Makarov, N. A. & Krasnikova, A. M. Suzdal elite: burial with weapons and equestrian equipment in the Gnezdilovo burial ground (Суздальская знать: Погребение с оружием и всадническим снаряжением в могильнике Гнездилово). *Russ. Archaeol. Российская Археология* **4**, 110–120 (2022).

30.	Syrovatko, A. S., Kleshchenko, E. A. & Svirkina, N. G. Lluzhki E - cremations and inhumations at one burial ground (Лужки Е – кремации и ингумации в одном могильнике). in *Archeology of the Moscow region: Materials of a scientific seminar (Археология Подмосковья: Материалы научного семинара)* vol. 17 98–109 (IA RAS (ИA РАН), Moscow, 2021).

31.	Nedoshvina N.G. Akatov burial mound of the XI-XII centuries in the Moscow region. (Акатовский курганный могильник XI-XII вв. в Подмосковье). in *Expedition of State Historical Museum (Экспедиции Государственного Исторического музея)* 214–227 (Moscow, 1969).

32.	Nefedov F.D. *Excavation of mounds in Kasimov district (Раскопки курганов в Касимовском уезде)*. (printing house of M.N. Lavrov (тип. М.Н. Лаврова), Moscow, 1878).

33.	Bogdanov A.P. *Prehistoric Tverites on kurgan excavations (Доисторические тверитяне по курганным раскопкам)*. (M.N. Lavrov and Co., Moscow, 1879).

34.	Kelsiev A.I. *Excavations carried out in Yaroslavl and Tver Provinces in the summer of 1878 by A.I. Kelsiev. (Раскопки, произведенные в Ярославской и Тверской губернии летом 1878 года А.И. Кельсиевым)*. (Publishin hous of M.N. Lavrov (тип. М. Н. Лаврова), Moscow, 1878).

35.	Ushakov Ya. A. *Excavations of mounds in the Uglich district, carried out in 1878 by Ya.A. Ushakov (Раскопки курганов в Угличском уезде, произведенные в 1878 году Я.А. Ушаковым)*. (Publishing house of M.N. Lavrov (тип. М.Н. Лаврова), Moscow, 1878).

36.	Zimina, M. P. *et al.* *Architectural and cultural heritage of Russia: Vladimir region (Архитектурно-культурное наследие России: Владимирская область)*. vol. 415 (1995).

37.	Bastamov, L. N. About the excavations he carried out in the Tver Province (О раскопках, произведенных им в Тверской губернии). in *Minutes of meetings of the Anthropological Department of the Imperial Society of Lovers of Natural History, Anthropology and Ethnography, affiliated with Moscow University (Протоколы заседаний Антропологического отдела Императорского Общества любителей естествознания, антропологии и этнографии, состоящего при Московском университете)* vol. 69 54–56 (Society of Lovers of Natural History, Anthropology and Ethnography (Общество любителей естествознания, антропологии и этнографии), Moscow, 1886).

38.	Kashkin. A.V. *Tver region (Тверская область)*. vol. 128 (Institute of Archeology RAS (Институт Археологии РАН), 2003).

39.	Nefedov, F. D. Excavations of mounds in Kostroma region, carried out in the summer of 1895 and 1896 (Раскопки курганов в Костромской губернии, произведенные летом 1895 и 1896 год). in *Materials on the archeology of the eastern provinces of Russia, collected and published by the Imperial Moscow Archaeological Society with the highest bestowed funds (Материалы по археологии восточных губерний России, собранные и изданные Императорским Московским Археологическим Обществом на высочайшие дарованные средства)* vol. III 161–234 (Publishing house of M.G. Volchaninova (тип. М. Г. Волчаниновой), Moscow, 1899).

40.	Gorodtsov, V.A. in *Archaeological research in Kolomna and Kashira districts (Археологические исследования Коломенского и Каширского районов)* 166 (Moscow, 1928).

41.	Rybakov B.A. About the excavations of Vyatichian mounds in Myakinino and Kremenye in 1927 (О раскопках вятичских курганов в Мякинине и Кременье в 1927 году). in *Collection Scientific Archaeological Society of the 1st Moscow State University (Сборник Научного археологического общества при 1-м Московском государственном университете)* (MSU (МГУ), Moscow, 1928).

42.	Rosenfeldt, R.L. *Report of the exploration team of the Moscow expedition of the IA of the USSR Academy of Sciences on conducting a survey of the state of archaeological sites in the Moscow region in 1976*. (1976).

50.	Buzhilova A.P. *Archaeology of ancient Yaroslavl. Riddles and discoveries (based on materials of the Yaroslavl expedition of the Institute of Archaeology of the Russian Academy of Sciences) (Археология древнего Ярославля. Загадки и открытия (по материалам Ярославской экспедиции ИА РАН))*. (Moscow, 2012).

51.	Zeifer, V. A., Mazurok, O. I., Leontiev, A. E., Stolyarova, E. K. & Saprykina, I. A. Issues of mansion development in Pereslavl – Zalessky (Pereslavl) in the XIII–XV centuries (based on research made during 2016) (К вопросу об усадебной застройке Переславля–Залесского (Переяславля) XIII–XV вв. (по материалам исследований 2016 года)). in vol. 15 132–149 (IA RAS (ИA РАН), Moscow, 2019).

52.	Rasskazova, A.V., Zeifer, V.A., Mazurok, O.I. Medieval mass burial in Pereslavl-Zalessky (Массовое средневековое захоронение в Переславле-Залесском). *Vestn. Arkheologii Antropol. Etnografii Вестник Археологии Антропологии И Этнографии* **4**, 138–150 (2021).

53.	Engovatova A.V., Antipina E.E., Vlasov D.V., Dobrovolskaya M.V., Karpukhin, A.A., Osipov D.O. The ninth collective burial in 1238 on the territory of the Rubleny city in Yaroslavl: Results of a comprehensive study (Девятое коллективное захоронение 1238 г. на территории Рубленого города в Ярославле (результаты комплексного исследования). in 185–208 (2012).

54.	Engovatova, A.V., Rasskazova, A.V., Zeifer, V.A. Dating and Interpretation of the Mass Sanitary Burial in Pereslavl-Zalessky. New AMS Dating Data. (Вопросы датировки и интерпретации массового санитарного захоронения в городе Переславле-Залесском. Новые данные AMS-датирования). *Archeology of the Moscow region (Археология Подмосковья )* vol. 20 408–416 (2024).

55.	*Old Ryazan: a major urban center on international trade routes (Старая Рязань: крупный городской центр на международных торговых путях)*. (2020).

56.	Loshina Yu.V., Sterligova I.A. A forgotten monument of ancient Russian cloisonné: small icon from the Old Ryazan collection of A.V. Selivanov (Забытый памятник древнерусской перегородчатой эмали: ‘образок’ из старорязанской коллекции А.В. Селиванова). *Bull. Russ. Mediev. Art Dep. Вестник Сектора Древнерусского Искусства* 12–22 (2023).

57.	Darkevich, V.P. Excavations of V. A. Gorodtsov in Old Ryazan (Раскопки В. А. Городцова в Старой Рязани). in *Problems of studying ancient cultures of Eurasia (Проблемы изучения древних культур Евразии)* 140–163 (Nauka (Наука), Moscow, 1991).

58.	Darkevich, V.P., Borisevich, G.V. *The ancient capital of the Ryazan land XI-XIII centuries (Древняя столица Рязанской земли XI-XIII вв)*. (Krug (Круг), Moscow, 1995).

64.	Spitsyn A.A. *Excavations in the Smolensk Province for 1892 (Раскопки в Смоленской губернии)*. 56–59 (1894).

65.	Eremenko P.M. Mounds excavated by P.M. Eremenko in Surazh District (Курганы, раскопанные П.М. Еременком в Суражском уезде). *Zap. Imperatorskogo Rus. Arkheologicheskogo Obshchestva Записки Императорского Русского Археологического Общества* **8**, 84–102 (1896).

66.	Bogdanov A.P. *Godichnoe zasedanie Imperatorskogo Obshchestva lyubiteley estestvoznaniya, antropologii i etnografii (Годичное заседание Императорского общества любителей естествознания, антропологии и этнографии)*. 31–32 (1867).

67.	Tyshkevich, K.P. About the excavations of the mounds of the Minsk province. Proceedings of the Anthropological Department. Book 2 (О раскопках курганов Минской губернии. Труды Антропологического отдела. Книга 2). *Proc. Imp. Soc. Lovers Nat. Hist. Anthropol. Ethnogr. Известия Императорского Общества Любителей Естествознания Антропологии И Этнографии* **20**, 30–31 (1876).

68.	Gorbachev, K.A. Protocol of excavations of burial mounds of Seletskaya volost in 1886 (Протокол раскопок курганных погребений Селецкой волости в 1886). *Proc. Soc. Lovers Nat. Hist. Anthropol. Ethnogr. Известия Общества Любителей Естествознания Антропологии И Этнографии* **XLIX**, 724. (1890).

69.	Pezhemsky, D.V. Burial mounds Selco in the upper Western Dvina according to paleoanthropology. (Могильник Сельцо в верховьях Западной Двины по данным палеоантропологии). *Bull. Tver State Univ. Вестник Тверского Государственного Университета* **№4**, 93–108 (2014).

70.	Stepanova Yu.V. Burial mounds Selco in the upper Western Dvina according to archeology (Могильник Сельцо в верховьях Западной Двины по данным археологии). *Bull. Tver State Univ. Вестник Тверского Государственного Университета* **4**, 155–164 (2014).

71.	Sedov V.V. Excavation of mounds on the Upper Podvinye (Раскопки курганов на Верхнем Подвинье). *KSIA КСИА* **129**, 54–60 (1972).

72.	Birkina N.A. *Report on archaeological excavations of the Forest-Steppe archaeological expedition of the Historical Museum at the complex of monuments of the Komarovka settlement and settlement in the Korenevsky district of the Kursk region in 2019 (Отчет об археологических раскопках Лесостепной археологической экспедиции Исторического музея на комплексе памятников городище и селище Комаровка в Кореневском районе Курской области в 2019 г. )*. (2019).

73.	Andreeva, T. V. *et al.* An Individual of the Volintsevo period from Kurilovka: The first Archaeogenetic data (Индивид волынцевского времени из Куриловки: первые археогенетические данные). *Russ. Archaeol. Российская Археология* **3**, 57–71 (2023).

74.	Alikhova A.E. Moiseevsky Settlement in the period of Kievan Rus’ (Моисеевское городище в период Киевской Руси). *KSIA КСИА* **87**, 82–87 (1962).

75.	Samokvasov, D. Ya. *Mogilnye drevnosti Severyanskoy Chernigovshchiny (Могильные древности Северянской Черниговщины)*. (Synodal Printing House (Синодальная типография), Moscow, 1916).

76.	Samokvasov, D. Ya. Mogily Russkoy zemli. Opisaniya arkheologicheskikh raskopok i sobraniya drevnostey professora D. Ya. Samokvasova (Могилы Русской земли. Описания археологическиъх раскопокъ и собрания древностей профессора Д.Я. Самоквасова). *Tr. Mosk. Kom. Po Ustroystvu Chernigovskogo Arkheologicheskogo Sezda Труды Московского Комитета По Устройству Черниговского Археологического Съезда* **3**, 188–192 (1908).

77.	Bogdanov A.P. Kurgan inhabitants of the Severnyanskaya land, according to excavations in the Chernigov province (Курганные жители Северянской земли, по раскопкам в Черниговской губернии). in *From the protocols of the anthropological exhibition (Из протоколов антропологической выставки)* 1–11 (Moscow, 1879).

78.	Belyashevsky, N.F. Excavations on Knyazhya gora in 1891 (Раскопки на Княжей горе в 1891 году). *Readings at the Historical Society of Nestor the Chronicler (Чтения в историческом обществе Нестора летописца)* vol. 6 21–23 (1892).

79.	Rybakov B.A. Excavations in Lubech in 1957. (Chernigov Region) (Раскопки в Любече в 1957 году. (Черниговская область)). *KSIIMK КСИИМК* **79**, 27–34 (1960).

80.	Rybakov, B.A. *Report on the work of the Chernigov expedition in 1958 in the Lubech and Chernigov. (Отчет о работе Черниговской экспедиции за 1958 год в пг. Любече и Чернигове)*. (1958).

81.	Rybakov, B.A. *Report of the Chernihiv expedition of the USSR IA for 1960 (excavations in Lyubech (Отчет Черниговской экспедиции ИА СССР за 1960 года (раскопки в Любече))*. (1960).

82.	Bogdanov, A.P. Delivery of kurgan skulls to the Committee by T.V. Kibalchich in Kiev, its excavations. (Доставление Комитету Т.В. Кибальчичем в Киев курганных черепов, его раскопки). *Izv. Imperatorskogo Obshchestva Lyubiteley Estestvozn. Antropol. Etnografii Известия Императорского Общества Любителей Естествознания Антропологии И Этнографии* **XXXI**, 97–98 (1879).

83.	Karger, M.K. *Ancient Kiev: Essays on the history of material culture of the ancient Rus’ city (Древний Киев: Очерки по истории материальной культуры древнерусского города)*. (1958).

84.	Spitsyn, A. A. Antiquities of the North-Western region: The Lucin burial ground (Древности Северо-Западного края: Люцинский могильник ). *Mater. Archeol. Russ. Publ. Imp. Archaeol. Comm. Материалы По Археологии России Известия Императорской Археологической Комиссии* **1**, 36 (1883).

85.	Zelentsova O.V. *Report on the archaeological security investigations of the Verbovsky (Podbolotyevsky) burial ground and settlement in the Murom District of the Vladimir Region in the area of the bridge crossing over the river Oka with a bypass of the Murom town (Stage II) in 2012. (Отчет о проведении охранных археологических исследованиях Вербовского (Подболотьевского) могильника и селища в Муромском районе Владимирской области в зоне строительства мостового перехода через р. Оку с обходом г. Муром (II этап) 2012 г.)*. (2013).

86.	Zelentsova O.V. *Report on the archaeological excavations of the Podbolotyevsky burial ground in Murom District of the Vladimir Region in 2014*. (2014).

87.	Zelentsova O.V. *Report on the scientific archaeological research of the Verbovsky (Podbolotyevsky) burial ground and settlement in the Murom District of the Vladimir Region in 2013. (Зеленцова О. В. Отчет о проведении научных археологических исследований Вербовского (Подболотьевского) могильника и селища в Муромском районе Владимирской области в 2013)*. (2014).

88.	Goryunova E. I. Pogiblovsky burial ground (based on collections stored in the State Museum) (Погибловский могильник (по коллекциям, хранящимся в Государственном Государственном Музее)). *Archaeological collection (Археологический сборник)* 88–111 (1948).

89.	Alikhova A.E. Muransky burial ground and settlement. Materials and research on the archeology of the USSR. (Муромский могильник и селище. Материалы и исследования по археологии СССР). *Research of the Kuibyshev archaeological expedition (Труды Куйбышевской археологической экспедиции)* 259–301 (1954).

90.	Polivanov V.N. *Muransky burial ground: an archaeological essay (Муранский могильник: археологический очерк)*. (Murakhovskaya Printing House (Типография О. В. Мураховского), Simbirsk, 1893).

98.	Engovatova, A., Zaiseva, G. & Cherkinsky, A. Dating of the defeay of Yaroslavl according to radiocarbon dating (Датировка разгрома Ярославля по данным радиоуглеродного датирования). in *Radiocarbon in archaeology and paleoecology: past, present, and future. Materials of the international conference (Радиоуглерод в археологии и палеоэкологии: прошлое, настоящее, будущее. Материалы международной конференции, посвященной 80-летию старшего научного сотрудника ИИМК РАН, кандидата химических наук Ганны Ивановны Зайцевой)* 114–118 (Samara State University of Social Sciences and Education, 2020). doi:10.31600/978-5-91867-213-6-114-118.

99.	Sedov V.V. The Beginning of Cities in Rus’ (Начало городов на Руси). *Proceedings of the Symposium Dedicated to the 1500th Anniversary of Kyiv (Материалы симпозиума, посвященного 1500-летию Киева)* 51–54 (1983).

100.	Nosov, E. N. The Ancient Rus’ city in Russian historical thought (Древнерусский город в отечественной исторической мысли). in *Cultural heritage of the Russia (Культурное наследие Российского государства)* ИПК «Вести» (IPC ‘Vesti’ (ИПК «Вести»), Saint Petersburg, 2000).

101.	Nosov, E. N. The problem of the origin of the first cities of Northern Rus’ (Проблема происхождения первых городов Северной Руси). in *Antiquities of the North-West of Rus’: Slavic-Finno-Ugric interaction, Rus’ cities of the Baltic (Древности Северо-Запада России: славяно-финно-угорское взаимодействие, русские города Балтики)* 59–78 (St. Petersburg Oriental Studies (Петербургское Востоковедение), Saint Petersburg, 1993).

102.	Margaryan, A. *et al.* Population genomics of the Viking world. *Nature* **585**, 390–396 (2020).

103.	Eremeev I.I. *Normans and the funeral rite of the Ilmen-Volkhov region (Норманы и погребальный обряд Ильмень-Волховского региона)*. vol. 2 (Saint Petersburg, 2023).

104.	Duczko W. *Viking Rus: Studies on the Presence of Scandinavians in Eastern Europe*. (Brill, Leiden, Boston, 2004).

105.	Jansson, I. Early Contacts between Scandinavia and the Orient. in *Untersuchungen zu Handel und Verkehr der vor- und fruhgeschichtlichen Zeit in Mittle- und Nordeuropa* 818 (Vandenhoeck & Ruprecht Gmbh & Co, Göttingen, 1987).

146.	Afanasyev A.N. *Folk Russian fairy tales. The complete edition in one volume (Народные русские сказки. Полное издание в одном томе)*. (Alpha-Kniga (Альфа-Книга), Moscow, 2010).

147.	Leo the Deacon. *The History (История)*. (Nauka (Наука), Moscow, 1988).

148.	Rakhno, K.Yu. The image of the Kievan Rus’ population in Byzantine sources (Внешность населения Руси в Византийских источниках). *Rusin Русин* 10–23 (2020) doi:10.17223/18572685/61/2.

149.	*Ancient Rus’ in Light of Foreign Sources: Anthology. Volume III: Eastern Sources (Древняя Русь в свете зарубежных источников: Хрестоматия. Том. III: Восточные источники)*. (Russian Foundation for the Promotion of Education and Science (Русский Фонд Содействия Образованию и Науке), Moscow, 2009).

150.	Garkavi, A. Ya. *The Stories of Muslim writers about Slavs and Russians: (from the middle of the VII century to the end of the X century AD) (Сказания мусульманских писателей о славянах и русских: с половины VII в. до конца X в. по Р. X.)*. (Saint Petersburg, Saint Petersburg, 1870).

151.	Drannikova, N. & Larsen, R. Representations of the Chudes in Norwegian and Russian Folklore. *Acta Borealia* **25**, 58–72 (2008).

152.	Zelenin, D.K. *Great Russian folk sayings as a material for ethnography (Великорусские народные присловья как материал для этнографии )*. (Indrik (Индрик), Moscow, 1994).

### Legends for Supplementary Tables

**Supplementary Table 1.** Archaeological data on newly reported samples

**Supplementary Table 2.** Library informations and sequencing statistics

**Supplementary Table 3.** Summary of samples sequencing statistics and quality control data

**Supplementary Table 4.** Genetic clustering, mitochondrial and Y-haplogroups information

**Supplementary Table 5.** New radiocarbon data.

**Supplementary Table 6.** Previously published samples used in analysis.

**Supplementary Table 7.** Previously published Viking Era samples genetically similar to ancient Rus' individuals

**Supplementary Table 8.** Outgroup f3-statistics of form *f3*(Ancient_Rus',BA/IA/MA;Mbuti). BA - BA group; IA - IA groups; MA - Medieval groups; Ancient_Rus' - Ancient Major Rus' or Ancient North Rus' genetic cluster

**Supplementary Table 9.** Outgroup *f*3-statistics of form *f3*(BA,Ancient_Rus';Mbuti). BA - BA group, Ancient_Rus' - Ancient Rus' genetic clusters (Rus_North_1, Rus_North_2, Rus_Core, Rus_Baltic, Rus_SWest).

**Supplementary Table 10.** Outgroup f3-statistics of form f3(Ancient_pops, Ancient_Rus';Mbuti). Ancient_pops - ancient groups spanning periods from the Early Iron Age to the Middle Ages, Ancient_Rus' - Ancient Rus' genetic clusters (Rus_North_1, Rus_North_2, Rus_Core, Rus_Baltic, Rus_SWest). Three chronological periods are indicated: IA - Iron Ages (1000-0 BCE), AA - Antiquity and Great Migration period of Common Era (0-700 CE), MA - Medieval period (700-1500 CE)

**Supplementary Table 11.** Outgroup f3-statistics of form f3(Ancient_pops,Rus';Mbuti). Ancient_pops - ancient groups spanning periods from the Early Iron Age to the Middle Ages, Rus' - Ancient_North_Rus' and Ancient_Major_Rus' genetic clusters. Three chronological periods are indicated: IA - Iron Ages (1000-0 BCE), AA - Antiquity and Great Migration period od Common Era (0-700 AD), MA - Medieval period (700-1500 AD)

**Supplementary Table 12.** Outgroup *f*3-statistics of form *f3*(Ancient_IA,Major_Rus;Mbuti). Ancient_IA - ancient IA groups preceding our Rus’ samples, Major_Rus' - Ancient Major Rus' genetic cluster (Rus_Core, Rus_Baltic, Rus_SWest).

**Supplementary Table 13.** *F*4-statistics of the form *f4(*Ancient_Rus',Mbuti;BA,BA). Ancient_Rus' - Ancient Rus' genetic clusters (Rus_North_1, Rus_North_2, Rus_Core, Rus_Baltic, Rus_SWest, Anc_Mordva), BA - Bronze age groups

**Supplementary Table 14.** *f*4-statistics of the form f4(Rus',Mbuti;VolgaOka_IA,Ancient_pop). Rus' - Major_Rus' or North_Rus' genetic clusters, Ancient_Pops - ancient populations of the first millennium CE.

**Supplementary Table 15.** *f*4-statistics in the form of *f4(*Krasnoyarsk_BA/Greece_Mycenian,Mbuti;Ancient_Rus',Latvia_BA).

**Supplementary Table 16.** *f*3-admixture results for Rus_Middle genetic cluster

**Supplementary Table 17.** *f*3-statistics of the form *f3*(Ancient_Rus’,Modern_pops;Mbuti) for ancient Rus' genetic clusters.

**Supplementary Table 18.** *f*3-statistics of the form *f3*(Ancient_Rus’,Modern_pops;Mbuti) for ancient Rus' genetic clusters. Only transversions were included in the analysis

**Supplementary Table 19.** Outgroup *f*3-statistics of form *f3*(Modern_pops,Ancient_Rus;Mbuti) for individual ancient samples. Transition and transversions were included in the analysis

**Supplementary Table 20.** Outgroup *f*3-statistics of form *f3*(Modern_pops,Ancient_Rus;Mbuti) for individual ancient samples. Only transversions were included in the analysis.

**Supplementary Table 21.** f3-statistics of the form f3(Ancient_pops,Modern_pops;Mbuti). Ancient_pops - ancient European groups dated to 1st millennium CE.

**Supplementary Table 22.** *f*4-cladality test of form *f4(*Mbuti, Modern_pops;Ancient_Rus',Target), where Modern_pops are present-day Eurasian and Northern-African populations, and Target is a set of geographically and genetically appropriated present-day Eastern-European populations. Only transversions were included in calculations

**Supplementary Table 23.** Distal qpAdm models with feasible proportions in the [0,1] range and p>0.05 with two or three sources for tested Rus' genetic groups

**Supplementary Table 24.** *qpAdm* models with feasible proportions in the [0,1] range and p>0.05 with two or three BA sources for tested Rus' genetic groups. BA sources: Armenia_EBA_KuraAraxes, Greece_BA_Mycenaean, Italy_Sardinia_EBA, Turkey_Alalakh_MLBA, Turkmenistan_Gonur_BA_1, Uzbekistan_Bustan_BA, Mongolia_LBA_Khovsgol_6, Russia_Krasnoyarsk_BA.SG, Russia_Bolshoy, Russia_Afanasievo, Andronovo, Russia_MLBA_Sintashta, Russia_Shamanka_EBA.SG, Russia_Srubnaya, Voronezh_LBA, Czech_CordedWare, Poland_GlobularAmphora, England_BellBeaker, Russia_Fatyanovo_BA, Sweden_Gotland_PittedWare_BattleAxe, Estonia_MN_CCC, Estonia_BA.SG, Lat.via_BA.Outgroups: Ethiopia_4500BP, Russia_DevilsCave_N, Russia_Karelia_HG, Iran_GanjDareh_N, Russia_Tyumen_HG, Germany_EN_LBK.

**Supplementary Table 25a.** *qpAdm* models with feasible proportions in the [0,1] range and p>0.05 with two or three BA sources for tested Rus' genetic groups. BA sources: Armenia_EBA_KuraAraxes, Greece_BA_Mycenaean, Italy_Sardinia_EBA, Turkey_Alalakh_MLBA, Turkmenistan_Gonur_BA_1, Uzbekistan_Bustan_BA, Mongolia_LBA_Khovsgol_6, Russia_Krasnoyarsk_BA.SG, Russia_Bolshoy, Russia_Afanasievo, Andronovo, Russia_MLBA_Sintashta, Russia_Shamanka_EBA.SG, Russia_Srubnaya, Voronezh_LBA, Czech_CordedWare, Poland_GlobularAmphora, England_BellBeaker, Russia_Fatyanovo_BA, Sweden_Gotland_PittedWare_BattleAxe, Estonia_MN_CCC, Estonia_BA.SG. Outgroups: Ethiopia_4500BP, Russia_DevilsCave_N, Russia_Karelia_HG, Iran_GanjDareh_N, Russia_Tyumen_HG, Germany_EN_LBK, and Latvia_BA.

**Supplementary Table 25b**. *qpAdm* models with feasible proportions in the [0,1] range and p>0.05 with two or three BA sources for tested Rus' genetic groups. BA sources: Armenia_EBA_KuraAraxes, Greece_BA_Mycenaean, Italy_Sardinia_EBA, Turkey_Alalakh_MLBA, Turkmenistan_Gonur_BA_1, Uzbekistan_Bustan_BA, Mongolia_LBA_Khovsgol_6, Russia_Krasnoyarsk_BA.SG, Russia_Bolshoy, Russia_Afanasievo, Andronovo, Russia_MLBA_Sintashta, Russia_Shamanka_EBA.SG, Russia_Srubnaya, Voronezh_LBA, Czech_CordedWare, Poland_GlobularAmphora, England_BellBeaker, Russia_Fatyanovo_BA, Sweden_Gotland_PittedWare_BattleAxe, Estonia_MN_CCC, Latvia_BA. Outgroups: Ethiopia_4500BP, Russia_DevilsCave_N, Russia_Karelia_HG, Iran_GanjDareh_N, Russia_Tyumen_HG, Germany_EN_LBK, and Estonia_BA.SG.

**Supplementary Table 25c.** *qpAdm* models with feasible proportions in the [0,1] range and p>0.05 with two or three BA sources for tested Rus' genetic groups. BA sources: Armenia_EBA_KuraAraxes, Greece_BA_Mycenaean, Italy_Sardinia_EBA, Turkey_Alalakh_MLBA, Turkmenistan_Gonur_BA_1, Uzbekistan_Bustan_BA, Mongolia_LBA_Khovsgol_6, Russia_Krasnoyarsk_BA.SG, Russia_Bolshoy, Russia_Afanasievo, Andronovo, Russia_MLBA_Sintashta, Russia_Shamanka_EBA.SG, Russia_Srubnaya, **Voronezh_LBA**, Czech_CordedWare, Poland_GlobularAmphora, England_BellBeaker, Sweden_Gotland_PittedWare_BattleAxe, Estonia_MN_CCC, PolanLatvia_BA, Estonia_BA.SG. Outgroups: Ethiopia_4500BP, Russia_DevilsCave_N, Russia_Karelia_HG, Iran_GanjDareh_N, Russia_Tyumen_HG, Germany_EN_LBK, and Russia_Fatyanovo_BA.

**Supplementary Table 25d.** *qpAdm* models with feasible proportions in the [0,1] range and p>0.05 with two or three BA sources for tested Rus' genetic groups. BA sources: Armenia_EBA_KuraAraxes, Greece_BA_Mycenaean, Italy_Sardinia_EBA, Turkey_Alalakh_MLBA, Turkmenistan_Gonur_BA_1, Uzbekistan_Bustan_BA, Mongolia_LBA_Khovsgol_6, Russia_Bolshoy, Russia_Afanasievo, Andronovo, Russia_MLBA_Sintashta, Russia_Shamanka_EBA.SG, Russia_Srubnaya, Voronezh_LBA, Czech_CordedWare, Poland_GlobularAmphora, England_BellBeaker, Russia_Fatyanovo_BA, Sweden_Gotland_PittedWare_BattleAxe, Estonia_MN_CCC, Latvia_BA, Estonia_BA.SG. Outgroups: Ethiopia_4500BP, Russia_DevilsCave_N, Russia_Karelia_HG, Iran_GanjDareh_N, Russia_Tyumen_HG, Germany_EN_LBK, and Russia_Krasnoyarsk_BA.SG.

**Supplementary Table 25e.** *qpAdm* models with feasible proportions in the [0,1] range and p>0.05 with two or three BA sources for tested Rus' genetic groups. BA sources: Armenia_EBA_KuraAraxes, Greece_BA_Mycenaean, Italy_Sardinia_EBA, Turkey_Alalakh_MLBA, Turkmenistan_Gonur_BA_1, Uzbekistan_Bustan_BA, Mongolia_LBA_Khovsgol_6, Russia_Krasnoyarsk_BA.SG, Russia_Afanasievo, Andronovo, Russia_MLBA_Sintashta, Russia_Shamanka_EBA.SG, Russia_Srubnaya, Voronezh_LBA, Czech_CordedWare, Poland_GlobularAmphora, England_BellBeaker, Russia_Fatyanovo_BA, Sweden_Gotland_PittedWare_BattleAxe, Estonia_MN_CCC, Latvia_BA, Estonia_BA.SG. Outgroups: Ethiopia_4500BP, Russia_DevilsCave_N, Russia_Karelia_HG, Iran_GanjDareh_N, Russia_Tyumen_HG, Germany_EN_LBK, and Russia_Bolshoy.

**Supplementary Table 25f.** *qpAdm* models with feasible proportions in the [0,1] range and p>0.05 with two or three BA sources for tested Rus' genetic groups. BA sources: Armenia_EBA_KuraAraxes, Italy_Sardinia_EBA, Turkey_Alalakh_MLBA, Turkmenistan_Gonur_BA_1, Uzbekistan_Bustan_BA, Mongolia_LBA_Khovsgol_6, Russia_Krasnoyarsk_BA.SG, Russia_Bolshoy, Russia_Afanasievo, Andronovo, Russia_MLBA_Sintashta, Russia_Shamanka_EBA.SG, Russia_Srubnaya, Voronezh_LBA, Czech_CordedWare, Poland_GlobularAmphora, England_BellBeaker, Russia_Fatyanovo_BA, Sweden_Gotland_PittedWare_BattleAxe, Estonia_MN_CCC, Latvia_BA, Estonia_BA.SG. Outgroups: Ethiopia_4500BP, Russia_DevilsCave_N, Russia_Karelia_HG, Iran_GanjDareh_N, Russia_Tyumen_HG, Germany_EN_LBK, and Greece_BA_Mycenaean.

**Supplementary Table 26**. *qpAdm* models with feasible proportions in the [0,1] range and p>0.05* with two or three selected BA sources for several previously published IA and medieval European groups. See Supplementary Note 2.4 for details. * - if no unrejected model were found, models with p<0.05 are represented.

**Supplementary Table 27.** Statistics of inferred runs of homozygosity (ROHs) longer than 4, 8, 12, and 20 cM.

**Supplementary Table 28.** IBD analysis results for ancient Rus' samples with mean genome coverage >0,25.

**Supplementary Table 29.** IBD analysis results for pairs containing one sample from our Rus' dataset and other previously published 438 European medieval samples/

**Supplementary Table 30.** Pairs of relatives estimated by READv2.

**Supplementary Table 31.** Degrees of relatedness and possible kinships for studied individuals based on their IBDP.

**Supplementary Table 32.** Eye, hair color and skin tone prediction based on DNA markers.

**Supplementary Table 33.** Frequencies for A/G allele rs12913832 in the Ancient Rus' population (samples from this study)

**Supplementary Table 34**. Prediction probabilities of eye color, hair color, and skin tone calculated using the HIrisPlex-S web tool.

**Supplementary Table 35.** Predicted phenotypes for ancient Rus' population (samples from this study) determined by HIrisPlex-S.
